# BTRR complex deficiency is a driver for genomic instability in Bloom syndrome

**DOI:** 10.1101/2025.04.17.649287

**Authors:** Ipek Ilgin Gönenç, Alexander Wolff, Alexandra Viktoria Busley, Angela Wieland, Andrea Tijhuis, Christian Müller, Rene Wardenaar, Loukas Argyriou, Silke Kaulfuß, Markus Räschle, Diana C. J. Spierings, Floris Foijer, Holger Bastians, Andrew N. Blackford, Gökhan Yigit, Arne Zibat, Lukas Cyganek, Bernd Wollnik

## Abstract

Biallelic loss-of-function (LoF) variants in the BTRR complex members *BLM*, *TOP3A, RMI1,* and *RMI2* cause Bloom syndrome (BSyn). The BTRR complex mainly acts on DNA replication and DNA repair processes, and dysfunction of this complex underlies the increased genomic instability and cancer predisposition associated with the BSyn phenotype. Here, we report CRISPR/Cas9-based genome-edited isogenic induced pluripotent stem cell (iPSC) models with compound heterozygous LoF variants in *BLM, TOP3A*, and *RMI1*. The cellular phenotype of all three BTRR-deficient iPSC lines included chromosome segregation defects, increased sister chromatid exchange rates, and impaired DNA single-strand template repair. Using single-cell whole genome sequencing, we showed that BTRR complex deficiency causes increased genome copy number alterations (CNAs) and, therefore, is a driver for genomic instability. CNA load was further induced by applying replication stress, and we observed that BTRR^KO^ iPSCs acquired fewer *de novo* CNA events compared to wild-type cells, suggesting a possible selection loss against cells with high levels of DNA damage. Importantly, stress-induced and non-stress-induced CNAs in single-cell genomes were not stochastically distributed throughout the genome, but instead enriched at fragile sites. This finding might offer an opportunity for the development of novel NGS-based approaches to measure rates of genomic instability in disease conditions.

## Introduction

Genomic instability refers to an increased incidence of aberrations in the genome, for example, as a direct consequence of inefficient DNA damage response (DDR), which is a hallmark of various genetic disorders with cancer predisposition^1^. Among them, Bloom syndrome (BSyn, MIM 210900) is a rare autosomal recessive disorder resulting from the impairment of DNA replication and repair mechanisms^2^. Clinical features of BSyn include primary microcephaly, growth deficiency, photosensitivity, and immunodeficiency^2,3^. Moreover, cancer predisposition is a prominent characteristic of BSyn associated with chromosomal instability and the occurrence of increased somatic variants^2^. Biallelic loss-of-function (LoF) variants in *BLM* have been identified in the majority of BSyn patients^4^. In addition, homozygous or compound heterozygous LoF variants in *TOP3A, RMI1,* and *RMI2* were recently reported in patients presenting a BSyn-like phenotype^5,6^.

The four BSyn-associated genes, *BLM*, *TOP3A*, *RMI1*, and *RMI2*, encode the members of the so-called BTRR complex^7–13^. In vitro, the BTRR complex can resolve potentially deleterious DNA intermediates such as G4 structures, D-loops, and double Holliday junctions (dHJs) forming during the activity of the homologous recombination (HR) pathway which can repair double-strand breaks (DSBs)^14–16^. The Bloom syndrome helicase (BLM), a member of the RecQ helicase family, is responsible for unwinding the DNA helix^17,18^. During HR, BLM activity prompts the migration of the dHJ branches, resulting in a hemicatenated DNA structure. This structure is then resolved by topoisomerase III alpha (TOP3A), a type IA topoisomerase, which is capable of nicking and re-ligating single-stranded DNA^19^. Further, RMI1 and RMI2 are accessory proteins important for the stability and enzymatic activity of the BTRR complex^12,13^. RMI1 binds to three members of the complex and stimulates decatenation activity of TOP3A^20^, whereas RMI2 is required to target BLM to chromatin via FANCM^12,13,21,22^.

The BTRR complex is the only protein complex identified so far that is able to dissolve dHJs without recombination between homologous strands^14,23^. As a result, BTRR complex dysfunction leads to endonuclease-mediated dHJ resolution, which can generate crossover products. The resulting hyper-recombination of chromosomes can be visualised as sister chromatid exchanges (SCEs), and a high SCE rate in metaphase chromosomes is one of the most prominent cellular hallmarks of BSyn^23,24^. Additional sources of genomic instability in BSyn include chromatid breaks, micronuclei, and chromosome segregation defects, resulting from unresolved DNA intermediates and replication stress^25–29^. Recently, recombination hotspots in BLM-deficient cells were shown to be enriched at G4-rich regions of transcribed genes^30^. However, a detailed description of genomic instability at the genome-wide level, specifically in relation to somatic CNAs associated with BTRR complex deficiency, has not yet been documented at single-cell resolution.

In this study, we provide a systematic characterization of the genomic instability profiles associated with BTRR complex deficiency using an isogenic induced pluripotent stem cell (iPSC) model system. We scrutinized cellular BSyn features in terms of chromosomal aberrations in CRISPR/Cas9-based knockout (KO) iPSC lines with biallelic LoF variants in *BLM, TOP3A,* and *RMI1*. First, we showed typical genomic hallmarks of BSyn such as increased SCE rates in the BTRR^KO^ iPSC lines and confirmed impaired homology-directed repair capacity by a novel CRISPR/Cas9-based approach. Next, single-cell whole genome sequencing (scWGS) identified significantly increased CNA formation associated with BTRR complex deficiency in genotype-specific, individual cells. Analysis of common fragile sites (CFS), defined as the late-replicating regions undergoing mitotic DNA synthesis^31,32^, revealed a tendency of CNAs to occur at CFS rather than randomly in the genome of BTRR-complex-deficient iPSCs. Moreover, moderate replication stress elevated both the number of CNA events and CNA/CFS overlaps in all iPSC lines. Taken together, our study provides insights into the genomic instability profile of BTRR complex deficiency at single-cell resolution, and our findings suggest that CNAs as a marker for genomic instability occur at fragile sites of the genome.

## Results

### Cellular hallmarks of BTRR-complex-deficient knockout iPSCs

To investigate the characteristics of genomic instability caused by impairment of BTRR complex members, we generated isogenic CRISPR/Cas9-based KO iPSC models (Fig. 1a). The iPSC model system provided a general platform to study BTRR complex deficiency in pluripotent stem cells. Reported BSyn patient mutations in *BLM*^33^*, TOP3A*^5^, and *RMI1*^5^ were chosen as reference positions for designing the guide RNA sequences to introduce LoF alterations (Supplementary Table 1, Supplementary Fig. 1 a, b). Independent transfections using the same wild-type (WT) iPSC line resulted in three isogenic KO clones (referred to as BLMc.715del2/c.716del_, *TOP3A*_c.2473ins/c.2475del7_, and *RMI1*_c.1228del7/c.1230del_) with biallelic LoF variants in_ the corresponding genes encoding three of the four members of the BTRR complex (Supplementary Fig. 1a, b). *RMI2,* which encodes the fourth member of the BTRR complex, was also targeted with two different guide RNAs, however, the observed genome editing efficiency was too low (<1%) to isolate RMI2 KO clones.

**Fig. 1.**
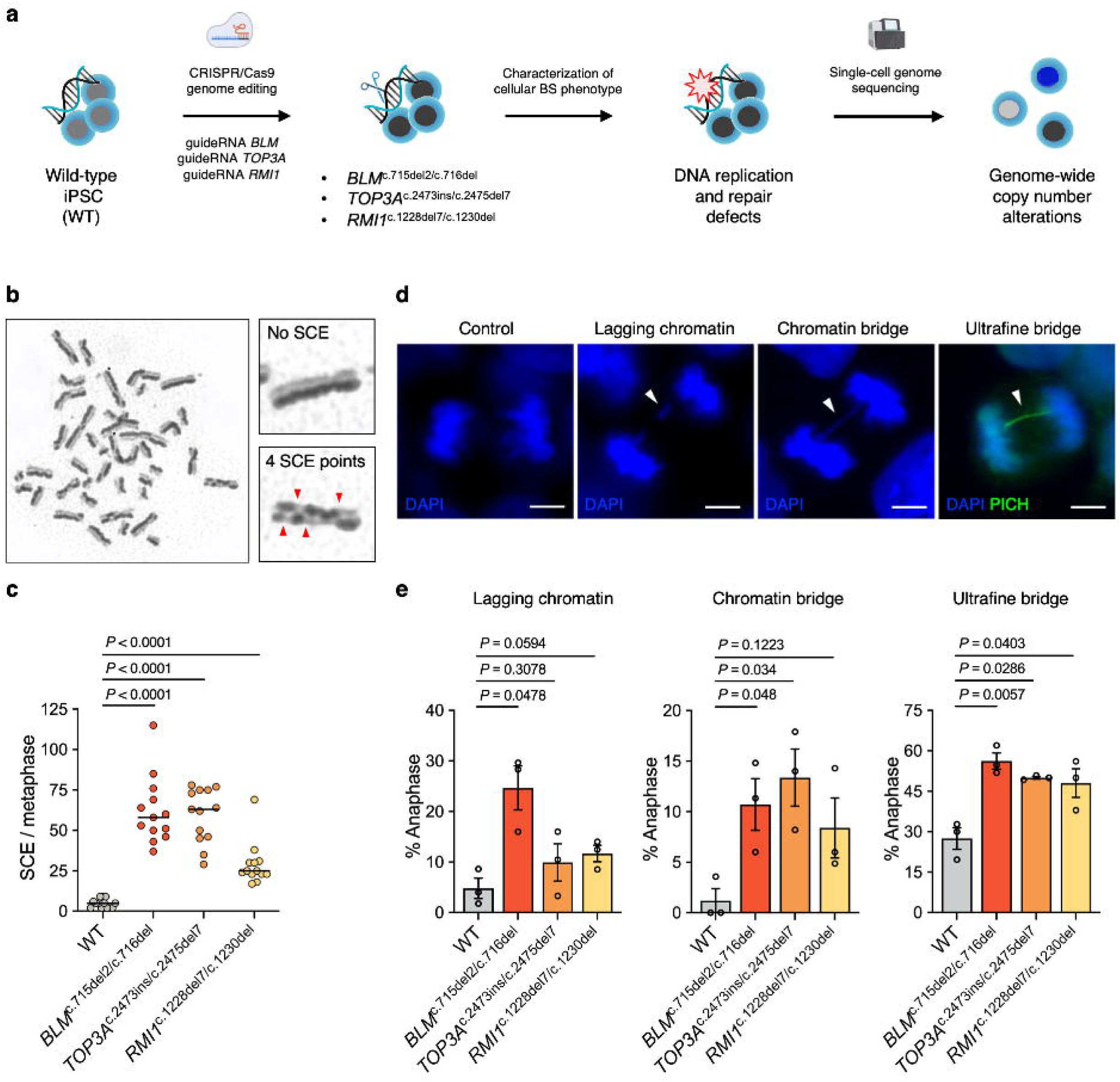
**CRISPR/Cas9-based knockout (KO) iPSCs exhibit cellular characteristics of Bloom syndrome**. **a** Schematic overview of the generation and characterization of BTRR^KO^ iPSCs. **b** BrdU strand-specific labelling of sister chromatids shows chromosomes with no sister chromatid exchange (SCE) or several SCE points (red arrows). The representative metaphase image was from an *RMI1*^c.1228del7/c.1230del^ cell. **c** Quantification of SCE frequencies revealed chromatid hyper-recombination in the BTRR^KO^ iPSCs. The median value was plotted with thirteen metaphase spreads per sample. Two-sided Mann-Whitney test was performed against parental wild type (WT). **d** Representative anaphase images of chromosome segregation abnormalities (white arrows). DNA was stained with DAPI. DAPI-negative ultrafine bridge was visualized with the PICH (ERCC6L) antibody. Scale bar: 2 µm. **e** Quantification of chromosome missegregation events revealed higher proportions of anaphases positive for the denoted events in BTRR^KO^ iPSCs. The assay was repeated three times and at least 50 anaphases per sample were scored. Error bars denote standard error of the mean. Pairwise two-sided t-test with Welch’s correction was performed. Image icons were created with BioRender.

Loss of full-length BLM, TOP3A, and RMI1 protein expression in the generated cell lines was determined by Western blot analysis (Supplementary Fig. 1c). Next, expression of the major pluripotency markers OCT3/4, SOX2, NANOG, SSEA4, and TRA-1-60 assessed by immunofluorescence and flow cytometry analyses confirmed the pluripotent state of all cell lines (Supplementary Fig. 2). In addition, BTRR^KO^ iPSCs did not acquire aneuploidy or gross chromosomal aberrations due to genome editing processes as evaluated by karyotype and arrayCGH analyses (Supplementary Fig. 3).

One of the well-defined roles of the BTRR complex is dHJ dissolution, and its dysfunction results in the resolution of dHJs through the action of endonucleases, which may yield crossover products that can be observed as SCEs^34^. Given that an elevated SCE rate is a hallmark of BSyn^24^, we analysed metaphase chromosomes of the BTRR^KO^ iPSCs. Newly synthesized DNA was labelled by BrdU incorporation and differential staining of metaphase chromosomes revealed highly significant, increased SCE rates in the BTRR^KO^ iPSCs (*P* <0.0001, Fig. 1b, c). SCE rates per chromosome numbers were twelve-fold higher in *_BLM_*c.715del2/c.716del _and *TOP3A*_c.2473ins/c.2475del7_, and six-fold higher in *RMI1*_c.1228del7/c.1230del _cells_ compared to WT cells, indicating that the generated BTRR^KO^ iPSCs represented suitable model systems for BSyn.

Another source of genomic instability is the failure of equal distribution of replicated DNA to daughter cells, which can originate from, e.g., lagging chromatin or DNA bridges^35^. To investigate the effect of BTRR complex deficiency on chromosome segregation, we analysed anaphase cells by immunofluorescence for abnormal events (Fig. 1d). Lagging chromatin can stem from faulty kinetochore attachment, chromatid breaks, or unrepaired DSBs associated with BLM deficiency^36–38^. The analysis of lagging chromatin showed a significantly higher frequency only in *BLM*^c.715del2/c.716del^ cells but not in *TOP3A*^c.2473ins/c.2475del7^ and *RMI1*^c.1228del7/c.1230del^ cells, although the individual values in all three BTRR^KO^ iPSC clones were higher than in WT cells (Fig. 1e). DNA bridges can occur during anaphase in form of (i) chromatin bridges or (ii) ultrafine bridges (UFBs) resulting from unresolved DNA intermediates^39^. Chromatin bridges have nucleosomes and can be imaged with conventional DNA dyes such as DAPI^40^, whereas UFBs are DAPI-negative DNA fibers that can be visualized only by labelling UFB-associated proteins such as BLM or PICH^40,41^. We observed significantly increased numbers of chromatin bridges in *BLM*^c.715del2/c.716del^ and in *TOP3A*^c.2473ins/c.2475del7^ cells (Fig. 1e). To quantify UFBs, we visualized them using a PICH antibody and we observed that the numbers of UFB-positive anaphases were approximately two-fold increased in all three BTRR^KO^ iPSC samples when compared to WT (*P* <0.05, Fig. 1e). The small number of anaphase UFBs in the WT sample confirmed formation of UFBs during cell division in healthy cells, which are subsequently resolved by the BTRR complex and PICH^42^. This result again suggests that lack of one of the four BTRR complex proteins impacts on complex function as seen by the impaired dissolution of UFBs, which is consistent with previous reports^40,42,43^. Taken together, the isogenic CRISPR/Cas9-based BTRR^KO^ iPSC lines presented cellular BSyn characteristics such as chromosome segregation defects, although impairment of individual members of the BTRR complex resulted in subtle variances between genotypes.

### Reduced DNA single-strand template-mediated repair in BTRR^KO^ iPSCs

BLM-deficient cells are generally proficient in homologous recombination^44,45^, but recent evidence suggests BTRR may promote homology-directed repair via the single-stranded template repair (SSTR) pathway^46^. To measure SSTR capacity in BTRR^KO^ iPSCs, we established a novel approach using CRISPR/Cas9 technology. A single-stranded oligonucleotide (ssODN) template featuring a specific *RYR2* variant (*RYR2*:c.7422G>C;p.Arg2474Ser) and the CRISPR/Cas9 complex designed for the variant locus were introduced into BTRR^KO^ iPSCs and the repair outcomes were analysed as a measure of DSB repair capacity at the specific *RYR2* locus by deep amplicon sequencing (Fig. 2a, b). In detail, we used the proportion of single-stranded template-mediated alleles (containing the G>C SNP) versus the NHEJ-mediated alleles (containing indels) to assess DNA repair efficiency. Given that NHEJ is the prominent DSB repair mechanism in G1^47,48^, we assessed cell cycle profiles of asynchronous cells used in the analysis which revealed similar proportions of cells in G1, S, and G2 phases, confirming an unbiased evaluation (Supplementary Fig. 4a, b). Analysis of each iPSC line at a sequencing depth of approximately 24,000x revealed a significantly decreased potential SSTR after Cas9-based DSB induction in the BTRR^KO^ iPSCs (Fig. 2c). In addition, the proportions of indel-containing alleles originating from NHEJ activity were increased in the BTRR^KO^ iPSCs in both transfections with and without ssODN template (Supplementary Fig. 4c, b), suggesting altered DNA repair efficiencies in BTRR complex deficiency.

**Fig. 2.**
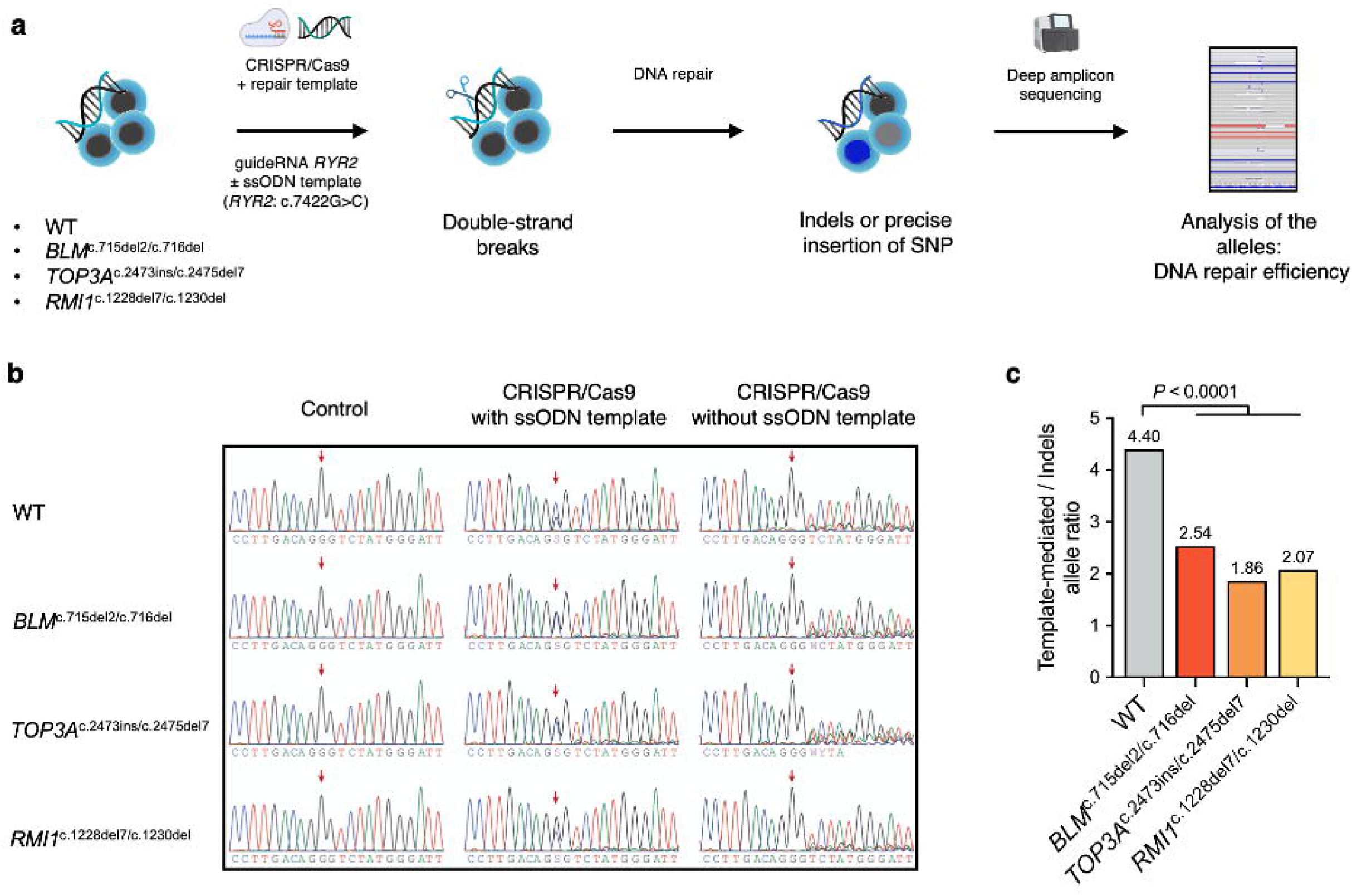
Impairment of the BTRR complex results in altered DSB repair efficiency. **a** Schematic representation of the strategy for determining DNA repair efficiency via a CRISPR/Cas9-based approach. Each cell line was transfected with Cas9 nuclease, *RYR2* guide RNA, and a single-stranded oligodeoxynucleotide (ssODN) containing a single nucleotide polymorphism (SNP) in the *RYR2* gene (NM_001035.3:c.7422G>C). Bulk cells were analyzed via deep amplicon sequencing after transfection and the generated alleles were evaluated with Cas-Analyzer . **b** Chromatograms of the corresponding region in *RYR2* after transfection. Red arrows show the location of the SNP. **c** The evaluation of the alleles (see *Methods*) revealed a decrease in the proportion of template-mediated alleles in the BTRR^KO^ iPSCs. At least 17,000 amplicons were analyzed for each cell line. Kruskal-Wallis test with Dunn’s correction was applied to the ratio values. Image icons were created with BioRender.

### Increased CNAs in single-cell genomes caused by BTRR complex deficiency

Next, we studied the load of genomic alterations as a marker for genome instability on a single-cell basis^49^ using low-coverage (∼0.1X) single-cell whole genome sequencing (scWGS) of the BTRR^KO^ iPSC model system and analysed the genomes from G1 cells with the Aneufinder tool^50^ with stringent quality control parameters. We aimed to use the lowest bin size to detect small CNAs, which was feasible with a 500 kb threshold (Fig. 3a). The CNAs that were identified in more than one WT cell were discarded from the data of BTRR^KO^ iPSCs to exclude inherited variants or so-called clonal CNAs^51^. The proportion of WT cells with one or more identified CNAs was 40.7%. This proportion significantly increased in *BLM*^c.715del2/c.716del^, *TOP3A*^c.2473ins/c.2475del7^, and *RMI1*^c.1228del7/c.1230del^ cell lines (Fig. 3b). Interestingly, *BLM*^c.715del2/c.716del^ and *RMI1*^c.1228del7/c.1230del^ genomes had similarly increased CNA load per cell ratios to the WT cells (Fig. 3c). The identified CNAs were further analysed individually with respect to size and type (i.e. gain and loss), which revealed a wider size distribution of CNAs that were mainly deletions in the BTRR^KO^ iPSCs. Overall, *TOP3A*^c.2473ins/c.2475del7^ and *RMI1*^c.1228del7/c.1230del^ cells had significantly different CNA size distribution compared to WT (Fig. 3d). Taken together, our results indicate that impairment of the BTRR complex is associated with an increase in CNAs in the genome of single cells.

**Fig. 3.**
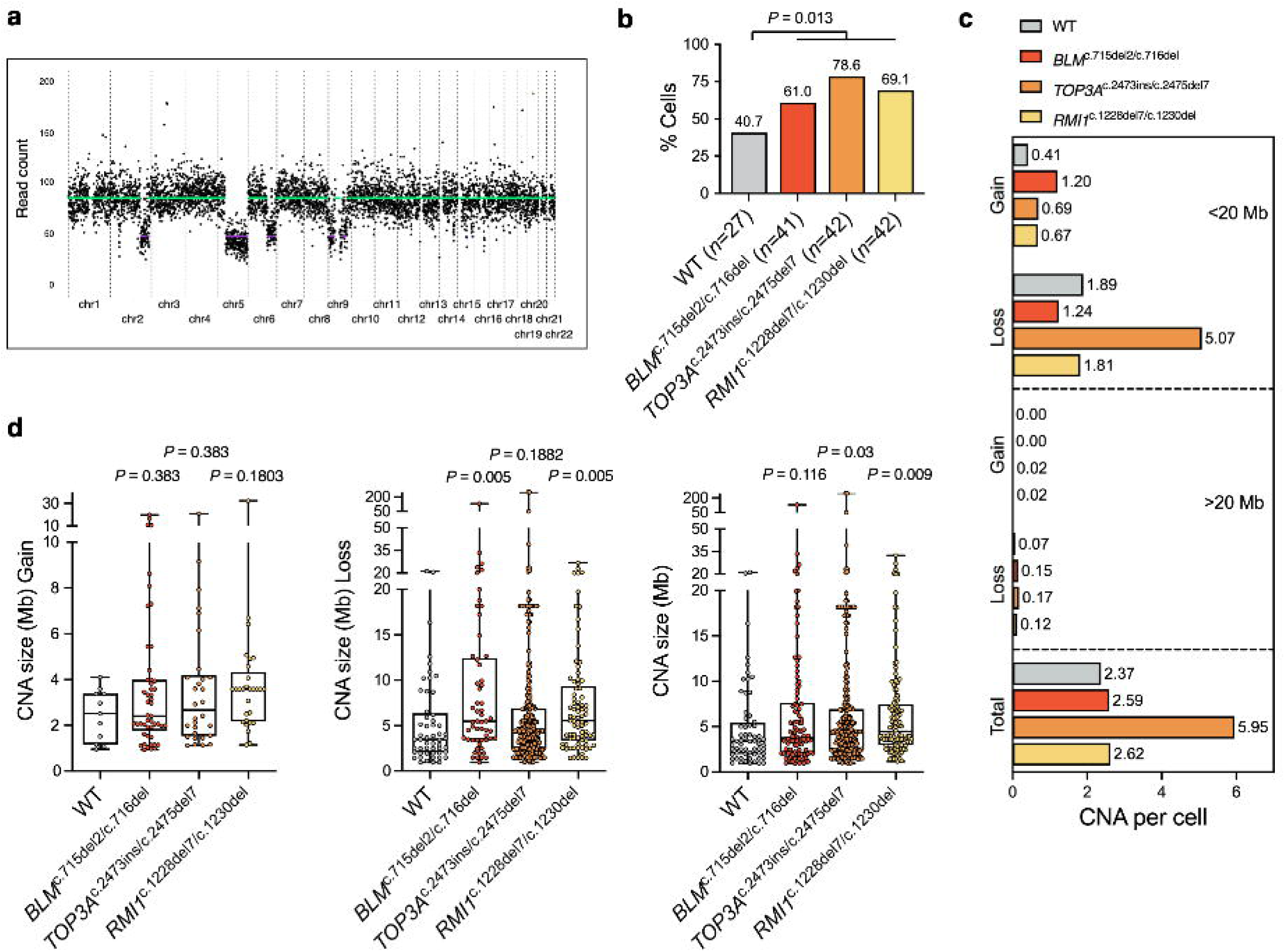
BTRR complex deficiency causes genome-wide copy number alterations (CNAs) in single cells. **a** Example histogram plot of a single *BLM*^c.715del2/c.716del^ cell with deletions identified with Aneufinder^50^. The bin size was set to 500 kb. **b** Proportion of cells with one or more CNA was higher in the BTRR-complex-deficient KO iPSCs than in wild-type cells. *n* indicates the total number of post-QC cells included in the analyses. Kruskal-Wallis test with Dunn’s correction was performed. **c** Rates of CNA based on size (20 Mb threshold) and type (gain or loss) per cell show increased CNA load in BTRR^KO^ iPSCs. **d** Size distribution of gains, losses, and both CNA types revealed that larger CNAs were enriched in BTRR-complex-deficient cells. *p*-values indicate adjusted values determined by Kruskal-Wallis test with Benjamini-Hochberg correction where individual values were compared against WT.

### CNAs are more likely to locate at fragile sites

Genomic loci that are prone to form breaks and structural variation under replication stress are termed fragile sites^52,53^. Common fragile sites (CFS) expression was observed on metaphase chromosomes after replication stress induction, which previously defined most of the CFS at a chromosome band resolution^54^. Genomic regions of fragile sites are associated with late replicating units, which might enter mitosis without completing replication, and this can trigger mitotic DNA synthesis (MiDAS)^38,55^. Moreover, genomic regions undergoing MiDAS were mapped by NGS-based, high-throughput sequencing^31,32^. As a result, the resolution of defined fragile sites substantially increased towards a kilobase scale, spanning approximately 8% of the whole genome (Supplementary Table 2). We studied the genomic localization of detected CNAs in WT and BTRR complex deficiency and analysed the proportion of CNAs mapped to the high-resolution CFS regions. Interestingly, our analysis revealed that in all our cell lines CNAs do not show a stochastic distribution in the genome, but instead are located more frequently at determined CFS of the genome (Fig. 4a). In addition, the proportion of CNAs mapped to the described fragile sites increased in the BTRR^KO^-iPSCs in comparison to WT-iPSCs (Fig. 4b, Supplementary Fig. 5a) and a higher proportion of CNAs associated with large expressed genes identified in the iPSC model (Supplementary Fig. 5b). As mentioned above, CFS were previously defined by cytogenetic methods at a chromosome band scale where recurrent breaks were identified upon replication stress^54,56^. In addition to the MiDAS-seq-defined CFS, we separately analysed the location of CNAs to CFS identified by cytogenetic techniques (Supplementary Table 3) and observed a similar tendency of CNAs predominantly locating at CFS (Supplementary Fig. 6a). The distribution of CNA/CFS overlapping regions showed a non-random distribution of CNAs in *BLM*^c.715del2/c.716del^ and *TOP3A*^c.2473ins/c.2475del7^ cells (Supplementary Fig. 6b).

**Fig. 4.**
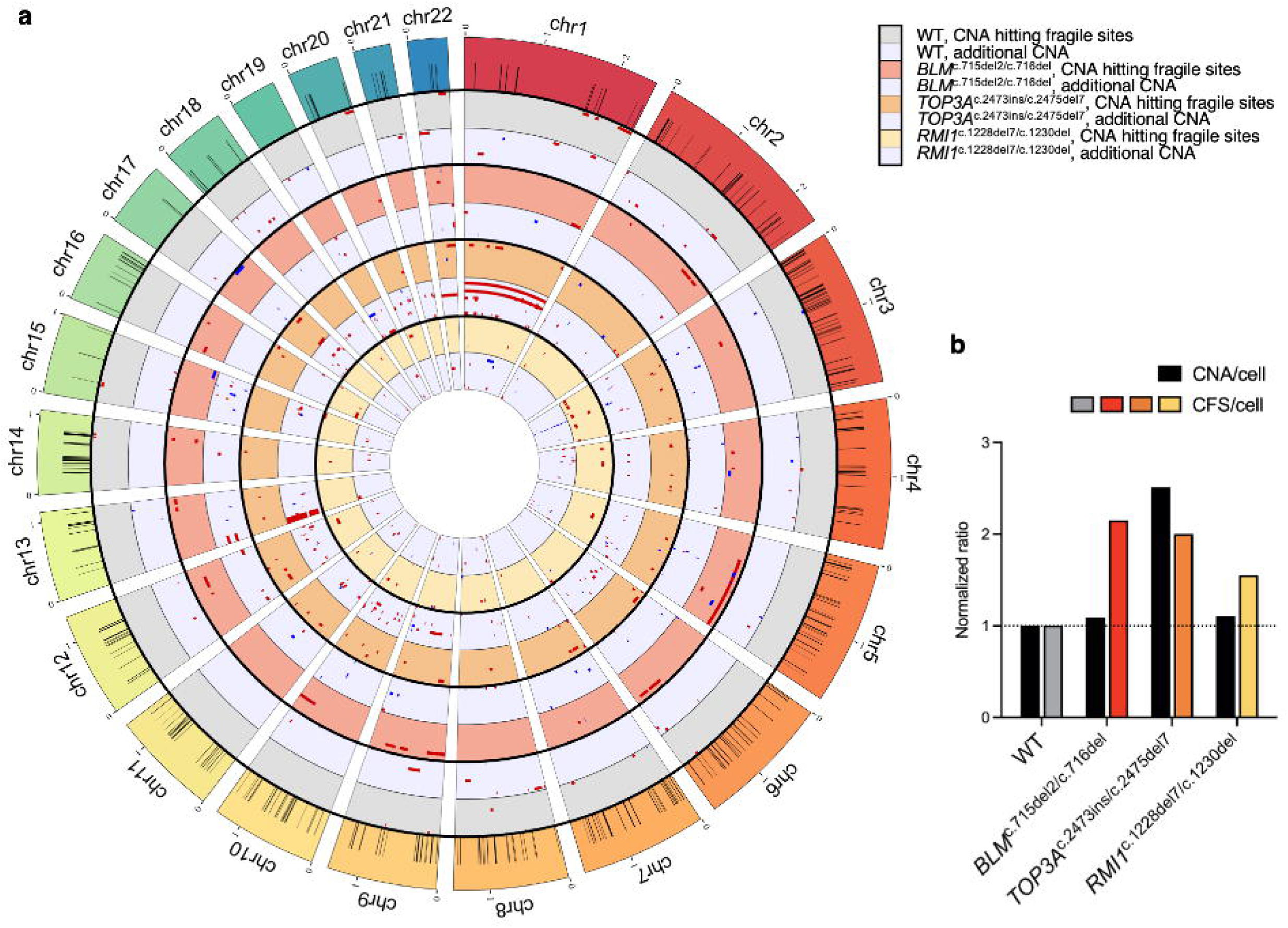
CNAs tend to locate at the common fragile sites (CFS). **a** Circos plot shows CNAs hitting fragile sites (gray, red, orange, and yellow-colored tracks for different samples) and the additional CNAs (light blue-colored tracks). Gains (blue) and losses (red) are shown in single cells as clusters based on genomic location and size. Black labels on the reference chromosomes show the previously identified^31,32^ CFS used in the analysis (see “Supplementary material” for interactive plots labeling each CNA and CFS). **b** The relation of CNA per cell to CFS per cell shows increased CFS affected by CNAs in BTRR^KO^ iPSCs. Ratios were normalized to WT for both values. Whole chromosome aneuploidies were excluded from the CFS analysis.

### Moderate replication stress induces CNA formation and CNA/CFS association

DNA replication stress refers to the slowing or stalling of the DNA replication fork caused by obstacles such as DNA breaks, unresolved secondary structures, or R-loops^57^ – all of which are potential substrates of the BTRR complex. We sought to determine the effect of replication stress on genomic stability in the BTRR-complex-deficient iPSCs by using a moderate concentration of aphidicolin, a well-known DNA polymerase inhibitor and inducer of replication stress^52,58^. The moderate aphidicolin treatment concentration (300 nM, 24 hours) allowed cells to continue cell cycle progression while this treatment also resulted in replication fork stalling by partial inhibition of the DNA polymerase as previously described^28^. We assessed the effect of aphidicolin on cell cycle profiles and observed that the majority of cells were blocked in S phase while a small population of G1 cells was still detectable for single cell sorting before whole genome sequencing (Supplementary Fig. 7a, b). Additionally, to assess replication-stress-associated DNA damage, we quantified 53BP1 foci in G1/early S phase cells after aphidicolin treatment, which is known to cause G1 53BP1 nuclear bodies due to replication stress in the previous S phase^59^. This analysis revealed induced DNA damage levels in G1/early S phase cells in the BTRR complex deficient cells (Supplementary Fig. 7c). We observed that the percentage of affected single-cell genomes was highly significantly increased in the aphidicolin-treated WT sample (89.9% vs 40.7%, *P* <0.0001), confirming the effect of moderate replication stress on CNA formation at single-genome resolution (Fig. 5a). Interestingly, none of the aphidicolin-treated BTRR^KO^ iPSCs showed an additional significant increase in the proportion of cells affected by CNA formation when compared to the non-treated cells. However, our analysis revealed an elevated proportion of CNAs in single-cell genomes of all BTRR-complex-deficient cell lines (Fig. 5b). Further analysis showed that large CNAs were present in the BTRR-complex-deficient and aphidicolin-treated cells (Fig. 5c), occurring mainly as deletions in the KO iPSCs (Supplementary Fig. 8). Complete loss of chromosome 5 was detected in the *BLM*^c.715del2/c.716del^ aphidicolin-treated sample and was the only whole chromosome aberration among aphidicolin-treated cells, suggesting that whole chromosome instability might not have been prominent as an effect of moderate replication stress based on our model system. The size distribution of CNAs in aphidicolin-treated WT, *TOP3A*^c.2473ins/c.2475del7^, and *RMI1*^c.1228del7/c.1230del^ revealed a significant variability when compared to non-treated WT (Fig. 5c). Taken together, our data showed that moderate replication stress strongly induces CNA formation in initially genomically stable WT cells, but that this effect is almost absent in already damaged, unstable BTRR-complex-deficient cells.

**Fig. 5.**
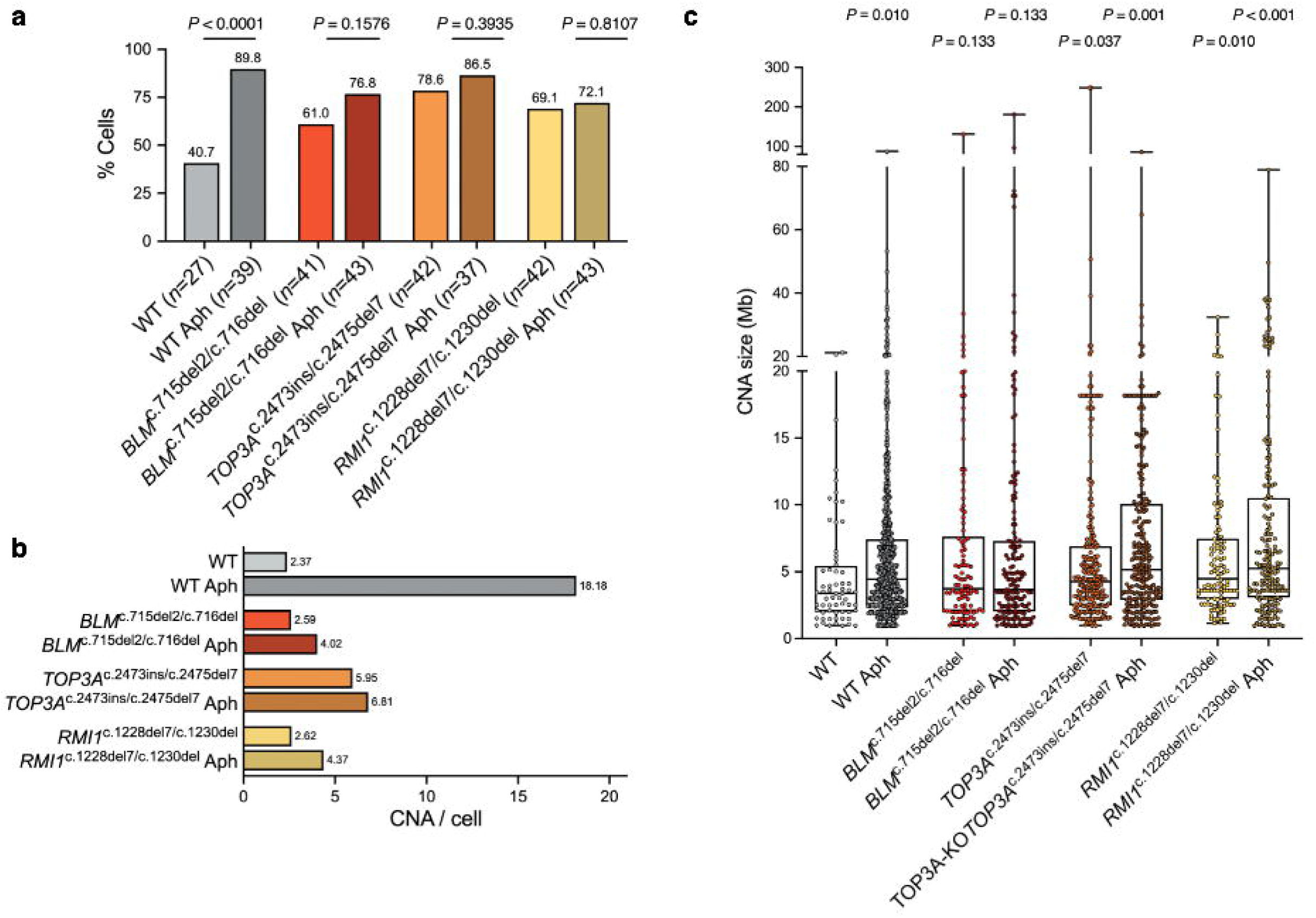
Moderate replication stress induced by aphidicolin results in elevation of CNAs in single cells. **a** Proportion of cells with at least one identified CNA after aphidicolin treatment with 300 nM for 24 hours. *n* indicates the number of post-QC single cells included in the analyses. Two-tailed Mann-Whitney test was performed between non-treated and aphidicolin-treated (Aph) sample pairs. **b** Total numbers of CNAs per single cell exhibit a sharp increase of CNAs in replication stress-induced wild-type cells among all samples. **c** Size distribution (Mb) of all CNAs revealed that large CNAs were present in BTRR-complex-deficient and aphidicolin-treated (Aph) samples. *p*-values indicate the adjusted values determined by Kruskal-Wallis test with Benjamini-Hochberg correction, performed against WT.

To further investigate the characteristics of the genomic regions affected by novel induced CNAs after replication stress, we analysed scWGS data of aphidicolin-treated samples for MiDAS-seq-defined CFS and showed that CNA/CFS linkage was more pronounced under replication stress (Fig. 6a). The most prominent CNA/CFS linkage occurred in the WT cells due to the strong increase of CNA numbers. The proportion of CNA/CFS also increased in the BTRR-complex-deficient cells (Fig. 6b), indicating that CNA formation more likely occurs at CFS under replication stress and at actively transcribing genes (Supplementary Fig. 9a, b). Importantly, replication stress inducers such as aphidicolin cause chromosome breaks at CFS^60^. We additionally analysed the replication-stress-associated CFS and observed a similar increase in CNA/CFS linkage, although this association was less specific due to the much lower chromosome band resolution of the reference CFS (Supplementary Fig. 10). Taken together, our results show that aphidicolin-induced replication stress promotes CNA formation and suggest that CNAs are more likely to occur at fragile sites of the genome.

**Fig. 6.**
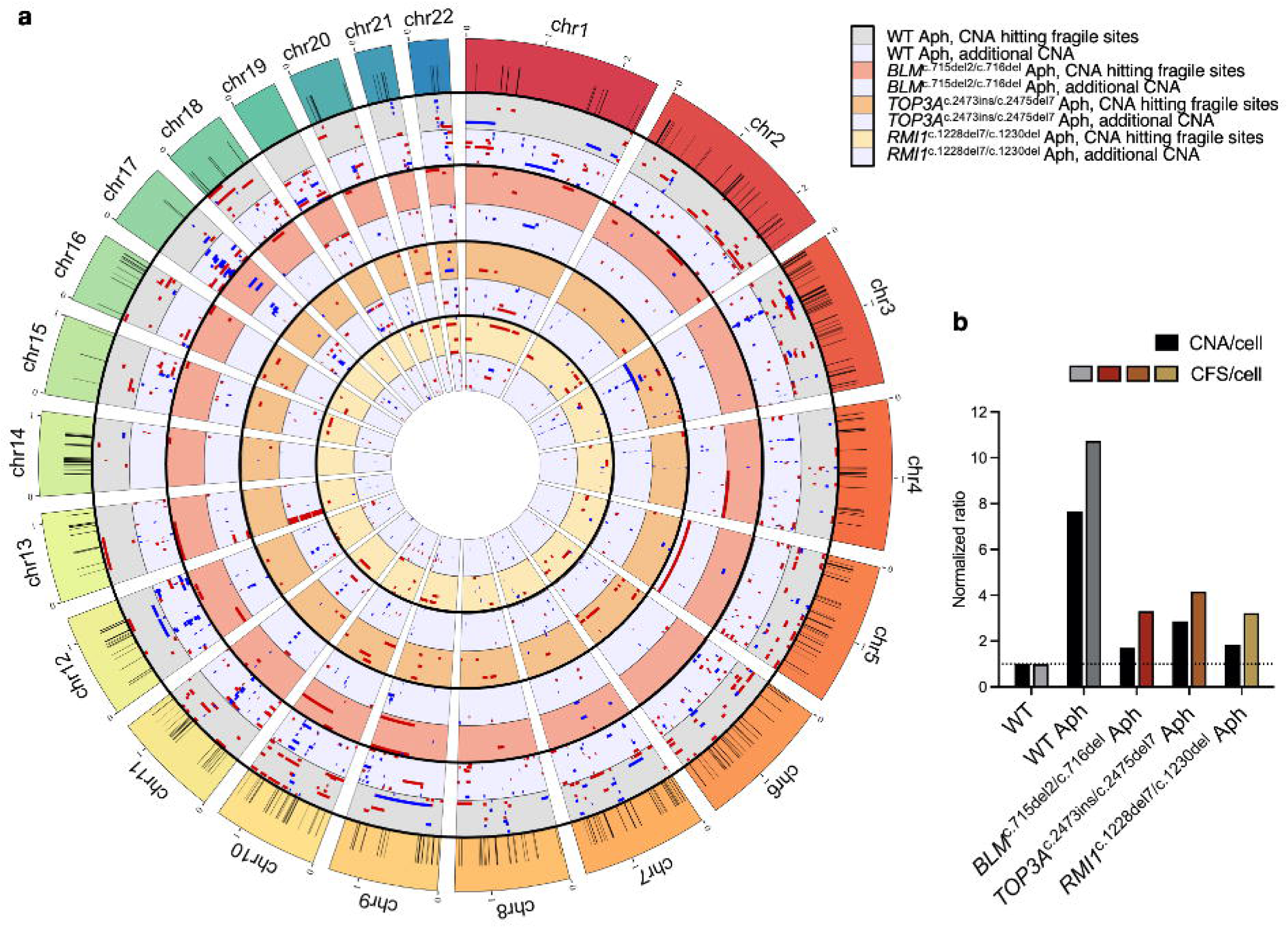
Moderate replication stress promotes CNA association with fragile sites. **a** Circos plot shows CNAs hitting fragile sites (gray, red, orange, and yellow-colored tracks for different samples) and the additional CNAs (light blue-colored tracks). Gains (blue) and losses (red) are shown in single cells as clusters based on genomic location and size. Black boxes on the reference chromosomes show previously identified CFS (see “Supplementary material” for interactive plots labeling each CNA and CFS**). b** The relation of CNA per cell to CFS per cell shows increased CFS affected by CNAs in the aphidicolin-treated (Aph) samples. Aphidicolin-treated samples were normalized to non-treated WT for both ratios. Whole chromosome aneuploidies were excluded from the CFS analysis.

## Discussion

Our study provides novel insights into the characteristics of genomic instability observed in patients with BSyn resulting from BTRR complex deficiency. Using CRISPR/Cas9-based knockout iPSC lines for three members of the BTRR complex, namely, BLM, TOP3A, and RMI1, we showed that genomic instability associated with an increased rate of chromosome segregation errors, favoured error-prone DNA repair, and resulted in *de novo* CNA formation. We further discovered that CNAs were not stochastically distributed in the genome but, instead, showed a tendency to specifically localise at fragile sites.

BSyn is caused by biallelic LoF variants in *BLM*, *TOP3A*, *RMI1,* and *RMI2*^4–6^. It is worth noting that BSyn patients with biallelic variants in these genes do present phenotypic variability potentially depending on the distinct molecular roles of each gene product^5,6,33^. RMI2 has been identified as a complex member and so far only two patients with a BSyn-like phenotype have been reported carrying a homozygous loss of the complete *RMI2* gene^6^. Previously, we reported the milder phenotypic characteristics of patients with *RMI1*-associated BSyn compared to BSyn caused by *BLM* mutations^5,33^. It was interesting to observe that in our isogenic iPSC model for the *RMI1*^c.1228del7/c.1230del^ bi-allelic LoF mutations, the functional consequences on genomic stability were milder compared to the effects seen in the BLM-deficient iPSCs, which could be resulting from the fact that RMI1 is transcribed from a single coding exon, allowing transcripts with premature stop codons to escape nonsense-mediated decay and potentially resulting in hypomorphic allele. Based on the analyses of SCE, lagging chromatin, and chromatin bridges, *RMI1*^c.1228del7/c.1230del^ cells showed a milder cellular BSyn phenotype than *BLM*^c.715del2/c.716del^ cells or *TOP3A*^c.2473ins/c.2475del7^ cells. In addition, we detected a putative truncated RMI1 protein expressed in the *RMI1*^c.1228del7/c.1230del^ cells which could point to a hypomorphic allele, consistent with the mild clinical phenotype reported for the only two *RMI1*-mutated BSyn-like cases described to date^5,33^. In addition, we assessed the repair efficiency after Cas9-induced DSBs which revealed impaired SSTR in the BTRR-complex-deficient cells in line with previous findings on factors required for single-strand template repair^61^. We observed only a few whole chromosome CNAs, which suggested that the lagging chromatin observed previously could have originated mostly from DNA breaks. Notably, the *TOP3A*^c.2473ins/c.2475del7^ cells exhibited the highest increase in CNAs among all samples. Besides its role within the BTRR complex, TOP3A is important for mitochondrial function, as it releases torsional strain and DNA interlinks on mitochondrial DNA for effective replication and transcription^62,63^. One hypothesis could be that mitochondrial dysfunction in *TOP3A*^c.2473ins/c.2475del7^ cells may exacerbate genomic DNA aberrations via increased production and release of reactive oxygen species leading to alterations in nuclear DNA. In addition, TOP3A may also perform S-phase–specific functions that are partly independent of the BTRR complex^64^, however, we observed only a mild increase in replication-associated DNA damage in TOP3A-deficient cells and a similar overall response to replication stress across all samples (Supplementary Fig. 11a, b; Supplementary Figure 7b). These observations suggest that the elevated CNAs in *TOP3A*^c.2473ins/c.2475del7^ cells likely reflect a combination of its nuclear role within the BTRR complex together with additional TOP3A-specific functions.

To further characterize the genomic instability profile of our BTRR^KO^ iPSC model system, we analysed CNAs in single-cell genomes. The applied low-coverage, single-cell WGS was able to detect and measure the heterogeneity of CNAs present in our cellular models, while in comparison, 30 x coverage, longread WGS (lrWGS) was not able to detect these differences due to the high heterogeneity of CNAs in single cells and increasing the coverage of bulk DNA lrWGS to e.g. 200 is currently not feasible due to technical limitations during the bioinformatic processing. The application of AneuFinder^50^ for scWGS data analysis has demonstrated both the strengths and limitations of this approach. One of the key advantages of scWGS is its ability to detect CNAs at the single-cell level, allowing for a more precise understanding of cellular heterogeneity, which would otherwise be masked in bulk sequencing. AneuFinder provides a robust framework for this analysis by binning aligned reads, correcting for biases such as GC content variation, and applying statistical models to classify chromosomal aneuploidies. Among the available models, we employed eDivisive, which represents the most refined and improved approach for segmentation. This model enhances the detection of CNAs by leveraging a statistical method that minimizes assumptions about underlying distributions, thereby increasing accuracy and reducing false-positive calls.

The strength of AneuFinder, particularly when using eDivisive, is its ability to provide high-resolution CNA detection while effectively managing biases introduced by sequencing artifacts and amplification noise. By using a wild type euploid reference, common artifacts are filtered out, ensuring that detected CNAs are meaningful and reproducible. Additionally, the method allows for clear visualization of results through heatmaps and histograms, aiding in the interpretation of chromosomal aberrations. However, despite these advantages, certain limitations exist. The reliance on a euploid reference dataset for bias correction can introduce errors if the reference is not well curated. Additionally, while eDivisive improves segmentation accuracy, scWGS as a whole remains susceptible to noise, particularly in highly repetitive genomic regions, which must often be excluded from analysis. This exclusion, while necessary, may limit the detection of certain structural variations. In addition, low coverage model of scWGS limits an in-depth analysis of the sequences flanking the genome segments with copy number losses or gains. Further development of high-coverage single-cell sequencing techniques will provide the opportunity to uncover the mechanisms of CNA formation triggered by BTRR complex dysfunction. In summary, AneuFinder, particularly when using the eDivisive model, proves to be a powerful and refined tool for CNA detection in single-cell sequencing, offering high accuracy and robust statistical modelling while facing inherent technical challenges that must be accounted for in data interpretation in future studies.

The generation of BLM-null murine models has proven to be challenging, with reports indicating embryonic lethality upon complete loss of BLM, although a hypomorphic allele is viable^65–67^. Likewise, Top3a-null mice are embryonically lethal^68^, and *TOP3A* alleles identified in BSyn-like patients are likely hypomorphic rather than complete loss-of-function variants^5^. These data further support the interpretation that our CRISPR-engineered *TOP3A* frameshift alleles may similarly represent hypomorphic variants rather than full null alleles. Therefore, our BTRR^KO^ iPSC model system represents a valuable cellular system to analyse genomic and molecular mechanisms underlying BSyn pathogenesis.

We observed a modest increase in CNA levels upon external replication stress induced by _aphidicolin treatment in *BLM*_c.715del2/c.716del_, *TOP3A*_c.2473ins/c.2475del7 _and *RMI1*_c.1228del7/c.1230del _single_ cells. Our flow cytometry data indicated that aphidicolin treatment resulted in a comparable shift in cell cycle profiles across all genotypes, suggesting a similar initial cell cycle response to replication stress. Therefore, the mild CNA induction observed in the knockout lines is unlikely to be explained by differences in the immediate response to aphidicolin. Instead, these cells may already experience substantial endogenous replication stress, leaving less capacity for additional CNA accumulation. Alternatively, cells with the highest levels of DNA damage may preferentially activate cell cycle checkpoints and fail to progress through the cell cycle before analysis. However, our experimental design did not include parallel quantification of cell viability, apoptosis, or cell cycle distribution at the point of nuclei harvest, and we therefore cannot formally exclude that selective loss of cells with very high CNA loads contributes to the apparent saturation observed in aphidicolin-treated knockout samples. These findings suggest that the limited CNA induction following aphidicolin treatment likely reflects a combination of intrinsic replication stress and selective loss of highly damaged cells.

Strikingly, in our scWGS studies, the BTRR-complex-deficient cells showed increased rates of CNA formation located predominantly at fragile sites. High levels of CFS-associated CNAs were also identified in the WT cells, clearly indicating that genomic alterations do not distribute evenly in the genome. Instead, they occur more often at fragile sites of the genome, which are representing “error-prone genomic regions”, e.g. mechanistically defined by repetitive structures, specific sequence combinations such as high GC contents, or under-replicated DNA^54^. The MiDAS-seq-defined CFS we used were originally mapped in cancer cell lines; despite known cell-type specificity of fragile-site expression, the reproducible CNA/CFS association observed across all three BTRR knockouts and under aphidicolin treatment support that a biologically conserved core of fragile regions is captured by this reference. In addition, large expressed genes were shown to result in fragile site expression in a cell-type specific manner^69,70^, which was supported by our data using an iPSC-specific transcriptional map. The susceptibility of regions containing CFS to harbor CNAs was in line with the alternative view of chromosomal evolution that suggests the predominant occurrence of breaks within fragile regions^71^. The underlying mechanism of increased CNA/CFS occurrence in BLM- and TOP3A-deficient cells could involve the compensatory activity of endonucleases, which process under-replicated DNA intermediates thereby leading to deletions and SCEs^42,44^. Furthermore, UFBs and micronuclei related to unresolved DNA structures from the CFS were identified in BLM-deficient cells^30,38^, however, TOP3A and RMI1 deficiency have not been associated with CNA/CFS occurrence so far. Of note, our analysis revealed similar rates of elevation of CNA/UFB formation in all three KO iPSCs, which suggests that these three BTRR complex members were equally important for UFB removal^40,42^. Based on our UFB and CFS findings, we propose that the observed formation of UFBs partially originated from CFS, regions acquiring aberrations during chromosome segregation, later manifesting as CNAs associated with CFS. In addition, although the bin size-dependent resolution of our scWGS data did not allow the analysis of specific sequence environment at CNA junctions, it is noteworthy that replication stress has been linked to POLQ-mediated end-joining (TMEJ)-mediated structural variant formation at common fragile sites^72^ and we cannot rule out the possibility that the identified CNAs were originating from TMEJ mechanism.

Applying moderate replication stress also increased the formation of novel CNAs, predominantly located at CFS. It is important to acknowledge that the reference CFS used in this study were regions undergoing mitotic DNA synthesis after replication stress induced by aphidicolin^31,32^. Most of the known fragile sites were identified by cytogenetic methods after aphidicolin-induced replication stress, which we also used for linking the CNA regions^54^. We stress that the genome contains further, as yet unknown fragile regions, which can, e.g., be induced by different factors such as hydroxyurea^51^ and which are additionally important for CNA formation. Consequently, our current study provides a single-cell-based molecular perspective into the predominant occurrence of CNA at known CFS identified mainly with aphidicolin treatment.

In summary, our work provides novel insights into changes of genomic stability caused by BTRR complex deficiency. iPSC models deficient for BLM, TOP3A, and RMI1 showed chromosome segregation aberrations and impaired HDR efficiency. Single-cell whole genome sequencing analyses revealed increased rates of CNA formation in BTRR-complex-deficient cells. Finally, our data show evidence of the predominant occurrence of CNAs at fragile sites as detected in single-cell genomes of the BTRR-complex-deficient cells with and without replication stress.

## Methods

### iPSC culture

Human iPSC lines were maintained on Matrigel^®^-coated (growth factor reduced, Corning Incorporated, 354230) 6-well plates in StemFlex™ medium (Thermo Fisher Scientific, A3349401) in a humidified incubator at 37°C supplemented with 5% CO_2_. Cells were split twice every week using Versene solution (Thermo Fisher Scientific, 15040066) and treated with 2 µM Thiazovivin (Merck Chemicals, 420220) for approximately 24 hours after the split. Medium was changed every other day. The wild-type iPSC line UMGi014-C clone 14, (isWT1.14, referred to as WT), was generated from fibroblasts of a healthy individual using integration-free Sendai virus and was described previously^73^. Knockout cell lines UMGi014-C-19 clone 1D1 (isWT1-BLM-KO.1D1, referred to as *BLM*^c.715del2/c.716del^), UMGi014-C-25 clone 1E10 (isWT1-TOP3A-KO.1E10, referred to as *TOP3A*^c.2473ins/c.2475del7^), and UMGi014-C-49 clone 1F11 (isWT1-RMI1-KO.1F11, referred to as *RMI1*^c.1228del7/c.1230del^) were derived from the wild-type cell line using CRISPR/Cas9. After genome editing, the pluripotency of cell lines was confirmed via immunocytochemistry and FACS analyses as previously described^74^. The exact passage numbers of all iPSC lines at the time of each experiment are listed in Supplementary Table 6 to ensure full transparency regarding cell stage and experimental comparability.

### CRISPR/Cas9-based genome editing

The guide RNA sequences for each member of the BTRR complex were designed using CRISPOR^75^ tool and the oligonucleotides were purchased from Integrated DNA Technologies (IDT™). Sequences of crRNAs are summarized in Supplementary Table 4. On the day of transfection, 80% confluent iPSCs were treated with 2 µM Thiazovivin for one hour before transfection. For genome editing, the Alt-R CRISPR-Cas9 crRNA and Alt-R CRISPR-Cas9 tracrRNA were mixed (1:1, vol/vol) and incubated at 95°C for 5 minutes. The guide RNA solution was mixed with Alt-R *S.p.* Cas9 Nuclease V3 (at a 3:1 molar ratio and incubated at room temperature (RT) for 20 minutes. 1 µM Electroporation Enhancer (IDT™) was added and the mixture was stored at 37°C until use. Two million iPSCs were transfected using Amaxa P3 Primary Cell 4D-Nucleofector kit (Lonza, V4XP-3024) with CA-137 setting according to manufacturer’s instructions. Transfected cells were seeded into StemFlex medium supplemented with 2 µM Thiazovivin. A medium change without Thiazovivin was performed on the third day. Confluent iPSC culture was cryopreserved in culture medium supplemented with 2 µM Thiazovivin and 10% DMSO. Genomic DNA of bulk-transfected cells was extracted using DNeasy Blood & Tissue kit (Qiagen, 69504). An approximately 350-bp DNA region spanning the PAM site was amplified with Multiplex PCR kit (Qiagen, 206143) and the PCR product was cleaned with Msb Spin PCRapace kit (Stratec Molecular). The NGS library was prepared with the PCR product using SureSelect QXT (Agilent Technologies, G9683A) and sequenced on an Illumina NextSeq550. Raw sequencing files were analysed with Cas-Analyzer^76^ and transfection efficiencies were determined according to the edited allele counts.

### Single-cell-derived clone analysis

Bulk-transfected iPSCs were thawed and passaged at least once before singularization. 70% confluent cell culture was treated with 1x Revita cell supplement (Thermo Fisher Scientific, A2644501) for half an hour and cells were dissociated using 1x TrypLE™ (Thermo Fisher Scientific, 12604013). Cell pellets were resuspended in cold DPBS and single cells were sorted with CellenONE (ScienION) system to 96-well plates containing StemFlex medium supplemented with 1x Penicillin-Streptomycin (Thermo Fisher Scientific, 15140122) and 1x Revita cell supplement. Plates were incubated at 37°C with 5% CO_2_ for at least one week before medium change. When colonies reached sufficient size, images of the colonies were taken with Incucyte S3 (Sartorius) and single-cell-derived clones were passaged to 12-well plates using Versene solution. At 80% confluency, cultures were cryopreserved and an aliquot was used for DNA extraction. DNA extraction was performed using DNeasy blood and tissue kit (Qiagen, 69504). The genomic DNA of each clone was evaluated with PCR amplification followed by Sanger sequencing as previously described^33^. The genotypes were analysed for the corresponding gene using CRISP-ID^77^. All primers used for PCR reactions are shown in Supplementary Table 5.

### Western blot

Western blot analysis was used for confirmation of the knockouts on protein level as described previously^78^. The following antibodies were used: anti-BLM (1:1000, Abcam, ab2179), anti-TOP3A (1:2000, Proteintech, 14525-1-AP), anti-RMI1 (1:1000, Proteintech, 14630-1-AP), anti-Tubulin (1:5000, Abcam, ab8505), anti-H3 (1:1000, Abcam, ab52866), anti-rabbit IgG-HRP (1:10.000, Santa Cruz, sc2004), anti-mouse IgG-HRP (1:10.000, Santa Cruz, sc2005). The membrane was developed using horseradish peroxidase-based chemiluminescence detection (WesternBright ECL HRP substrate, Advansta K-12045-D20) and visualized using the ChemiDoc™ Touch Imaging system (Bio-Rad).

### Pluripotency analyses

Immunofluorescence and flow cytometry-based pluripotency analyses of generated KO iPSCs were performed as previously described^74^. In brief, for immunocytochemistry, colonies grown on Matrigel-coated coverslips were fixed in 4% paraformaldehyde followed by blocking in 1% bovine serum albumin (BSA) and permeabilization in 0.2% Triton X100. Primary antibodies were incubated on coverslips overnight at 4°C. The pluripotency markers and the dilutions used are as follows: anti-SSEA4 FITC (Miltenyi Biotec, 130-098-371), 1:30; anti-OCT3/4 PE (Miltenyi Biotec, 130-120-236), 1:50; anti-SOX2 PE (Miltenyi Biotec, 130-104-941), 1:50; anti-NANOG mouse IgG1 (Thermo Fisher Scientific, MA1-017), 1:100; and anti-TRA-1-60 mouse IgM (Abcam, ab16288), 1:200. Secondary antibodies were anti-mouse IgG Alexa Fluor 555 (Thermo Fisher Scientific, A-2142) and anti-mouse IgM Alexa Fluor 488 (Abcam, ab150121) used with 1:1000 dilution. DNA was counterstained with 43.75 µg/ml DAPI and coverslips were mounted with ProLong™ Gold Antifade Mountant (Invitrogen, P10144). Images were taken with Revolve fluorescent microscope (Echo). For flow cytometry analysis, fixed single cells were blocked, permeabilized and incubated with fluorescence-conjugated OCT4 and TRA-1-60 antibodies as described^74^. DNA was labeled with Hoechst 3342 (Thermo Fisher Scientific, H3570). Stained samples were analysed in the LSRII flow cytometer X20 (BD Biosciences) using BD FACSDiva software. The raw data were analysed with Floreada.io online tool (https://floreada.io/).

### Sister chromatid exchange assay

Cells were treated with 10 μM BrdU (Thermo Fisher Scientific, B23151) for 42 hours before harvesting. In the last 3 hours, 1 μM colcemid (Merck Chemicals, 234109) was added to the medium. Cells were then collected using 1x TrpLE (Thermo Fisher Scientific, 12604021) and centrifuged with 400g for 5 minutes. Cell pellets were resuspended in 0.075 M KCl and incubated for 13 minutes at 37°C followed by centrifugation. Pelleted cells were fixed in ice-cold methanol/acetic acid (3:1 vol/vol) fixative solution, spread onto clean slides and dried for at least two days. Slides were incubated in 1 μg/ml Hoechst 33258 (Thermo Fisher Scientific, H3569) for 15 minutes in the dark, rinsed in water, and irradiated with UV-C (0.260J/cm^2^) in freshly prepared 2x SSC buffer. Slides were incubated at 65°C for two hours in 2x SSC buffer followed by a 5% Giemsa staining (Merck Chemicals, 109204). Air-dried slides were analysed using Zeiss Axioplan microscope. Exchanges at the centromeres were not counted.

### Immunofluorescence assays

For detection of ultrafine bridges, iPSC colonies were cultured on Matrigel-coated (1:80 dilution) 1N coverslips in 6-well plates until 80% confluency. Unsynchronized iPSCs on coverslips were washed once with DPBS and fixed in 4% paraformaldehyde containing 0.1% Triton-X100 for 20 minutes at RT. Fixed cells were washed two times with DPBS and incubated in 0.15 M NH_4_Cl for 20 minutes at RT. Coverslips were washed three times with DPBS and blocked in 2% BSA in DPBS for 20 minutes. Anti-ERCC6L (Abnova, H00054821-M01) antibody with 1:150 dilution was added to the coverslips and incubated overnight at 4°C. Next day, coverslips were washed twice with 2% BSA and incubated with secondary antibody anti-mouse IgG(H+L) Alexa Fluor^TM^ 488 (1 µg/mL, Thermo Fisher Scientific, A11017) for 45 minutes in the dark at RT. Cells were washed twice with DPBS and coverslips were dipped into 100% ethanol followed by 43.75 µg/ml DAPI and 10 µg/ml RNase incubation for five minutes. Coverslips were washed with DPBS, H_2_O, and, finally, with 0.01M NaN_3_. They were mounted to clean microscopy slides using ProLong™ Gold Antifade Mountant (Invitrogen, P10144) and visualized with Revolve fluorescent microscope (Echo). Three repeats were prepared and at least 50 anaphases were quantified for the each analysis. Images were analysed with ImageJ v2.1 (FIJI). DNA damage levels at G1/early S phase were analysed by quantitative image-based cytometry using ScanR (Olympus) microscope and its software using segmentation based on DAPI and automated 53BP1 foci detection. Cells grown on cover slips were fixed, permeabilized and blocked as described above. Primary antibody incubation with anti-53BP1 (Merck, MAB3802, 1:500) was performed for 1 h at room temperature, followed by two washes in PBST (0.2% Tween-20 in DPBS) and incubation with Alexa Fluor Plus 488-conjugated anti-mouse IgG secondary antibody (Thermo Fisher Scientific, A32723; 1:1000) in blocking buffer containing 0.5 μg/mL DAPI for 1 h at room temperature in the dark. Coverslips were washed five times in PBST, dipped into dH_2_O and air dried before mounting in 10% Mowiol 4-88 (Merck), 25% glycerol, 100 mM Tris-HCl, pH 8.5 solution. For the assessment of DNA damage levels in S-phase cells, same procedure was applied with the exception of an additional Click reaction after primary antibody incubation. In brief, cells were incubated with 10 µM EdU 30 min before harvesting and EdU was labeled with Alexa Fluor 647 using a Click-iT EdU Cell Proliferation Kit for Imaging (C10340, Thermo Fisher Scientific) according to the manufacturer’s instructions. Coverslips were imaged using IX60 (Olympus) microscope and images were analysed using FIJI with custom macros for segmentation and automated foci quantification.

### Flow cytometry analyses for cell cycle profiling

10 µM EdU was added to asynchronous and aphidicolin (24 hours, 300 nM)-treated samples 30 minutes before harvesting. Cells were dissociated with Versene, resuspended in DPBS and pelleted with centrifugation (500G, 5 min). Pellets were resuspended and fixed in 4 & PFA for 15 minutes at room temperature, permeabilized in 0.2 % Triton X-100 (5 min, room temperature) and blocked in 2 % BSA for 1 hour at room temperature. Cells were pelleted by centrifugation and resuspended in Click reaction mix prepared according to Click-iT EdU Cell Proliferation Kit Alexa Fluor™ 488 dye (C10637, Thermo Fisher Scientific) manufacturer’s instructions. Cells were washed two times in DPBS and resuspended in 0.5 μg/mL DAPI solution in PBS. Samples were analysed with CytoFlex (Beckman Coulter) and FlowJo software (version 11).

### Cas9-induced DSB repair efficiency assay

To evaluate the potential of single-strand template repair (SSTR) versus non-homologous end joining (NHEJ) in the individual knockout iPSC lines, CRISPR/Cas9 technology was used to introduce a specific gene variant c.7422G>C/p.Arg2474Ser into the *RYR2* gene. The ribonucleoprotein-based CRISPR/Cas9 complex targeting the *RYR2* gene locus was combined with or without a single-stranded oligonucleotide template containing the desired base pair exchange and introduced into iPSCs by nucleofection, as described above. Three days post-transfection, genomic DNA from bulk-transfected cells was extracted and analysed by amplicon sequencing for precise template-mediated editing (SNP introduction) versus NHEJ editing (indels), as described above. Sequence of crRNA is shown in Supplementary Table 4.

### Single-cell whole genome sequencing

For sequencing, iPSC samples were detached using TrypLE™, washed once with DPBS, and pelleted. To generate nuclei, cells were resuspended in cell lysis buffer (100 mM Tris-HCl pH 7.4, 154 mM NaCl, 1 mM CaCl_2_, 500 µM MgCl_2_, 0.2% BSA, 0.1% NP-40, 10 µg/mL Hoechst 33358, 2 µg/mL propidium iodide in ultra-pure water) and incubated on ice in the dark for 15 minutes to complete lysis. Resulting cell nuclei were gated for the G1 phase (as determined by Hoechst and propidium iodide staining) and sorted into wells of 96-well plates containing freezing buffer on a MoFlo Astrios cell sorter (Beckman Coulter). For single-cell sequencing, a single nucleus was deposited per well. 96-well plates containing nuclei and freezing buffer were stored at −80°C until further processing. Automated library preparation was then performed as previously described^79^. Libraries were sequenced on a NextSeq 500 (Illumina; up to 77 cycles – single end) and the raw data was generated.

For the pre-processing of scWGS data, first the input fastq files were systematically screened for potential quality or bias issues with fastqc (version 0.11.8)^80^. Data quality parameters included (1) per base sequence quality, (2) per base GC content, (3) per base N content, (4) sequence duplication levels, and (5) overrepresented sequences and adapter contaminations. In the second step, files were screened for potential contaminations of different species or organisms using fastq-screen (version 0.13.0)^81^. As each of the quality parameters and potential sequence contaminations were per sample level, all quality samples were aggregated into a single report using MultiQC (version v1.10.1)^82^. The proportion of insufficient read quality bases or contaminated sequences were then trimmed out by identifying the poor-quality read proportions individually adapted towards the dataset. The provided quality control metrics from fastqc were further processed via an in-house R-script to be applicable for individual trimming of reads via bbduk (version 38.69)^83^. The quality-controlled input files were then aligned against a human reference, using bowtie2 (version 2.3.5.1)^84^. Duplicated reads introduced on sequence amplification step were removed using Picard Tools (version 2.20.2) (http://broadinstitute.github.io/picard/) and the read order was fixed with samtools sort (version 1.9) .

After the complete preprocessing of the input data, AneuFinder tool package (version 1.14)^50^ was used for identifying CNAs in single cells. It counted the aligned reads into 500 kb bins, corrected for over- and under-representation of sequence reads, and addressed the GC-biases. This was achieved by selecting the wild-type cell line (WT) as the euploid control. Artifacts in highly repetitive regions e.g. centromeric regions and regions with extremely high or low read counts identified in the euploid reference were excluded from the analysis by a blacklist file. Afterwards, an adaption of a hidden Markov model was applied to the binned reads ranging from several biological states e.g. nullisomy up to decasomy. With the blacklist-masked euploid reference, the quality-filtered data were then analysed for chromosomal aneuploidies via histograms and heatmaps. CNAs identified in two or more wild-type cells were defined as clonal CNAs and excluded from the KO samples once identified. X and Y chromosomes were excluded from the analysis.

### Analysis of common fragile sites

CFS regions determined by MiDAS-seq^31,32^ were used in the analysis of CNAs. Alternatively, a custom-made fragile site database was constructed from previous reports based on cytogenetic assays^70,86–89^ and by using the band coordinates from UCSC^90^ (http://genome.ucsc.edu, hg38, Supplementary Table 3). The criteria for CNA/CFS interaction was at least 1 bp overlap between the two regions. Circos plots were generated with BioCircos^91^.

### Single-cell transcriptome sequencing

iPSC samples were detached using TrypLE, washed once with DPBS, and pelleted. Sample preparation and data processing for single-cell RNA sequencing were performed as previously described^78^. In brief, cells were stained with Hoechst 33342 and Propidium Iodide (ReadyProbes Cell Viability Imaging Kit, Thermo Fisher Scientific) and single viable cells were isolated using the ICELL8 Single-Cell System (Takara Bio). Following cell lysis, cDNA synthesis was performed using the SMART-Seq Single Cell Kit (Takara Bio), and sequencing libraries were prepared using the Nextera XT DNA Library Preparation Kit (Illumina). Libraries were sequenced on an Illumina HiSeq4000 platform. Sequencing reads were demultiplexed using the demux script of the CogentAP analysis pipeline version 1.0 (Cogent NGS Analysis Pipeline, Takara Bio). The single-end 51-bp reads were processed using the CogentAP workflow, including adapter and quality trimming with Cutadapt (version 1.18). Reads were aligned to the GRCh38.12 human reference genome (Ensembl release 94) using STAR (version 2.7.8a), and gene-level expression counts were generated with featureCounts (version 2.0.1). Cells with fewer than 10,000 reads or fewer than 300 detected genes were excluded. Additional filtering removed cells with total counts exceeding three median absolute deviations from the median log10 library size, as well as cells with mitochondrial read fractions >0.2 or intergenic read fractions >0.1.

### Statistical analyses

Rates of sister chromatid exchanges were compared using two-tailed Mann-Whitney test. Analysis of chromosome segregation events included three biological replicates for each sample and pairwise two-tailed t-test with Welch’s correction was performed for statistics. Proportions of single cells with at least one identified CNA were compared by assigning binominal values (0 and 1) to each type of cells. The assigned values were then subjected to two-tailed t-test with Welch’s correction. Size distribution of identified CNA were compared using Kruskal-Wallis test with Benjamini-Hochberg correction. All tests were performed using GraphPad Prism 8.3.1.

## Data availability

The data that support the findings of this study are available from the corresponding author upon reasonable request. Sequencing data is deposited in NIH/SRA platform and accessible under SUB15809629.

## Code availability

Custom codes established in this study have been deposited on GitHub (https://gitlab.gwdg.de/wolff16/scwgs).

## Acknowledgments

We thank Karin Boss for critically reading the manuscript. This work was supported by the German Research Foundation (DFG) under Germany’s Excellence Strategy (EXC 2067/1-390729940) and FOR2800 to B.W. and H.B., by the German Federal Ministry of Education and Research (BMBF)/German Center for Cardiovascular Research (DZHK) to B.W., by the Federal Ministry of Education and Research (BMBF) as part of the German Center for Child and Adolescent Health (DZKJ) under the funding code 01GL2402A, by a Cancer Research UK Senior Cancer Research Fellowship to A.N.B. (RCCSCF-May23/100001) and a Novo Nordisk Foundation Hallas-Møller Ascending Investigator award to A.N.B. (NNF23OC0082165).

## Author contributions

B.W. designed the study. I.I.G. performed and analysed the iPSC experiments. A.Wo. analysed the scWGS data. A.V.B. contributed to the generation of the iPSC lines. A.Wi. and M.R. contributed to the UFB experiment design and analysis. A.T. contributed to the single-cell sorting for scWGS. R.W. contributed to scWGS raw data analysis. C.M., L.A., S.K., G.Y., and A.Z. provided support in cytogenetics and NGS-based molecular genetics experiments. D.C.J.S. and F.F. contributed to the design and sequencing of scWGS. H.B. designed the replication stress-associated experimental set-up. G.Y. and A.Z. contributed to scWGS data analysis. L.C. designed the CRISPR/Cas9-based experiments. B.W. and A.N.B. supervised the project. I.I.G. and B.W. wrote the manuscript with input from all co-authors.

## Supplementary information

The description of supplementary material is given in the online version.

## References

1. Jackson, S. P. & Bartek, J. The DNA-damage response in human biology and disease. Nature 461, 1071–1078 (2009).

2. Cunniff, C., Bassetti, J. A. & Ellis, N. A. Bloom’s syndrome: Clinical spectrum, molecular pathogenesis, and cancer predisposition. Mol. Syndromol. 8, 4–23 (2017).

3. Taylor, A. M. R. et al. Chromosome instability syndromes. Nat. Rev. Dis. Primer 5, 64 (2019).

4. German, J., Sanz, M. M., Ciocci, S., Ye, T. Z. & Ellis, N. A. Syndrome-causing mutations of the BLM gene in persons in the Bloom’s Syndrome Registry. Hum. Mutat. 28, 743– 753 (2007).

5. Martin, C. A. et al. Mutations in TOP3A cause a Bloom syndrome-like disorder. Am. J. Hum. Genet. 103, 221–231 (2018).

6. Hudson, D. F. et al. Loss of RMI2 increases genome instability and causes a Bloom-like syndrome. PLoS Genet. 12, 1–24 (2016).

7. Wu, L. et al. The Bloom’s syndrome gene product interacts with Topoisomerase III. J. Biol. Chem. 275, 9636–9644 (2000).

8. Hu, P. et al. Evidence for BLM and Topoisomerase IIIα interaction in genomic stability. Hum. Mol. Genet. 10, 1287–1298 (2001).

9. Raynard, S., Bussen, W. & Sung, P. A double holliday junction dissolvasome comprising BLM, topoisomerase IIIα, and BLAP75. J. Biol. Chem. 281, 13861–13864 (2006).

10. Yin, J. et al. BLAP75, an essential component of Bloom’s syndrome protein complexes that maintain genome integrity. EMBO J. 24, 1465–1476 (2005).

11. Wu, L. et al. BLAP75/RMI1 promotes the BLM-dependent dissolution of homologous recombination intermediates. Proc. Natl. Acad. Sci. U. S. A. 103, 4068–4073 (2006).

12. Singh, T. R. et al. BLAP18/RMI2, a novel OB-fold-containing protein, is an essential component of the Bloom helicase-double Holliday junction dissolvasome. Genes Dev. 22, 2856–2868 (2008).

13. Xu, D. et al. RMI, a new OB-fold complex essential for Bloom syndrome protein to maintain genome stability. Genes Dev. 22, 2843–2855 (2008).

14. Wu, L. & Hickson, I. D. The Bloom’s syndrome helicase suppresses crossing over during homologous recombination. Nature 426, 870–874 (2003).

15. Mankouri, H. W. & Hickson, I. D. The RecQ helicase–topoisomerase III–Rmi1 complex: a DNA structure-specific ‘dissolvasome’? Trends Biochem. Sci. 32, 538–546 (2007).

16. Harami, G. M. et al. The toposiomerase IIIalpha-RMI1-RMI2 complex orients human Bloom’s syndrome helicase for efficient disruption of D-loops. Nat. Commun. 13, 654 (2022).

17. van Brabant, A. J., Stan, R. & Ellis, N. A. DNA helicases, genomic instability, and human genetic disease. Annu. Rev. Genomics Hum. Genet. 1, 409–459 (2000).

18. Ellis, N. A. et al. The Bloom’s syndrome gene product is homologous to RecQ helicases. Cell 83, 655–666 (1995).

19. Champoux, J. J. DNA topoisomerases: Structure, function, and mechanism. Annu. Rev. Biochem. 70, 369–413 (2001).

20. Bocquet, N. et al. Structural and mechanistic insight into Holliday-junction dissolution by topoisomerase IIIα and RMI1. Nat. Struct. Mol. Biol. 21, 261–268 (2014).

21. Deans, A. J. & West, S. C. FANCM connects the genome instability disorders Bloom’s Syndrome and Fanconi Anemia. Mol. Cell 36, 943–953 (2009).

22. Hoadley, K. A. et al. Defining the molecular interface that connects the Fanconi anemia protein FANCM to the Bloom syndrome dissolvasome. Proc. Natl. Acad. Sci. 109, 4437– 4442 (2012).

23. Bythell-Douglas, R. & Deans, A. J. A structural guide to the Bloom syndrome complex. Structure 29, 99–113 (2021).

24. Chaganti, R. S. K., Schonberg, S. & German, J. A manyfold increase in sister chromatid exchanges in Bloom’s syndrome lymphocytes. Proc. Natl. Acad. Sci. U. S. A. 71, 4508– 4512 (1974).

25. Wechsler, T., Newman, S. & West, S. C. Aberrant chromosome morphology in human cells defective for Holliday junction resolution. Nature 471, 642–646 (2011).

26. Rosin, M. P. & German, J. Evidence for chromosome instability in vivo in bloom syndrome: Increased numbers of micronuclei in exfoliated cells. Hum. Genet. 71, 187– 191 (1985).

27. Yankiwski, V., Marciniak, R. A., Guarente, L. & Neff, N. F. Nuclear structure in normal and Bloom syndrome cells. Proc. Natl. Acad. Sci. 97, 5214–5219 (2000).

28. Böhly, N., Kistner, M. & Bastians, H. Mild replication stress causes aneuploidy by deregulating microtubule dynamics in mitosis. Cell Cycle Georget. Tex 18, 2770–2783 (2019).

29. Böhly, N. et al. Increased replication origin firing links replication stress to whole chromosomal instability in human cancer. Cell Rep. 41, (2022).

30. Van Wietmarschen, N. et al. BLM helicase suppresses recombination at G-quadruplex motifs in transcribed genes. Nat. Commun. 9, 1–12 (2018).

31. Macheret, M. et al. High-resolution mapping of mitotic DNA synthesis regions and common fragile sites in the human genome through direct sequencing. Cell Res. 30, 997–1008 (2020).

32. Ji, F. et al. Genome-wide high-resolution mapping of mitotic DNA synthesis sites and common fragile sites by direct sequencing. Cell Res. 30, 1009–1023 (2020).

33. Gönenc, I. I. et al. Phenotypic spectrum of BLM- and RMI1–related Bloom syndrome. Clin. Genet. 101, 559–564 (2022).

34. Bizard, A. H. & Hickson, I. D. The dissolution of double Holliday junctions. Cold Spring Harb. Perspect. Biol. 6, a016477 (2014).

35. Bizard, A. H. & Hickson, I. D. Anaphase: a fortune-teller of genomic instability. Cell Nucl. 52, 112–119 (2018).

36. Cimini, D. & Degrassi, F. Aneuploidy: a matter of bad connections. Trends Cell Biol. 15, 442–451 (2005).

37. Bakhoum, S. F. et al. The mitotic origin of chromosomal instability. Curr. Biol. 24, R148– R149 (2014).

38. Chan, K. L., Palmai-Pallag, T., Ying, S. & Hickson, I. D. Replication stress induces sister-chromatid bridging at fragile site loci in mitosis. Nat. Cell Biol. 11, 753–760 (2009).

39. Chan, Y. W., Fugger, K. & West, S. C. Unresolved recombination intermediates lead to ultra-fine anaphase bridges, chromosome breaks and aberrations. Nat. Cell Biol. 20, 92– 103 (2018).

40. Chan, K. L., North, P. S. & Hickson, I. D. BLM is required for faithful chromosome segregation and its localization defines a class of ultrafine anaphase bridges. EMBO J. 26, 3397–3409 (2007).

41. Baumann, C., Körner, R., Hofmann, K. & Nigg, E. A. PICH, a centromere-associated SNF2 family ATPase, is regulated by Plk1 and required for the spindle checkpoint. Cell 128, 101–114 (2007).

42. Sarlós, K. et al. Reconstitution of anaphase DNA bridge recognition and disjunction. Nat. Struct. Mol. Biol. 25, 868–876 (2018).

43. Hamann, T. E. et al. Mitotic BLM functions are required to maintain genomic stability. Nucleic Acids Res. 54, gkag199 (2026).

44. Tsukada, K. et al. BLM and BRCA1-BARD1 coordinate complementary mechanisms of joint DNA molecule resolution. Mol. Cell 84, 640–658 (2024).

45. Regan, S. B. et al. Megabase-scale loss of heterozygosity provoked by CRISPR-Cas9 DNA double-strand breaks. Mol. Cell 85, 4119–4137 (2025).

46. Richardson, C. D. et al. CRISPR–Cas9 genome editing in human cells occurs via the Fanconi anemia pathway. Nat. Genet. 50, 1132–1139 (2018).

47. van Gent, D. C., Hoeijmakers, J. H. J. & Kanaar, R. Chromosomal stability and the DNA double-stranded break connection. Nat. Rev. Genet. 2, 196–206 (2001).

48. Johnson, R. D. & Jasin, M. Sister chromatid gene conversion is a prominent double-strand break repair pathway in mammalian cells. EMBO J. 19, 3398–3407 (2000).

49. Pikor, L., Thu, K., Vucic, E. & Lam, W. The detection and implication of genome instability in cancer. Cancer Metastasis Rev. 32, 341–352 (2013).

50. Bakker, B. et al. Single-cell sequencing reveals karyotype heterogeneity in murine and human malignancies. Genome Biol. 17, 115 (2016).

51. Shaikh, N. et al. Replication stress generates distinctive landscapes of DNA copy number alterations and chromosome scale losses. Genome Biol. 23, 223 (2022).

52. Sarni, D. & Kerem, B. The complex nature of fragile site plasticity and its importance in cancer. Cell Nucl. 40, 131–136 (2016).

53. Bhowmick, R. & Hickson, I. D. The” enemies within”: regions of the genome that are inherently difficult to replicate. F1000Research 6, (2017).

54. Ji, F. et al. New Era of Mapping and Understanding Common Fragile Sites: An Updated Review on Origin of Chromosome Fragility. Front. Genet. 13, 906957 (2022).

55. Minocherhomji, S. et al. Replication stress activates DNA repair synthesis in mitosis. Nature 528, 286–290 (2015).

56. Minocherhomji, S. & Hickson, I. D. Structure-specific endonucleases: guardians of fragile site stability. Trends Cell Biol. 24, 321–327 (2014).

57. Zeman, M. K. & Cimprich, K. A. Causes and consequences of replication stress. Nat. Cell Biol. 16, 2–9 (2014).

58. Vesela, E., Chroma, K., Turi, Z. & Mistrik, M. Common chemical inductors of replication stress: focus on cell-based studies. Biomolecules 7, 19 (2017).

59. Lukas, C. et al. 53BP1 nuclear bodies form around DNA lesions generated by mitotic transmission of chromosomes under replication stress. Nat. Cell Biol. 13, 243–253 (2011).

60. Glover, T. W., Berger, C., Coyle, J. & Echo, B. DNA polymerase α inhibition by aphidicolin induces gaps and breaks at common fragile sites in human chromosomes. Hum. Genet. 67, 136–142 (1984).

61. Richardson, C. D. et al. CRISPR–Cas9 genome editing in human cells occurs via the Fanconi anemia pathway. Nat. Genet. 50, 1132–1139 (2018).

62. Nicholls, T. J. et al. Topoisomerase 3α Is Required for Decatenation and Segregation of Human mtDNA. Mol. Cell 69, 9–23.e6 (2018).

63. Hangas, A. et al. Top3α is the replicative topoisomerase in mitochondrial DNA replication. Nucleic Acids Res. 50, 8733–8748 (2022).

64. Saha, L. K. & Pommier, Y. TOP3A coupling with replication forks and repair of TOP3A cleavage complexes. Cell Cycle 23, 115–130 (2024).

65. Chester, N., Kuo, F., Kozak, C., O’Hara, C. D. & Leder, P. Stage-specific apoptosis, developmental delay, and embryonic lethality in mice homozygous for a targeted disruption in the murine Bloom’s syndrome gene. Genes Dev. 12, 3382–3393 (1998).

66. Goss, K. H. et al. Enhanced tumor formation in mice heterozygous for Blm mutation. Science 297, 2051–2053 (2002).

67. McDaniel, L. D. et al. Chromosome instability and tumor predisposition inversely correlate with BLM protein levels. DNA Repair 2, 1387–1404 (2003).

68. Li, W. & Wang, J. C. Mammalian DNA topoisomerase IIIα is essential in early embryogenesis. Proc. Natl. Acad. Sci. 95, 1010–1013 (1998).

69. Wilson, T. E. et al. Large transcription units unify copy number variants and common fragile sites arising under replication stress. Genome Res. 25, 189–200 (2015).

70. Brison, O. et al. Transcription-mediated organization of the replication initiation program across large genes sets common fragile sites genome-wide. Nat. Commun. 10, 1–12 (2019).

71. Bailey, J. A., Baertsch, R., Kent, W. J., Haussler, D. & Eichler, E. E. Hotspots of mammalian chromosomal evolution. Genome Biol. 5, R23 (2004).

72. Wilson, T. E., Ahmed, S., Winningham, A. & Glover, T. W. Replication stress induces POLQ-mediated structural variant formation throughout common fragile sites after entry into mitosis. Nat. Commun. 15, 9582 (2024).

73. Rössler, U. et al. Efficient generation of osteoclasts from human induced pluripotent stem cells and functional investigations of lethal CLCN7-related osteopetrosis. J. Bone Miner. Res. 36, 1621–1635 (2021).

74. Hanses, U. et al. Intronic CRISPR Repair in a Preclinical Model of Noonan Syndrome-Associated Cardiomyopathy. Circulation 142, 1059–1076 (2020).

75. Concordet, J.-P. & Haeussler, M. CRISPOR: Intuitive guide selection for CRISPR/Cas9 genome editing experiments and screens. Nucleic Acids Res. 46, W242–W245 (2018).

76. Park, J., Lim, K., Kim, J.-S. & Bae, S. Cas-analyzer: An online tool for assessing genome editing results using NGS data. Bioinforma. Oxf. Engl. 33, 286–288 (2017).

77. Dehairs, J., Talebi, A., Cherifi, Y. & Swinnen, J. V. CRISP-ID: Decoding CRISPR mediated indels by Sanger sequencing. Sci. Rep. 6, 28973 (2016).

78. Gönenc, I. I. et al. Single-cell transcription profiles in Bloom syndrome patients link BLM deficiency with altered condensin complex expression signatures. Hum. Mol. Genet. 31, 2185–2193 (2022).

79. van den Bos, H., et al. Quantification of Aneuploidy in Mammalian Systems BT - Cellular Senescence: Methods and Protocols. in (ed. Demaria, M.) 159–190 (Springer New York, New York, NY, 2019). doi:10.1007/978-1-4939-8931-7_15.

80. Andrews, S. FastQC: A quality control tool for high throughput sequence data. (2010).

81. Wingett, S. W. & Andrews, S. FastQ Screen: A tool for multi-genome mapping and quality control. F1000Research 7, 1338–1338 (2018).

82. Ewels, P., Magnusson, M., Lundin, S. & Käller, M. MultiQC: summarize analysis results for multiple tools and samples in a single report. Bioinformatics 32, 3047–3048 (2016).

83. Bushnell, B., Rood, J. & Singer, E. BBMerge - Accurate paired shotgun read merging via overlap. PloS One 12, e0185056–e0185056 (2017).

84. Langmead, B. & Salzberg, S. L. Fast gapped-read alignment with Bowtie 2. Nat. Methods 9, 357–359 (2012).

85. Li, H. et al. The Sequence Alignment/Map format and SAMtools. Bioinformatics 25, 2078–2079 (2009).

86. Mirceta, M., Shum, N., Schmidt, M. H. M. & Pearson, C. E. Fragile sites, chromosomal lesions, tandem repeats, and disease. Front. Genet. 13, 1–45 (2022).

87. Kumar, R. et al. HumCFS: A database of fragile sites in human chromosomes. BMC Genomics 19, 1–8 (2019).

88. Georgakilas, A. G. et al. Are common fragile sites merely structural domains or highly organized “functional” units susceptible to oncogenic stress? Cell. Mol. Life Sci. 71, 4519–4544 (2014).

89. Debacker, K. & Frank Kooy, R. Fragile sites and human disease. Hum. Mol. Genet. 16, 150–158 (2007).

90. Kent, W. J. et al. The human genome browser at UCSC. Genome Res. 12, 996–1006 (2002).

91. Cui, Y. et al. BioCircos. js: an interactive Circos JavaScript library for biological data visualization on web applications. Bioinformatics 32, 1740–1742 (2016).

