## Supplementary Materials for "BTRR complex deficiency is a driver for genomic instability in Bloom syndrome"

### Supplementary Figures and Tables

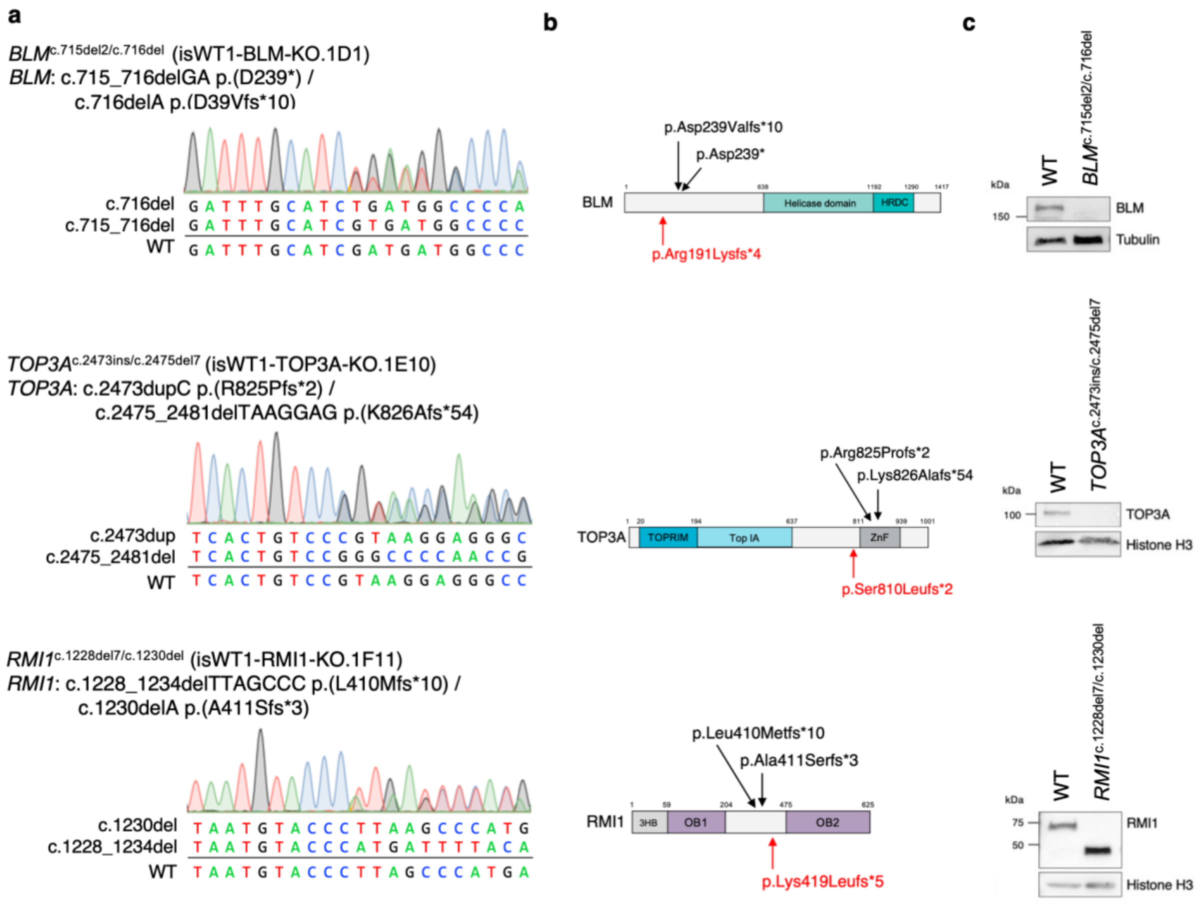


**Supplementary Fig. 1 Knockout (KO) confirmation of the CRISPR/Cas9 genome-edited iPSCs. a** Biallelic frameshift mutations in the corresponding genes were identified in iPSC lines via Sanger sequencing. **b** The positions of frameshift mutations with respect to the schematic representations of the corresponding proteins are marked in black, the positions of the corresponding mutations identified in BSyn and BSyn-like patients are marked in red. **c** Full length BLM, TOP3A, and RMI1 proteins were not detected in the cell lysates of KO iPSC lines of the corresponding genes by Western blot, while a putative truncated RMI1 variant was detected in the *RMI1*$^{c.1228del7/c.1230del}$ clone. Uncropped images are provided in the Supplementary Information.

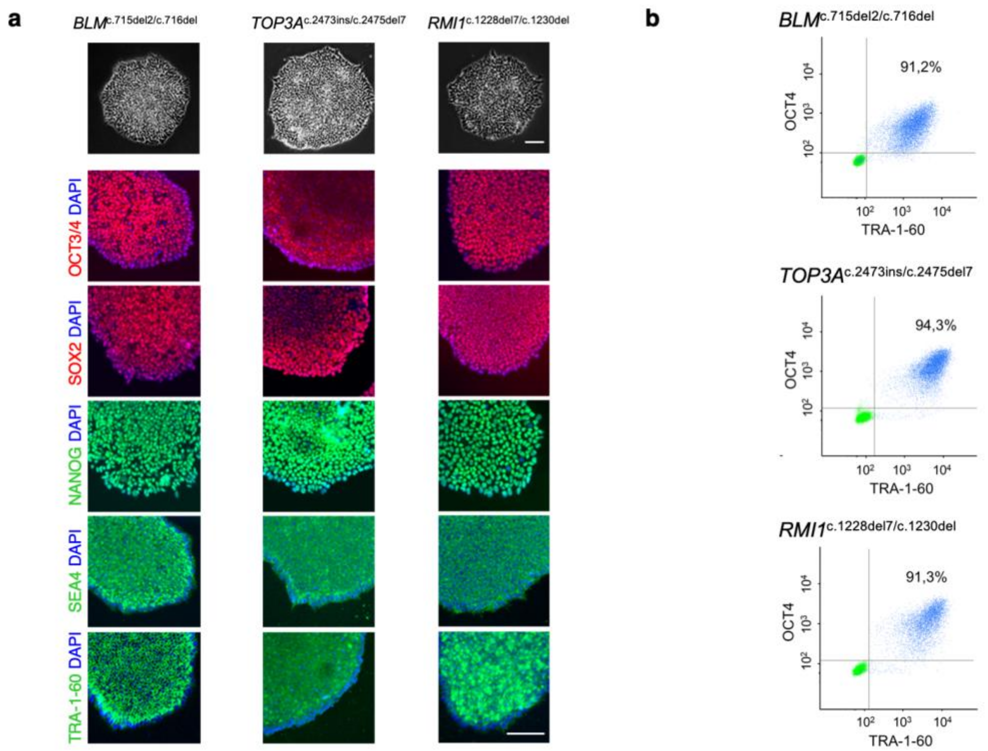


**Supplementary Fig. 2 Pluripotency characterization of the CRISPR/Cas9 genome-edited iPSCs. a** Generated BTRR$^{KO}$ iPSC lines had a typical human stem cell-like colony morphology. All three BTRR$^{KO}$ iPSC lines expressed the major pluripotency markers (SOX2, OCT3/4, in red; TRA-1-60, SSEA4, NANOG, in green). DNA was counter-stained with DAPI. Scale bar: 100 µm **b** Flow cytometry analysis of pluripotency markers OCT4 and TRA-1-60 revealed more than 90% of cells (blue dots) were pluripotent in BTRR$^{KO}$ iPSC lines. Green points represent the negative controls.

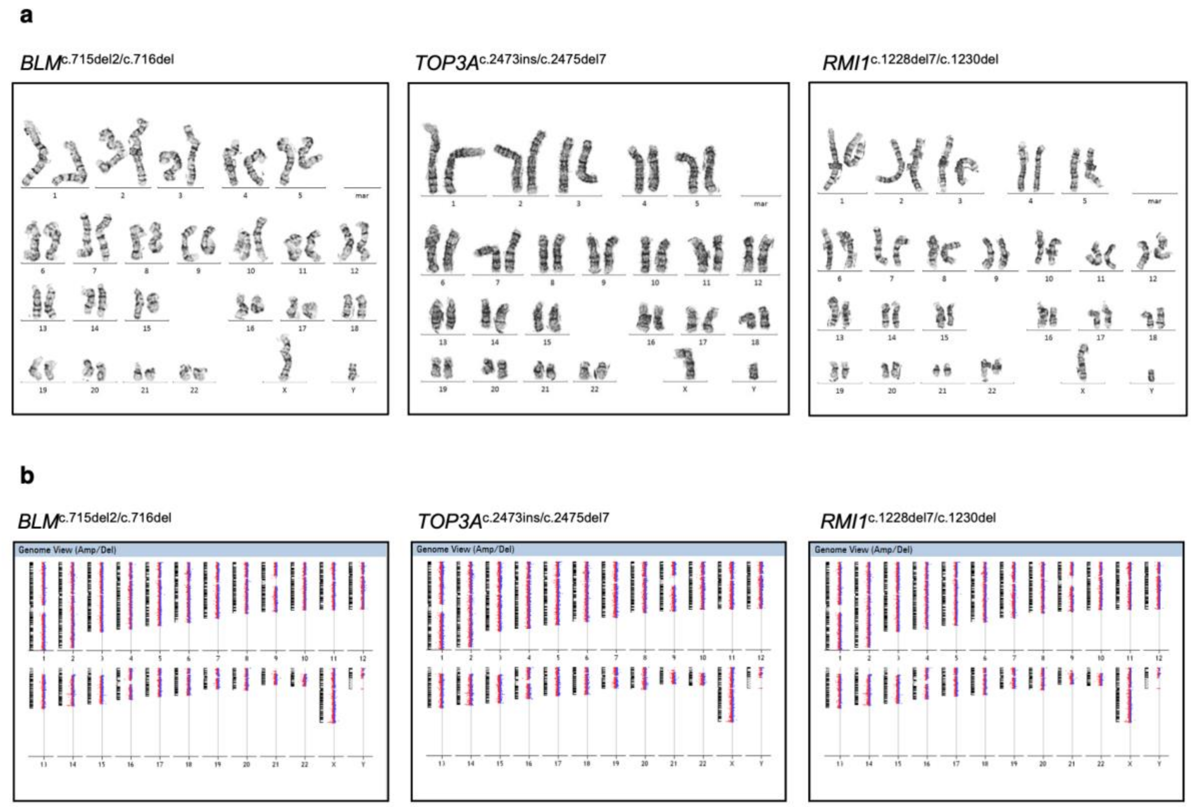


**Supplementary Fig. 3 Metaphase karyotype and structural variant analyses of KO iPSCs. a** Representative karyograms of each $BTRR^{KO}$ iPSC line are shown revealing normal 46, XY karyotype for all three cell lines. For WT 12, for *BLM*$^{c.715del2/c.716del}$ 6, and for *RMI1*$^{c.1228del7/c.1230del}$ 13 cells were karyotyped. For *TOP3A*$^{c.2473ins/c.2475del7}$, 13 cells were karyotyped; 12 showed a normal karyotype, and one cell exhibited loss of the Y chromosome. **b** Array CGH analysis showed normal results for the three $BTRR^{KO}$ iPSC lines i.e., no pathogenic structural variant was identified in the bulk samples.

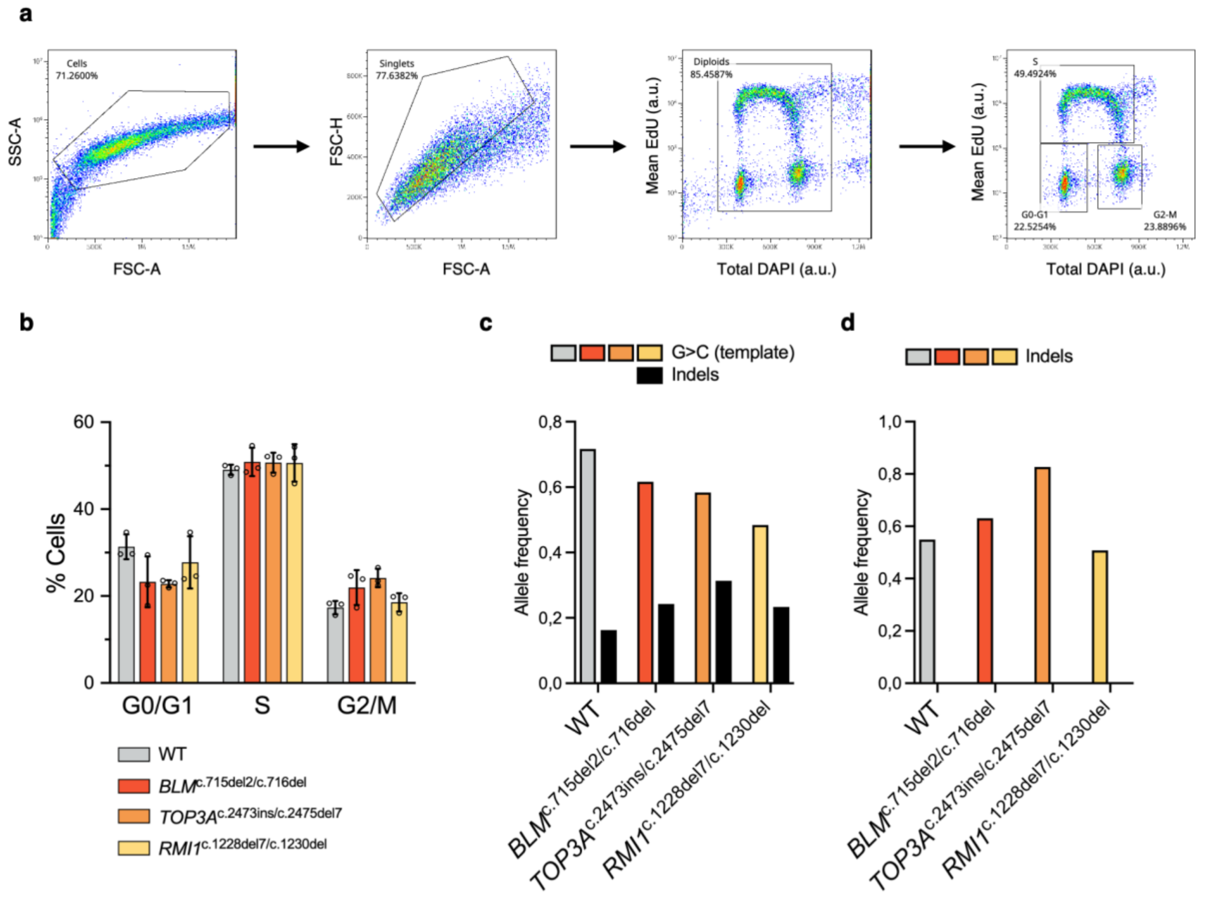


**Supplementary Fig. 4 Cell cycle profiling and analyses of DNA repair outcomes in BTRR$^{KO}$ iPSCs. a** Gating strategy for cell cycle profiling with flow cytometry. **b** The percentage of cells in the different cell cycle phases in wild-type and BTRR$^{KO}$ iPSCs. Error bars indicate mean with S.E.M from 3 independent flow cytometry experiments using samples from different passage numbers. **c** Frequencies of alleles generated by single-stranded template repair (G>C associated template) or NHEJ (indels) after Cas9-induced DSB identified with NGS-based deep amplicon sequencing with approximately 24 000x coverage of the targeted site. **d** Frequencies of NHEJ-mediated alleles after Cas9-induced DSB without single-stranded template provided, analyzed as in (c).

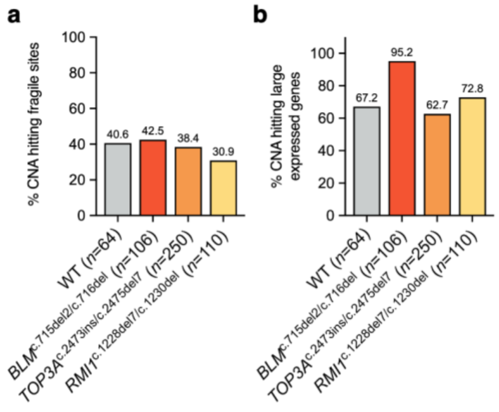


**Supplementary Fig. 5 Percentages of CNAs hitting certain genomic regions. a** CNA percentages hitting MiDAS-seq-defined fragile sites. *n* denotes total number of identified CNAs. **b** CNA percentages hitting expressed large (>300 kb) genes identified in the iPSC model by single-cell transcriptome sequencing. A complete list of expressed genes is provided as Source data. Overlapping criteria were at least one base pair overlap among the denoted regions.

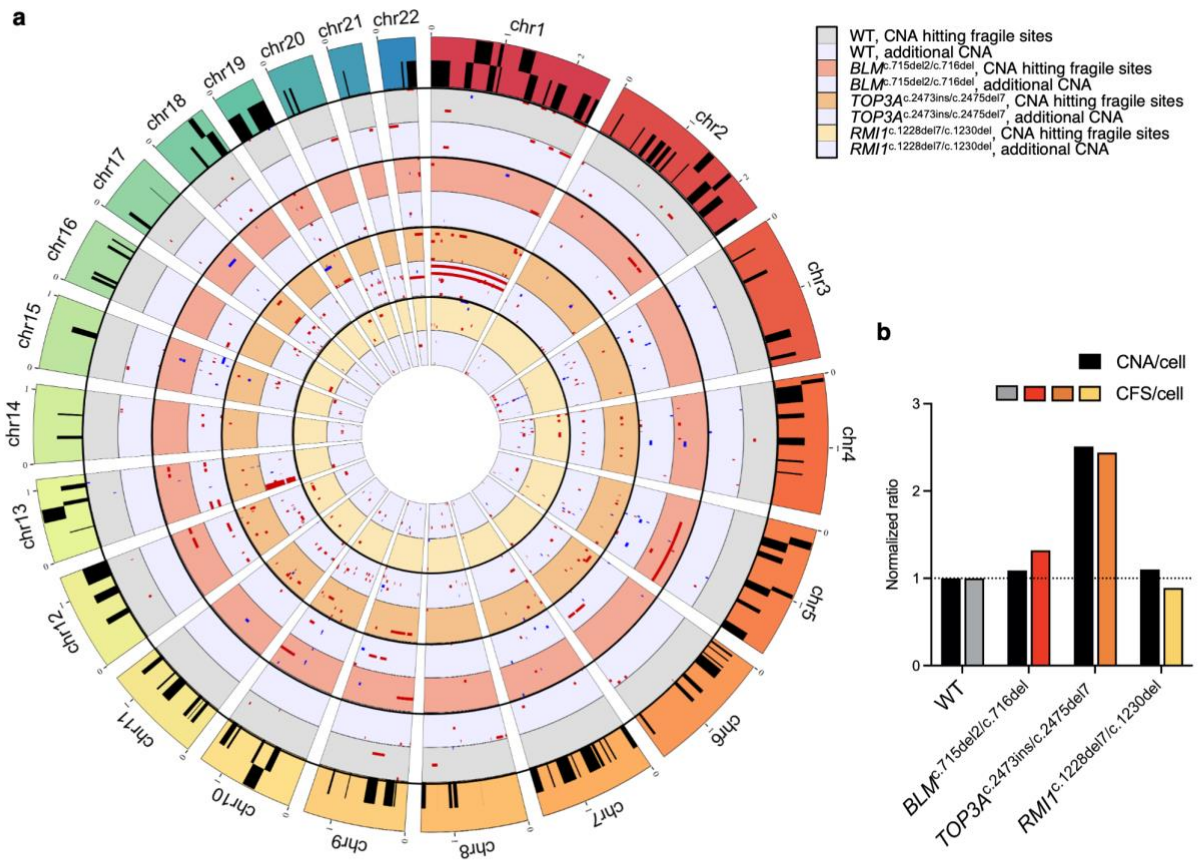


**Supplementary Fig. 6 CNAs associated with common fragile sites (CFS) previously identified by cytogenetics. a** Circos plot shows CNAs hitting fragile sites (gray, red, orange, and yellow-colored tracks for different samples) and the additional CNAs (light blue-colored tracks). The criteria for overlaps were at least 10 % length of the CNA to correspond with a fragile site. Gains (blue) and losses (red) are shown in single cells based on genomic location and size. Black boxes on the outer reference track indicate the CFS regions (two tracks for overlayed CFS) based on a custom-made database (Supplementary Table 5). **b** The relation of CNA per cell to CFS per cell shows increased CFS affected by CNAs. Ratios were normalized to WT for both values. Whole chromosome aneuploidies were excluded from the CFS analysis.

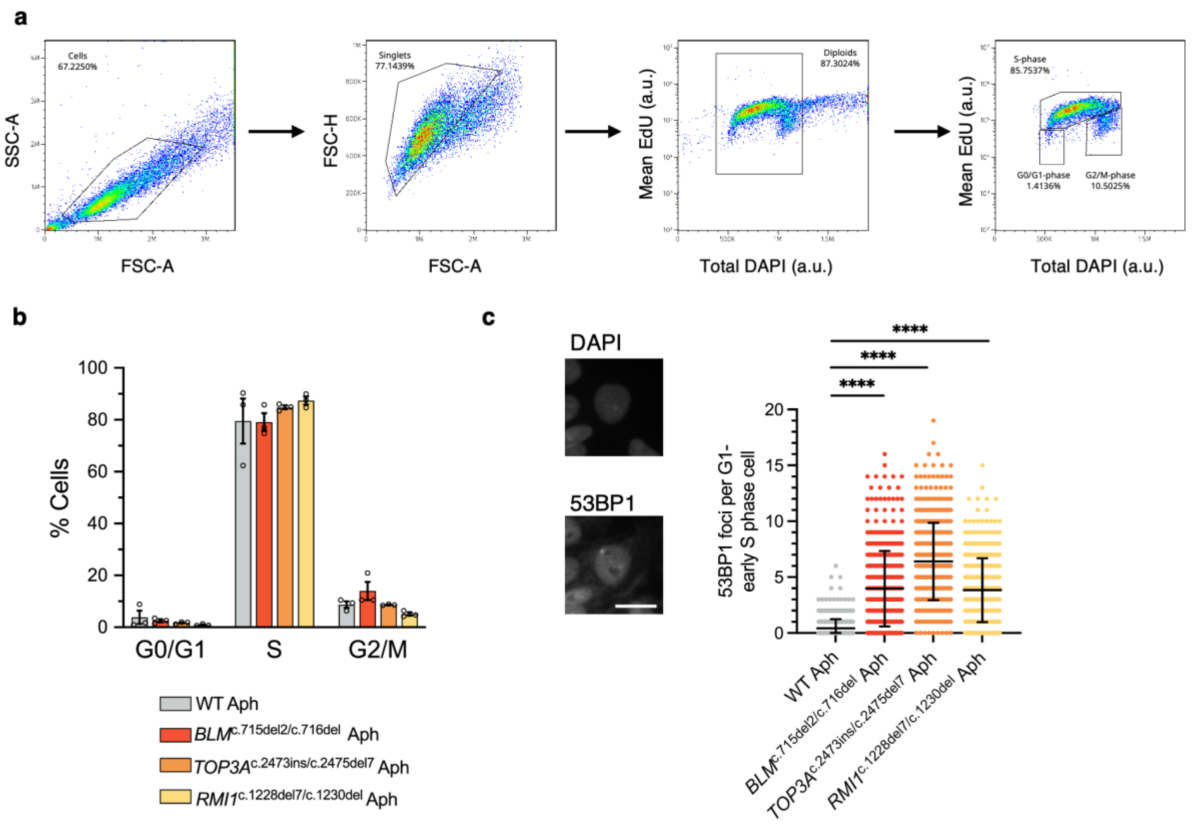


**Supplementary Fig. 7 Assessment of the effects of aphidicolin in the iPSCs model. a** Gating strategy for the cell cycle profiling evaluated by EdU incorporation and DAPI signals. **b** Analyses of percentages of cells in each cell cycle stage revealed a high proportion of S-phase cells after aphidicolin (300 nM, 24h) treatment with no major differences among wild-type and knockout cell lines. **c** Analyses of DNA damage identified by 53BP1 foci count in G1 cells after aphidicolin treatment revealed slightly increased DNA damage in G1-early S phase cells of BTRR-complex-deficient cell lines. Representative image of nuclei (DAPI) and 53BP1 foci in WT cells (left panel) and quantification of identified foci by quantitative image-based cytometry (right panel). Scale bar 10 µm. 500 nuclei were analyzed for three experiments and data from one repeat is plotted. Error bars indicate mean with standard deviation. Kruskal-Wallis test with Benjamini-Hochberg correction was performed. ****p<0.0001

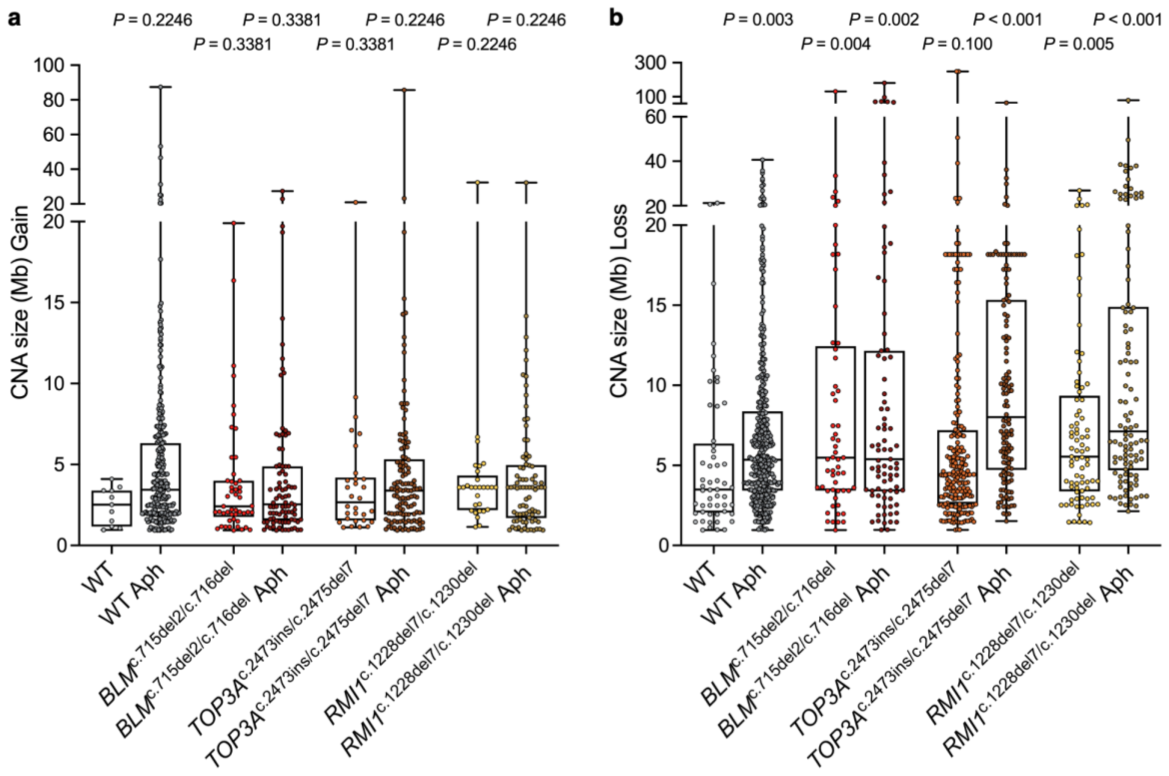


**Supplementary Fig. 8 Size distribution of CNAs identified in all samples including replication-stress induced samples.** Larger CNAs were present in BTRR complex deficient-iPSC and replication stress-induced cells including the wild type for both **a** gains and **b** losses (right). Kruskal-Wallis test with Benjamini-Hochberg correction was performed and pairwise comparisons were performed against untreated WT cells.

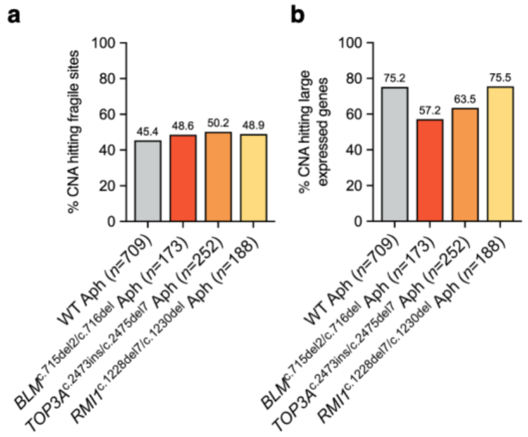


**Supplementary Fig. 9 Percentages of replication-stress-induced CNAs hitting certain genomic regions. a** Percentage of CNAs hitting MiDASseq-defined fragile sites after aphidicolin treatment (300 nM, 24h). n denotes total number of identified CNAs. **b** CNA percentages hitting expressed large (>300 kb) genes identified in the iPSC model. Overlapping criteria were at least one base pair overlap among the denoted regions.

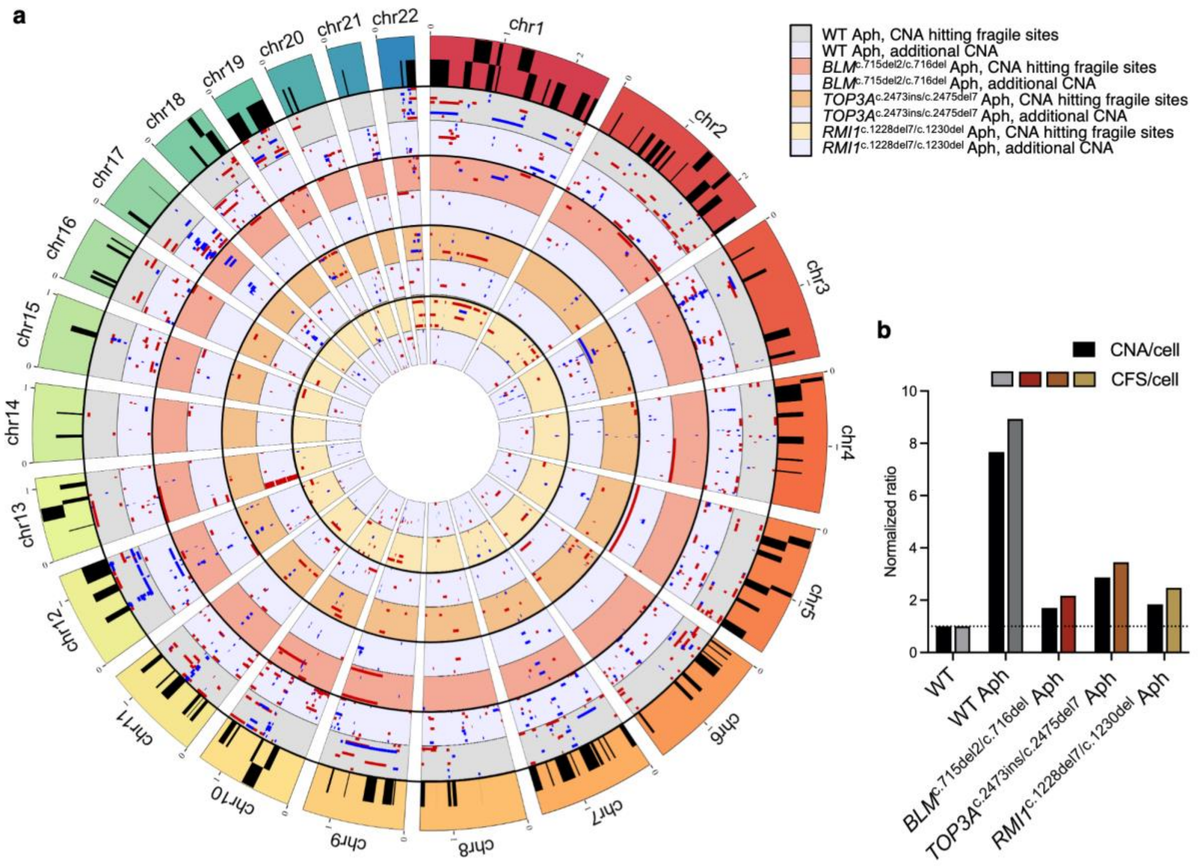


**Supplementary Fig. 10 Moderate replication stress-induced CNAs associated with CFS identified by cytogenetics. a** Circos plot indicates CNA hitting fragile sites (gray, red, orange, and yellow-colored tracks for different samples) and additional CNA (light blue-colored tracks) in the aphidicolin (Aph)-treated samples. The criteria for overlaps were at least 10 % length of CNA to correspond with a fragile site. Gains (blue) and losses (red) are shown in single cells. Black boxes on the outer track indicate the CFS regions. **b** The relation of CNA per cell to CFS per cell shows increased CFS affected by CNAs in the aphidicolin-treated samples. Aphidicolin-treated samples were normalized to non-treated WT for both ratios. Whole chromosome aneuploidies were excluded from the CFS analysis.

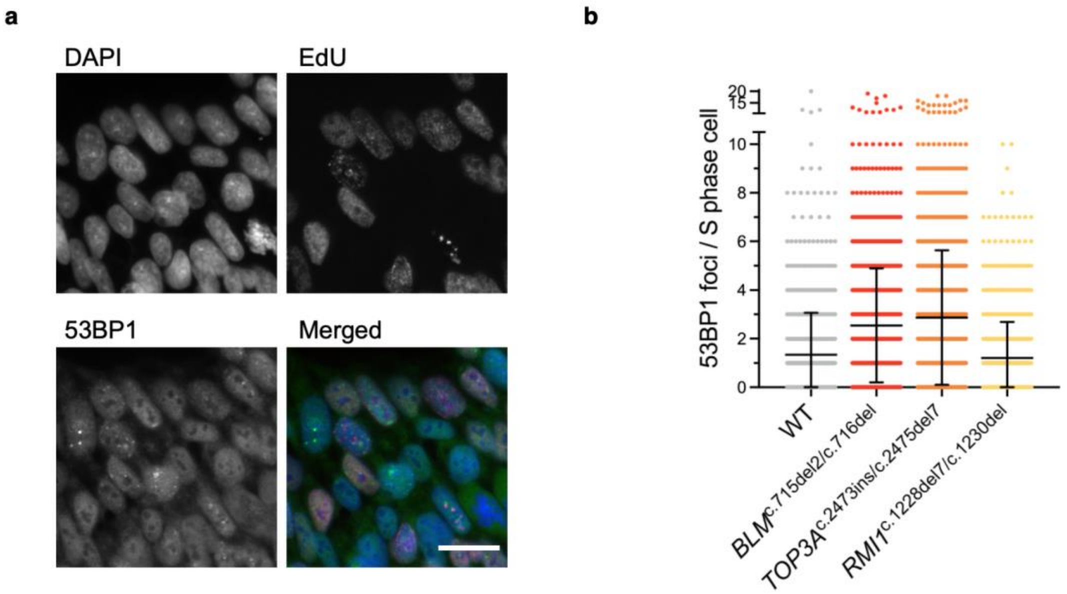


**Supplementary Fig. 11 DNA damage levels in S-phase cells. a** Representative images of S phase WT cells identified by EdU incorporation and analyzed for 53BP1 foci as a DNA damage marker. Scale bar 20 µm. **b** Quantification of 53BP1 foci in EdU-positive nuclei. Cells were segmented using the DAPI signal, and EdU-positive nuclei were selected prior to automated 53BP1 foci count in ImageJ. 1000 nuclei were analyzed for each repeat (*n*=3), data from one representative experiment are plotted. Error bars indicate mean with standard deviation.

### Supplementary Information

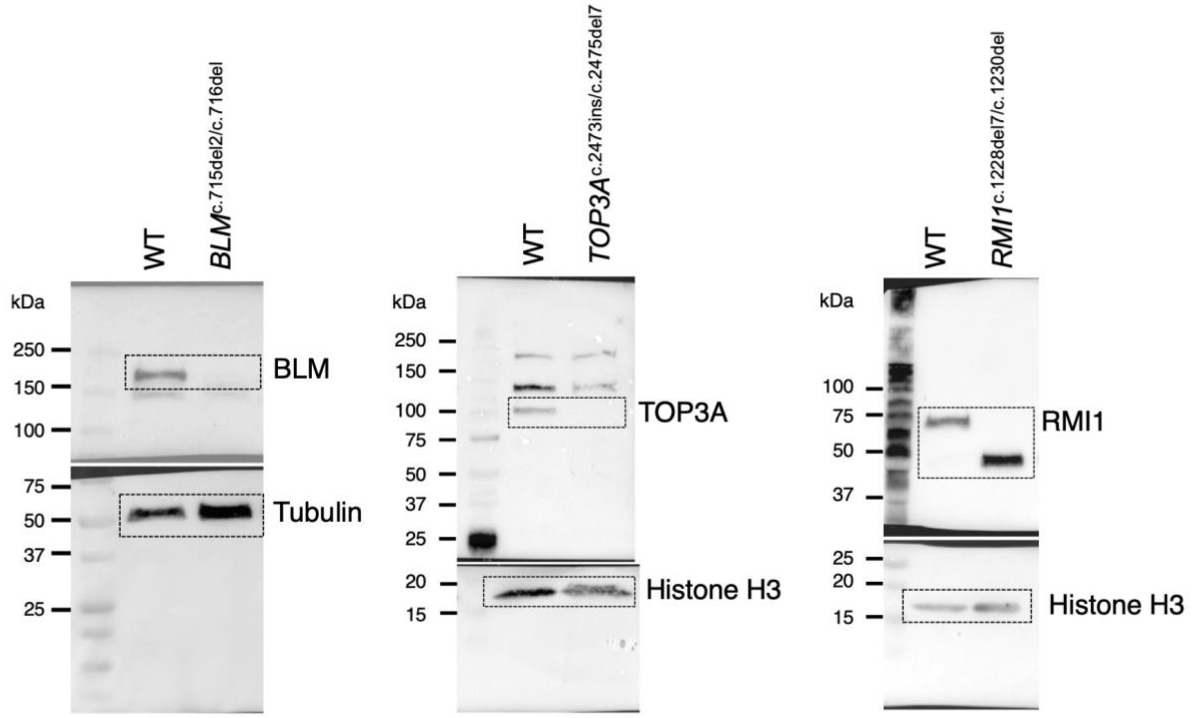


**Supplementary Data 1** Uncropped Western blots for evaluating full length BLM, TOP3A and RMI1 in the wild type (WT) and the CRISPR/Cas9 genome-edited knockout clones. The antibodies with the working dilutions were as the follows: anti-BLM (Abcam, ab2179), 1:1000; anti-TOP3A (Proteintech, 14525-1-AP), 1:2000; anti-RMI1 (Proteintech, 14630-1-AP), 1:1000, anti-Tubulin (Abcam, ab52866), 1:5000; anti-Histone-H3 (Abcam, ab8505), 1:1000.

### Supplementary Tables

**Supplementary Table 1. Reference BSyn patient mutations**

| Gene | Homozygous mutation | Reference |
|---|---|---|
| *BLM* (NM_000057.4) | c.572_573del;p.Arg191Lysfs*4 | Gönenc et al. [1] |
| *TOP3A* (NM_004618.5) | c.2428del;p.Ser810Leufs*2 | Martin et al. [2] |
| *RMI1* (NM_024945.2) | c.1255_1259del;p.Lys419Leufs*5 | Martin et al. [2] |

**Supplementary Table 2. Reference common fragile sites (CFS) determined previously by MiDASseq in U2-OS cells.** Coordinates are based on hg38 assembly. Regions were merged from Macheret et al.[3], and Ji et al.[4]. Chr, chromosome.

| Chr | Start | End |
|---|---|---|
| chr1 | 13853505 | 14800000 |
| chr1 | 49000000 | 50104328 |
| chr1 | 57094327 | 58100000 |
| chr1 | 71264317 | 72874317 |
| chr1 | 80504315 | 81124315 |
| chr1 | 97124444 | 97784444 |
| chr1 | 174320862 | 174940863 |
| chr1 | 176200000 | 177020864 |
| chr1 | 187080868 | 187660868 |
| chr1 | 192210870 | 192670870 |
| chr1 | 192920870 | 193460870 |
| chr1 | 237236700 | 237636700 |
| chr1 | 245696698 | 246356698 |
| chr2 | 13349875 | 15689876 |
| chr2 | 18638734 | 19180239 |
| chr2 | 21317128 | 21937128 |
| chr2 | 32600000 | 33000000 |
| chr2 | 53742863 | 54102863 |
| chr2 | 63100000 | 63992866 |
| chr2 | 79612874 | 80112874 |
| chr2 | 81452876 | 81932876 |
| chr2 | 115122423 | 115842424 |
| chr2 | 116500000 | 117682424 |
| chr2 | 121872424 | 122812424 |
| chr2 | 132412427 | 133862429 |
| chr2 | 135182430 | 135722430 |
| chr2 | 136722430 | 137712430 |
| chr2 | 138102430 | 138500000 |
| chr2 | 139252430 | 140000000 |
| chr2 | 140300000 | 140400000 |
| chr2 | 140700000 | 140900000 |

| | | |
|---|---|---|
| chr2 | 141482431 | 142042431 |
| chr2 | 147892431 | 148432431 |
| chr2 | 153600000 | 154453488 |
| chr2 | 157800000 | 158300000 |
| chr2 | 159400000 | 159700000 |
| chr2 | 167253490 | 168183490 |
| chr2 | 183245272 | 183765273 |
| chr2 | 185445273 | 186565273 |
| chr2 | 204625277 | 205535276 |
| chr2 | 211700000 | 212465276 |
| chr2 | 213455276 | 214500000 |
| chr2 | 220100000 | 220765280 |
| chr2 | 235661356 | 236041356 |
| chr3 | 1998316 | 3068316 |
| chr3 | 3268316 | 4428316 |
| chr3 | 6448313 | 7678313 |
| chr3 | 8200000 | 8700000 |
| chr3 | 17038508 | 17758508 |
| chr3 | 19088508 | 19748508 |
| chr3 | 20028508 | 20888508 |
| chr3 | 24898509 | 25598509 |
| chr3 | 29318509 | 29998509 |
| chr3 | 41318509 | 41778508 |
| chr3 | 53885973 | 55145973 |
| chr3 | 55345972 | 56525972 |
| chr3 | 59644274 | 61374326 |
| chr3 | 61474326 | 62434325 |
| chr3 | 65264325 | 65964325 |
| chr3 | 67039576 | 67839576 |
| chr3 | 71030849 | 71590849 |
| chr3 | 74170849 | 74670849 |
| chr3 | 75820849 | 76660849 |
| chr3 | 77100000 | 77200000 |
| chr3 | 78560850 | 79650850 |
| chr3 | 81180849 | 81740849 |
| chr3 | 96721156 | 97961156 |
| chr3 | 104271156 | 104791156 |
| chr3 | 109501153 | 110121153 |
| chr3 | 114391153 | 115091153 |
| chr3 | 115821153 | 116381153 |
| chr3 | 117300000 | 117700000 |
| chr3 | 121041153 | 121461153 |
| chr3 | 130941156 | 131700000 |
| chr3 | 158052211 | 158872211 |
| chr3 | 168382212 | 168782212 |
| chr3 | 169152212 | 169532212 |

| | | |
|---|---|---|
| chr3 | 173472210 | 174252210 |
| chr3 | 174552210 | 175712212 |
| chr3 | 188252212 | 189300000 |
| chr3 | 192182211 | 192782211 |
| chr4 | 11300000 | 11400000 |
| chr4 | 12600000 | 13458376 |
| chr4 | 18808377 | 19988377 |
| chr4 | 21300000 | 21600000 |
| chr4 | 32100000 | 32200000 |
| chr4 | 32900000 | 34600000 |
| chr4 | 35658378 | 36498378 |
| chr4 | 36900000 | 37000000 |
| chr4 | 61194282 | 62004282 |
| chr4 | 84800000 | 84900000 |
| chr4 | 85700000 | 86100000 |
| chr4 | 86400000 | 86900000 |
| chr4 | 88800000 | 88900000 |
| chr4 | 92438849 | 93748849 |
| chr4 | 94538849 | 95408849 |
| chr4 | 97138849 | 98200000 |
| chr4 | 112818844 | 113598844 |
| chr4 | 142088847 | 142748847 |
| chr4 | 143448847 | 144028847 |
| chr4 | 144538848 | 145098848 |
| chr4 | 147868849 | 148428848 |
| chr4 | 150300000 | 150848848 |
| chr4 | 163578848 | 164278848 |
| chr5 | 7439887 | 7919887 |
| chr5 | 10989888 | 11929888 |
| chr5 | 12049888 | 12959888 |
| chr5 | 15519891 | 15979891 |
| chr5 | 20889891 | 22629891 |
| chr5 | 23859891 | 24299891 |
| chr5 | 28369893 | 29499893 |
| chr5 | 59144173 | 59600000 |
| chr5 | 59800000 | 59900000 |
| chr5 | 60600000 | 61200000 |
| chr5 | 62800000 | 63000000 |
| chr5 | 65200000 | 65300000 |
| chr5 | 65500000 | 66100000 |
| chr5 | 81000000 | 81300000 |
| chr5 | 82924181 | 83344181 |
| chr5 | 83934181 | 84534182 |
| chr5 | 93424294 | 94100000 |
| chr5 | 94454295 | 95354296 |
| chr5 | 104124299 | 105354299 |
| chr5 | 107134299 | 107534299 |
| chr5 | 108864299 | 109244299 |

| | | |
|---|---|---|
| chr5 | 116184303 | 116564304 |
| chr5 | 119884305 | 120774305 |
| chr5 | 129400000 | 130400000 |
| chr5 | 136900000 | 137400000 |
| chr5 | 145200000 | 145400000 |
| chr5 | 162692994 | 163532994 |
| chr5 | 167900000 | 168100000 |
| chr6 | 1609765 | 2309766 |
| chr6 | 4829766 | 5839767 |
| chr6 | 20589769 | 21249769 |
| chr6 | 37932224 | 38672224 |
| chr6 | 61590095 | 62370095 |
| chr6 | 68650108 | 69310108 |
| chr6 | 76780283 | 77300283 |
| chr6 | 77770283 | 78530283 |
| chr6 | 82730283 | 83350281 |
| chr6 | 92030282 | 92670282 |
| chr6 | 98300000 | 98792124 |
| chr6 | 101442124 | 102022125 |
| chr6 | 103900000 | 104822125 |
| chr6 | 114048836 | 114918836 |
| chr6 | 120008854 | 121228854 |
| chr6 | 125898854 | 126838855 |
| chr6 | 128018855 | 128378855 |
| chr6 | 128988855 | 129548855 |
| chr6 | 141498863 | 141898863 |
| chr6 | 147668864 | 148200000 |
| chr6 | 161800000 | 162800000 |
| chr7 | 3360368 | 4280368 |
| chr7 | 7550369 | 8170370 |
| chr7 | 11300000 | 11700000 |
| chr7 | 13150375 | 13710375 |
| chr7 | 14210375 | 14810375 |
| chr7 | 15820375 | 16360375 |
| chr7 | 18090377 | 18910377 |
| chr7 | 31620386 | 32240388 |
| chr7 | 33310388 | 33800000 |
| chr7 | 36920395 | 37380397 |
| chr7 | 40010401 | 40940401 |
| chr7 | 69725014 | 70805014 |
| chr7 | 78000000 | 79410684 |
| chr7 | 90610686 | 91390685 |
| chr7 | 109900000 | 111579944 |
| chr7 | 111849944 | 112149945 |
| chr7 | 119399946 | 119879946 |
| chr7 | 122379946 | 122819946 |
| chr7 | 126409946 | 127169946 |
| chr7 | 133315246 | 134235248 |

| | | |
|---|---|---|
| chr7 | 136665253 | 137300000 |
| chr7 | 141500000 | 141800000 |
| chr8 | 2612910 | 3112478 |
| chr8 | 33732482 | 34392482 |
| chr8 | 35332482 | 35692482 |
| chr8 | 47367432 | 47847439 |
| chr8 | 51347440 | 51887440 |
| chr8 | 62257441 | 62797441 |
| chr8 | 63587442 | 64167443 |
| chr8 | 70657765 | 71297765 |
| chr8 | 71907765 | 72407765 |
| chr8 | 87947772 | 88647771 |
| chr8 | 88877771 | 90047772 |
| chr8 | 99017772 | 99817772 |
| chr8 | 103600000 | 104000000 |
| chr8 | 105407772 | 105927772 |
| chr8 | 108867771 | 109307771 |
| chr8 | 109867771 | 110717771 |
| chr8 | 112227771 | 112647771 |
| chr8 | 136587757 | 137147757 |
| chr8 | 139747757 | 140329901 |
| chr9 | 2030000 | 2790000 |
| chr9 | 9110000 | 10650000 |
| chr9 | 16000000 | 16400000 |
| chr9 | 17250002 | 17790002 |
| chr9 | 23120001 | 24230002 |
| chr9 | 27000000 | 27100000 |
| chr9 | 27900000 | 29530002 |
| chr9 | 31870002 | 32470002 |
| chr9 | 73555084 | 73955084 |
| chr9 | 78345084 | 79085084 |
| chr9 | 83195085 | 83915085 |
| chr9 | 105667719 | 106727719 |
| chr9 | 116227721 | 117337721 |
| chr9 | 119500000 | 120200000 |
| chr10 | 9200000 | 9828037 |
| chr10 | 18181071 | 18681071 |
| chr10 | 19811071 | 20351071 |
| chr10 | 34261072 | 34581072 |
| chr10 | 50940240 | 52320240 |
| chr10 | 54100000 | 54500000 |
| chr10 | 55200000 | 56210239 |
| chr10 | 57190240 | 57810240 |
| chr10 | 64360240 | 64800243 |
| chr10 | 66300000 | 66700000 |
| chr10 | 66900000 | 67400000 |
| chr10 | 75800000 | 76300000 |
| chr10 | 81900244 | 82970244 |

| | | |
|---|---|---|
| chr10 | 85600243 | 86220243 |
| chr10 | 106620242 | 107120242 |
| chr10 | 107700000 | 108500000 |
| chr10 | 112530241 | 113100000 |
| chr10 | 115500000 | 115800000 |
| chr10 | 127021736 | 127361736 |
| chr11 | 16108454 | 16468453 |
| chr11 | 20748454 | 21528454 |
| chr11 | 24598454 | 24998454 |
| chr11 | 28058453 | 28700000 |
| chr11 | 29000000 | 29100000 |
| chr11 | 30888453 | 31718452 |
| chr11 | 38498450 | 39158450 |
| chr11 | 78838955 | 79338955 |
| chr11 | 79600000 | 79700000 |
| chr11 | 83478957 | 84378957 |
| chr11 | 84700000 | 85500000 |
| chr11 | 87600000 | 87900000 |
| chr11 | 88546832 | 89436832 |
| chr11 | 90226832 | 91006832 |
| chr11 | 92326834 | 92766834 |
| chr11 | 99109270 | 99529269 |
| chr11 | 106900000 | 107100000 |
| chr11 | 116200000 | 116400000 |
| chr11 | 126270105 | 127010105 |
| chr11 | 131800000 | 131900000 |
| chr12 | 19600000 | 19700000 |
| chr12 | 20700000 | 21300000 |
| chr12 | 23637066 | 24407066 |
| chr12 | 43856197 | 44396217 |
| chr12 | 59736219 | 60136219 |
| chr12 | 66426220 | 66926220 |
| chr12 | 77546220 | 78186220 |
| chr12 | 78896220 | 79476220 |
| chr12 | 81186221 | 81686221 |
| chr12 | 82696221 | 83196221 |
| chr12 | 85916222 | 87456223 |
| chr12 | 99000000 | 100326222 |
| chr12 | 102016222 | 103026222 |
| chr13 | 25535862 | 25915862 |
| chr13 | 35035863 | 35555863 |
| chr13 | 56185866 | 56725866 |
| chr13 | 57275866 | 57975866 |
| chr13 | 58325866 | 58765866 |
| chr13 | 63495867 | 64385868 |
| chr13 | 66255868 | 67235868 |
| chr13 | 67925868 | 68725868 |
| chr13 | 80800000 | 81500000 |

| | | |
|---|---|---|
| chr13 | 88587745 | 89107746 |
| chr13 | 91427746 | 92787747 |
| chr13 | 93347747 | 94337746 |
| chr13 | 107177652 | 107967652 |
| chr14 | 26680794 | 27840794 |
| chr14 | 29800000 | 30000000 |
| chr14 | 32100000 | 32750794 |
| chr14 | 37000000 | 37930795 |
| chr14 | 39700000 | 40100000 |
| chr14 | 40520796 | 41500000 |
| chr14 | 42810797 | 44780797 |
| chr14 | 44800000 | 46000000 |
| chr14 | 46400000 | 48000797 |
| chr14 | 48800000 | 48900000 |
| chr14 | 49100000 | 49800000 |
| chr14 | 65400000 | 65700000 |
| chr14 | 66300000 | 67400000 |
| chr14 | 67800000 | 68500000 |
| chr14 | 78233657 | 80883656 |
| chr14 | 82000000 | 82400000 |
| chr14 | 83600000 | 83700000 |
| chr15 | 26984853 | 27384854 |
| chr15 | 60477801 | 61197801 |
| chr15 | 97046770 | 97486770 |
| chr16 | 7700000 | 8309998 |
| chr16 | 46300000 | 46400000 |
| chr16 | 47196089 | 47736089 |
| chr16 | 53716088 | 54016088 |
| chr16 | 63046096 | 63546096 |
| chr16 | 72046101 | 73116101 |
| chr16 | 76076102 | 76756103 |
| chr16 | 78006103 | 79100000 |
| chr16 | 82596395 | 83816395 |
| chr18 | 3820000 | 4500000 |
| chr18 | 7470002 | 8410002 |
| chr18 | 36600000 | 37200000 |
| chr18 | 39100000 | 39730036 |
| chr18 | 52473630 | 54133630 |
| chr18 | 69382764 | 70142764 |
| chr20 | 8169353 | 8869353 |
| chr20 | 12939352 | 13479353 |
| chr20 | 14289354 | 15949355 |
| chr20 | 61244944 | 61944944 |
| chr21 | 15967680 | 16497680 |
| chr21 | 20987682 | 21507679 |
| chr21 | 27327681 | 28500000 |
| chr21 | 30900000 | 31547687 |
| chr22 | 24700000 | 24800000 |

| | | |
|---|---|---|
| chr22 | 25500000 | 25900000 |
| chr22 | 27900000 | 28600000 |
| chr22 | 33264014 | 33864012 |
| chr22 | 35100000 | 35300000 |
| chrX | 6621959 | 7361959 |
| chrX | 7701959 | 8311959 |
| chrX | 10621960 | 11531880 |
| chrX | 26831883 | 27411883 |
| chrX | 28661883 | 29201883 |
| chrX | 55713567 | 56393567 |
| chrX | 57100000 | 57200000 |
| chrX | 63250123 | 63770120 |
| chrX | 64860120 | 65480120 |
| chrX | 67670158 | 68410158 |
| chrX | 75130165 | 75750165 |
| chrX | 81124501 | 81504501 |
| chrX | 86234997 | 86774997 |
| chrX | 91175001 | 92665001 |
| chrX | 94205001 | 94725001 |
| chrX | 95455001 | 96445001 |
| chrX | 96715001 | 98000000 |
| chrX | 99415002 | 99935002 |
| chrX | 121116146 | 122096147 |
| chrX | 124516150 | 125156151 |
| chrX | 125656002 | 126486017 |
| chrX | 127286017 | 127786019 |
| chrX | 128300000 | 129486023 |
| chrX | 138617839 | 139097838 |
| chrX | 141902214 | 142722214 |
| chrX | 148258480 | 148898470 |
| chr17 | 58700000 | 59000000 |
| chr17 | 59200000 | 59300000 |
| chr17 | 60600000 | 61300000 |

**Supplementary Table 3. Reference fragile sites (FS) determined previously by cytogenetic techniques.** Band coordinates were confirmed from UCSC GRCh38 (hg38) genome database.

| **Chr #** | **Name** | **Band** | **Start** | **End** | **Reference 1** | **Reference 2** |
|---|---|---|---|---|---|---|
| chr1 | FRA1A | 1p36 | 1 | 27.600.000 | Kumar et al.[1] | |
| chr1 | FRA1B | 1p32 | 50.200.001 | 60.800.000 | Kumar et al.[1] | |
| chr1 | FRA1D | 1p22 | 84.400.001 | 94.300.000 | Kumar et al.[1] | Georgakilas et al.[2] |
| chr1 | FRA1E | 1p21.2 | 99.300.001 | 101.800.000 | Kumar et al.[1] | Georgakilas et al.[2] |
| chr1 | FRA1F | 1q21 | 143.200.001 | 155.100.000 | Kumar et al.[1] | Georgakilas et al.[2] |
| chr1 | FRA1G | 1q25.1 | 173.000.001 | 176.100.000 | Kumar et al.[1] | Georgakilas et al.[2] |
| chr1 | FRA1H | 1q42 | 223.900.001 | 236.400.000 | Kumar et al.[1] | Georgakilas et al.[2] |
| chr1 | FRA1I | 1q44 | 243.500.001 | 248.956.422 | Kumar et al.[1] | Georgakilas et al.[2] |
| chr1 | FRA1J | 1q12 | 125.100.001 | 143.200.000 | Kumar et al.[1] | Georgakilas et al.[2] |
| chr1 | FRA1K | 1q31 | 185.800.001 | 198.700.000 | Kumar et al.[1] | Georgakilas et al.[2] |
| chr1 | FRA1L | 1p31 | 60.800.001 | 84.400.000 | Kumar et al.[1] | Georgakilas et al.[2] |
| chr1 | FRA1M | 1p21.3 | 94.300.001 | 99.300.000 | Kumar et al.[1] | Mirceta et al. [3] |
| chr2 | FRA2A | 2q11.2 | 96.000.001 | 102.100.000 | Kumar et al.[1] | Georgakilas et al.[2] |
| chr2 | FRA2B | 2q13 | 108.700.001 | 112.200.000 | Kumar et al.[1] | Georgakilas et al.[2] |
| chr2 | FRA2C | 2p24.2 | 16.500.001 | 19.000.000 | Kumar et al.[1] | Brison et al.[4] |
| chr2 | FRA2D | 2p16.2 | 52.600.001 | 54.700.000 | Kumar et al.[1] | Brison et al.[4] |
| chr2 | FRA2E | 2p13 | 68.400.001 | 74.800.000 | Kumar et al.[1] | Georgakilas et al.[2] |
| chr2 | FRA2F | 2q21.3 | 134.300.001 | 136.100.000 | Smith et al. [5] | Georgakilas et al.[2] |
| chr2 | FRA2G | 2q31 | 168.900.001 | 182.100.000 | Debacker and Kooy[6] | |
| chr2 | FRA2H | 2q32.1 | 182.100.001 | 188.500.000 | Kumar et al.[1] | Brison et al.[4] |
| chr2 | FRA2I | 2q33 | 196.600.001 | 208.200.000 | Kumar et al.[1] | Georgakilas et al.[2] |
| chr2 | FRA2J | 2q37.3 | 236.400.001 | 242.193.529 | Kumar et al.[1] | Georgakilas et al.[2] |
| chr2 | FRA2L | 2p11.2 | 83.100.001 | 91.800.000 | Schuffenhauer et al. [7] | |
| chr2 | FRA2S | 2q22.3–q23.3 | 143.400.001 | 154.000.000 | Pelliccia et al. [8] | |
| chr3 | FRA3A | 3p24.2 | 23.800.001 | 26.300.000 | Kumar et al.[1] | Georgakilas et al.[2] |
| chr3 | FRA3B | 3p14.2 | 58.600.001 | 63.800.000 | Kumar et al.[1] | Brison et al.[4] |
| chr3 | FRA3C | 3q27 | 183.000.001 | 188.200.000 | Kumar et al.[1] | Georgakilas et al.[2] |

| | | | | | | |
|---|---|---|---|---|---|---|
| chr3 | FRA3D | 3q25 | 149.200.001 | 161.000.000 | Debacker and Kooy[6] | Georgakilas et al.[2] |
| chr4 | FRA4A | 4p16.1 | 6.000.001 | 11.300.000 | Kumar et al.[1] | Georgakilas et al.[2] |
| chr4 | FRA4B | 4q12 | 51.800.001 | 58.500.000 | Kumar et al.[1] | Georgakilas et al.[2] |
| chr4 | FRA4C | 4q31.1 | 138.500.001 | 140.600.000 | Kumar et al.[1] | Georgakilas et al.[2] |
| chr4 | FRA4D | 4p15 | 11.300.001 | 35.800.000 | Kumar et al.[1] | Brison et al.[4] |
| chr4 | FRA4E | 4q27 | 119.900.001 | 122.800.000 | Debacker and Kooy[6] | |
| chr4 | FRA4F | 4q22 | 87.100.001 | 97.900.000 | Rozier et al. [9] | Georgakilas et al.[2] |
| chr5 | FRA5A | 5p13 | 28.900.001 | 42.500.000 | Kumar et al.[1] | Georgakilas et al.[2] |
| chr5 | FRA5C | 5q31.1 | 131.200.001 | 136.900.000 | Kumar et al.[1] | Georgakilas et al.[2] |
| chr5 | FRA5D | 5q15 | 93.000.001 | 98.900.000 | Debacker and Kooy[6] | |
| chr5 | FRA5E | 5p14 | 18.400.001 | 28.900.000 | Debacker and Kooy[6] | Brison et al.[4] |
| chr5 | FRA5F | 5q21 | 98.900.001 | 110.200.000 | Georgakilas et al.[2] | Georgakilas et al.[2] |
| chr5 | FRA5G | 5q35 | 169.000.001 | 181.538.259 | Debacker and Kooy[6] | |
| chr5 | FRA5H | 5q11.2 | 51.400.001 | 59.600.000 | Kumar et al.[1] | |
| chr6 | FRA6A | 6p23 | 13.400.001 | 15.200.000 | Kumar et al.[1] | Georgakilas et al.[2] |
| chr6 | FRA6b | 6p25.1 | 4.200.001 | 7.100.000 | Kumar et al.[1] | |
| chr6 | FRA6C | 6p22.2 | 25.200.001 | 27.100.000 | Kumar et al.[1] | Brison et al.[4] |
| chr6 | FRA6D | 6q13 | 69.200.001 | 75.200.000 | Kumar et al.[1] | Georgakilas et al.[2] |
| chr6 | FRA6E | 6q26 | 160.600.001 | 164.100.000 | Debacker and Kooy[6] | |
| chr6 | FRA6F | 6q21 | 4.200.001 | 7.100.000 | Morelli et al.[10] | Georgakilas et al.[2] |
| chr6 | FRA6G | 6q15 | 87.300.001 | 92.500.000 | Georgakilas et al. [2] | |
| chr6 | FRA6H | 6p21 | 30.500.001 | 46.200.000 | Fechter et al. [11] | |
| chr7 | FRA7A | 7p11.2 | 53.900.001 | 58.100.000 | Kumar et al.[1] | Georgakilas et al.[2] |
| chr7 | FRA7b | 7p22 | 1 | 7.200.000 | Kumar et al.[1] | Georgakilas et al.[2] |
| chr7 | FRA7C | 7p14.2 | 34.900.001 | 37.100.000 | Kumar et al.[1] | Georgakilas et al.[2] |
| chr7 | FRA7D | 7p13 | 43.300.001 | 45.400.000 | Kumar et al.[1] | Georgakilas et al.[2] |
| chr7 | FRA7E | 7q21.2 | 91.500.001 | 93.300.000 | Kumar et al.[1] | Georgakilas et al.[2] |
| chr7 | FRA7f | 7p22 | 1 | 7.200.000 | Kumar et al.[1] | Georgakilas et al.[2] |
| chr7 | FRA7G | 7q31.2 | 115.000.001 | 117.700.000 | Brison et al.[4] | |
| chr7 | FRA7H | 7q32.3 | 130.800.001 | 132.900.000 | Brison et al.[4] | |
| chr7 | FRA7I | 7q36 | 148.200.001 | 159.345.973 | Kumar et al.[1] | Georgakilas et al.[2] |

| | | | | | | |
|---|---|---|---|---|---|---|
| chr7 | FRA7J | 7q11 | 60.100.001 | 77.900.000 | Georgakilas et al.[2] | |
| chr7 | FRA7K | 7q22-7q31.1 | 98.400.001 | 115.000.000 | Brison et al.[4] | |
| chr8 | FRA8A | 8q22.3 | 100.500.001 | 105.100.000 | Kumar et al.[1] | Georgakilas et al.[2] |
| chr8 | FRA8B | 8q22.1 | 92.300.001 | 97.900.000 | Kumar et al.[1] | Georgakilas et al.[2] |
| chr8 | FRA8C | 8q24.1 | 121.612.116 | 121.641.440 | Brison et al.[4] | |
| chr8 | FRA8C | 8q24.1 | 56.160.909 | 56.211.273 | Brison et al.[4] | |
| chr8 | FRA8D | 8q24.3 | 138.900.001 | 145.138.636 | Ferber et al. [12] | |
| chr8 | FRA8E | 8q24.1 | 121.612.116 | 121.641.440 | Brison et al.[4] | |
| chr8 | FRA8E | 8q24.1 | 56.160.909 | 56.211.273 | Brison et al.[4] | |
| chr9 | FRA9B | 9q32 | 112.100.001 | 114.900.000 | Mirceta et al. [3] | |
| chr9 | FRA9C | 9p21 | 19.900.001 | 33.200.000 | Kumar et al.[1] | Georgakilas et al.[2] |
| chr9 | FRA9D | 9q22.1 | 87.800.001 | 89.200.000 | Kumar et al.[1] | Georgakilas et al.[2] |
| chr9 | FRA9E | 9q32 | 112.100.001 | 114.900.000 | Kumar et al.[1] | Georgakilas et al.[2] |
| chr9 | FRA9F | 9q12 | 45.500.001 | 61.500.000 | Kumar et al.[1] | Georgakilas et al.[2] |
| chr9 | FRA9G | 9p22.2 | 16.600.001 | 18.500.000 | Kumar et al.[1] | Georgakilas et al.[2] |
| chr10 | Fra10A | 10q23.3 | 93.060.798 | 93.069.540 | Kumar et al.[1] | Mirceta et al. [3] |
| chr10 | Fra10B | 10q25.2 | 110.100.001 | 113.100.000 | Kumar et al.[1] | Mirceta et al. [3] |
| chr10 | Fra10C | 10q21 | 51.100.001 | 68.800.000 | Kumar et al.[1] | Brison et al.[4] |
| chr10 | Fra10D | 10q22.1 | 68.800.001 | 73.100.000 | Kumar et al.[1] | Georgakilas et al.[2] |
| chr10 | Fra10E | 10q25.2 | 110.100.001 | 113.100.000 | Kumar et al.[1] | Georgakilas et al.[2] |
| chr10 | Fra10F | 10q26.11-10q26.13 | 117.300.001 | 125.700.000 | Kumar et al.[1] | |
| chr10 | Fra10G | 10q11.21-10q11.23 | 41.600.001 | 51.100.000 | Dillon et al. [13] | |
| chr11 | FRA11b | 11q23.3 | 114.600.001 | 121.300.000 | Kumar et al.[1] | Mirceta et al. [3] |
| chr11 | FRA11C | 11p15.1 | 16.900.001 | 22.000.000 | Kumar et al.[1] | Brison et al.[4] |
| chr11 | FRA11D | 11p14.2 | 26.200.001 | 27.200.000 | Kumar et al.[1] | Brison et al.[4] |
| chr11 | FRA11E | 11p13 | 31.000.001 | 36.400.000 | Kumar et al.[1] | Georgakilas et al.[2] |
| chr11 | FRA11F | 11q14.2 | 85.900.001 | 88.600.000 | Kumar et al.[1] | Georgakilas et al.[2] |
| chr11 | FRA11G | 11q23.3 | 114.600.001 | 121.300.000 | Kumar et al.[1] | Brison et al.[4] |
| chr11 | FRA11EH | 11q13 | 63.600.001 | 77.400.000 | Kumar et al.[1] | |
| chr11 | FRA11I | 11p15.1 | 16.900.001 | 22.000.000 | Kumar et al.[1] | Mirceta et al. [3] |
| chr12 | FRA12A | 12q13.11-12q13.13 | 46.000.001 | 54.500.000 | Debacker and Kooy[6] | Mirceta et al. [3] |

| | | | | | | |
|---|---|---|---|---|---|---|
| chr12 | FRA12b | 12q21.31-12q21.33 | 79.900.001 | 92.200.000 | Debacker and Kooy[6] | Brison et al.[4] |
| chr12 | FRA12C | 12q24 | 108.600.001 | 133.275.309 | Debacker and Kooy[6] | Mirceta et al. [3] |
| chr12 | FRA12E | 12q24 | 108.600.001 | 133.275.309 | Kumar et al.[1] | Georgakilas et al.[2] |
| chr13 | FRA13A | 13q13.2 | 33.400.001 | 34.900.000 | Kumar et al.[1] | Brison et al.[4] |
| chr13 | FRA13B | 13q21 | 54.700.001 | 72.800.000 | Kumar et al.[1] | Georgakilas et al.[2] |
| chr13 | FRA13D | 13q32 | 94.400.001 | 101.100.000 | Kumar et al.[1] | Georgakilas et al.[2] |
| chr13 | FRA13E | 13q22 | 72.800.001 | 78.500.000 | Kumar et al.[1] | Fechter et al. [11] |
| chr14 | FRA14B | 14q13 | 32.900.001 | 37.400.000 | Brison et al.[4] | |
| chr14 | FRA14C | 14q24.1 | 67.400.001 | 69.800.000 | Kumar et al.[1] | Georgakilas et al.[2] |
| chr15 | FRA15A | 15q22 | 58.800.001 | 67.200.000 | Kumar et al.[1] | Georgakilas et al.[2] |
| chr16 | FRA16A | 16p12.3 | 16.700.001 | 21.200.000 | Mirceta et al. [3] | |
| chr16 | FRA16B | 16q22.1 | 66.600.001 | 70.800.000 | Kumar et al.[1] | Mirceta et al. [3] |
| chr16 | FRA16C | 16q22.1 | 63.917.503 | 63.934.965 | Kumar et al.[1] | Georgakilas et al.[2] |
| chr16 | FRA16D | 16q23.2 | 79.200.001 | 81.600.000 | Brison et al.[4] | |
| chr16 | FRA16E | 16p12.1 | 24.200.001 | 28.500.000 | Kumar et al.[1] | Mirceta et al. [3] |
| chr17 | FRA17A | 17p12 | 10.800.001 | 16.100.000 | Kumar et al.[1] | Mirceta et al. [3] |
| chr17 | FRA17B | 17q23.1 | 59.500.001 | 60.200.000 | Kumar et al.[1] | Georgakilas et al.[2] |
| chr18 | FRA18A | 18q12.2 | 35.100.001 | 39.500.000 | Kumar et al.[1] | Brison et al.[4] |
| chr18 | FRA18B | 18q21.31-18q21.33 | 56.200.001 | 63.900.000 | Brison et al.[4] | |
| chr18 | FRA18C | 18q22 | 63.900.001 | 75.400.000 | Brison et al.[4] | |
| chr19 | FRA19A | 19q13 | 31.900.001 | 58.617.616 | Kumar et al.[1] | Brison et al.[4] |
| chr19 | FRA19B | 19p13.1 | 1 | 19.900.000 | Mirceta et al. [3] | |
| chr20 | FRA20A | 20p11.23 | 17.900.001 | 21.300.000 | Kumar et al.[1] | Georgakilas et al.[2] |
| chr20 | FRA20B | 20p12.2 | 9.200.001 | 12.000.000 | Kumar et al.[1] | Georgakilas et al.[2] |
| chr21 | FRA21A | 21q11.2 | 13.000.001 | 15.000.000 | Kumar et al.[1] | Georgakilas et al.[2] |
| chr22 | FRA22A | 22q13 | 37.200.001 | 50.818.468 | Kumar et al.[1] | Mirceta et al. [3] |
| chr22 | FRA22B | 22q12.2 | 29.200.001 | 31.800.000 | Kumar et al.[1] | Brison et al.[4] |
| **Fragile sites excluded from analysis due to another CFS covering the same region** | | | | | | |
| chr1 | FRA1C | 1p31.2 | 68.500.001 | 69.300.000 | Kumar et al.[1] | Georgakilas et al.[2] |
| chr2 | FRA2K | 2q22.3 | 143.400.001 | 147.900.000 | Kumar et al.[1] | Georgakilas et al.[2] |

| | | | | | | |
|---|---|---|---|---|---|---|
| chr5 | FRA5B | 5q15 | 93.000.001 | 98.900.000 | Georgakilas et al.[2] | |
| chr9 | FRA9A | 9p21 | 19.900.001 | 33.200.000 | Kähkönen [14] | Georgakilas et al.[2] |
| chr11 | FRA11A | 11q13.3 | 68.700.001 | 70.500.000 | Kumar et al.[1] | Georgakilas et al.[2] |
| chr12 | FRA12D | 12q24.13 | 111.900.001 | 113.900.000 | Mirceta et al. [3] | |
| chr13 | FRA13C | 13q21.2 | 59.000.001 | 61.800.000 | Kumar et al.[1] | Brison et al.[4] |

**References for Supplementary Table 3**


1. Kumar, R. *et al.* HumCFS: A database of fragile sites in human chromosomes. *BMC Genomics* **19**, 1–8 (2019).
2. Georgakilas, A. G. *et al.* Are common fragile sites merely structural domains or highly organized ‘functional’ units susceptible to oncogenic stress? *Cell. Mol. Life Sci.* **71**, 4519–4544 (2014).
3. Mirceta, M., Shum, N., Schmidt, M. H. M. & Pearson, C. E. Fragile sites, chromosomal lesions, tandem repeats, and disease. *Front. Genet.* **13**, 1–45 (2022).
4. Brison, O. *et al.* Transcription-mediated organization of the replication initiation program across large genes sets common fragile sites genome-wide. *Nat. Commun.* **10**, 1–12 (2019).
5. Smith, D. I., Zhu, Y., McAvoy, S. & Kuhn, R. Common fragile sites, extremely large genes, neural development and cancer. *FRAGILOME - Common Fragile Sites Genomic Instab. Cancer* **232**, 48–57 (2006).
6. Debacker, K. & Frank Kooy, R. Fragile sites and human disease. *Hum. Mol. Genet.* **16**, 150–158 (2007).
7. Schuffenhauer, S., Lederer, G. & Murken, J. A heritable folate-sensitive fragile site on chromosome 2p11. 2 (FRA2L). *Chromosome Res.* **4**, 252–254 (1996).
8. Pelliccia, F., Bosco, N. & Rocchi, A. Breakages at common fragile sites set boundaries of amplified regions in two leukemia cell lines K562 – Molecular characterization of FRA2H and localization of a new CFS FRA2S. *Cancer Lett.* **299**, 37–44 (2010).
9. Rozier, L., El-Achkar, E., Apiou, F. & Debatisse, M. Characterization of a conserved aphidicolin-sensitive common fragile site at human 4q22 and mouse 6C1: possible association with an inherited disease and cancer. *Oncogene* **23**, 6872–6880 (2004).
10. Morelli, C. *et al.* Characterization of a 4-Mb region at chromosome 6q21 harboring a replicative senescence gene. *Cancer Res.* **57**, 4153–4157 (1997).
11. Fechter, A., Buettel, I., Kuehnel, E., Schwab, M. & Savelyeva, L. Cloning of genetically tagged chromosome break sequences reveals new fragile sites at 6p21 and 13q22. *Int. J. Cancer* **120**, 2359–2367 (2007).
12. Ferber, M. J. *et al.* Positioning of cervical carcinoma and Burkitt lymphoma translocation breakpoints with respect to the human papillomavirus integration cluster in FRA8C at 8q24.13. *Cancer Genet. Cytogenet.* **154**, 1–9 (2004).
13. Dillon, L. W., Pierce, L. C. T., Ng, M. C. Y. & Wang, Y.-H. Role of DNA secondary structures in fragile site breakage along human chromosome 10. *Hum. Mol. Genet.* **22**, 1443–1456 (2013).
14. Kähkönen, M. Population cytogenetics of folate-sensitive fragile sites: I. Common fragile sites. *Hum. Genet.* **80**, 344–348 (1988).

**Supplementary Table 4. Guide RNA sequences**

| sgRNA ID | Sequence (5'-3') PAM |
|---|---|
| BLM_Ex3_cr1 | GATGTGATTTGCATCGATGA TGG |
| TOP3A_Ex18_cr1 | TGTGCTGCTCACTGTCCGTA AGG |
| RMI1_Ex3_cr1 | GGAATGGCTATCTGAACTGC TGG |
| RMI2_Ex1_cr2 | AGCGGGGACTTCTCGGTCCG CGG |
| RMI2_Ex1_cr3 | AGGGTAGTGATGGCGGACCG CGG |
| RYR-R2474Sins-crRNA | TATTCCTTGACAGGGTCTAT GGG |

**Supplementary Table 5. List of genotyping primer sequences**

| Gene | Primer ID | Sequence (5'-3') |
|---|---|---|
| *BLM* | E4-2_for | CACCAACTGTAAAGAAATCCC |
| *BLM* | E4-2_rev | TCATCAGTACTGCCTGCCTC |
| *TOP3A* | E18_for | CTGGGTCCTCAAAGGCTCTG |
| *TOP3A* | E18_rev | CATCCTTCTGCACAGTCCGT |
| *RMI1* | E3_for | CTTTTCTTTGACTTGCAAAAATGG |
| *RMI1* | E3_rev | AGGGTGGAGAATACAAATCCATG |

**Supplementary Table 6. Development stages of iPSC samples for each assay.** p indicates passage number.

| Assay | WT | *BLM*$^{c.715del2/c.716del}$ | *TOP3A*$^{c.2473ins/c.2475del7}$ | *RMI1*$^{c.1228del7/c.1230del}$ |
|---|---|---|---|---|
| Induced pluripotency | p1 | - | - | - |
| Genotyping and confirmation of knockouts | - | p26 | p26 | p26 |
| Karyotype analysis | - | p30 | p30 | p30 |
| Array CGH | - | p31 | p31 | p31 |
| SCE | p31 | p31 | p31 | p31 |
| Chromatin bridge/lagging DNA/UFBs | p31/p32/p33 | p30/p31/p30* | p31/p32/p32* | p31/p31*/p31* |
| scWGS | p35 | p37 | p34 | p34 |

*different freezing stocks were used for the same passage numbers.

### References


1. Gönenc, I. I. *et al.* Phenotypic spectrum of BLM- and RMI1–related Bloom syndrome. *Clin. Genet.* **101**, 559–564 (2022).
2. Martin, C. A. *et al.* Mutations in TOP3A cause a Bloom syndrome-like disorder. *Am. J. Hum. Genet.* **103**, 221–231 (2018).
3. Macheret, M. *et al.* High-resolution mapping of mitotic DNA synthesis regions and common fragile sites in the human genome through direct sequencing. *Cell Res.* **30**, 997–1008 (2020).
4. Ji, F. *et al.* Genome-wide high-resolution mapping of mitotic DNA synthesis sites and common fragile sites by direct sequencing. *Cell Res.* **30**, 1009–1023 (2020).
