## Supplementary Material to Figure 4 and Figure 6 for "BTRR complex deficiency is a driver for genomic instability in Bloom syndrome": biocircos-1.1.2.js

```
/**
* BioCircos.js is an open source interactive Javascript library which 
* provides an easy way to interactive display biological data on the web.
* It implements a raster-based SVG visualization using the open source 
* Javascript framework jquery.js. BioCircos.js is multiplatform and works 
* in all major internet browsers (Internet Explorer, Mozilla Firefox, 
* Google Chrome, Safari, Opera). Its speed is determined by the client's 
* hardware and internet browser. For smoothest user experience, we recommend 
* Google Chrome.
*
* Source code, tutorial, documentation, and example data are freely available
* from BioCircos.js website "http://bioinfo.ibp.ac.cn/biocircos/".
* 
* @author <a href="">Ya Cui</a>, <a href="">Xiaowei Chen</a>
* @version 1.1.2+
*
* @example 
*      var BioCircosGenome = [
*         ["chr1" , 249250621],
*         ["chr2" , 243199373]
*      ];
*      BioCircos01 = new BioCircos(BioCircosGenome,{
*         target : "biocircos",
*         svgWidth : 900,
*         svgHeight : 600
*      });
*      BioCircos01.draw_genome(BioCircos01.genomeLength);
*
**/

var BioCircos;

(function($){

  BioCircos = function(){
      var self = this;

      if(arguments.length >= 2){
            self.argumentsBiocircosSettings=arguments[arguments.length-1];
            self.argumentsBiocircosGenome=arguments[arguments.length-2];

            self.CNV = new Array();
            self.CNVConfig = new Array();
            for (var n=0; n< arguments.length; n++){
                var reg=/^CNV/;
                if(reg.test(arguments[n][0])){
                    self.CNVConfig.push(arguments[n][1]);
                    self.CNV.push(arguments[n][2]);
                }
            }

            self.HEATMAP = new Array();
            self.HEATMAPConfig = new Array();
            for (var n=0; n< arguments.length; n++){
                var reg=/^HEATMAP/;
                if(reg.test(arguments[n][0])){
                    self.HEATMAPConfig.push(arguments[n][1]);
                    self.HEATMAP.push(arguments[n][2]);
                }
            }

            self.SNP = new Array();
            self.SNPConfig = new Array();
            for (var n=0; n< arguments.length; n++){
                var reg=/^SNP/;
                if(reg.test(arguments[n][0])){
                    self.SNPConfig.push(arguments[n][1]);
                    self.SNP.push(arguments[n][2]);
                }
            }

            self.LINK = new Array();
            self.LINKConfig = new Array();
            for (var n=0; n< arguments.length; n++){
                var reg=/^LINK/;
                if(reg.test(arguments[n][0])){
                    self.LINKConfig.push(arguments[n][1]);
                    self.LINK.push(arguments[n][2]);
                }
            }

            self.HISTOGRAM = new Array();
            self.HISTOGRAMConfig = new Array();
            for (var n=0; n< arguments.length; n++){
                var reg=/^HISTOGRAM/;
                if(reg.test(arguments[n][0])){
                    self.HISTOGRAMConfig.push(arguments[n][1]);
                    self.HISTOGRAM.push(arguments[n][2]);
                }
            }

            self.LINE = new Array();
            self.LINEConfig = new Array();
            for (var n=0; n< arguments.length; n++){
                var reg=/^LINE/;
                if(reg.test(arguments[n][0])){
                    self.LINEConfig.push(arguments[n][1]);
                    self.LINE.push(arguments[n][2]);
                }
            }

            self.SCATTER = new Array();
            self.SCATTERConfig = new Array();
            for (var n=0; n< arguments.length; n++){
                var reg=/^SCATTER/;
                if(reg.test(arguments[n][0])){
                    self.SCATTERConfig.push(arguments[n][1]);
                    self.SCATTER.push(arguments[n][2]);
                }
            }

            self.BACKGROUND = new Array();
            self.BACKGROUNDConfig = new Array();
            for (var n=0; n< arguments.length; n++){
                var reg=/^BACKGROUND/;
                if(reg.test(arguments[n][0])){
                    self.BACKGROUNDConfig.push(arguments[n][1]);
                    self.BACKGROUND.push(arguments[n][2]);
                }
            }

            self.TEXT = new Array();
            self.TEXTConfig = new Array();
            for (var n=0; n< arguments.length; n++){
                var reg=/^TEXT/;
                if(reg.test(arguments[n][0])){
                    self.TEXTConfig.push(arguments[n][1]);
                    self.TEXT.push(arguments[n][2]);
                }
            }

            self.ARC = new Array();
            self.ARCConfig = new Array();
            for (var n=0; n< arguments.length; n++){
                var reg=/^ARC/;
                if(reg.test(arguments[n][0])){
                    self.ARCConfig.push(arguments[n][1]);
                    self.ARC.push(arguments[n][2]);
                }
            }

      }else{
            document.getElementById(self.settings.target).innerHTML='Arguments Error: at least two arguments must supplied.<br>example: new BioCircos([FUSION01,CNV01,SNP01,]BioCircosGenome,{target : "biocircos",zoom : true})';
      }

      self.settings = {
          "target" : "biocircos",
          "svgWidth" : 900,
          "svgHeight" : 600,
          "chrPad" : 0.04,
          "innerRadius" : 246,
          "outerRadius" : 270,
          "zoom" : false,
          "genomeFillColor" : ["rgb(153,102,0)", "rgb(102,102,0)", "rgb(153,153,30)", "rgb(204,0,0)","rgb(255,0,0)", "rgb(255,0,204)", "rgb(255,204,204)", "rgb(255,153,0)", "rgb(255,204,0)", "rgb(255,255,0)", "rgb(204,255,0)", "rgb(0,255,0)","rgb(53,128,0)", "rgb(0,0,204)", "rgb(102,153,255)", "rgb(153,204,255)", "rgb(0,255,255)", "rgb(204,255,255)", "rgb(153,0,204)", "rgb(204,51,255)","rgb(204,153,255)", "rgb(102,102,102)", "rgb(153,153,153)", "rgb(204,204,204)"],
          "CNVMouseEvent" : true,
          "CNVMouseClickDisplay" : false,
          "CNVMouseClickColor" : "red",
          "CNVMouseClickArcOpacity" : 1.0,
          "CNVMouseClickArcStrokeColor" : "#F26223",
          "CNVMouseClickArcStrokeWidth" : 0,
          "CNVMouseClickTextFromData" : "fourth",   //first,second,third,fourth column
          "CNVMouseClickTextOpacity" : 1,
          "CNVMouseClickTextColor" : "red",
          "CNVMouseClickTextSize" : 8,
          "CNVMouseClickTextPostionX" : 0,
          "CNVMouseClickTextPostionY" : 0,
          "CNVMouseClickTextDrag" : true,
          "CNVMouseDownDisplay" : false,
          "CNVMouseDownColor" : "green",
          "CNVMouseDownArcOpacity" : 1.0,
          "CNVMouseDownArcStrokeColor" : "#F26223",
          "CNVMouseDownArcStrokeWidth" : 0,
          "CNVMouseEnterDisplay" : false,
          "CNVMouseEnterColor" : "yellow",
          "CNVMouseEnterArcOpacity" : 1.0,
          "CNVMouseEnterArcStrokeColor" : "#F26223",
          "CNVMouseEnterArcStrokeWidth" : 0,
          "CNVMouseLeaveDisplay" : false,
          "CNVMouseLeaveColor" : "pink",
          "CNVMouseLeaveArcOpacity" : 1.0,
          "CNVMouseLeaveArcStrokeColor" : "#F26223",
          "CNVMouseLeaveArcStrokeWidth" : 0,
          "CNVMouseMoveDisplay" : false,
          "CNVMouseMoveColor" : "red",
          "CNVMouseMoveArcOpacity" : 1.0,
          "CNVMouseMoveArcStrokeColor" : "#F26223",
          "CNVMouseMoveArcStrokeWidth" : 0,
          "CNVMouseOutDisplay" : false,
          "CNVMouseOutAnimationTime" : 500,
          "CNVMouseOutColor" : "red",
          "CNVMouseOutArcOpacity" : 1.0,
          "CNVMouseOutArcStrokeColor" : "red",
          "CNVMouseOutArcStrokeWidth" : 0,
          "CNVMouseUpDisplay" : false,
          "CNVMouseUpColor" : "grey",
          "CNVMouseUpArcOpacity" : 1.0,
          "CNVMouseUpArcStrokeColor" : "#F26223",
          "CNVMouseUpArcStrokeWidth" : 0,
          "CNVMouseOverDisplay" : false,
          "CNVMouseOverColor" : "red",
          "CNVMouseOverArcOpacity" : 1.0,
          "CNVMouseOverArcStrokeColor" : "#F26223",
          "CNVMouseOverArcStrokeWidth" : 3,
          "CNVMouseOverTooltipsHtml01" : "chr : ",
          "CNVMouseOverTooltipsHtml02" : "<br>start : ",
          "CNVMouseOverTooltipsHtml03" : "<br>end : ",
          "CNVMouseOverTooltipsHtml04" : "<br>value : ",
          "CNVMouseOverTooltipsHtml05" : "",
          "CNVMouseOverTooltipsPosition" : "absolute",
          "CNVMouseOverTooltipsBackgroundColor" : "white",
          "CNVMouseOverTooltipsBorderStyle" : "solid",
          "CNVMouseOverTooltipsBorderWidth" : 0,
          "CNVMouseOverTooltipsPadding" : "3px",
          "CNVMouseOverTooltipsBorderRadius" : "3px",
          "CNVMouseOverTooltipsOpacity" : 0.8,
          "HEATMAPMouseEvent" : true,
          "HEATMAPMouseClickDisplay" : false,
          "HEATMAPMouseClickColor" : "green",            //"none","red"
          "HEATMAPMouseClickOpacity" : 1.0,            //"none",1.0
          "HEATMAPMouseClickStrokeColor" : "none",  //"none","#F26223"
          "HEATMAPMouseClickStrokeWidth" : "none",          //"none",3
          "HEATMAPMouseDownDisplay" : false,
          "HEATMAPMouseDownColor" : "green",            //"none","red"
          "HEATMAPMouseDownOpacity" : 1.0,            //"none",1.0
          "HEATMAPMouseDownStrokeColor" : "none",  //"none","#F26223"
          "HEATMAPMouseDownStrokeWidth" : "none",          //"none",3
          "HEATMAPMouseEnterDisplay" : false,
          "HEATMAPMouseEnterColor" : "green",            //"none","red"
          "HEATMAPMouseEnterOpacity" : 1.0,            //"none",1.0
          "HEATMAPMouseEnterStrokeColor" : "none",  //"none","#F26223"
          "HEATMAPMouseEnterStrokeWidth" : "none",          //"none",3
          "HEATMAPMouseLeaveDisplay" : false,
          "HEATMAPMouseLeaveColor" : "green",            //"none","red"
          "HEATMAPMouseLeaveOpacity" : 1.0,            //"none",1.0
          "HEATMAPMouseLeaveStrokeColor" : "none",  //"none","#F26223"
          "HEATMAPMouseLeaveStrokeWidth" : "none",          //"none",3
          "HEATMAPMouseMoveDisplay" : false,
          "HEATMAPMouseMoveColor" : "green",            //"none","red"
          "HEATMAPMouseMoveOpacity" : 1.0,            //"none",1.0
          "HEATMAPMouseMoveStrokeColor" : "none",  //"none","#F26223"
          "HEATMAPMouseMoveStrokeWidth" : "none",          //"none",3
          "HEATMAPMouseOutDisplay" : false,
          "HEATMAPMouseOutAnimationTime" : 500,
          "HEATMAPMouseOutColor" : "green",            //"none","red"
          "HEATMAPMouseOutOpacity" : 1.0,            //"none",1.0
          "HEATMAPMouseOutStrokeColor" : "none",  //"none","#F26223"
          "HEATMAPMouseOutStrokeWidth" : "none",          //"none",3
          "HEATMAPMouseUpDisplay" : false,
          "HEATMAPMouseUpColor" : "green",            //"none","red"
          "HEATMAPMouseUpOpacity" : 1.0,            //"none",1.0
          "HEATMAPMouseUpStrokeColor" : "none",  //"none","#F26223"
          "HEATMAPMouseUpStrokeWidth" : "none",          //"none",3
          "HEATMAPMouseOverDisplay" : false,
          "HEATMAPMouseOverColor" : "none",            //"none","red"
          "HEATMAPMouseOverOpacity" : 1.0,            //"none",1.0
          "HEATMAPMouseOverStrokeColor" : "none",  //"none","#F26223"
          "HEATMAPMouseOverStrokeWidth" : "none",          //"none",3
          "HEATMAPMouseOverTooltipsHtml01" : "chr : ",
          "HEATMAPMouseOverTooltipsHtml02" : "<br>position: ",
          "HEATMAPMouseOverTooltipsHtml03" : "-",
          "HEATMAPMouseOverTooltipsHtml04" : "<br>name : ",
          "HEATMAPMouseOverTooltipsHtml05" : "<br>value : ",
          "HEATMAPMouseOverTooltipsHtml06" : "",
          "HEATMAPMouseOverTooltipsPosition" : "absolute",
          "HEATMAPMouseOverTooltipsBackgroundColor" : "white",
          "HEATMAPMouseOverTooltipsBorderStyle" : "solid",
          "HEATMAPMouseOverTooltipsBorderWidth" : 0,
          "HEATMAPMouseOverTooltipsPadding" : "3px",
          "HEATMAPMouseOverTooltipsBorderRadius" : "3px",
          "HEATMAPMouseOverTooltipsOpacity" : 0.8,
          "SNPMouseEvent" : true,
          "SNPMouseClickDisplay" : false,
          "SNPMouseClickColor" : "red",
          "SNPMouseClickCircleSize" : 4,
          "SNPMouseClickCircleOpacity" : 1.0,
          "SNPMouseClickCircleStrokeColor" : "#F26223",
          "SNPMouseClickCircleStrokeWidth" : 0,
          "SNPMouseClickTextFromData" : "fourth",   //first,second,third,fourth column
          "SNPMouseClickTextOpacity" : 1.0,
          "SNPMouseClickTextColor" : "red",
          "SNPMouseClickTextSize" : 8,
          "SNPMouseClickTextPostionX" : 1.0,
          "SNPMouseClickTextPostionY" : 10.0,
          "SNPMouseClickTextDrag" : true,
          "SNPMouseDownDisplay" : false,
          "SNPMouseDownColor" : "green",
          "SNPMouseDownCircleSize" : 4,
          "SNPMouseDownCircleOpacity" : 1.0,
          "SNPMouseDownCircleStrokeColor" : "#F26223",
          "SNPMouseDownCircleStrokeWidth" : 0,
          "SNPMouseEnterDisplay" : false,
          "SNPMouseEnterColor" : "yellow",
          "SNPMouseEnterCircleSize" : 4,
          "SNPMouseEnterCircleOpacity" : 1.0,
          "SNPMouseEnterCircleStrokeColor" : "#F26223",
          "SNPMouseEnterCircleStrokeWidth" : 0,
          "SNPMouseLeaveDisplay" : false,
          "SNPMouseLeaveColor" : "pink",
          "SNPMouseLeaveCircleSize" : 4,
          "SNPMouseLeaveCircleOpacity" : 1.0,
          "SNPMouseLeaveCircleStrokeColor" : "#F26223",
          "SNPMouseLeaveCircleStrokeWidth" : 0,
          "SNPMouseMoveDisplay" : false,
          "SNPMouseMoveColor" : "red",
          "SNPMouseMoveCircleSize" : 2,
          "SNPMouseMoveCircleOpacity" : 1.0,
          "SNPMouseMoveCircleStrokeColor" : "#F26223",
          "SNPMouseMoveCircleStrokeWidth" : 0,
          "SNPMouseOutDisplay" : false,
          "SNPMouseOutAnimationTime" : 500,
          "SNPMouseOutColor" : "red",
          "SNPMouseOutCircleSize" : 2,
          "SNPMouseOutCircleOpacity" : 1.0,
          "SNPMouseOutCircleStrokeColor" : "red",
          "SNPMouseOutCircleStrokeWidth" : 0,
          "SNPMouseUpDisplay" : false,
          "SNPMouseUpColor" : "grey",
          "SNPMouseUpCircleSize" : 4,
          "SNPMouseUpCircleOpacity" : 1.0,
          "SNPMouseUpCircleStrokeColor" : "#F26223",
          "SNPMouseUpCircleStrokeWidth" : 0,
          "SNPMouseOverDisplay" : false,
          "SNPMouseOverColor" : "red",
          "SNPMouseOverCircleSize" : 2,
          "SNPMouseOverCircleOpacity" : 1.0,
          "SNPMouseOverCircleStrokeColor" : "#F26223",
          "SNPMouseOverCircleStrokeWidth" : 3,
          "SNPMouseOverTooltipsHtml01" : "chr : ",
          "SNPMouseOverTooltipsHtml02" : "<br>position : ",
          "SNPMouseOverTooltipsHtml03" : "<br>value : ",
          "SNPMouseOverTooltipsHtml04" : "<br>des : ",
          "SNPMouseOverTooltipsHtml05" : "",
          "SNPMouseOverTooltipsPosition" : "absolute",
          "SNPMouseOverTooltipsBackgroundColor" : "white",
          "SNPMouseOverTooltipsBorderStyle" : "solid",
          "SNPMouseOverTooltipsBorderWidth" : 0,
          "SNPMouseOverTooltipsPadding" : "3px",
          "SNPMouseOverTooltipsBorderRadius" : "3px",
          "SNPMouseOverTooltipsOpacity" : 0.8,
          "TEXTModuleDragEvent" : true,
          "LINKMouseEvent" : true,
          "LINKMouseClickDisplay" : false,
          "LINKMouseClickOpacity" : 1.0,
          "LINKMouseClickStrokeColor" : "green",
          "LINKMouseClickStrokeWidth" : 4,
          "LINKMouseDownDisplay" : false,
          "LINKMouseDownOpacity" : 1.0,
          "LINKMouseDownStrokeColor" : "#F26223",
          "LINKMouseDownStrokeWidth" : 4,
          "LINKMouseEnterDisplay" : false,
          "LINKMouseEnterOpacity" : 1.0,
          "LINKMouseEnterStrokeColor" : "#F26223",
          "LINKMouseEnterStrokeWidth" : 4,
          "LINKMouseLeaveDisplay" : false,
          "LINKMouseLeaveOpacity" : 1.0,
          "LINKMouseLeaveStrokeColor" : "#F26223",
          "LINKMouseLeaveStrokeWidth" : 4,
          "LINKMouseMoveDisplay" : false,
          "LINKMouseMoveOpacity" : 1.0,
          "LINKMouseMoveStrokeColor" : "#F26223",
          "LINKMouseMoveStrokeWidth" : 4,
          "LINKMouseOutDisplay" : false,
          "LINKMouseOutAnimationTime" : 500,
          "LINKMouseOutOpacity" : 1.0,
          "LINKMouseOutStrokeColor" : "red",
          "LINKMouseOutStrokeWidth" : 4,
          "LINKMouseUpDisplay" : false,
          "LINKMouseUpOpacity" : 1.0,
          "LINKMouseUpStrokeColor" : "#F26223",
          "LINKMouseUpStrokeWidth" : 4,
          "LINKMouseOverDisplay" : false,
          "LINKMouseOverOpacity" : 1.0,
          "LINKMouseOverStrokeColor" : "#F26223",
          "LINKMouseOverStrokeWidth" : 3,
          "LINKMouseOverTooltipsHtml01" : "Link : ",
          "LINKMouseOverTooltipsHtml02" : "",
          "LINKMouseOverTooltipsPosition" : "absolute",
          "LINKMouseOverTooltipsBackgroundColor" : "white",
          "LINKMouseOverTooltipsBorderStyle" : "solid",
          "LINKMouseOverTooltipsBorderWidth" : 0,
          "LINKMouseOverTooltipsPadding" : "3px",
          "LINKMouseOverTooltipsBorderRadius" : "3px",
          "LINKMouseOverTooltipsOpacity" : 0.8,
          "LINKLabelDragEvent" : false,
          "HISTOGRAMMouseEvent" : true,
          "HISTOGRAMMouseClickDisplay" : false,
          "HISTOGRAMMouseClickColor" : "red",            //"none","red"
          "HISTOGRAMMouseClickOpacity" : 1.0,            //"none",1.0
          "HISTOGRAMMouseClickStrokeColor" : "none",  //"none","#F26223"
          "HISTOGRAMMouseClickStrokeWidth" : "none",          //"none",3
          "HISTOGRAMMouseDownDisplay" : false,
          "HISTOGRAMMouseDownColor" : "red",            //"none","red"
          "HISTOGRAMMouseDownOpacity" : 1.0,            //"none",1.0
          "HISTOGRAMMouseDownStrokeColor" : "none",  //"none","#F26223"
          "HISTOGRAMMouseDownStrokeWidth" : "none",          //"none",3
          "HISTOGRAMMouseEnterDisplay" : false,
          "HISTOGRAMMouseEnterColor" : "red",            //"none","red"
          "HISTOGRAMMouseEnterOpacity" : 1.0,            //"none",1.0
          "HISTOGRAMMouseEnterStrokeColor" : "none",  //"none","#F26223"
          "HISTOGRAMMouseEnterStrokeWidth" : "none",          //"none",3
          "HISTOGRAMMouseLeaveDisplay" : false,
          "HISTOGRAMMouseLeaveColor" : "red",            //"none","red"
          "HISTOGRAMMouseLeaveOpacity" : 1.0,            //"none",1.0
          "HISTOGRAMMouseLeaveStrokeColor" : "none",  //"none","#F26223"
          "HISTOGRAMMouseLeaveStrokeWidth" : "none",          //"none",3
          "HISTOGRAMMouseMoveDisplay" : false,
          "HISTOGRAMMouseMoveColor" : "red",            //"none","red"
          "HISTOGRAMMouseMoveOpacity" : 1.0,            //"none",1.0
          "HISTOGRAMMouseMoveStrokeColor" : "none",  //"none","#F26223"
          "HISTOGRAMMouseMoveStrokeWidth" : "none",          //"none",3
          "HISTOGRAMMouseOutDisplay" : false,
          "HISTOGRAMMouseOutAnimationTime" : 500,
          "HISTOGRAMMouseOutColor" : "red",            //"none","red"
          "HISTOGRAMMouseOutOpacity" : 1.0,            //"none",1.0
          "HISTOGRAMMouseOutStrokeColor" : "none",  //"none","#F26223"
          "HISTOGRAMMouseOutStrokeWidth" : "none",          //"none",3
          "HISTOGRAMMouseUpDisplay" : false,
          "HISTOGRAMMouseUpColor" : "red",            //"none","red"
          "HISTOGRAMMouseUpOpacity" : 1.0,            //"none",1.0
          "HISTOGRAMMouseUpStrokeColor" : "none",  //"none","#F26223"
          "HISTOGRAMMouseUpStrokeWidth" : "none",          //"none",3
          "HISTOGRAMMouseOverDisplay" : false,
          "HISTOGRAMMouseOverColor" : "red",            //"none","red"
          "HISTOGRAMMouseOverOpacity" : 1.0,            //"none",1.0
          "HISTOGRAMMouseOverStrokeColor" : "none",  //"none","#F26223"
          "HISTOGRAMMouseOverStrokeWidth" : "none",          //"none",3
          "HISTOGRAMMouseOverTooltipsHtml01" : "chr :",
          "HISTOGRAMMouseOverTooltipsHtml02" : "<br>position: ",
          "HISTOGRAMMouseOverTooltipsHtml03" : "-",
          "HISTOGRAMMouseOverTooltipsHtml04" : "<br>name : ",
          "HISTOGRAMMouseOverTooltipsHtml05" : "<br>value : ",
          "HISTOGRAMMouseOverTooltipsHtml06" : "",
          "HISTOGRAMMouseOverTooltipsPosition" : "absolute",
          "HISTOGRAMMouseOverTooltipsBackgroundColor" : "white",
          "HISTOGRAMMouseOverTooltipsBorderStyle" : "solid",
          "HISTOGRAMMouseOverTooltipsBorderWidth" : 0,
          "HISTOGRAMMouseOverTooltipsPadding" : "3px",
          "HISTOGRAMMouseOverTooltipsBorderRadius" : "3px",
          "HISTOGRAMMouseOverTooltipsOpacity" : 0.8,
          "LINEMouseEvent" : true,
          "LINEMouseClickDisplay" : false,
          "LINEMouseClickLineOpacity" : 1,           //"none"
          "LINEMouseClickLineStrokeColor" : "red",   //"none"
          "LINEMouseClickLineStrokeWidth" : "none",  //"none"
          "LINEMouseDownDisplay" : false,
          "LINEMouseDownLineOpacity" : 1,           //"none"
          "LINEMouseDownLineStrokeColor" : "red",   //"none"
          "LINEMouseDownLineStrokeWidth" : "none",  //"none"
          "LINEMouseEnterDisplay" : false,
          "LINEMouseEnterLineOpacity" : 1,           //"none"
          "LINEMouseEnterLineStrokeColor" : "red",   //"none"
          "LINEMouseEnterLineStrokeWidth" : "none",  //"none"
          "LINEMouseLeaveDisplay" : false,
          "LINEMouseLeaveLineOpacity" : 1,           //"none"
          "LINEMouseLeaveLineStrokeColor" : "red",   //"none"
          "LINEMouseLeaveLineStrokeWidth" : "none",  //"none"
          "LINEMouseMoveDisplay" : false,
          "LINEMouseMoveLineOpacity" : 1,           //"none"
          "LINEMouseMoveLineStrokeColor" : "red",   //"none"
          "LINEMouseMoveLineStrokeWidth" : "none",  //"none"
          "LINEMouseOutDisplay" : false,
          "LINEMouseOutAnimationTime" : 500,
          "LINEMouseOutLineOpacity" : 1.0,   //"none"
          "LINEMouseOutLineStrokeColor" : "red",    //"none"
          "LINEMouseOutLineStrokeWidth" : "none",    //"none"
          "LINEMouseUpDisplay" : false,
          "LINEMouseUpLineOpacity" : 1,           //"none"
          "LINEMouseUpLineStrokeColor" : "red",   //"none"
          "LINEMouseUpLineStrokeWidth" : "none",  //"none"
          "LINEMouseOverDisplay" : false,
          "LINEMouseOverLineOpacity" : 1,           //"none"
          "LINEMouseOverLineStrokeColor" : "red",   //"none"
          "LINEMouseOverLineStrokeWidth" : "none",  //"none"
          "LINEMouseOverTooltipsHtml01" : "Line",
          "LINEMouseOverTooltipsPosition" : "absolute",
          "LINEMouseOverTooltipsBackgroundColor" : "white",
          "LINEMouseOverTooltipsBorderStyle" : "solid",
          "LINEMouseOverTooltipsBorderWidth" : 0,
          "LINEMouseOverTooltipsPadding" : "3px",
          "LINEMouseOverTooltipsBorderRadius" : "3px",
          "LINEMouseOverTooltipsOpacity" : 0.8,
          "SCATTERMouseEvent" : true,
          "SCATTERMouseClickDisplay" : false,
          "SCATTERMouseClickColor" : "red",
          "SCATTERMouseClickCircleSize" : 4,
          "SCATTERMouseClickCircleOpacity" : 1.0,
          "SCATTERMouseClickCircleStrokeColor" : "#F26223",
          "SCATTERMouseClickCircleStrokeWidth" : 0,
          "SCATTERMouseClickTextFromData" : "fourth",   //first,second,third,fourth column
          "SCATTERMouseClickTextOpacity" : 1,
          "SCATTERMouseClickTextColor" : "red",
          "SCATTERMouseClickTextSize" : 8,
          "SCATTERMouseClickTextPostionX" : 1.0,
          "SCATTERMouseClickTextPostionY" : 10.0,
          "SCATTERMouseClickTextDrag" : true,
          "SCATTERMouseDownDisplay" : false,
          "SCATTERMouseDownColor" : "green",
          "SCATTERMouseDownCircleSize" : 4,
          "SCATTERMouseDownCircleOpacity" : 1.0,
          "SCATTERMouseDownCircleStrokeColor" : "#F26223",
          "SCATTERMouseDownCircleStrokeWidth" : 0,
          "SCATTERMouseEnterDisplay" : false,
          "SCATTERMouseEnterColor" : "yellow",
          "SCATTERMouseEnterCircleSize" : 4,
          "SCATTERMouseEnterCircleOpacity" : 1.0,
          "SCATTERMouseEnterCircleStrokeColor" : "#F26223",
          "SCATTERMouseEnterCircleStrokeWidth" : 0,
          "SCATTERMouseLeaveDisplay" : false,
          "SCATTERMouseLeaveColor" : "pink",
          "SCATTERMouseLeaveCircleSize" : 4,
          "SCATTERMouseLeaveCircleOpacity" : 1.0,
          "SCATTERMouseLeaveCircleStrokeColor" : "#F26223",
          "SCATTERMouseLeaveCircleStrokeWidth" : 0,
          "SCATTERMouseMoveDisplay" : false,
          "SCATTERMouseMoveColor" : "red",
          "SCATTERMouseMoveCircleSize" : 2,
          "SCATTERMouseMoveCircleOpacity" : 1.0,
          "SCATTERMouseMoveCircleStrokeColor" : "#F26223",
          "SCATTERMouseMoveCircleStrokeWidth" : 0,
          "SCATTERMouseOutDisplay" : false,
          "SCATTERMouseOutAnimationTime" : 500,
          "SCATTERMouseOutColor" : "red",
          "SCATTERMouseOutCircleSize" : 2,
          "SCATTERMouseOutCircleOpacity" : 1.0,
          "SCATTERMouseOutCircleStrokeColor" : "red",
          "SCATTERMouseOutCircleStrokeWidth" : 0,
          "SCATTERMouseUpDisplay" : false,
          "SCATTERMouseUpColor" : "grey",
          "SCATTERMouseUpCircleSize" : 4,
          "SCATTERMouseUpCircleOpacity" : 1.0,
          "SCATTERMouseUpCircleStrokeColor" : "#F26223",
          "SCATTERMouseUpCircleStrokeWidth" : 0,
          "SCATTERMouseOverDisplay" : false,
          "SCATTERMouseOverColor" : "red",
          "SCATTERMouseOverCircleSize" : 2,
          "SCATTERMouseOverCircleOpacity" : 1.0,
          "SCATTERMouseOverCircleStrokeColor" : "#F26223",
          "SCATTERMouseOverCircleStrokeWidth" : 3,
          "SCATTERMouseOverTooltipsHtml01" : "item : ",
          "SCATTERMouseOverTooltipsHtml02" : "<br>start : ",
          "SCATTERMouseOverTooltipsHtml03" : "<br>end : ",
          "SCATTERMouseOverTooltipsHtml04" : "<br>name : ",
          "SCATTERMouseOverTooltipsHtml05" : "<br>des : ",
          "SCATTERMouseOverTooltipsHtml06" : "",
          "SCATTERMouseOverTooltipsPosition" : "absolute",
          "SCATTERMouseOverTooltipsBackgroundColor" : "white",
          "SCATTERMouseOverTooltipsBorderStyle" : "solid",
          "SCATTERMouseOverTooltipsBorderWidth" : 0,
          "SCATTERMouseOverTooltipsPadding" : "3px",
          "SCATTERMouseOverTooltipsBorderRadius" : "3px",
          "SCATTERMouseOverTooltipsOpacity" : 0.8,
          "ARCMouseEvent" : true,
          "ARCMouseClickDisplay" : false,
          "ARCMouseClickColor" : "red",
          "ARCMouseClickArcOpacity" : 1.0,
          "ARCMouseClickArcStrokeColor" : "#F26223",
          "ARCMouseClickArcStrokeWidth" : 1,
          "ARCMouseClickTextFromData" : "fourth",   //first,second,third,fourth column
          "ARCMouseClickTextOpacity" : 1,
          "ARCMouseClickTextColor" : "red",
          "ARCMouseClickTextSize" : 8,
          "ARCMouseClickTextPostionX" : 0,
          "ARCMouseClickTextPostionY" : 0,
          "ARCMouseClickTextDrag" : true,
          "ARCMouseDownDisplay" : false,
          "ARCMouseDownColor" : "green",
          "ARCMouseDownArcOpacity" : 1.0,
          "ARCMouseDownArcStrokeColor" : "#F26223",
          "ARCMouseDownArcStrokeWidth" : 0,
          "ARCMouseEnterDisplay" : false,
          "ARCMouseEnterColor" : "yellow",
          "ARCMouseEnterArcOpacity" : 1.0,
          "ARCMouseEnterArcStrokeColor" : "#F26223",
          "ARCMouseEnterArcStrokeWidth" : 0,
          "ARCMouseLeaveDisplay" : false,
          "ARCMouseLeaveColor" : "pink",
          "ARCMouseLeaveArcOpacity" : 1.0,
          "ARCMouseLeaveArcStrokeColor" : "#F26223",
          "ARCMouseLeaveArcStrokeWidth" : 0,
          "ARCMouseMoveDisplay" : false,
          "ARCMouseMoveColor" : "red",
          "ARCMouseMoveArcOpacity" : 1.0,
          "ARCMouseMoveArcStrokeColor" : "#F26223",
          "ARCMouseMoveArcStrokeWidth" : 0,
          "ARCMouseOutDisplay" : false,
          "ARCMouseOutAnimationTime" : 500,
          "ARCMouseOutColor" : "red",
          "ARCMouseOutArcOpacity" : 1.0,
          "ARCMouseOutArcStrokeColor" : "red",
          "ARCMouseOutArcStrokeWidth" : 0,
          "ARCMouseUpDisplay" : false,
          "ARCMouseUpColor" : "grey",
          "ARCMouseUpArcOpacity" : 1.0,
          "ARCMouseUpArcStrokeColor" : "#F26223",
          "ARCMouseUpArcStrokeWidth" : 0,
          "ARCMouseOverDisplay" : false,
          "ARCMouseOverColor" : "red",
          "ARCMouseOverArcOpacity" : 1.0,
          "ARCMouseOverArcStrokeColor" : "#F26223",
          "ARCMouseOverArcStrokeWidth" : 3,
          "ARCMouseOverTooltipsHtml01" : "item : ",
          "ARCMouseOverTooltipsHtml02" : "<br>start : ",
          "ARCMouseOverTooltipsHtml03" : "<br>end : ",
          "ARCMouseOverTooltipsHtml04" : "<br>des : ",
          "ARCMouseOverTooltipsHtml05" : "",
          "ARCMouseOverTooltipsPosition" : "absolute",
          "ARCMouseOverTooltipsBackgroundColor" : "white",
          "ARCMouseOverTooltipsBorderStyle" : "solid",
          "ARCMouseOverTooltipsBorderWidth" : 0,
          "ARCMouseOverTooltipsPadding" : "3px",
          "ARCMouseOverTooltipsBorderRadius" : "3px",
          "ARCMouseOverTooltipsOpacity" : 0.8,
          "genomeBorder" : {
             "display" : true,
             "borderColor" : "#000",
             "borderSize" : 0.5
          },
          "ticks" : {
             "display" : true,
             "len" : 5,
             "color" : "#000",
             "textSize" : 10,
             "textColor" : "#000",
             "scale" : 30000000
          },
          "genomeLabel" : {
             "display" : true,
             "textSize" : 15,
             "textColor" : "#000",
             "dx" : 0.028,
             "dy" : 15,
             "rotation" : 0
          }
      };

      self.CNVsettings = {
          "maxRadius": 200,
          "minRadius": 190,
          "CNVwidth": 10,
          "CNVColor": "#CAE1FF"
      };

      self.HEATMAPsettings = {
          "innerRadius": -100,
          "outerRadius": -100,
          "maxColor": "red",
          "minColor": "green"
      };

      self.SNPsettings = {
          "maxRadius": 200,
          "minRadius": 190,
          "SNPFillColor": "red",
          "PointType": "circle", //circle,rect
          "circleSize": 2,
          "rectWidth": 2,
          "rectHeight": 2,
          "SNPAnimationDisplay": false,
          "SNPAnimationTime": 2000,
          "SNPAnimationDelay": 20,
          "SNPAnimationType": "bounce",  //linear,circle,elastic,bounce
      };

      self.LINKsettings = {
          "LinkRadius": 108,
          "LinkFillColor": "red",
          "LinkWidth": 3,
          "displayLinkAxis": true,
          "LinkAxisColor": "#B8B8B8",
          "LinkAxisWidth": 0.5,
          "LinkAxisPad": 3,
          "displayLinkLabel": true,
          "LinkLabelColor": "red",
          "LinkLabelSize": 13,
          "LinkLabelPad": 8
      };

      self.HISTOGRAMsettings = {
          "maxRadius": 108,
          "minRadius": 95,
          "histogramFillColor": "red"
      };

      self.LINEsettings = {
          "maxRadius": 108,
          "minRadius": 95,
          "LineFillColor": "red",
          "LineWidth": 0.5
      };

      self.SCATTERsettings = {
          "SCATTERRadius": 140,
          "innerCircleSize": 1,
          "outerCircleSize": 5,
          "innerCircleColor": "#F26223",
          "outerCircleColor": "#F26223",
          "innerPointType": "circle", //circle,rect
          "outerPointType": "circle", //circle,rect
          "innerrectWidth": 2,
          "innerrectHeight": 2,
          "outerrectWidth": 2,
          "outerrectHeight": 2,
          "outerCircleOpacity": 1,
          "random_data": 0,
          "SCATTERAnimationDisplay": false,
          "SCATTERAnimationTime": 2000,
          "SCATTERAnimationDelay": 20,
          "SCATTERAnimationType": "bounce",  //linear,circle,elastic,bounce
      };

      self.BACKGROUNDsettings = {
          "BginnerRadius": 180,
          "BgouterRadius": 230,
          "BgFillColor": "none",
          "BgborderColor": "#000",
          "BgborderSize" : 0.5,
          "axisShow": "false",
          "axisWidth": 0.3,
          "axisColor": "#000",
          "axisOpacity": 0.5,
          "axisNum": 4
      };

      self.TEXTsettings = {
          "x": 20,
          "y": 20,
          "textSize": 10,
          "textWeight": "normal", //normal,bold,bolder,lighter,100,200,300,400,500,600,700,800,900
          "textColor": "#000",
          "textOpacity": 1.0
      };

      self.ARCsettings = {
          "innerRadius": -30,
          "outerRadius": -30
      };

      self.update_settings(self.argumentsBiocircosSettings)

      self.target = "#" + self.settings.target;
      self.svgWidth = self.settings.svgWidth;
      self.svgHeight = self.settings.svgHeight;
      self.chrPad = self.settings.chrPad;
      self.innerRadius = self.settings.innerRadius;
      self.outerRadius = self.settings.outerRadius;
      self.zoom = self.settings.zoom;
      self.testTip = self.settings.testTip;
      self.genomeFillColor = self.settings.genomeFillColor;
      self.genomeBorderDisplay=self.settings.genomeBorder.display;
      self.genomeBorderColor=self.settings.genomeBorder.borderColor;
      self.genomeBorderSize=self.settings.genomeBorder.borderSize;
      self.genome = self.argumentsBiocircosGenome;
      self.genome_matrix(self.argumentsBiocircosGenome)
      self.ticksDisplay=self.settings.ticks.display;
      self.ticksLength=self.settings.ticks.len;
      self.ticksColor=self.settings.ticks.color;
      self.ticksTextSize=self.settings.ticks.textSize;
      self.ticksTextColor=self.settings.ticks.textColor;
      self.ticksScale=self.settings.ticks.scale;
      self.genomeTextDisplay=self.settings.genomeLabel.display;
      self.genomeTextSize=self.settings.genomeLabel.textSize;
      self.genomeTextColor=self.settings.genomeLabel.textColor;
      self.genomeTextDx=self.settings.genomeLabel.dx;
      self.genomeTextDy=self.settings.genomeLabel.dy;
      self.genomeTextRotation=self.settings.genomeLabel.rotation;

      var labeli= self.genomeLabel.length;
      var initGenome = new Object();
      for(var labelk=0;labelk<labeli;labelk++){
          var labelInit=self.genomeLabel[labelk];
          initGenome[labelInit]=labelk;
      }
      self.initGenome = initGenome;

  }

  BioCircos.prototype.update_settings = function(settings_object){
    var self = this;
    $.extend(self.settings, settings_object);
  }

  BioCircos.prototype.update_CNVsettings = function(settings_object){
    var self = this;
    $.extend(self.CNVsettings, settings_object);
  }

  BioCircos.prototype.update_HEATMAPsettings = function(settings_object){
    var self = this;
    $.extend(self.HEATMAPsettings, settings_object);
  }

  BioCircos.prototype.update_SNPsettings = function(settings_object){
    var self = this;
    $.extend(self.SNPsettings, settings_object);
  }

  BioCircos.prototype.update_LINKsettings = function(settings_object){
    var self = this;
    $.extend(self.LINKsettings, settings_object);
  }

  BioCircos.prototype.update_HISTOGRAMsettings = function(settings_object){
    var self = this;
    $.extend(self.HISTOGRAMsettings, settings_object);
  }

  BioCircos.prototype.update_LINEsettings = function(settings_object){
    var self = this;
    $.extend(self.LINEsettings, settings_object);
  }

  BioCircos.prototype.update_SCATTERsettings = function(settings_object){
    var self = this;
    $.extend(self.SCATTERsettings, settings_object);
  }

  BioCircos.prototype.update_BACKGROUNDsettings = function(settings_object){
    var self = this;
    $.extend(self.BACKGROUNDsettings, settings_object);
  }

  BioCircos.prototype.update_TEXTsettings = function(settings_object){
    var self = this;
    $.extend(self.TEXTsettings, settings_object);
  }

  BioCircos.prototype.update_ARCsettings = function(settings_object){
    var self = this;
    $.extend(self.ARCsettings, settings_object);
  }

  BioCircos.prototype.init_CNVsettings = function(){
    var self = this;
    self.CNVsettings = {
          "maxRadius": 200,
          "minRadius": 190,
          "CNVwidth": 10,
          "CNVColor": "#CAE1FF"
    };
  }

  BioCircos.prototype.init_HEATMAPsettings = function(){
    var self = this;
    self.HEATMAPsettings = {
          "innerRadius": -100,
          "outerRadius": -100,
          "maxColor": "red",
          "minColor": "green"
    };
  }

  BioCircos.prototype.init_SNPsettings = function(){
    var self = this;
    self.SNPsettings = {
          "maxRadius": 200,
          "minRadius": 190,
          "SNPFillColor": "red",
          "PointType": "circle",
          "circleSize": 2,
          "rectWidth": 2,
          "rectHeight": 2,
          "SNPAnimationDisplay": false,
          "SNPAnimationTime": 2000,
          "SNPAnimationDelay": 20,
          "SNPAnimationType": "bounce",  //linear,circle,elastic,bounce
    };
  }

  BioCircos.prototype.init_LINKsettings = function(){
    var self = this;
    self.LINKsettings = {
          "LinkRadius": 108,
          "LinkFillColor": "red",
          "LinkWidth": 3,
          "displayLinkAxis": true,
          "LinkAxisColor": "#B8B8B8",
          "LinkAxisWidth": 0.5,
          "LinkAxisPad": 3,
          "displayLinkLabel": true,
          "LinkLabelColor": "red",
          "LinkLabelSize": 13,
          "LinkLabelPad": 8
    };
  }


  BioCircos.prototype.init_HISTOGRAMsettings = function(){
    var self = this;
    self.HISTOGRAMsettings = {
          "maxRadius": 108,
          "minRadius": 95,
          "histogramFillColor": "red"
    };
  }

  BioCircos.prototype.init_LINEsettings = function(){
    var self = this;
    self.LINEsettings = {
          "maxRadius": 108,
          "minRadius": 95,
          "LineFillColor": "red",
          "LineWidth": 0.5
    };
  }

  BioCircos.prototype.init_SCATTERsettings = function(){
    var self = this;
    self.SCATTERsettings = {
          "SCATTERRadius": 140,
          "innerCircleSize": 1,
          "outerCircleSize": 5,
          "innerCircleColor": "#F26223",
          "outerCircleColor": "#F26223",
          "innerPointType": "circle", //circle,rect
          "outerPointType": "circle", //circle,rect
          "innerrectWidth": 2,
          "innerrectHeight": 2,
          "outerrectWidth": 2,
          "outerrectHeight": 2,
          "outerCircleOpacity": 1,
          "random_data": 0,
          "SCATTERAnimationDisplay": false,
          "SCATTERAnimationTime": 2000,
          "SCATTERAnimationDelay": 20,
          "SCATTERAnimationType": "bounce",  //linear,circle,elastic,bounce
      };
  }

  BioCircos.prototype.init_BACKGROUNDsettings = function(){
    var self = this;
    self.BACKGROUNDsettings = {
          "BginnerRadius": 180,
          "BgouterRadius": 230,
          "BgFillColor": "none",
          "BgborderColor": "#000",
          "BgborderSize" : 0.5,
          "axisShow": "false",
          "axisWidth": 0.3,
          "axisColor": "#000",
          "axisOpacity": 0.5,
          "axisNum": 4
    };
  }

  BioCircos.prototype.init_TEXTsettings = function(){
    var self = this;
    self.TEXTsettings = {
          "x": 20,
          "y": 20,
          "textSize": 10,
          "textColor": "#000",
          "textWeight": "normal", //normal,bold,bolder,lighter,100,200,300,400,500,600,700,800,900
          "textOpacity": 1.0
    };
  }

  BioCircos.prototype.init_ARCsettings = function(){
    var self = this;
    self.ARCsettings = {
          "innerRadius": -100,
          "outerRadius": -100
    };
  }

  BioCircos.prototype.genome_matrix = function(genome){
      var self = this;
      var i=self.genome.length;
      var genomeLabel = new Array();
      var genomeLength = new Array();
      for(var k=0;k<i;k++){
          genomeLabel[k]=self.genome[k][0];
      }
      for(var k=0;k<i;k++){
          genomeLength[k]=self.genome[k][1];
      }
      var i=genomeLength.length;
      var p=genomeLength.length;
      var genome = new Array();
      for(var k=0;k<i;k++){ 
         genome[k]=new Array();
           for(var j=0;j<p;j++){
              genome[k][j]=0;
           }
      }
      for(var k=0;k<i;k++){
         genome[k][0]=genomeLength[k];
      }
      self.genomeLabel = genomeLabel;
      self.genomeLength = genome;
  }

  BioCircos.prototype.heatmap_value_maxmin = function(heatmapIn){
      var self = this;
      var i=heatmapIn.length;
      var heatmapValueList = new Array();
      for(var k=0;k<i;k++){
          heatmapValueList[k]=heatmapIn[k].value;
      }
      Array.max=function(array){
          return Math.max.apply(Math,array);
      }
      Array.min=function(array){
         return Math.min.apply(Math,array);
      }
      var heatmapValueMax = Array.max(heatmapValueList);
      var heatmapValueMin = Array.min(heatmapValueList);
      var heatmapValueMaxmin = new Array();
      heatmapValueMaxmin[0]=heatmapValueMax;
      heatmapValueMaxmin[1]=heatmapValueMin;
      return heatmapValueMaxmin;
  }

  BioCircos.prototype.snp_value_maxmin = function(snpIn){
      var self = this;
      var i=snpIn.length;
      var snpValueList = new Array();
      for(var k=0;k<i;k++){
          snpValueList[k]=snpIn[k].value;
      }
      Array.max=function(array){
          return Math.max.apply(Math,array);
      }
      Array.min=function(array){
         return Math.min.apply(Math,array);
      }
      var snpValueMax = Array.max(snpValueList);
      var snpValueMin = Array.min(snpValueList);
      var snpValueMaxmin = new Array();
      snpValueMaxmin[0]=snpValueMax;
      snpValueMaxmin[1]=snpValueMin;
      return snpValueMaxmin;
  }

  BioCircos.prototype.cnv_value_maxmin = function(cnvIn){
      var self = this;
      var i=cnvIn.length;
      var cnvValueList = new Array();
      for(var k=0;k<i;k++){
          cnvValueList[k]=cnvIn[k].value;
      }
      Array.max=function(array){
          return Math.max.apply(Math,array);
      }
      Array.min=function(array){
         return Math.min.apply(Math,array);
      }
      var cnvValueMax = Array.max(cnvValueList);
      var cnvValueMin = Array.min(cnvValueList);
      var cnvValueMaxmin = new Array();
      cnvValueMaxmin[0]=cnvValueMax;
      cnvValueMaxmin[1]=cnvValueMin;
      return cnvValueMaxmin;
  }

  BioCircos.prototype.histogram_value_maxmin = function(histogramIn){
      var self = this;
      var i=histogramIn.length;
      var histogramValueList = new Array();
      for(var k=0;k<i;k++){
          histogramValueList[k]=histogramIn[k].value;
      }
      Array.max=function(array){
          return Math.max.apply(Math,array);
      }
      Array.min=function(array){
         return Math.min.apply(Math,array);
      }
      var histogramValueMax = Array.max(histogramValueList);
      var histogramValueMin = Array.min(histogramValueList);
      var histogramValueMaxmin = new Array();
      histogramValueMaxmin[0]=histogramValueMax;
      histogramValueMaxmin[1]=histogramValueMin;
      return histogramValueMaxmin;
  }

  BioCircos.prototype.line_value_maxmin = function(lineIn){
      var self = this;
      var i=lineIn.length;
      var lineValueList = new Array();
      for(var k=0;k<i;k++){
          lineValueList[k]=lineIn[k].value;
      }
      Array.max=function(array){
          return Math.max.apply(Math,array);
      }
      Array.min=function(array){
         return Math.min.apply(Math,array);
      }
      var lineValueMax = Array.max(lineValueList);
      var lineValueMin = Array.min(lineValueList);
      var lineValueMaxmin = new Array();
      lineValueMaxmin[0]=lineValueMax;
      lineValueMaxmin[1]=lineValueMin;
      return lineValueMaxmin;
  }

  BioCircos.prototype.draw_genome = function(genome){
    var self = this;

    var chord = d3.layout.chord()
      .padding(self.chrPad)
      .sortSubgroups(d3.descending)
      .matrix(genome);

    var width = self.svgWidth,
      height = self.svgHeight,
      innerRadius = self.innerRadius,
      outerRadius = self.outerRadius;

    var fill = d3.scale.ordinal()
        .domain(d3.range(4))
        .range(self.genomeFillColor);

    if(self.zoom == true){
        function zoom() {
            a=d3.event.translate[0]+width / 2
            b=d3.event.translate[1]+height / 2
            svg.attr("transform", "translate(" 
                + a +","+ b 
                + ")scale(" + d3.event.scale + ")");
        }
        var svg = d3.select(self.target).append("svg")
            .attr("width", width)
            .attr("height", height)
            .call(
                 d3.behavior.zoom()
                 .scaleExtent([0.9, 10])
                 .on("zoom", zoom)
            )
          .append("g")
            .attr("transform", "translate(" + width / 2 + "," + height / 2 + ")");

    }else{
        var svg = d3.select(self.target).append("svg")
            .attr("width", width)
            .attr("height", height)
          .append("g")
            .attr("transform", "translate(" + width / 2 + "," + height / 2 + ")");
    }

    if(self.genomeBorderDisplay == true){
        svg.append("g").selectAll("path")
            .data(chord.groups)
          .enter().append("path")
            .style("fill", function(d) { return fill(d.index); })
            .style("stroke", self.genomeBorderColor)
            .style("stroke-width", self.genomeBorderSize)
            .attr("d", d3.svg.arc().innerRadius(innerRadius).outerRadius(outerRadius))
            .attr("name", function(d) { return d.index+1; });
    }else{
        svg.append("g").selectAll("path")
            .data(chord.groups)
          .enter().append("path")
            .style("fill", function(d) { return fill(d.index); })
            .style("stroke", function(d) { return fill(d.index); })
            .attr("d", d3.svg.arc().innerRadius(innerRadius).outerRadius(outerRadius))
            .attr("name", function(d) { return d.index+1; });
    }

    if(self.genomeTextDisplay == true){
        svg.append("g").selectAll("text")
            .data(chord.groups)
          .enter().append("text")
            .style("fill", self.genomeTextColor)
            .style("font-size", self.genomeTextSize)
	    .each( function(d,i) { 
               d.angle = (d.startAngle + d.endAngle) / 2 - self.genomeTextDx;
               d.name = self.genomeLabel[i];
               d.rotation = self.genomeTextRotation;
               d.dy = self.genomeTextDy;
            })
      .attr("text-anchor", "middle") // Rotate around horizontal center of text box
      .attr("alignment-baseline", "middle") // Rotate around vertical center of text box
      .attr("vertical-align", "middle")
	    .attr("transform", function(d){
	       return "rotate(" + (d.angle * 180 / Math.PI ) + ")" +
	       "translate(0,"+ -1.0*(outerRadius+10+d.dy) +")" +
	       ( ( d.angle > Math.PI*2 && d.angle < Math.PI*0 ) ? "rotate(180)" : "") +
         "rotate(" + d.rotation +")";
	    })
	    .text(function(d){
	       return d.name;
	    });
    }

    if(self.ticksDisplay == true){
        function groupTicks(d) {
          var k = (d.endAngle - d.startAngle) / d.value;
          return d3.range(0, d.value, self.ticksScale).map(function(v, i) {
            return {
              angle: v * k + d.startAngle,
              label: v / self.ticksScale + ""
            };
          });
        }
		
        var ticks = svg.append("g").selectAll("g")
            .data(chord.groups)
          .enter().append("g").selectAll("g")
            .data(groupTicks)
          .enter().append("g")
            .attr("transform", function(d) {
              return "rotate(" + (d.angle * 180 / Math.PI - 90) + ")"
                  + "translate(" + (outerRadius - 0) + ",0)";
            });

        ticks.append("line")
            .attr("x1", 1)
            .attr("y1", 0)
            .attr("x2", self.ticksLength)
            .attr("y2", 0)
            .style("stroke", self.ticksColor);

        ticks.append("text")
            .attr("x", 8)
            .attr("dy", "0.35em")
            .style("font-size", self.ticksTextSize)
            .style("fill", self.ticksTextColor)
            .attr("transform", function(d) { return d.angle > Math.PI ? "rotate(180)translate(-16)" : null; })
            .style("text-anchor", function(d) { return d.angle > Math.PI ? "end" : null; })
            .text(function(d) { return d.label; });

    }

    var drag = d3.behavior.drag()
              .on("drag", dragmove);

    function dragmove(d) {
        d3.select(this)
          .attr("x", d3.event.x )
          .attr("y", d3.event.y );
    }

    //var draglinklabel = d3.behavior.drag()
    //          .on("drag", draglinkmove);

    //function draglinkmove(d) {
    //    d3.select(this)
    //      .attr("x", d3.event.x )
    //      .attr("y", d3.event.y );
    //}

    if(self.BACKGROUND.length > 0){
        for(var backgroundi=0; backgroundi<self.BACKGROUND.length; backgroundi++){
            self.update_BACKGROUNDsettings(self.BACKGROUNDConfig[backgroundi]);

            svg.append("g").selectAll("path")
                .data(chord.groups)
              .enter().append("path")
                .style("fill", self.BACKGROUNDsettings.BgFillColor)
                .style("stroke", self.BACKGROUNDsettings.BgborderColor)
                .style("stroke-width", self.BACKGROUNDsettings.BgborderSize)
                .attr("d", d3.svg.arc().innerRadius(self.BACKGROUNDsettings.BginnerRadius).outerRadius(self.BACKGROUNDsettings.BgouterRadius));

            if(self.BACKGROUNDsettings.axisShow=="true"){
                for(i=1;i<=self.BACKGROUNDsettings.axisNum;i++){
                    svg.append("g").selectAll("path")
                        .data(chord.groups)
                      .enter().append("path")
                        .style("fill", "none")
                        .style("opacity",self.BACKGROUNDsettings.axisOpacity)
                        .style("stroke", self.BACKGROUNDsettings.axisColor)
                        .style("stroke-width", self.BACKGROUNDsettings.axisWidth)
                        .attr("d", d3.svg.arc().innerRadius(self.BACKGROUNDsettings.BginnerRadius+(self.BACKGROUNDsettings.BgouterRadius-self.BACKGROUNDsettings.BginnerRadius)/(self.BACKGROUNDsettings.axisNum+1)*i).outerRadius(self.BACKGROUNDsettings.BginnerRadius+(self.BACKGROUNDsettings.BgouterRadius-self.BACKGROUNDsettings.BginnerRadius)/(self.BACKGROUNDsettings.axisNum+1)*i+self.BACKGROUNDsettings.axisWidth));
                }
            }

            self.init_BACKGROUNDsettings();

        }
    }

    if(self.ARC.length > 0){
            function BioCircosArc(d) {
              return self.ARC[arci].map(function(v, i) {
                var arc_k = (d[self.initGenome[v.chr]].endAngle - d[self.initGenome[v.chr]].startAngle) / d[self.initGenome[v.chr]].value;
                return {
                  startAngle: v.start * arc_k + d[self.initGenome[v.chr]].startAngle,
                  endAngle: v.end * arc_k + d[self.initGenome[v.chr]].startAngle,
                  arc_chr: v.chr,
                  arc_start: v.start,
                  arc_end: v.end,
                  arc_color: v.color,
                  arc_des: v.des,
                  arc_link: v.link,
                  arc_opacity: v.opacity,
                  arc_click_label: "arc"+arci+"_"+i,
                };
              });
            }
        for(var arci=0; arci<self.ARC.length; arci++){
            self.update_ARCsettings(self.ARCConfig[arci]);

            var arc_objects = BioCircosArc(chord.groups())

            var arc = d3.svg.arc()
                  .innerRadius(innerRadius+self.ARCsettings.innerRadius)
                  .outerRadius(outerRadius+self.ARCsettings.outerRadius);

            svg.append("g")
                .attr("class", "BioCircosARC")
                .selectAll("path.BioCircosARC")
                  .data(arc_objects)
                  .enter()
                .append("a")
                .attr("xlink:href", function(d){return d.arc_link})
                .append("path")
                .attr("class", "BioCircosARC")
                .attr("fill", function(d,i) { return d.arc_color; })
                .attr("d", function(d,i) { return arc(d,i); })
                .attr("opacity", function(d,i){ return d.arc_opacity; });
                if(self.settings.ARCMouseClickTextFromData=="first"){
                    svg.append("g")
                        .attr("class", "BioCircosARClabel")
                      .selectAll("text")
                        .data(arc_objects)
                        .enter().append("text")
                        .attr("class", "dragText")
                        .attr("id", function(d,i) { return "arc"+arci+"_"+i; })
                        .text(function(d) { return d.arc_chr; })
                        .attr("x", -1000)
                        .attr("y", -1000)
                        .style("opacity", 0)
                        .style("font-size", 1)
                        .attr("fill", self.ARCsettings.ARCFillColor);
                }
                if(self.settings.ARCMouseClickTextFromData=="second"){
                    svg.append("g")
                        .attr("class", "BioCircosARClabel")
                      .selectAll("text")
                        .data(arc_objects)
                        .enter().append("text")
                        .attr("class", "dragText")
                        .attr("id", function(d,i) { return "arc"+arci+"_"+i; })
                        .text(function(d) { return d.arc_start; })
                        .attr("x", -1000)
                        .attr("y", -1000)
                        .style("opacity", 0)
                        .style("font-size", 1)
                        .attr("fill", self.ARCsettings.ARCFillColor);
                }
                if(self.settings.ARCMouseClickTextFromData=="third"){
                    svg.append("g")
                        .attr("class", "BioCircosARClabel")
                      .selectAll("text")
                        .data(arc_objects)
                        .enter().append("text")
                        .attr("class", "dragText")
                        .attr("id", function(d,i) { return "arc"+arci+"_"+i; })
                        .text(function(d) { return d.arc_end; })
                        .attr("x", -1000)
                        .attr("y", -1000)
                        .style("opacity", 0)
                        .style("font-size", 1)
                        .attr("fill", self.ARCsettings.ARCFillColor);
                }
                if(self.settings.ARCMouseClickTextFromData=="fifth"){
                    svg.append("g")
                        .attr("class", "BioCircosARClabel")
                      .selectAll("text")
                        .data(arc_objects)
                        .enter().append("text")
                        .attr("class", "dragText")
                        .attr("id", function(d,i) { return "arc"+arci+"_"+i; })
                        .text(function(d) { return d.arc_des; })
                        .attr("x", -1000)
                        .attr("y", -1000)
                        .style("opacity", 0)
                        .style("font-size", 1)
                        .attr("fill", self.ARCsettings.ARCFillColor);
                }

            self.init_ARCsettings();

        }

        if(self.settings.ARCMouseEvent==true){
            var ARCMouseOnTooltip = d3.select("body")
                .append("div")
                .attr("class","BioCircosARCTooltip")
                .attr("id","BioCircosARCTooltip")
                .style("opacity",0);

            var ARCMouseOn = svg.selectAll("path.BioCircosARC");
            if(self.settings.ARCMouseOverDisplay==true){
                   ARCMouseOn.on("mouseover",function(d){
                       ARCMouseOnTooltip.html(self.settings.ARCMouseOverTooltipsHtml01+d.arc_chr+self.settings.ARCMouseOverTooltipsHtml02+d.arc_start+self.settings.ARCMouseOverTooltipsHtml03+d.arc_end+self.settings.ARCMouseOverTooltipsHtml04+d.arc_des+self.settings.ARCMouseOverTooltipsHtml05)
                              .style("left", (d3.event.pageX) + "px")
                              .style("top", (d3.event.pageY + 20) + "px")
                              .style("position", self.settings.ARCMouseOverTooltipsPosition)
                              .style("background-color", self.settings.ARCMouseOverTooltipsBackgroundColor)
                              .style("border-style", self.settings.ARCMouseOverTooltipsBorderStyle)
                              .style("border-width", self.settings.ARCMouseOverTooltipsBorderWidth)
                              .style("padding", self.settings.ARCMouseOverTooltipsPadding)
                              .style("border-radius", self.settings.ARCMouseOverTooltipsBorderRadius)
                              .style("opacity", self.settings.ARCMouseOverTooltipsOpacity)
                           d3.select(this)
                              .style("fill",  function(d,i) { if(self.settings.ARCMouseOverColor=="none"){return "";}else{return self.settings.ARCMouseOverColor;} })
                              .style("opacity",  function(d,i) { if(self.settings.ARCMouseOverArcOpacity=="none"){return "";}else{return self.settings.ARCMouseOverArcOpacity;} })
                              .style("stroke", function(d,i) { if(self.settings.ARCMouseOverArcStrokeColor=="none"){return "";}else{return self.settings.ARCMouseOverArcStrokeColor;} })
                              .style("stroke-width", function(d,i) { if(self.settings.ARCMouseOverArcStrokeWidth=="none"){return "";}else{return self.settings.ARCMouseOverArcStrokeWidth;} });
                   })
            }
            if(self.settings.ARCMouseClickDisplay==true){
                   ARCMouseOn.on("click",function(d){
                       d3.select(this)
                           .style("fill",  function(d,i) { if(self.settings.ARCMouseClickColor=="none"){return "";}else{return self.settings.ARCMouseClickColor;} })
                           .style("opacity",  function(d,i) { if(self.settings.ARCMouseClickArcOpacity=="none"){return "";}else{return self.settings.ARCMouseClickArcOpacity;} })
                           .style("stroke", function(d,i) { if(self.settings.ARCMouseClickArcStrokeColor=="none"){return "";}else{return self.settings.ARCMouseClickArcStrokeColor;} })
                           .style("stroke-width", function(d,i) { if(self.settings.ARCMouseClickArcStrokeWidth=="none"){return "";}else{return self.settings.ARCMouseClickArcStrokeWidth;} });
                       d3.select("#"+d.arc_click_label)
                           .style("opacity", self.settings.ARCMouseClickTextOpacity)
                           .style("fill", self.settings.ARCMouseClickTextColor)
                           .style("font-size", self.settings.ARCMouseClickTextSize)
                           .attr("x", d3.event.x - self.svgWidth/2 + self.settings.ARCMouseClickTextPostionX)
                           .attr("y", d3.event.y - self.svgHeight/2 + self.settings.ARCMouseClickTextPostionY);
                   })
            }

            if(self.settings.ARCMouseClickTextDrag==true){
                svg.selectAll("text.dragText").call(drag);
            }

            if(self.settings.ARCMouseDownDisplay==true){
                   ARCMouseOn.on("mousedown",function(d){
                      d3.select(this)
                          .style("fill",  function(d,i) { if(self.settings.ARCMouseDownColor=="none"){return "";}else{return self.settings.ARCMouseDownColor;} })
                          .style("opacity",  function(d,i) { if(self.settings.ARCMouseDownArcOpacity=="none"){return "";}else{return self.settings.ARCMouseDownArcOpacity;} })
                          .style("stroke", function(d,i) { if(self.settings.ARCMouseDownArcStrokeColor=="none"){return "";}else{return self.settings.ARCMouseDownArcStrokeColor;} })
                          .style("stroke-width", function(d,i) { if(self.settings.ARCMouseDownArcStrokeWidth=="none"){return "";}else{return self.settings.ARCMouseDownArcStrokeWidth;} });
                  })
            }

            if(self.settings.ARCMouseEnterDisplay==true){
                  ARCMouseOn.on("mouseenter",function(d){
                      d3.select(this)
                          .style("fill", function(d,i) { if(self.settings.ARCMouseEnterColor=="none"){return "";}else{return self.settings.ARCMouseEnterColor;} })
                          .style("opacity", function(d,i) { if(self.settings.ARCMouseEnterArcOpacity=="none"){return "";}else{return self.settings.ARCMouseEnterArcOpacity;} })
                          .style("stroke", function(d,i) { if(self.settings.ARCMouseEnterArcStrokeColor=="none"){return "";}else{return self.settings.ARCMouseEnterArcStrokeColor;} })
                          .style("stroke-width", function(d,i) { if(self.settings.ARCMouseEnterArcStrokeWidth=="none"){return "";}else{return self.settings.ARCMouseEnterArcStrokeWidth;} });
                  })
            }

            if(self.settings.ARCMouseLeaveDisplay==true){
                  ARCMouseOn.on("mouseleave",function(d){
                      ARCMouseOnTooltip.style("opacity",0.0);
                      d3.select(this)
                          .style("fill",  function(d,i) { if(self.settings.ARCMouseLeaveColor=="none"){return "";}else{return self.settings.ARCMouseLeaveColor;} })
                          .style("opacity",  function(d,i) { if(self.settings.ARCMouseLeaveArcOpacity=="none"){return "";}else{return self.settings.ARCMouseLeaveArcOpacity;} })
                          .style("stroke", function(d,i) { if(self.settings.ARCMouseLeaveArcStrokeColor=="none"){return "";}else{return self.settings.ARCMouseLeaveArcStrokeColor;} })
                          .style("stroke-width", function(d,i) { if(self.settings.ARCMouseLeaveArcStrokeWidth=="none"){return "";}else{return self.settings.ARCMouseLeaveArcStrokeWidth;} });
                  })
            }

            if(self.settings.ARCMouseUpDisplay==true){
                   ARCMouseOn.on("mouseup",function(d){
                       d3.select(this)
                          .style("fill",  function(d,i) { if(self.settings.ARCMouseUpColor=="none"){return "";}else{return self.settings.ARCMouseUpColor;} })
                          .style("opacity",  function(d,i) { if(self.settings.ARCMouseUpArcOpacity=="none"){return "";}else{return self.settings.ARCMouseUpArcOpacity;} })
                          .style("stroke", function(d,i) { if(self.settings.ARCMouseUpArcStrokeColor=="none"){return "";}else{return self.settings.ARCMouseUpArcStrokeColor;} })
                          .style("stroke-width", function(d,i) { if(self.settings.ARCMouseUpArcStrokeWidth=="none"){return "";}else{return self.settings.ARCMouseUpArcStrokeWidth;} });
                   })
            }

            if(self.settings.ARCMouseMoveDisplay==true){
                   ARCMouseOn.on("mousemove",function(d){
                       d3.select(this)
                          .style("fill",  function(d,i) { if(self.settings.ARCMouseMoveColor=="none"){return "";}else{return self.settings.ARCMouseMoveColor;} })
                          .style("opacity",  function(d,i) { if(self.settings.ARCMouseMoveArcOpacity=="none"){return "";}else{return self.settings.ARCMouseMoveArcOpacity;} })
                          .style("stroke", function(d,i) { if(self.settings.ARCMouseMoveArcStrokeColor=="none"){return "";}else{return self.settings.ARCMouseMoveArcStrokeColor;} })
                          .style("stroke-width", function(d,i) { if(self.settings.ARCMouseMoveArcStrokeWidth=="none"){return "";}else{return self.settings.ARCMouseMoveArcStrokeWidth;} });
                       ARCMouseOnTooltip.style("left", (d3.event.pageX) + "px")
                       .style("top", (d3.event.pageY + 20) + "px");
                   })
            }

            if(self.settings.ARCMouseOutDisplay==true){
                   ARCMouseOn.on("mouseout",function(d){
                       ARCMouseOnTooltip.style("opacity",0.0);
                       d3.select(this)
                           .transition()
                           .duration(self.settings.ARCMouseOutAnimationTime)
                          .style("fill",  function(d,i) { if(self.settings.ARCMouseOutColor=="none"){return "";}else{return self.settings.ARCMouseOutColor;} })
                          .style("opacity",  function(d,i) { if(self.settings.ARCMouseOutArcOpacity=="none"){return "";}else{return self.settings.ARCMouseOutArcOpacity;} })
                          .style("stroke", function(d,i) { if(self.settings.ARCMouseOutArcStrokeColor=="none"){return "";}else{return self.settings.ARCMouseOutArcStrokeColor;} })
                          .style("stroke-width", function(d,i) { if(self.settings.ARCMouseOutArcStrokeWidth=="none"){return "";}else{return self.settings.ARCMouseOutArcStrokeWidth;} });
                   });
            }
        }

    }

    if(self.HISTOGRAM.length > 0){
            function BioCircosHISTOGRAM(d) {
              return self.HISTOGRAM[histogrami].map(function(v, i) {
                var histogram_k = (d[self.initGenome[v.chr]].endAngle - d[self.initGenome[v.chr]].startAngle) / d[self.initGenome[v.chr]].value;
                return {
                  startAngle: v.start * histogram_k + d[self.initGenome[v.chr]].startAngle,
                  endAngle: v.end * histogram_k + d[self.initGenome[v.chr]].startAngle,
                  histogram_chr: v.chr,
                  histogram_start: v.start,
                  histogram_end: v.end,
                  histogram_name: v.name,
                  histogram_link: v.link,
                  histogram_value: v.value
                };
              });
            }
        for(var histogrami=0; histogrami<self.HISTOGRAM.length; histogrami++){
            self.update_HISTOGRAMsettings(self.HISTOGRAMConfig[histogrami]);

            var histogram_objects = BioCircosHISTOGRAM(chord.groups())
            
            if(self.HISTOGRAMsettings.range){
              var histogram_value_maxmin_instance = self.HISTOGRAMsettings.range;
              histogram_value_maxmin_instance.reverse();
            }
            else{
              var histogram_value_maxmin_instance = self.histogram_value_maxmin(self.HISTOGRAM[histogrami]);
            }
            
            svg.append("g")
                .attr("class", "BioCircosHISTOGRAM")
                .selectAll("path.BioCircosHISTOGRAM")
                  .data(histogram_objects)
                  .enter()
                .append("a")
                .attr("xlink:href", function(d){return d.histogram_link})
                .append("path")
                .attr("class", "BioCircosHISTOGRAM")
                .attr("fill", self.HISTOGRAMsettings.histogramFillColor)
                .attr("d", d3.svg.arc().innerRadius(self.HISTOGRAMsettings.minRadius).outerRadius(function(d) {return self.HISTOGRAMsettings.minRadius + ((d.histogram_value-histogram_value_maxmin_instance[1])*(self.HISTOGRAMsettings.maxRadius-self.HISTOGRAMsettings.minRadius)/( histogram_value_maxmin_instance[0] - histogram_value_maxmin_instance[1]));}));
            self.init_HISTOGRAMsettings();

        }

        if(self.settings.HISTOGRAMMouseEvent==true){
            var HISTOGRAMMouseOnTooltip = d3.select("body")
                .append("div")
                .attr("class","BioCircosHISTOGRAMTooltip")
                .attr("id","BioCircosHISTOGRAMTooltip")
                .style("opacity",0);

            var HISTOGRAMMouseOn = svg.selectAll("path.BioCircosHISTOGRAM");

            if(self.settings.HISTOGRAMMouseOverDisplay==true){
                HISTOGRAMMouseOn.on("mouseover",function(d){
                      HISTOGRAMMouseOnTooltip.html(self.settings.HISTOGRAMMouseOverTooltipsHtml01+d.histogram_chr+self.settings.HISTOGRAMMouseOverTooltipsHtml02+d.histogram_start+self.settings.HISTOGRAMMouseOverTooltipsHtml03+d.histogram_end+self.settings.HISTOGRAMMouseOverTooltipsHtml04+d.histogram_name+self.settings.HISTOGRAMMouseOverTooltipsHtml05+d.histogram_value+self.settings.HISTOGRAMMouseOverTooltipsHtml06)
                       .style("left", (d3.event.pageX) + "px")
                       .style("top", (d3.event.pageY + 20) + "px")
                       .style("position", self.settings.HISTOGRAMMouseOverTooltipsPosition)
                       .style("background-color", self.settings.HISTOGRAMMouseOverTooltipsBackgroundColor)
                       .style("border-style", self.settings.HISTOGRAMMouseOverTooltipsBorderStyle)
                       .style("border-width", self.settings.HISTOGRAMMouseOverTooltipsBorderWidth)
                       .style("padding", self.settings.HISTOGRAMMouseOverTooltipsPadding)
                       .style("border-radius", self.settings.HISTOGRAMMouseOverTooltipsBorderRadius)
                       .style("opacity", self.settings.HISTOGRAMMouseOverTooltipsOpacity)
                    d3.select(this)
                       .style("fill",  function(d,i) { if(self.settings.HISTOGRAMMouseOverColor=="none"){return "";}else{return self.settings.HISTOGRAMMouseOverColor;} })
                       .style("opacity",  function(d,i) { if(self.settings.HISTOGRAMMouseOverOpacity=="none"){return "";}else{return self.settings.HISTOGRAMMouseOverOpacity;} })
                       .style("stroke", function(d,i) { if(self.settings.HISTOGRAMMouseOverStrokeColor=="none"){return "";}else{return self.settings.HISTOGRAMMouseOverStrokeColor;} })
                       .style("stroke-width", function(d,i) { if(self.settings.HISTOGRAMMouseOverStrokeWidth=="none"){return "";}else{return self.settings.HISTOGRAMMouseOverStrokeWidth;} });
                })
            }
            if(self.settings.HISTOGRAMMouseClickDisplay==true){
                HISTOGRAMMouseOn.on("click",function(d){
                    d3.select(this)
                       .style("fill",  function(d,i) { if(self.settings.HISTOGRAMMouseClickColor=="none"){return "";}else{return self.settings.HISTOGRAMMouseClickColor;} })
                       .style("opacity",  function(d,i) { if(self.settings.HISTOGRAMMouseClickOpacity=="none"){return "";}else{return self.settings.HISTOGRAMMouseClickOpacity;} })
                       .style("stroke", function(d,i) { if(self.settings.HISTOGRAMMouseClickStrokeColor=="none"){return "";}else{return self.settings.HISTOGRAMMouseClickStrokeColor;} })
                       .style("stroke-width", function(d,i) { if(self.settings.HISTOGRAMMouseClickStrokeWidth=="none"){return "";}else{return self.settings.HISTOGRAMMouseClickStrokeWidth;} });
                })
            }
            if(self.settings.HISTOGRAMMouseDownDisplay==true){
               HISTOGRAMMouseOn.on("mousedown",function(d){
                   d3.select(this)
                       .style("fill",  function(d,i) { if(self.settings.HISTOGRAMMouseDownColor=="none"){return "";}else{return self.settings.HISTOGRAMMouseDownColor;} })
                       .style("opacity",  function(d,i) { if(self.settings.HISTOGRAMMouseDownOpacity=="none"){return "";}else{return self.settings.HISTOGRAMMouseDownOpacity;} })
                       .style("stroke", function(d,i) { if(self.settings.HISTOGRAMMouseDownStrokeColor=="none"){return "";}else{return self.settings.HISTOGRAMMouseDownStrokeColor;} })
                       .style("stroke-width", function(d,i) { if(self.settings.HISTOGRAMMouseDownStrokeWidth=="none"){return "";}else{return self.settings.HISTOGRAMMouseDownStrokeWidth;} });
               })
            }
            if(self.settings.HISTOGRAMMouseEnterDisplay==true){
               HISTOGRAMMouseOn.on("mouseenter",function(d){
                   d3.select(this)
                       .style("fill",  function(d,i) { if(self.settings.HISTOGRAMMouseEnterColor=="none"){return "";}else{return self.settings.HISTOGRAMMouseEnterColor;} })
                       .style("opacity",  function(d,i) { if(self.settings.HISTOGRAMMouseEnterOpacity=="none"){return "";}else{return self.settings.HISTOGRAMMouseEnterOpacity;} })
                       .style("stroke", function(d,i) { if(self.settings.HISTOGRAMMouseEnterStrokeColor=="none"){return "";}else{return self.settings.HISTOGRAMMouseEnterStrokeColor;} })
                       .style("stroke-width", function(d,i) { if(self.settings.HISTOGRAMMouseEnterStrokeWidth=="none"){return "";}else{return self.settings.HISTOGRAMMouseEnterStrokeWidth;} });
               })
            }
            if(self.settings.HISTOGRAMMouseLeaveDisplay==true){
               HISTOGRAMMouseOn.on("mouseleave",function(d){
                   HISTOGRAMMouseOnTooltip.style("opacity",0.0);
                   d3.select(this)
                       .style("fill",  function(d,i) { if(self.settings.HISTOGRAMMouseLeaveColor=="none"){return "";}else{return self.settings.HISTOGRAMMouseLeaveColor;} })
                       .style("opacity",  function(d,i) { if(self.settings.HISTOGRAMMouseLeaveOpacity=="none"){return "";}else{return self.settings.HISTOGRAMMouseLeaveOpacity;} })
                       .style("stroke", function(d,i) { if(self.settings.HISTOGRAMMouseLeaveStrokeColor=="none"){return "";}else{return self.settings.HISTOGRAMMouseLeaveStrokeColor;} })
                       .style("stroke-width", function(d,i) { if(self.settings.HISTOGRAMMouseLeaveStrokeWidth=="none"){return "";}else{return self.settings.HISTOGRAMMouseLeaveStrokeWidth;} });
               })
            }
            if(self.settings.HISTOGRAMMouseUpDisplay==true){
               HISTOGRAMMouseOn.on("mouseup",function(d){
                   d3.select(this)
                       .style("fill",  function(d,i) { if(self.settings.HISTOGRAMMouseUpColor=="none"){return "";}else{return self.settings.HISTOGRAMMouseUpColor;} })
                       .style("opacity",  function(d,i) { if(self.settings.HISTOGRAMMouseUpOpacity=="none"){return "";}else{return self.settings.HISTOGRAMMouseUpOpacity;} })
                       .style("stroke", function(d,i) { if(self.settings.HISTOGRAMMouseUpStrokeColor=="none"){return "";}else{return self.settings.HISTOGRAMMouseUpStrokeColor;} })
                       .style("stroke-width", function(d,i) { if(self.settings.HISTOGRAMMouseUpStrokeWidth=="none"){return "";}else{return self.settings.HISTOGRAMMouseUpStrokeWidth;} });
               })
            }
            if(self.settings.HISTOGRAMMouseMoveDisplay==true){
               HISTOGRAMMouseOn.on("mousemove",function(d){
                   d3.select(this)
                       .style("fill",  function(d,i) { if(self.settings.HISTOGRAMMouseMoveColor=="none"){return "";}else{return self.settings.HISTOGRAMMouseMoveColor;} })
                       .style("opacity",  function(d,i) { if(self.settings.HISTOGRAMMouseMoveOpacity=="none"){return "";}else{return self.settings.HISTOGRAMMouseMoveOpacity;} })
                       .style("stroke", function(d,i) { if(self.settings.HISTOGRAMMouseMoveStrokeColor=="none"){return "";}else{return self.settings.HISTOGRAMMouseMoveStrokeColor;} })
                       .style("stroke-width", function(d,i) { if(self.settings.HISTOGRAMMouseMoveStrokeWidth=="none"){return "";}else{return self.settings.HISTOGRAMMouseMoveStrokeWidth;} });
                   HISTOGRAMMouseOnTooltip.style("left", (d3.event.pageX) + "px")
                   .style("top", (d3.event.pageY + 20) + "px");
               })
            }
            if(self.settings.HISTOGRAMMouseOutDisplay==true){
               HISTOGRAMMouseOn.on("mouseout",function(d){
                   HISTOGRAMMouseOnTooltip.style("opacity",0.0);
                   d3.select(this)
                       .transition()
                       .duration(self.settings.HISTOGRAMMouseOutAnimationTime)
                       .style("fill",  function(d,i) { if(self.settings.HISTOGRAMMouseOutColor=="none"){return "";}else{return self.settings.HISTOGRAMMouseOutColor;} })
                       .style("opacity",  function(d,i) { if(self.settings.HISTOGRAMMouseOutOpacity=="none"){return "";}else{return self.settings.HISTOGRAMMouseOutOpacity;} })
                       .style("stroke", function(d,i) { if(self.settings.HISTOGRAMMouseOutStrokeColor=="none"){return "";}else{return self.settings.HISTOGRAMMouseOutStrokeColor;} })
                       .style("stroke-width", function(d,i) { if(self.settings.HISTOGRAMMouseOutStrokeWidth=="none"){return "";}else{return self.settings.HISTOGRAMMouseOutStrokeWidth;} });
               });
            }

        }

    }

    if(self.LINE.length > 0){
            function BioCircosLINE(d) {
              return self.LINE[linei].map(function(v, i) {
                var line_k = (d[self.initGenome[v.chr]].endAngle - d[self.initGenome[v.chr]].startAngle) / d[self.initGenome[v.chr]].value;
                return {
                  line_angle: v.pos * line_k + d[self.initGenome[v.chr]].startAngle,
                  line_chr: v.chr,
                  line_pos: v.pos,
                  line_des: v.des,
                  line_value: v.value,
                  line_color: self.LINEsettings.LineColor,
                  line_width: self.LINEsettings.LineWidth,
                  x: (0 + Math.sin(v.pos * line_k + d[self.initGenome[v.chr]].startAngle) * ((self.LINEsettings.minRadius + ( (v.value-self.line_value_maxmin(self.LINE[linei])[1])/(self.line_value_maxmin(self.LINE[linei])[0]-self.line_value_maxmin(self.LINE[linei])[1])*(self.LINEsettings.maxRadius-self.LINEsettings.minRadius) )))),
                  y: (0 - Math.cos(v.pos * line_k + d[self.initGenome[v.chr]].startAngle) * ((self.LINEsettings.minRadius + ( (v.value-self.line_value_maxmin(self.LINE[linei])[1])/(self.line_value_maxmin(self.LINE[linei])[0]-self.line_value_maxmin(self.LINE[linei])[1])*(self.LINEsettings.maxRadius-self.LINEsettings.minRadius) ))))
                };
              });
            }
        for(var linei=0; linei<self.LINE.length; linei++){
            self.update_LINEsettings(self.LINEConfig[linei]);

            var line_objects = BioCircosLINE(chord.groups())

            for(var chri=0; chri<self.genomeLabel.length; chri++){
                var line_objects_a_chr = line_objects.filter(function(element,pos){return element.line_chr==self.genomeLabel[chri]})
                if(line_objects_a_chr.length != 0){
                    svg.append("g")
                        .attr("class", "BioCircosLINE")
                        .append("path")
                          .datum(line_objects_a_chr)
                        .attr("class", "BioCircosLINE")
                        .attr("fill", "none")
                        .attr("stroke",self.LINEsettings.LineColor)
                        .attr("stroke-width",self.LINEsettings.LineWidth)
                        .attr("d",  d3.svg.line()
                            .x(function(d) { return d.x; })
                            .y(function(d) { return d.y; })
                        );
                }
            }
            self.init_LINEsettings();

        }

        if(self.settings.LINEMouseEvent==true){
            var LINEMouseOnTooltip = d3.select("body")
                .append("div")
                .attr("class","BioCircosLINETooltip")
                .attr("id","BioCircosLINETooltip")
                .style("opacity",0);

            var LINEMouseOn = svg.selectAll("path.BioCircosLINE");

            if(self.settings.LINEMouseOverDisplay==true){
                LINEMouseOn.on("mouseover",function(d){
                      LINEMouseOnTooltip.html(self.settings.LINEMouseOverTooltipsHtml01)
                       .style("left", (d3.event.pageX) + "px")
                       .style("top", (d3.event.pageY + 20) + "px")
                       .style("position", self.settings.LINEMouseOverTooltipsPosition)
                       .style("background-color", self.settings.LINEMouseOverTooltipsBackgroundColor)
                       .style("border-style", self.settings.LINEMouseOverTooltipsBorderStyle)
                       .style("border-width", self.settings.LINEMouseOverTooltipsBorderWidth)
                       .style("padding", self.settings.LINEMouseOverTooltipsPadding)
                       .style("border-radius", self.settings.LINEMouseOverTooltipsBorderRadius)
                       .style("opacity", self.settings.LINEMouseOverTooltipsOpacity)
                    d3.select(this)
                       .style("opacity",  function(d,i) { if(self.settings.LINEMouseOverLineOpacity=="none"){return "";}else{return self.settings.LINEMouseOverLineOpacity;} })
                       .style("stroke", function(d,i) { if(self.settings.LINEMouseOverLineStrokeColor=="none"){return "";}else{return self.settings.LINEMouseOverLineStrokeColor;} })
                       .style("stroke-width", function(d,i) { if(self.settings.LINEMouseOverLineStrokeWidth=="none"){return "";}else{return self.settings.LINEMouseOverLineStrokeWidth;} });
                })
            }
            if(self.settings.LINEMouseClickDisplay==true){
                LINEMouseOn.on("click",function(d){
                    d3.select(this)
                       .style("opacity",  function(d,i) { if(self.settings.LINEMouseClickLineOpacity=="none"){return "";}else{return self.settings.LINEMouseClickLineOpacity;} })
                       .style("stroke", function(d,i) { if(self.settings.LINEMouseClickLineStrokeColor=="none"){return "";}else{return self.settings.LINEMouseClickLineStrokeColor;} })
                       .style("stroke-width", function(d,i) { if(self.settings.LINEMouseClickLineStrokeWidth=="none"){return "";}else{return self.settings.LINEMouseClickLineStrokeWidth;} });
                })
            }
            if(self.settings.LINEMouseDownDisplay==true){
               LINEMouseOn.on("mousedown",function(d){
                   d3.select(this)
                       .style("opacity",  function(d,i) { if(self.settings.LINEMouseDownLineOpacity=="none"){return "";}else{return self.settings.LINEMouseDownLineOpacity;} })
                       .style("stroke", function(d,i) { if(self.settings.LINEMouseDownLineStrokeColor=="none"){return "";}else{return self.settings.LINEMouseDownLineStrokeColor;} })
                       .style("stroke-width", function(d,i) { if(self.settings.LINEMouseDownLineStrokeWidth=="none"){return "";}else{return self.settings.LINEMouseDownLineStrokeWidth;} });
               })
            }
            if(self.settings.LINEMouseEnterDisplay==true){
               LINEMouseOn.on("mouseenter",function(d){
                   d3.select(this)
                       .style("opacity",  function(d,i) { if(self.settings.LINEMouseEnterLineOpacity=="none"){return "";}else{return self.settings.LINEMouseEnterLineOpacity;} })
                       .style("stroke", function(d,i) { if(self.settings.LINEMouseEnterLineStrokeColor=="none"){return "";}else{return self.settings.LINEMouseEnterLineStrokeColor;} })
                       .style("stroke-width", function(d,i) { if(self.settings.LINEMouseEnterLineStrokeWidth=="none"){return "";}else{return self.settings.LINEMouseEnterLineStrokeWidth;} });
               })
            }
            if(self.settings.LINEMouseLeaveDisplay==true){
               LINEMouseOn.on("mouseleave",function(d){
                   LINEMouseOnTooltip.style("opacity",0.0);
                   d3.select(this)
                       .style("opacity",  function(d,i) { if(self.settings.LINEMouseLeaveLineOpacity=="none"){return "";}else{return self.settings.LINEMouseLeaveLineOpacity;} })
                       .style("stroke", function(d,i) { if(self.settings.LINEMouseLeaveLineStrokeColor=="none"){return "";}else{return self.settings.LINEMouseLeaveLineStrokeColor;} })
                       .style("stroke-width", function(d,i) { if(self.settings.LINEMouseLeaveLineStrokeWidth=="none"){return "";}else{return self.settings.LINEMouseLeaveLineStrokeWidth;} });
               })
            }
            if(self.settings.LINEMouseUpDisplay==true){
               LINEMouseOn.on("mouseup",function(d){
                   d3.select(this)
                       .style("opacity",  function(d,i) { if(self.settings.LINEMouseUpLineOpacity=="none"){return "";}else{return self.settings.LINEMouseUpLineOpacity;} })
                       .style("stroke", function(d,i) { if(self.settings.LINEMouseUpLineStrokeColor=="none"){return "";}else{return self.settings.LINEMouseUpLineStrokeColor;} })
                       .style("stroke-width", function(d,i) { if(self.settings.LINEMouseUpLineStrokeWidth=="none"){return "";}else{return self.settings.LINEMouseUpLineStrokeWidth;} });
               })
            }
            if(self.settings.LINEMouseMoveDisplay==true){
               LINEMouseOn.on("mousemove",function(d){
                   d3.select(this)
                       .style("opacity",  function(d,i) { if(self.settings.LINEMouseMoveLineOpacity=="none"){return "";}else{return self.settings.LINEMouseMoveLineOpacity;} })
                       .style("stroke", function(d,i) { if(self.settings.LINEMouseMoveLineStrokeColor=="none"){return "";}else{return self.settings.LINEMouseMoveLineStrokeColor;} })
                       .style("stroke-width", function(d,i) { if(self.settings.LINEMouseMoveLineStrokeWidth=="none"){return "";}else{return self.settings.LINEMouseMoveLineStrokeWidth;} });
                   LINEMouseOnTooltip.style("left", (d3.event.pageX) + "px")
                   .style("top", (d3.event.pageY + 20) + "px");
               })
            }
            if(self.settings.LINEMouseOutDisplay==true){
               LINEMouseOn.on("mouseout",function(d){
                   LINEMouseOnTooltip.style("opacity",0.0);
                   d3.select(this)
                       .transition()
                       .duration(self.settings.LINEMouseOutAnimationTime)
                       .style("opacity",  function(d,i) { if(self.settings.LINEMouseOutLineOpacity=="none"){return "";}else{return self.settings.LINEMouseOutLineOpacity;} })
                       .style("stroke", function(d,i) { if(self.settings.LINEMouseOutLineStrokeColor=="none"){return "";}else{return self.settings.LINEMouseOutLineStrokeColor;} })
                       .style("stroke-width", function(d,i) { if(self.settings.LINEMouseOutLineStrokeWidth=="none"){return "";}else{return self.settings.LINEMouseOutLineStrokeWidth;} });
               });
            }

        }

    }

    if(self.TEXT.length > 0){
        for(var texti=0; texti<self.TEXT.length; texti++){
            self.update_TEXTsettings(self.TEXTConfig[texti]);
                svg.append("text")
                 .attr("x", self.TEXTsettings.x)
                 .attr("y", self.TEXTsettings.y)
                 .style("opacity", self.TEXTsettings.textOpacity)
                 .style("font-size", self.TEXTsettings.textSize)
                 .style("font-weight", self.TEXTsettings.textWeight) //normal,bold,bolder,lighter,100,200,300,400,500,600,700,800,900
                 .attr("fill", self.TEXTsettings.textColor)
                 .attr("class", "dragText")
                 .text(self.TEXTsettings.text);
            self.init_TEXTsettings();
        }
        if(self.settings.TEXTModuleDragEvent==true){
            svg.selectAll("text.dragText").call(drag);
        }

    }

    if(self.CNV.length > 0){
            function BioCircosCnv(d) {
              return self.CNV[cnvi].map(function(v, i) {
                var cnv_k = (d[self.initGenome[v.chr]].endAngle - d[self.initGenome[v.chr]].startAngle) / d[self.initGenome[v.chr]].value;
                // Check if a range is given
                if(self.CNVsettings.range){
                  var cnv_value_maxmin_instance = self.CNVsettings.range;
                }
                else{
                  var cnv_value_maxmin_instance = self.cnv_value_maxmin(self.CNV[cnvi]);
                }
                return {
                  startAngle: v.start * cnv_k + d[self.initGenome[v.chr]].startAngle,
                  endAngle: v.end * cnv_k + d[self.initGenome[v.chr]].startAngle,
                  cnv_chr: v.chr,
                  cnv_start: v.start,
                  cnv_end: v.end,
                  cnv_val: v.value,
                  cnv_link: v.link,
                  cnv_click_label: "cnv"+cnvi+"_"+i,
                  cnv_deviation: (v.value-cnv_value_maxmin_instance[1])/(cnv_value_maxmin_instance[0]-cnv_value_maxmin_instance[1])*(self.CNVsettings.maxRadius-self.CNVsettings.minRadius)

                };
              });
            }
        for(var cnvi=0; cnvi<self.CNV.length; cnvi++){
            self.update_CNVsettings(self.CNVConfig[cnvi]);

            var cnv_objects = BioCircosCnv(chord.groups())

            svg.append("g")
                .attr("class", "BioCircosCNV")
                .selectAll("path.BioCircosCNV")
                  .data(cnv_objects)
                  .enter()
                .append("a")
                .attr("xlink:href", function(d){return d.cnv_link})
                .append("path")
                .attr("class", "BioCircosCNV")
                .attr("transform", function(d) {
                  return "rotate(0)"
                      + "translate(0,0)";
                })
                .attr("fill", function(d,i) { return self.CNVsettings.CNVColor; })
                .attr("d", function(d,i) { var cnv = d3.svg.arc().innerRadius(self.CNVsettings.minRadius+d.cnv_deviation).outerRadius(self.CNVsettings.minRadius+self.CNVsettings.CNVwidth+d.cnv_deviation); return cnv(d,i); });
                if(self.settings.CNVMouseClickTextFromData=="first"){
                    svg.append("g")
                        .attr("class", "BioCircosCNVlabel")
                      .selectAll("text")
                        .data(cnv_objects)
                        .enter().append("text")
                        .attr("class", "dragText")
                        .attr("id", function(d,i) { return "cnv"+cnvi+"_"+i; })
                        .text(function(d) { return d.cnv_chr; })
                        .attr("x", 0)
                        .attr("y", 0)
                        .style("opacity", 0)
                        .style("font-size", 1)
                        .attr("fill", self.CNVsettings.CNVFillColor);
                }
                if(self.settings.CNVMouseClickTextFromData=="second"){
                    svg.append("g")
                        .attr("class", "BioCircosCNVlabel")
                      .selectAll("text")
                        .data(cnv_objects)
                        .enter().append("text")
                        .attr("class", "dragText")
                        .attr("id", function(d,i) { return "cnv"+cnvi+"_"+i; })
                        .text(function(d) { return d.cnv_start; })
                        .attr("x", 0)
                        .attr("y", 0)
                        .style("opacity", 0)
                        .style("font-size", 1)
                        .attr("fill", self.CNVsettings.CNVFillColor);
                }
                if(self.settings.CNVMouseClickTextFromData=="third"){
                    svg.append("g")
                        .attr("class", "BioCircosCNVlabel")
                      .selectAll("text")
                        .data(cnv_objects)
                        .enter().append("text")
                        .attr("class", "dragText")
                        .attr("id", function(d,i) { return "cnv"+cnvi+"_"+i; })
                        .text(function(d) { return d.cnv_end; })
                        .attr("x", 0)
                        .attr("y", 0)
                        .style("opacity", 0)
                        .style("font-size", 1)
                        .attr("fill", self.CNVsettings.CNVFillColor);
                }
                if(self.settings.CNVMouseClickTextFromData=="fourth"){
                    svg.append("g")
                        .attr("class", "BioCircosCNVlabel")
                      .selectAll("text")
                        .data(cnv_objects)
                        .enter().append("text")
                        .attr("class", "dragText")
                        .attr("id", function(d,i) { return "cnv"+cnvi+"_"+i; })
                        .text(function(d) { return d.cnv_val; })
                        .attr("x", 0)
                        .attr("y", 0)
                        .style("opacity", 0)
                        .style("font-size", 1)
                        .attr("fill", self.CNVsettings.CNVFillColor);
                }

            self.init_CNVsettings();

        }

        if(self.settings.CNVMouseEvent==true){
            var CNVMouseOnTooltip = d3.select("body")
                .append("div")
                .attr("class","BioCircosCNVTooltip")
                .attr("id","BioCircosCNVTooltip")
                .style("opacity",0);

            var CNVMouseOn = svg.selectAll("path.BioCircosCNV");

            if(self.settings.CNVMouseOverDisplay==true){
                CNVMouseOn.on("mouseover",function(d){
                      CNVMouseOnTooltip.html(self.settings.CNVMouseOverTooltipsHtml01+d.cnv_chr+self.settings.CNVMouseOverTooltipsHtml02+d.cnv_start+self.settings.CNVMouseOverTooltipsHtml03+d.cnv_end+self.settings.CNVMouseOverTooltipsHtml04+d.cnv_val+self.settings.CNVMouseOverTooltipsHtml05)
                       .style("left", (d3.event.pageX) + "px")
                       .style("top", (d3.event.pageY + 20) + "px")
                       .style("position", self.settings.CNVMouseOverTooltipsPosition)
                       .style("background-color", self.settings.CNVMouseOverTooltipsBackgroundColor)
                       .style("border-style", self.settings.CNVMouseOverTooltipsBorderStyle)
                       .style("border-width", self.settings.CNVMouseOverTooltipsBorderWidth)
                       .style("padding", self.settings.CNVMouseOverTooltipsPadding)
                       .style("border-radius", self.settings.CNVMouseOverTooltipsBorderRadius)
                       .style("opacity", self.settings.CNVMouseOverTooltipsOpacity)
                    d3.select(this)
                       .style("fill",  function(d,i) { if(self.settings.CNVMouseOverColor=="none"){return "";}else{return self.settings.CNVMouseOverColor;} })
                       .style("opacity",  function(d,i) { if(self.settings.CNVMouseOverArcOpacity=="none"){return "";}else{return self.settings.CNVMouseOverArcOpacity;} })
                       .style("stroke", function(d,i) { if(self.settings.CNVMouseOverArcStrokeColor=="none"){return "";}else{return self.settings.CNVMouseOverArcStrokeColor;} })
                       .style("stroke-width", function(d,i) { if(self.settings.CNVMouseOverArcStrokeWidth=="none"){return "";}else{return self.settings.CNVMouseOverArcStrokeWidth;} });
                })
            }

            if(self.settings.CNVMouseClickDisplay==true){
                CNVMouseOn.on("click",function(d){
                    d3.select(this)
                       .style("fill",  function(d,i) { if(self.settings.CNVMouseClickColor=="none"){return "";}else{return self.settings.CNVMouseClickColor;} })
                       .style("opacity",  function(d,i) { if(self.settings.CNVMouseClickArcOpacity=="none"){return "";}else{return self.settings.CNVMouseClickArcOpacity;} })
                       .style("stroke", function(d,i) { if(self.settings.CNVMouseClickArcStrokeColor=="none"){return "";}else{return self.settings.CNVMouseClickArcStrokeColor;} })
                       .style("stroke-width", function(d,i) { if(self.settings.CNVMouseClickArcStrokeWidth=="none"){return "";}else{return self.settings.CNVMouseClickArcStrokeWidth;} });
                    d3.select("#"+d.cnv_click_label)
                        .style("opacity", self.settings.CNVMouseClickTextOpacity)
                        .style("fill", self.settings.CNVMouseClickTextColor)
                        .style("font-size", self.settings.CNVMouseClickTextSize)
                        .attr("x", d3.event.x - self.svgWidth/2 + self.settings.ARCMouseClickTextPostionX)
                        .attr("y", d3.event.y - self.svgHeight/2 + self.settings.ARCMouseClickTextPostionY);
                })
            }

            if(self.settings.CNVMouseClickTextDrag==true){
                svg.selectAll("text.dragText").call(drag);
            }
            if(self.settings.CNVMouseDownDisplay==true){
               CNVMouseOn.on("mousedown",function(d){
                   d3.select(this)
                       .style("fill",  function(d,i) { if(self.settings.CNVMouseDownColor=="none"){return "";}else{return self.settings.CNVMouseDownColor;} })
                       .style("opacity",  function(d,i) { if(self.settings.CNVMouseDownCircleOpacity=="none"){return "";}else{return self.settings.CNVMouseDownCircleOpacity;} })
                       .style("stroke", function(d,i) { if(self.settings.CNVMouseDownCircleStrokeColor=="none"){return "";}else{return self.settings.CNVMouseDownCircleStrokeColor;} })
                       .style("stroke-width", function(d,i) { if(self.settings.CNVMouseDownCircleStrokeWidth=="none"){return "";}else{return self.settings.CNVMouseDownCircleStrokeWidth;} });
               })
            }
            if(self.settings.CNVMouseEnterDisplay==true){
               CNVMouseOn.on("mouseenter",function(d){
                   d3.select(this)
                       .style("fill",  function(d,i) { if(self.settings.CNVMouseEnterColor=="none"){return "";}else{return self.settings.CNVMouseEnterColor;} })
                       .style("opacity",  function(d,i) { if(self.settings.CNVMouseEnterCircleOpacity=="none"){return "";}else{return self.settings.CNVMouseEnterCircleOpacity;} })
                       .style("stroke", function(d,i) { if(self.settings.CNVMouseEnterCircleStrokeColor=="none"){return "";}else{return self.settings.CNVMouseEnterCircleStrokeColor;} })
                       .style("stroke-width", function(d,i) { if(self.settings.CNVMouseEnterCircleStrokeWidth=="none"){return "";}else{return self.settings.CNVMouseEnterCircleStrokeWidth;} });
               })
            }
            if(self.settings.CNVMouseLeaveDisplay==true){
               CNVMouseOn.on("mouseleave",function(d){
                   CNVMouseOnTooltip.style("opacity",0.0);
                   d3.select(this)
                       .style("fill",  function(d,i) { if(self.settings.CNVMouseLeaveColor=="none"){return "";}else{return self.settings.CNVMouseLeaveColor;} })
                       .style("opacity",  function(d,i) { if(self.settings.CNVMouseLeaveCircleOpacity=="none"){return "";}else{return self.settings.CNVMouseLeaveCircleOpacity;} })
                       .style("stroke", function(d,i) { if(self.settings.CNVMouseLeaveCircleStrokeColor=="none"){return "";}else{return self.settings.CNVMouseLeaveCircleStrokeColor;} })
                       .style("stroke-width", function(d,i) { if(self.settings.CNVMouseLeaveCircleStrokeWidth=="none"){return "";}else{return self.settings.CNVMouseLeaveCircleStrokeWidth;} });
               })
            }
            if(self.settings.CNVMouseUpDisplay==true){
               CNVMouseOn.on("mouseup",function(d){
                   d3.select(this)
                       .style("fill",  function(d,i) { if(self.settings.CNVMouseUpColor=="none"){return "";}else{return self.settings.CNVMouseUpColor;} })
                       .style("opacity",  function(d,i) { if(self.settings.CNVMouseUpCircleOpacity=="none"){return "";}else{return self.settings.CNVMouseUpCircleOpacity;} })
                       .style("stroke", function(d,i) { if(self.settings.CNVMouseUpCircleStrokeColor=="none"){return "";}else{return self.settings.CNVMouseUpCircleStrokeColor;} })
                       .style("stroke-width", function(d,i) { if(self.settings.CNVMouseUpCircleStrokeWidth=="none"){return "";}else{return self.settings.CNVMouseUpCircleStrokeWidth;} });
               })
            }
            if(self.settings.CNVMouseMoveDisplay==true){
               CNVMouseOn.on("mousemove",function(d){
                   d3.select(this)
                       .style("fill",  function(d,i) { if(self.settings.CNVMouseMoveColor=="none"){return "";}else{return self.settings.CNVMouseMoveColor;} })
                       .style("opacity",  function(d,i) { if(self.settings.CNVMouseMoveCircleOpacity=="none"){return "";}else{return self.settings.CNVMouseMoveCircleOpacity;} })
                       .style("stroke", function(d,i) { if(self.settings.CNVMouseMoveCircleStrokeColor=="none"){return "";}else{return self.settings.CNVMouseMoveCircleStrokeColor;} })
                       .style("stroke-width", function(d,i) { if(self.settings.CNVMouseMoveCircleStrokeWidth=="none"){return "";}else{return self.settings.CNVMouseMoveCircleStrokeWidth;} });
                   CNVMouseOnTooltip.style("left", (d3.event.pageX) + "px")
                   .style("top", (d3.event.pageY + 20) + "px");
               })
            }
            if(self.settings.CNVMouseOutDisplay==true){
               CNVMouseOn.on("mouseout",function(d){
                   CNVMouseOnTooltip.style("opacity",0.0);
                   d3.select(this)
                       .transition()
                       .duration(self.settings.CNVMouseOutAnimationTime)
                       .style("fill",  function(d,i) { if(self.settings.CNVMouseOutColor=="none"){return "";}else{return self.settings.CNVMouseOutColor;} })
                       .style("opacity",  function(d,i) { if(self.settings.CNVMouseOutCircleOpacity=="none"){return "";}else{return self.settings.CNVMouseOutCircleOpacity;} })
                       .style("stroke", function(d,i) { if(self.settings.CNVMouseOutCircleStrokeColor=="none"){return "";}else{return self.settings.CNVMouseOutCircleStrokeColor;} })
                       .style("stroke-width", function(d,i) { if(self.settings.CNVMouseOutCircleStrokeWidth=="none"){return "";}else{return self.settings.CNVMouseOutCircleStrokeWidth;} });
               });
            }
        }

    }

    if(self.HEATMAP.length > 0){
            function BioCircosHeatmap(d) {
              return self.HEATMAP[heatmapi].map(function(v, i) {
                var heatmap_k = (d[self.initGenome[v.chr]].endAngle - d[self.initGenome[v.chr]].startAngle) / d[self.initGenome[v.chr]].value;
                return {
                  startAngle: v.start * heatmap_k + d[self.initGenome[v.chr]].startAngle,
                  endAngle: v.end * heatmap_k + d[self.initGenome[v.chr]].startAngle,
                  heatmap_chr: v.chr,
                  heatmap_start: v.start,
                  heatmap_end: v.end,
                  heatmap_name: v.name,
                  heatmap_value: v.value,
                };
              });
            }
        for(var heatmapi=0; heatmapi<self.HEATMAP.length; heatmapi++){
            self.update_HEATMAPsettings(self.HEATMAPConfig[heatmapi]);

            var heatmap_objects = BioCircosHeatmap(chord.groups())

            var HeatmapMaxColor = d3.rgb(self.HEATMAPsettings.maxColor);
            var HeatmapMinColor = d3.rgb(self.HEATMAPsettings.minColor);
            var HeatmapValue2Color = d3.interpolate(HeatmapMinColor,HeatmapMaxColor);

            if(self.HEATMAPsettings.range){
              var heatmap_value_maxmin_instance = self.HEATMAPsettings.range;
              heatmap_value_maxmin_instance.reverse();
            }
            else{
              var heatmap_value_maxmin_instance = self.heatmap_value_maxmin(self.HEATMAP[heatmapi]);
            }

            var heatmap = d3.svg.arc().innerRadius(innerRadius+self.HEATMAPsettings.innerRadius).outerRadius(outerRadius+self.HEATMAPsettings.outerRadius);
            svg.append("g")
                .attr("class", "BioCircosHEATMAP")
                .selectAll("path.BioCircosHEATMAP")
                  .data(heatmap_objects)
                  .enter()
                .append("path")
                .attr("class", "BioCircosHEATMAP")
                .attr("fill", function(d,i) { return HeatmapValue2Color((d.heatmap_value - heatmap_value_maxmin_instance[1])/(heatmap_value_maxmin_instance[0]-heatmap_value_maxmin_instance[1])); })
                .attr("d", function(d,i) { return heatmap(d,i); });
            self.init_HEATMAPsettings();

        }

        if(self.settings.HEATMAPMouseEvent==true){
            var HEATMAPMouseOnTooltip = d3.select("body")
                .append("div")
                .attr("class","BioCircosHeatmapTooltip")
                .attr("id","BioCircosHeatmapTooltip")
                .style("opacity",0);

            var HEATMAPMouseOn = svg.selectAll("path.BioCircosHEATMAP");

            if(self.settings.HEATMAPMouseOverDisplay==true){
                HEATMAPMouseOn.on("mouseover",function(d){
                      HEATMAPMouseOnTooltip.html(self.settings.HEATMAPMouseOverTooltipsHtml01+d.heatmap_chr+self.settings.HEATMAPMouseOverTooltipsHtml02+d.heatmap_start+self.settings.HEATMAPMouseOverTooltipsHtml03+d.heatmap_end+self.settings.HEATMAPMouseOverTooltipsHtml04+d.heatmap_name+self.settings.HEATMAPMouseOverTooltipsHtml05+d.heatmap_value+self.settings.HEATMAPMouseOverTooltipsHtml06)
                       .style("left", (d3.event.pageX) + "px")
                       .style("top", (d3.event.pageY + 20) + "px")
                       .style("position", self.settings.HEATMAPMouseOverTooltipsPosition)
                       .style("background-color", self.settings.HEATMAPMouseOverTooltipsBackgroundColor)
                       .style("border-style", self.settings.HEATMAPMouseOverTooltipsBorderStyle)
                       .style("border-width", self.settings.HEATMAPMouseOverTooltipsBorderWidth)
                       .style("padding", self.settings.HEATMAPMouseOverTooltipsPadding)
                       .style("border-radius", self.settings.HEATMAPMouseOverTooltipsBorderRadius)
                       .style("opacity", self.settings.HEATMAPMouseOverTooltipsOpacity)
                    d3.select(this)
                       .style("fill",  function(d,i) { if(self.settings.HEATMAPMouseOverColor=="none"){return "";}else{return self.settings.HEATMAPMouseOverColor;} })
                       .style("opacity",  function(d,i) { if(self.settings.HEATMAPMouseOverOpacity=="none"){return "";}else{return self.settings.HEATMAPMouseOverOpacity;} })
                       .style("stroke", function(d,i) { if(self.settings.HEATMAPMouseOverStrokeColor=="none"){return "";}else{return self.settings.HEATMAPMouseOverStrokeColor;} })
                       .style("stroke-width", function(d,i) { if(self.settings.HEATMAPMouseOverStrokeWidth=="none"){return "";}else{return self.settings.HEATMAPMouseOverStrokeWidth;} });
                })
            }
            if(self.settings.HEATMAPMouseClickDisplay==true){
                HEATMAPMouseOn.on("click",function(d){
                    d3.select(this)
                       .style("fill",  function(d,i) { if(self.settings.HEATMAPMouseClickColor=="none"){return "";}else{return self.settings.HEATMAPMouseClickColor;} })
                       .style("opacity",  function(d,i) { if(self.settings.HEATMAPMouseClickOpacity=="none"){return "";}else{return self.settings.HEATMAPMouseClickOpacity;} })
                       .style("stroke", function(d,i) { if(self.settings.HEATMAPMouseClickStrokeColor=="none"){return "";}else{return self.settings.HEATMAPMouseClickStrokeColor;} })
                       .style("stroke-width", function(d,i) { if(self.settings.HEATMAPMouseClickStrokeWidth=="none"){return "";}else{return self.settings.HEATMAPMouseClickStrokeWidth;} });
                })
            }
            if(self.settings.HEATMAPMouseDownDisplay==true){
               HEATMAPMouseOn.on("mousedown",function(d){
                   d3.select(this)
                       .style("fill",  function(d,i) { if(self.settings.HEATMAPMouseDownColor=="none"){return "";}else{return self.settings.HEATMAPMouseDownColor;} })
                       .style("opacity",  function(d,i) { if(self.settings.HEATMAPMouseDownOpacity=="none"){return "";}else{return self.settings.HEATMAPMouseDownOpacity;} })
                       .style("stroke", function(d,i) { if(self.settings.HEATMAPMouseDownStrokeColor=="none"){return "";}else{return self.settings.HEATMAPMouseDownStrokeColor;} })
                       .style("stroke-width", function(d,i) { if(self.settings.HEATMAPMouseDownStrokeWidth=="none"){return "";}else{return self.settings.HEATMAPMouseDownStrokeWidth;} });
               })
            }
            if(self.settings.HEATMAPMouseEnterDisplay==true){
               HEATMAPMouseOn.on("mouseenter",function(d){
                   d3.select(this)
                       .style("fill",  function(d,i) { if(self.settings.HEATMAPMouseEnterColor=="none"){return "";}else{return self.settings.HEATMAPMouseEnterColor;} })
                       .style("opacity",  function(d,i) { if(self.settings.HEATMAPMouseEnterOpacity=="none"){return "";}else{return self.settings.HEATMAPMouseEnterOpacity;} })
                       .style("stroke", function(d,i) { if(self.settings.HEATMAPMouseEnterStrokeColor=="none"){return "";}else{return self.settings.HEATMAPMouseEnterStrokeColor;} })
                       .style("stroke-width", function(d,i) { if(self.settings.HEATMAPMouseEnterStrokeWidth=="none"){return "";}else{return self.settings.HEATMAPMouseEnterStrokeWidth;} });
               })
            }
            if(self.settings.HEATMAPMouseLeaveDisplay==true){
               HEATMAPMouseOn.on("mouseleave",function(d){
                   HEATMAPMouseOnTooltip.style("opacity",0.0);
                   d3.select(this)
                       .style("fill",  function(d,i) { if(self.settings.HEATMAPMouseLeaveColor=="none"){return "";}else{return self.settings.HEATMAPMouseLeaveColor;} })
                       .style("opacity",  function(d,i) { if(self.settings.HEATMAPMouseLeaveOpacity=="none"){return "";}else{return self.settings.HEATMAPMouseLeaveOpacity;} })
                       .style("stroke", function(d,i) { if(self.settings.HEATMAPMouseLeaveStrokeColor=="none"){return "";}else{return self.settings.HEATMAPMouseLeaveStrokeColor;} })
                       .style("stroke-width", function(d,i) { if(self.settings.HEATMAPMouseLeaveStrokeWidth=="none"){return "";}else{return self.settings.HEATMAPMouseLeaveStrokeWidth;} });
               })
            }
            if(self.settings.HEATMAPMouseUpDisplay==true){
               HEATMAPMouseOn.on("mouseup",function(d){
                   d3.select(this)
                       .style("fill",  function(d,i) { if(self.settings.HEATMAPMouseUpColor=="none"){return "";}else{return self.settings.HEATMAPMouseUpColor;} })
                       .style("opacity",  function(d,i) { if(self.settings.HEATMAPMouseUpOpacity=="none"){return "";}else{return self.settings.HEATMAPMouseUpOpacity;} })
                       .style("stroke", function(d,i) { if(self.settings.HEATMAPMouseUpStrokeColor=="none"){return "";}else{return self.settings.HEATMAPMouseUpStrokeColor;} })
                       .style("stroke-width", function(d,i) { if(self.settings.HEATMAPMouseUpStrokeWidth=="none"){return "";}else{return self.settings.HEATMAPMouseUpStrokeWidth;} });
               })
            }
            if(self.settings.HEATMAPMouseMoveDisplay==true){
               HEATMAPMouseOn.on("mousemove",function(d){
                   d3.select(this)
                       .style("fill",  function(d,i) { if(self.settings.HEATMAPMouseMoveColor=="none"){return "";}else{return self.settings.HEATMAPMouseMoveColor;} })
                       .style("opacity",  function(d,i) { if(self.settings.HEATMAPMouseMoveOpacity=="none"){return "";}else{return self.settings.HEATMAPMouseMoveOpacity;} })
                       .style("stroke", function(d,i) { if(self.settings.HEATMAPMouseMoveStrokeColor=="none"){return "";}else{return self.settings.HEATMAPMouseMoveStrokeColor;} })
                       .style("stroke-width", function(d,i) { if(self.settings.HEATMAPMouseUpStrokeWidth=="none"){return "";}else{return self.settings.HEATMAPMouseMoveStrokeWidth;} });
                   HEATMAPMouseOnTooltip.style("left", (d3.event.pageX) + "px")
                   .style("top", (d3.event.pageY + 20) + "px");
               })
            }
            if(self.settings.HEATMAPMouseOutDisplay==true){
               HEATMAPMouseOn.on("mouseout",function(d){
                   HEATMAPMouseOnTooltip.style("opacity",0.0);
                   d3.select(this)
                       .transition()
                       .duration(self.settings.HEATMAPMouseOutAnimationTime)
                       .style("fill",  function(d,i) { if(self.settings.HEATMAPMouseOutColor=="none"){return "";}else{return self.settings.HEATMAPMouseOutColor;} })
                       .style("opacity",  function(d,i) { if(self.settings.HEATMAPMouseOutOpacity=="none"){return "";}else{return self.settings.HEATMAPMouseOutOpacity;} })
                       .style("stroke", function(d,i) { if(self.settings.HEATMAPMouseOutStrokeColor=="none"){return "";}else{return self.settings.HEATMAPMouseOutStrokeColor;} })
                       .style("stroke-width", function(d,i) { if(self.settings.HEATMAPMouseUpStrokeWidth=="none"){return "";}else{return self.settings.HEATMAPMouseOutStrokeWidth;} });
               });
            }

        }

    }

    if(self.SCATTER.length > 0){
            function BioCircosSCATTER(d) {
              return self.SCATTER[scatteri].map(function(v, i) {
                var random_data = Math.random()*self.SCATTERsettings.random_data
                var scatter_k = (d[self.initGenome[v.chr]].endAngle - d[self.initGenome[v.chr]].startAngle) / d[self.initGenome[v.chr]].value;
                return {
                  scatter_angle: (v.start/2+v.end/2) * scatter_k + d[self.initGenome[v.chr]].startAngle,
                  scatter_chr: v.chr,
                  scatter_start: v.start,
                  scatter_end: v.end,
                  scatter_name: v.name,
                  scatter_des: v.des,
                  scatter_link: v.link,
                  scatter_click_label: "scatter"+scatteri+"_"+i,
                  x: (0 + Math.sin((v.start/2+v.end/2) * scatter_k + d[self.initGenome[v.chr]].startAngle) * (self.SCATTERsettings.SCATTERRadius - random_data)),
                  y: (0 - Math.cos((v.start/2+v.end/2) * scatter_k + d[self.initGenome[v.chr]].startAngle) * (self.SCATTERsettings.SCATTERRadius - random_data))
                };
              });
            }
        for(var scatteri=0; scatteri<self.SCATTER.length; scatteri++){
            self.update_SCATTERsettings(self.SCATTERConfig[scatteri]);
            var scatter_objects = BioCircosSCATTER(chord.groups())

            if(self.SCATTERsettings.outerPointType=="circle"){
              if(self.SCATTERsettings.SCATTERAnimationDisplay==false){
                svg.append("g")
                    .attr("class", "BioCircosSCATTER")
                  .selectAll("circle")
                    .data(scatter_objects)
                    .enter()
                    .append("a")
                    .attr("xlink:href", function(d){return d.scatter_link})
                    .append("circle")
                    .attr("id", "BioCircosSCATTEROut")
                    .attr("fill", self.SCATTERsettings.outerCircleColor)
                    .attr("opacity", self.SCATTERsettings.outerCircleOpacity)
                    .attr("r", self.SCATTERsettings.outerCircleSize)
                    .attr("cx", function(d) { return d.x; })
                    .attr("cy", function(d) { return d.y; });
              }
              if(self.SCATTERsettings.SCATTERAnimationDisplay==true){
                svg.append("g")
                    .attr("class", "BioCircosSCATTER")
                  .selectAll("circle")
                    .data(scatter_objects)
                    .enter()
                    .append("a")
                    .attr("xlink:href", function(d){return d.scatter_link})
                    .append("circle")
                    .attr("id", "BioCircosSCATTEROut")
                    .attr("fill", self.SCATTERsettings.outerCircleColor)
                    .attr("opacity", self.SCATTERsettings.outerCircleOpacity)
                    .attr("r", self.SCATTERsettings.outerCircleSize)
		    .attr("cx",function(d){
			    return 0;
		    })
		    .attr("cy",function(d){
			    return 0;
		    })
		    .transition()
		    .delay(function(d,i){
			    return i * self.SCATTERsettings.SCATTERAnimationDelay;
		    })
		    .duration(self.SCATTERsettings.SCATTERAnimationTime)
		    .ease(self.SCATTERsettings.SCATTERAnimationType)
                    .attr("cx", function(d) { return d.x; })
                    .attr("cy", function(d) { return d.y; });
               }

                if(self.settings.SCATTERMouseClickTextFromData=="first"){
                    svg.append("g")
                        .attr("class", "BioCircosSCATTERlabel")
                      .selectAll("text")
                        .data(scatter_objects)
                        .enter().append("text")
                        .attr("class", "dragText")
                        .attr("id", function(d,i) { return "scatter"+scatteri+"_"+i; })
                        .text(function(d) { return d.scatter_chr; })
                        .attr("x", -1000)
                        .attr("y", -1000)
                        .style("opacity", 0)
                        .style("font-size", 1)
                        .attr("fill", self.SCATTERsettings.SCATTERFillColor);
                }
                if(self.settings.SCATTERMouseClickTextFromData=="second"){
                    svg.append("g")
                        .attr("class", "BioCircosSCATTERlabel")
                      .selectAll("text")
                        .data(scatter_objects)
                        .enter().append("text")
                        .attr("class", "dragText")
                        .attr("id", function(d,i) { return "scatter"+scatteri+"_"+i; })
                        .text(function(d) { return d.scatter_pos; })
                        .attr("x", -1000)
                        .attr("y", -1000)
                        .style("opacity", 0)
                        .style("font-size", 1)
                        .attr("fill", self.SCATTERsettings.SCATTERFillColor);
                }
                if(self.settings.SCATTERMouseClickTextFromData=="third"){
                    svg.append("g")
                        .attr("class", "BioCircosSCATTERlabel")
                      .selectAll("text")
                        .data(scatter_objects)
                        .enter().append("text")
                        .attr("class", "dragText")
                        .attr("id", function(d,i) { return "scatter"+scatteri+"_"+i; })
                        .text(function(d) { return d.scatter_val; })
                        .attr("x", -1000)
                        .attr("y", -1000)
                        .style("opacity", 0)
                        .style("font-size", 1)
                        .attr("fill", self.SCATTERsettings.SCATTERFillColor);
                }
                if(self.settings.SCATTERMouseClickTextFromData=="fourth"){
                    svg.append("g")
                        .attr("class", "BioCircosSCATTERlabel")
                      .selectAll("text")
                        .data(scatter_objects)
                        .enter().append("text")
                        .attr("class", "dragText")
                        .attr("id", function(d,i) { return "scatter"+scatteri+"_"+i; })
                        .text(function(d) { return d.scatter_name; })
                        .attr("x", -1000)
                        .attr("y", -1000)
                        .style("opacity", 0)
                        .style("font-size", 1)
                        .attr("fill", self.SCATTERsettings.SCATTERFillColor);
                }
                if(self.settings.SCATTERMouseClickTextFromData=="fifth"){
                    svg.append("g")
                        .attr("class", "BioCircosSCATTERlabel")
                      .selectAll("text")
                        .data(scatter_objects)
                        .enter().append("text")
                        .attr("class", "dragText")
                        .attr("id", function(d,i) { return "scatter"+scatteri+"_"+i; })
                        .text(function(d) { return d.scatter_des; })
                        .attr("x", -1000)
                        .attr("y", -1000)
                        .style("opacity", 0)
                        .style("font-size", 1)
                        .attr("fill", self.SCATTERsettings.SCATTERFillColor);
                }
            }

            if(self.SCATTERsettings.outerPointType=="rect"){
                svg.append("g")
                    .attr("class", "BioCircosSCATTER")
                  .selectAll("rect")
                    .data(scatter_objects)
                    .enter()
                    .append("a")
                    .attr("xlink:href", function(d){return d.scatter_link})
                    .append("rect")
                    .attr("id", "BioCircosSCATTEROut")
                    .attr("x", function(d) { return d.x - self.SCATTERsettings.outerrectWidth/2; })
                    .attr("y", function(d) { return d.y - self.SCATTERsettings.outerrectHeight/2; })
                    .attr("fill", self.SCATTERsettings.outerCircleColor)
                    .attr("width", self.SCATTERsettings.outerrectWidth)
                    .attr("height", self.SCATTERsettings.outerrectHeight)
                    .attr("opacity", 0.5);
            }

            if(self.SCATTERsettings.innerPointType=="circle"){
              if(self.SCATTERsettings.SCATTERAnimationDisplay==false){
                svg.append("g")
                    .attr("class", "BioCircosSCATTEROut")
                  .selectAll("circle")
                    .data(scatter_objects)
                    .enter()
                    .append("a")
                    .attr("xlink:href", function(d){return d.scatter_link})
                    .append("circle")
                    .attr("id", "BioCircosSCATTEROut")  //out
                    .attr("fill", self.SCATTERsettings.innerCircleColor)
                    .attr("r", self.SCATTERsettings.innerCircleSize)
                    .attr("cx", function(d) { return d.x; })
                    .attr("cy", function(d) { return d.y; });
               }
              if(self.SCATTERsettings.SCATTERAnimationDisplay==true){
                svg.append("g")
                    .attr("class", "BioCircosSCATTEROut")
                  .selectAll("circle")
                    .data(scatter_objects)
                    .enter()
                    .append("a")
                    .attr("xlink:href", function(d){return d.scatter_link})
                    .append("circle")
                    .attr("id", "BioCircosSCATTEROut")  //out
                    .attr("fill", self.SCATTERsettings.innerCircleColor)
                    .attr("r", self.SCATTERsettings.innerCircleSize)
		    .attr("cx",function(d){
			    return 0;
		    })
		    .attr("cy",function(d){
			    return 0;
		    })
		    .transition()
		    .delay(function(d,i){
			    return i * self.SCATTERsettings.SCATTERAnimationDelay;
		    })
		    .duration(self.SCATTERsettings.SCATTERAnimationTime)
		    .ease(self.SCATTERsettings.SCATTERAnimationType)
                    .attr("cx", function(d) { return d.x; })
                    .attr("cy", function(d) { return d.y; });
               }
                if(self.settings.SCATTERMouseClickTextFromData=="first"){
                    svg.append("g")
                        .attr("class", "BioCircosSCATTERlabel")
                      .selectAll("text")
                        .data(scatter_objects)
                        .enter().append("text")
                        .attr("class", "dragText")
                        .attr("id", function(d,i) { return "scatter"+scatteri+"_"+i; })
                        .text(function(d) { return d.scatter_chr; })
                        .attr("x", -1000)
                        .attr("y", -1000)
                        .style("opacity", 0)
                        .style("font-size", 1)
                        .attr("fill", self.SCATTERsettings.SCATTERFillColor);
                }
                if(self.settings.SCATTERMouseClickTextFromData=="second"){
                    svg.append("g")
                        .attr("class", "BioCircosSCATTERlabel")
                      .selectAll("text")
                        .data(scatter_objects)
                        .enter().append("text")
                        .attr("class", "dragText")
                        .attr("id", function(d,i) { return "scatter"+scatteri+"_"+i; })
                        .text(function(d) { return d.scatter_start; })
                        .attr("x", -1000)
                        .attr("y", -1000)
                        .style("opacity", 0)
                        .style("font-size", 1)
                        .attr("fill", self.SCATTERsettings.SCATTERFillColor);
                }
                if(self.settings.SCATTERMouseClickTextFromData=="third"){
                    svg.append("g")
                        .attr("class", "BioCircosSCATTERlabel")
                      .selectAll("text")
                        .data(scatter_objects)
                        .enter().append("text")
                        .attr("class", "dragText")
                        .attr("id", function(d,i) { return "scatter"+scatteri+"_"+i; })
                        .text(function(d) { return d.scatter_end; })
                        .attr("x", -1000)
                        .attr("y", -1000)
                        .style("opacity", 0)
                        .style("font-size", 1)
                        .attr("fill", self.SCATTERsettings.SCATTERFillColor);
                }
                if(self.settings.SCATTERMouseClickTextFromData=="fourth"){
                    svg.append("g")
                        .attr("class", "BioCircosSCATTERlabel")
                      .selectAll("text")
                        .data(scatter_objects)
                        .enter().append("text")
                        .attr("class", "dragText")
                        .attr("id", function(d,i) { return "scatter"+scatteri+"_"+i; })
                        .text(function(d) { return d.scatter_name; })
                        .attr("x", -1000)
                        .attr("y", -1000)
                        .style("opacity", 0)
                        .style("font-size", 1)
                        .attr("fill", self.SCATTERsettings.SCATTERFillColor);
                }
                if(self.settings.SCATTERMouseClickTextFromData=="fifth"){
                    svg.append("g")
                        .attr("class", "BioCircosSCATTERlabel")
                      .selectAll("text")
                        .data(scatter_objects)
                        .enter().append("text")
                        .attr("class", "dragText")
                        .attr("id", function(d,i) { return "scatter"+scatteri+"_"+i; })
                        .text(function(d) { return d.scatter_des; })
                        .attr("x", -1000)
                        .attr("y", -1000)
                        .style("opacity", 0)
                        .style("font-size", 1)
                        .attr("fill", self.SCATTERsettings.SCATTERFillColor);
                }
            }

            if(self.SCATTERsettings.innerPointType=="rect"){
                svg.append("g")
                    .attr("class", "BioCircosSCATTEROut")
                  .selectAll("rect")
                    .data(scatter_objects)
                    .enter()
                    .append("a")
                    .attr("xlink:href", function(d){return d.scatter_link})
                    .append("rect")
                    .attr("id", "BioCircosSCATTEROut")  //out
                    .attr("x", function(d) { return d.x - self.SCATTERsettings.innerrectWidth/2; })
                    .attr("y", function(d) { return d.y - self.SCATTERsettings.innerrectHeight/2; })
                    .attr("width", self.SCATTERsettings.innerrectWidth)
                    .attr("height", self.SCATTERsettings.innerrectHeight)
                    .attr("fill", self.SCATTERsettings.innerCircleColor);
            }

            self.init_SCATTERsettings();

        }

        if(self.settings.SCATTERMouseEvent==true){
            var SCATTERMouseOnTooltip = d3.select("body")
                .append("div")
                .attr("class","BioCircosSCATTERTooltip")
                .attr("id","BioCircosSCATTERTooltip")
                .style("opacity",0);

            var SCATTERMouseOn = svg.selectAll("#BioCircosSCATTEROut");

            if(self.settings.SCATTERMouseOverDisplay==true){
                SCATTERMouseOn.on("mouseover",function(d){
                      SCATTERMouseOnTooltip.html(self.settings.SCATTERMouseOverTooltipsHtml01+d.scatter_chr+self.settings.SCATTERMouseOverTooltipsHtml02+d.scatter_start+self.settings.SCATTERMouseOverTooltipsHtml03+d.scatter_end+self.settings.SCATTERMouseOverTooltipsHtml04+d.scatter_name+self.settings.SCATTERMouseOverTooltipsHtml05+d.scatter_des+self.settings.SCATTERMouseOverTooltipsHtml06)
                       .style("left", (d3.event.pageX) + "px")
                       .style("top", (d3.event.pageY + 20) + "px")
                       .style("position", self.settings.SCATTERMouseOverTooltipsPosition)
                       .style("background-color", self.settings.SCATTERMouseOverTooltipsBackgroundColor)
                       .style("border-style", self.settings.SCATTERMouseOverTooltipsBorderStyle)
                       .style("border-width", self.settings.SCATTERMouseOverTooltipsBorderWidth)
                       .style("padding", self.settings.SCATTERMouseOverTooltipsPadding)
                       .style("border-radius", self.settings.SCATTERMouseOverTooltipsBorderRadius)
                       .style("opacity", self.settings.SCATTERMouseOverTooltipsOpacity)
                    d3.select(this)
                       .style("r",  function(d,i) { if(self.settings.SCATTERMouseOverCircleSize=="none"){return "";}else{return self.settings.SCATTERMouseOverCircleSize;} })
                       .style("fill",  function(d,i) { if(self.settings.SCATTERMouseOverColor=="none"){return "";}else{return self.settings.SCATTERMouseOverColor;} })
                       .style("opacity",  function(d,i) { if(self.settings.SCATTERMouseOverCircleOpacity=="none"){return "";}else{return self.settings.SCATTERMouseOverCircleOpacity;} })
                       .style("stroke", function(d,i) { if(self.settings.SCATTERMouseOverCircleStrokeColor=="none"){return "";}else{return self.settings.SCATTERMouseOverCircleStrokeColor;} })
                       .style("stroke-width", function(d,i) { if(self.settings.SCATTERMouseOverCircleStrokeWidth=="none"){return "";}else{return self.settings.SCATTERMouseOverCircleStrokeWidth;} });
                })
            }
            if(self.settings.SCATTERMouseClickDisplay==true){
                SCATTERMouseOn.on("click",function(d){
                    d3.select(this)
                       .style("r",  function(d,i) { if(self.settings.SCATTERMouseClickCircleSize=="none"){return "";}else{return self.settings.SCATTERMouseClickCircleSize;} })
                       .style("fill",  function(d,i) { if(self.settings.SCATTERMouseClickColor=="none"){return "";}else{return self.settings.SCATTERMouseClickColor;} })
                       .style("opacity",  function(d,i) { if(self.settings.SCATTERMouseClickCircleOpacity=="none"){return "";}else{return self.settings.SCATTERMouseClickCircleOpacity;} })
                       .style("stroke", function(d,i) { if(self.settings.SCATTERMouseClickCircleStrokeColor=="none"){return "";}else{return self.settings.SCATTERMouseClickCircleStrokeColor;} })
                       .style("stroke-width", function(d,i) { if(self.settings.SCATTERMouseClickCircleStrokeWidth=="none"){return "";}else{return self.settings.SCATTERMouseClickCircleStrokeWidth;} });
                    d3.select("#"+d.scatter_click_label)
                        .style("opacity", self.settings.SCATTERMouseClickTextOpacity)
                        .style("fill", self.settings.SCATTERMouseClickTextColor)
                        .style("font-size", self.settings.SCATTERMouseClickTextSize)
                        .attr("x", d.x+self.settings.SCATTERMouseClickTextPostionX)
                        .attr("y", d.y+self.settings.SCATTERMouseClickTextPostionY);
                })
            }
            if(self.settings.SCATTERMouseClickTextDrag==true){
                svg.selectAll("text.dragText").call(drag);
            }
            if(self.settings.SCATTERMouseDownDisplay==true){
               SCATTERMouseOn.on("mousedown",function(d){
                   d3.select(this)
                       .style("r", function(d,i) { if(self.settings.SCATTERMouseDownCircleSize=="none"){return "";}else{return self.settings.SCATTERMouseDownCircleSize;} })
                       .style("fill", function(d,i) { if(self.settings.SCATTERMouseDownColor=="none"){return "";}else{return self.settings.SCATTERMouseDownColor;} })
                       .style("opacity",  function(d,i) { if(self.settings.SCATTERMouseDownCircleOpacity=="none"){return "";}else{return self.settings.SCATTERMouseDownCircleOpacity;} })
                       .style("stroke", function(d,i) { if(self.settings.SCATTERMouseDownCircleStrokeColor=="none"){return "";}else{return self.settings.SCATTERMouseDownCircleStrokeColor;} })
                       .style("stroke-width", function(d,i) { if(self.settings.SCATTERMouseDownCircleStrokeWidth=="none"){return "";}else{return self.settings.SCATTERMouseDownCircleStrokeWidth;} });
               })
            }
            if(self.settings.SCATTERMouseEnterDisplay==true){
               SCATTERMouseOn.on("mouseenter",function(d){
                   d3.select(this)
                       .style("r", function(d,i) { if(self.settings.SCATTERMouseEnterCircleSize=="none"){return "";}else{return self.settings.SCATTERMouseEnterCircleSize;} })
                       .style("fill", function(d,i) { if(self.settings.SCATTERMouseEnterColor=="none"){return "";}else{return self.settings.SCATTERMouseEnterColor;} })
                       .style("opacity",  function(d,i) { if(self.settings.SCATTERMouseEnterCircleOpacity=="none"){return "";}else{return self.settings.SCATTERMouseEnterCircleOpacity;} })
                       .style("stroke", function(d,i) { if(self.settings.SCATTERMouseEnterCircleStrokeColor=="none"){return "";}else{return self.settings.SCATTERMouseEnterCircleStrokeColor;} })
                       .style("stroke-width", function(d,i) { if(self.settings.SCATTERMouseEnterCircleStrokeWidth=="none"){return "";}else{return self.settings.SCATTERMouseEnterCircleStrokeWidth;} });
               })
            }
            if(self.settings.SCATTERMouseLeaveDisplay==true){
               SCATTERMouseOn.on("mouseleave",function(d){
                   SCATTERMouseOnTooltip.style("opacity",0.0);
                   d3.select(this)
                       .style("r", function(d,i) { if(self.settings.SCATTERMouseLeaveCircleSize=="none"){return "";}else{return self.settings.SCATTERMouseLeaveCircleSize;} })
                       .style("fill", function(d,i) { if(self.settings.SCATTERMouseLeaveColor=="none"){return "";}else{return self.settings.SCATTERMouseLeaveColor;} })
                       .style("opacity",  function(d,i) { if(self.settings.SCATTERMouseLeaveCircleOpacity=="none"){return "";}else{return self.settings.SCATTERMouseLeaveCircleOpacity;} })
                       .style("stroke", function(d,i) { if(self.settings.SCATTERMouseLeaveCircleStrokeColor=="none"){return "";}else{return self.settings.SCATTERMouseLeaveCircleStrokeColor;} })
                       .style("stroke-width", function(d,i) { if(self.settings.SCATTERMouseLeaveCircleStrokeWidth=="none"){return "";}else{return self.settings.SCATTERMouseLeaveCircleStrokeWidth;} });
               })
            }
            if(self.settings.SCATTERMouseUpDisplay==true){
               SCATTERMouseOn.on("mouseup",function(d){
                   d3.select(this)
                       .style("r", function(d,i) { if(self.settings.SCATTERMouseUpCircleSize=="none"){return "";}else{return self.settings.SCATTERMouseUpCircleSize;} })
                       .style("fill", function(d,i) { if(self.settings.SCATTERMouseUpColor=="none"){return "";}else{return self.settings.SCATTERMouseUpColor;} })
                       .style("opacity",  function(d,i) { if(self.settings.SCATTERMouseUpCircleOpacity=="none"){return "";}else{return self.settings.SCATTERMouseUpCircleOpacity;} })
                       .style("stroke", function(d,i) { if(self.settings.SCATTERMouseUpCircleStrokeColor=="none"){return "";}else{return self.settings.SCATTERMouseUpCircleStrokeColor;} })
                       .style("stroke-width", function(d,i) { if(self.settings.SCATTERMouseUpCircleStrokeWidth=="none"){return "";}else{return self.settings.SCATTERMouseUpCircleStrokeWidth;} });
               })
            }
            if(self.settings.SCATTERMouseMoveDisplay==true){
               SCATTERMouseOn.on("mousemove",function(d){
                   d3.select(this)
                       .style("r", function(d,i) { if(self.settings.SCATTERMouseMoveCircleSize=="none"){return "";}else{return self.settings.SCATTERMouseMoveCircleSize;} })
                       .style("fill", function(d,i) { if(self.settings.SCATTERMouseMoveColor=="none"){return "";}else{return self.settings.SCATTERMouseMoveColor;} })
                       .style("opacity",  function(d,i) { if(self.settings.SCATTERMouseMoveCircleOpacity=="none"){return "";}else{return self.settings.SCATTERMouseMoveCircleOpacity;} })
                       .style("stroke", function(d,i) { if(self.settings.SCATTERMouseMoveCircleStrokeColor=="none"){return "";}else{return self.settings.SCATTERMouseMoveCircleStrokeColor;} })
                       .style("stroke-width", function(d,i) { if(self.settings.SCATTERMouseMoveCircleStrokeWidth=="none"){return "";}else{return self.settings.SCATTERMouseMoveCircleStrokeWidth;} });
                   SCATTERMouseOnTooltip.style("left", (d3.event.pageX) + "px")
                   .style("top", (d3.event.pageY + 20) + "px");
               })
            }
            if(self.settings.SCATTERMouseOutDisplay==true){
               SCATTERMouseOn.on("mouseout",function(d){
                   SCATTERMouseOnTooltip.style("opacity",0.0);
                   d3.select(this)
                       .transition()
                       .duration(self.settings.SCATTERMouseOutAnimationTime)
                       .style("r", function(d,i) { if(self.settings.SCATTERMouseOutCircleSize=="none"){return "";}else{return self.settings.SCATTERMouseOutCircleSize;} })
                       .style("fill", function(d,i) { if(self.settings.SCATTERMouseOutColor=="none"){return "";}else{return self.settings.SCATTERMouseOutColor;} })
                       .style("opacity",  function(d,i) { if(self.settings.SCATTERMouseOutCircleOpacity=="none"){return "";}else{return self.settings.SCATTERMouseOutCircleOpacity;} })
                       .style("stroke", function(d,i) { if(self.settings.SCATTERMouseOutCircleStrokeColor=="none"){return "";}else{return self.settings.SCATTERMouseOutCircleStrokeColor;} })
                       .style("stroke-width", function(d,i) { if(self.settings.SCATTERMouseOutCircleStrokeWidth=="none"){return "";}else{return self.settings.SCATTERMouseOutCircleStrokeWidth;} });
               });
            }
        }

    }


    if(self.SNP.length > 0){
            function BioCircosSNP(d) {
              return self.SNP[snpi].map(function(v, i) {
                var snp_k = (d[self.initGenome[v.chr]].endAngle - d[self.initGenome[v.chr]].startAngle) / d[self.initGenome[v.chr]].value;
                // Check if a range is given
                if(self.SNPsettings.range){
                  var snp_value_maxmin_instance = self.SNPsettings.range;
                }
                else{
                  var snp_value_maxmin_instance = self.snp_value_maxmin(self.SNP[snpi]);
                }
                return {
                  snp_angle: v.pos * snp_k + d[self.initGenome[v.chr]].startAngle,
                  snp_chr: v.chr,
                  snp_pos: v.pos,
                  snp_val: v.value,
                  snp_des: v.des,
                  snp_color: v.color,
                  snp_link: v.link,
                  snp_opacity: v.opacity,
                  snp_click_label: "snp"+snpi+"_"+i,
                  x: (0 + Math.sin(v.pos * snp_k + d[self.initGenome[v.chr]].startAngle) * (self.SNPsettings.minRadius + ( (v.value-snp_value_maxmin_instance[1])/(snp_value_maxmin_instance[0]-snp_value_maxmin_instance[1])*(self.SNPsettings.maxRadius-self.SNPsettings.minRadius) ))),  //snp_value_maxmin_instance[0] max
                  y: (0 - Math.cos(v.pos * snp_k + d[self.initGenome[v.chr]].startAngle) * (self.SNPsettings.minRadius + ( (v.value-snp_value_maxmin_instance[1])/(snp_value_maxmin_instance[0]-snp_value_maxmin_instance[1])*(self.SNPsettings.maxRadius-self.SNPsettings.minRadius) )))
                };
              });
            }
        for(var snpi=0; snpi<self.SNP.length; snpi++){
            self.update_SNPsettings(self.SNPConfig[snpi]);

            var snp_objects = BioCircosSNP(chord.groups())

            if(self.SNPsettings.PointType=="circle"){
              if(self.SNPsettings.SNPAnimationDisplay==false){
                svg.append("g")
                    .attr("class", "BioCircosSNP")
                  .selectAll("circle")
                    .data(snp_objects)
                    .enter()
                    .append("a")
                    .attr("xlink:href", function(d){return d.scatter_link})
                    .append("circle")
                    .attr("id", "BioCircosSNP")
                    .attr("fill", function(d,i) { if(d.snp_color!=undefined){return d.snp_color;}else{return self.SNPsettings.SNPFillColor;} })
                    .attr("r", self.SNPsettings.circleSize)
                    .attr("cx", function(d) { return d.x; })
                    .attr("cy", function(d) { return d.y; })
                    .attr("opacity", function(d,i) { return d.snp_opacity; });
               }
              if(self.SNPsettings.SNPAnimationDisplay==true){
                svg.append("g")
                    .attr("class", "BioCircosSNP")
                  .selectAll("circle")
                    .data(snp_objects)
                    .enter()
                    .append("a")
                    .attr("xlink:href", function(d){return d.scatter_link})
                    .append("circle")
                    .attr("id", "BioCircosSNP")
                    .attr("fill", function(d,i) { if(d.snp_color!=undefined){return d.snp_color;}else{return self.SNPsettings.SNPFillColor;} })
                    .attr("r", self.SNPsettings.circleSize)
                    .attr("opacity", function(d,i) { return d.snp_opacity; })
		    .attr("cx",function(d){
			    return 0;
		    })
		    .attr("cy",function(d){
			    return 0;
		    })
		    .transition()
		    .delay(function(d,i){
			    return i * self.SNPsettings.SNPAnimationDelay;
		    })
		    .duration(self.SNPsettings.SNPAnimationTime)
		    .ease(self.SNPsettings.SNPAnimationType)
                    .attr("cx", function(d) { return d.x; })
                    .attr("cy", function(d) { return d.y; });
               }

                if(self.settings.SNPMouseClickTextFromData=="first"){
                    svg.append("g")
                        .attr("class", "BioCircosSNPlabel")
                      .selectAll("text")
                        .data(snp_objects)
                        .enter().append("text")
                        .attr("class", "dragText")
                        .attr("id", function(d,i) { return "snp"+snpi+"_"+i; })
                        .text(function(d) { return d.snp_chr; })
                        .attr("x", -1000)
                        .attr("y", -1000)
                        .style("opacity", 0)
                        .style("font-size", 1)
                        .attr("fill", self.SNPsettings.SNPFillColor);
                }
                if(self.settings.SNPMouseClickTextFromData=="second"){
                    svg.append("g")
                        .attr("class", "BioCircosSNPlabel")
                      .selectAll("text")
                        .data(snp_objects)
                        .enter().append("text")
                        .attr("class", "dragText")
                        .attr("id", function(d,i) { return "snp"+snpi+"_"+i; })
                        .text(function(d) { return d.snp_pos; })
                        .attr("x", -1000)
                        .attr("y", -1000)
                        .style("opacity", 0)
                        .style("font-size", 1)
                        .attr("fill", self.SNPsettings.SNPFillColor);
                }
                if(self.settings.SNPMouseClickTextFromData=="third"){
                    svg.append("g")
                        .attr("class", "BioCircosSNPlabel")
                      .selectAll("text")
                        .data(snp_objects)
                        .enter().append("text")
                        .attr("class", "dragText")
                        .attr("id", function(d,i) { return "snp"+snpi+"_"+i; })
                        .text(function(d) { return d.snp_val; })
                        .attr("x", -1000)
                        .attr("y", -1000)
                        .style("opacity", 0)
                        .style("font-size", 1)
                        .attr("fill", self.SNPsettings.SNPFillColor);
                }
                if(self.settings.SNPMouseClickTextFromData=="fourth"){
                    svg.append("g")
                        .attr("class", "BioCircosSNPlabel")
                      .selectAll("text")
                        .data(snp_objects)
                        .enter().append("text")
                        .attr("class", "dragText")
                        .attr("id", function(d,i) { return "snp"+snpi+"_"+i; })
                        .text(function(d) { return d.snp_des; })
                        .attr("x", -1000)
                        .attr("y", -1000)
                        .style("opacity", 0)
                        .style("font-size", 1)
                        .attr("fill", self.SNPsettings.SNPFillColor);
                }
            }

            if(self.SNPsettings.PointType=="rect"){
                svg.append("g")
                    .attr("class", "BioCircosSNP")
                  .selectAll("rect")
                    .data(snp_objects)
                    .enter()
                    .append("a")
                    .attr("xlink:href", function(d){return d.scatter_link})
                    .append("rect")
                    .attr("id", "BioCircosSNP")
                    .attr("x", function(d) { return d.x; })
                    .attr("y", function(d) { return d.y; })
                    .attr("width", self.SNPsettings.rectWidth)
                    .attr("height", self.SNPsettings.rectHeight)
                    //.attr("fill", self.SNPsettings.SNPFillColor);
                    .attr("opacity", function(d,i) { return d.snp_opacity; })
                    .attr("fill", function(d,i) { if(d.snp_color!=undefined){return d.snp_color;}else{return self.SNPsettings.SNPFillColor;} });
            }

                 self.init_SNPsettings();

        }

        if(self.settings.SNPMouseEvent==true){
            var SNPMouseOnTooltip = d3.select("body")
                .append("div")
                .attr("class","BioCircosSNPTooltip")
                .attr("id","BioCircosSNPTooltip")
                .style("opacity",0);

            var SNPMouseOn = svg.selectAll("#BioCircosSNP");

            if(self.settings.SNPMouseOverDisplay==true){
                SNPMouseOn.on("mouseover",function(d){
                      SNPMouseOnTooltip.html(self.settings.SNPMouseOverTooltipsHtml01+d.snp_chr+self.settings.SNPMouseOverTooltipsHtml02+d.snp_pos+self.settings.SNPMouseOverTooltipsHtml03+d.snp_val+self.settings.SNPMouseOverTooltipsHtml04+d.snp_des+self.settings.SNPMouseOverTooltipsHtml05)
                       .style("left", (d3.event.pageX) + "px")
                       .style("top", (d3.event.pageY + 20) + "px")
                       .style("position", self.settings.SNPMouseOverTooltipsPosition)
                       .style("background-color", self.settings.SNPMouseOverTooltipsBackgroundColor)
                       .style("border-style", self.settings.SNPMouseOverTooltipsBorderStyle)
                       .style("border-width", self.settings.SNPMouseOverTooltipsBorderWidth)
                       .style("padding", self.settings.SNPMouseOverTooltipsPadding)
                       .style("border-radius", self.settings.SNPMouseOverTooltipsBorderRadius)
                       .style("opacity", self.settings.SNPMouseOverTooltipsOpacity)
                    d3.select(this)
                       .style("r",  function(d,i) { if(self.settings.SNPMouseOverCircleSize=="none"){return "";}else{return self.settings.SNPMouseOverCircleSize;} })
                       .style("fill",  function(d,i) { if(self.settings.SNPMouseOverColor=="none"){return "";}else{return self.settings.SNPMouseOverColor;} })
                       .style("opacity",  function(d,i) { if(self.settings.SNPMouseOverCircleOpacity=="none"){return "";}else{return self.settings.SNPMouseOverCircleOpacity;} })
                       .style("stroke", function(d,i) { if(self.settings.SNPMouseOverCircleStrokeColor=="none"){return "";}else{return self.settings.SNPMouseOverCircleStrokeColor;} })
                       .style("stroke-width", function(d,i) { if(self.settings.SNPMouseOverCircleStrokeWidth=="none"){return "";}else{return self.settings.SNPMouseOverCircleStrokeWidth;} });
                })
            }
            if(self.settings.SNPMouseClickDisplay==true){
                SNPMouseOn.on("click",function(d){
                    d3.select(this)
                       .style("r",  function(d,i) { if(self.settings.SNPMouseClickCircleSize=="none"){return "";}else{return self.settings.SNPMouseClickCircleSize;} })
                       .style("fill",  function(d,i) { if(self.settings.SNPMouseClickColor=="none"){return "";}else{return self.settings.SNPMouseClickColor;} })
                       .style("opacity",  function(d,i) { if(self.settings.SNPMouseClickCircleOpacity=="none"){return "";}else{return self.settings.SNPMouseClickCircleOpacity;} })
                       .style("stroke", function(d,i) { if(self.settings.SNPMouseClickCircleStrokeColor=="none"){return "";}else{return self.settings.SNPMouseClickCircleStrokeColor;} })
                       .style("stroke-width", function(d,i) { if(self.settings.SNPMouseClickCircleStrokeWidth=="none"){return "";}else{return self.settings.SNPMouseClickCircleStrokeWidth;} });
                    d3.select("#"+d.snp_click_label)
                        .style("opacity", self.settings.SNPMouseClickTextOpacity)
                        .style("fill", self.settings.SNPMouseClickTextColor)
                        .style("font-size", self.settings.SNPMouseClickTextSize)
                        .attr("x", d.x+self.settings.SNPMouseClickTextPostionX)
                        .attr("y", d.y+self.settings.SNPMouseClickTextPostionY);
                })
            }

            if(self.settings.SNPMouseClickTextDrag==true){
                svg.selectAll("text.dragText").call(drag);
            }

            if(self.settings.SNPMouseDownDisplay==true){
               SNPMouseOn.on("mousedown",function(d){
                   d3.select(this)
                       .style("r", function(d,i) { if(self.settings.SNPMouseDownCircleSize=="none"){return "";}else{return self.settings.SNPMouseDownCircleSize;} })
                       .style("fill", function(d,i) { if(self.settings.SNPMouseDownColor=="none"){return "";}else{return self.settings.SNPMouseDownColor;} })
                       .style("opacity", function(d,i) { if(self.settings.SNPMouseDownCircleOpacity=="none"){return "";}else{return self.settings.SNPMouseDownCircleOpacity;} })
                       .style("stroke", function(d,i) { if(self.settings.SNPMouseDownCircleStrokeColor=="none"){return "";}else{return self.settings.SNPMouseDownCircleStrokeColor;} })
                       .style("stroke-width", function(d,i) { if(self.settings.SNPMouseDownCircleStrokeWidth=="none"){return "";}else{return self.settings.SNPMouseDownCircleStrokeWidth;} });
               })
            }
            if(self.settings.SNPMouseEnterDisplay==true){
               SNPMouseOn.on("mouseenter",function(d){
                   d3.select(this)
                       .style("r", function(d,i) { if(self.settings.SNPMouseEnterCircleSize=="none"){return "";}else{return self.settings.SNPMouseEnterCircleSize;} })
                       .style("fill", function(d,i) { if(self.settings.SNPMouseEnterColor=="none"){return "";}else{return self.settings.SNPMouseEnterColor;} })
                       .style("opacity", function(d,i) { if(self.settings.SNPMouseEnterCircleOpacity=="none"){return "";}else{return self.settings.SNPMouseEnterCircleOpacity;} })
                       .style("stroke", function(d,i) { if(self.settings.SNPMouseEnterCircleStrokeColor=="none"){return "";}else{return self.settings.SNPMouseEnterCircleStrokeColor;} })
                       .style("stroke-width", function(d,i) { if(self.settings.SNPMouseEnterCircleStrokeWidth=="none"){return "";}else{return self.settings.SNPMouseEnterCircleStrokeWidth;} });
               })
            }
            if(self.settings.SNPMouseLeaveDisplay==true){
               SNPMouseOn.on("mouseleave",function(d){
                   SNPMouseOnTooltip.style("opacity",0.0);
                   d3.select(this)
                       .style("r", function(d,i) { if(self.settings.SNPMouseLeaveCircleSize=="none"){return "";}else{return self.settings.SNPMouseLeaveCircleSize;} })
                       .style("fill", function(d,i) { if(self.settings.SNPMouseLeaveColor=="none"){return "";}else{return self.settings.SNPMouseLeaveColor;} })
                       .style("opacity", function(d,i) { if(self.settings.SNPMouseLeaveCircleOpacity=="none"){return "";}else{return self.settings.SNPMouseLeaveCircleOpacity;} })
                       .style("stroke", function(d,i) { if(self.settings.SNPMouseLeaveCircleStrokeColor=="none"){return "";}else{return self.settings.SNPMouseLeaveCircleStrokeColor;} })
                       .style("stroke-width", function(d,i) { if(self.settings.SNPMouseLeaveCircleStrokeWidth=="none"){return "";}else{return self.settings.SNPMouseLeaveCircleStrokeWidth;} });
               })
            }
            if(self.settings.SNPMouseUpDisplay==true){
               SNPMouseOn.on("mouseup",function(d){
                   d3.select(this)
                       .style("r", function(d,i) { if(self.settings.SNPMouseUpCircleSize=="none"){return "";}else{return self.settings.SNPMouseUpCircleSize;} })
                       .style("fill", function(d,i) { if(self.settings.SNPMouseUpColor=="none"){return "";}else{return self.settings.SNPMouseUpColor;} })
                       .style("opacity", function(d,i) { if(self.settings.SNPMouseUpCircleOpacity=="none"){return "";}else{return self.settings.SNPMouseUpCircleOpacity;} })
                       .style("stroke", function(d,i) { if(self.settings.SNPMouseUpCircleStrokeColor=="none"){return "";}else{return self.settings.SNPMouseUpCircleStrokeColor;} })
                       .style("stroke-width", function(d,i) { if(self.settings.SNPMouseUpCircleStrokeWidth=="none"){return "";}else{return self.settings.SNPMouseUpCircleStrokeWidth;} });
               })
            }
            if(self.settings.SNPMouseMoveDisplay==true){
               SNPMouseOn.on("mousemove",function(d){
                   d3.select(this)
                       .style("r", function(d,i) { if(self.settings.SNPMouseMoveCircleSize=="none"){return "";}else{return self.settings.SNPMouseMoveCircleSize;} })
                       .style("fill", function(d,i) { if(self.settings.SNPMouseMoveColor=="none"){return "";}else{return self.settings.SNPMouseMoveColor;} })
                       .style("opacity", function(d,i) { if(self.settings.SNPMouseMoveCircleOpacity=="none"){return "";}else{return self.settings.SNPMouseMoveCircleOpacity;} })
                       .style("stroke", function(d,i) { if(self.settings.SNPMouseMoveCircleStrokeColor=="none"){return "";}else{return self.settings.SNPMouseMoveCircleStrokeColor;} })
                       .style("stroke-width", function(d,i) { if(self.settings.SNPMouseMoveCircleStrokeWidth=="none"){return "";}else{return self.settings.SNPMouseMoveCircleStrokeWidth;} });
                   SNPMouseOnTooltip.style("left", (d3.event.pageX) + "px")
                   .style("top", (d3.event.pageY + 20) + "px");
               })
            }
            if(self.settings.SNPMouseOutDisplay==true){
               SNPMouseOn.on("mouseout",function(d){
                   SNPMouseOnTooltip.style("opacity",0.0);
                   d3.select(this)
                       .transition()
                       .duration(self.settings.SNPMouseOutAnimationTime)
                       .style("r", function(d,i) { if(self.settings.SNPMouseOutCircleSize=="none"){return "";}else{return self.settings.SNPMouseOutCircleSize;} })
                       .style("fill", function(d,i) { if(self.settings.SNPMouseOutColor=="none"){return "";}else{return self.settings.SNPMouseOutColor;} })
                       .style("opacity", function(d,i) { if(self.settings.SNPMouseOutCircleOpacity=="none"){return "";}else{return self.settings.SNPMouseOutCircleOpacity;} })
                       .style("stroke", function(d,i) { if(self.settings.SNPMouseOutCircleStrokeColor=="none"){return "";}else{return self.settings.SNPMouseOutCircleStrokeColor;} })
                       .style("stroke-width", function(d,i) { if(self.settings.SNPMouseOutCircleStrokeWidth=="none"){return "";}else{return self.settings.SNPMouseOutCircleStrokeWidth;} });
               });
            }
        }
    }

    if(self.LINK.length > 0){
            function BioCircosLINK(d) {
              return self.LINK[linki].map(function(v, i) {
                var start_k = (d[self.initGenome[v.g1chr]].endAngle - d[self.initGenome[v.g1chr]].startAngle) / d[self.initGenome[v.g1chr]].value;
                var end_k = (d[self.initGenome[v.g2chr]].endAngle - d[self.initGenome[v.g2chr]].startAngle) / d[self.initGenome[v.g2chr]].value;
                return {
                  link_angle1: (v.g1start/2+v.g1end/2) * start_k + d[self.initGenome[v.g1chr]].startAngle,
                  link_angle2: (v.g2start/2+v.g2end/2) * end_k + d[self.initGenome[v.g2chr]].startAngle,
                  link_label1: v.g1name,
                  link_label2: v.g2name,
                  link_pair: v.fusion,
                  link_width: self.LINKsettings.LinkWidth,
                  link_X1: (0 + Math.sin((v.g1start/2+v.g1end/2) * start_k + d[self.initGenome[v.g1chr]].startAngle) * (self.LINKsettings.LinkRadius)),
                  link_Y1: (0 - Math.cos((v.g1start/2+v.g1end/2) * start_k + d[self.initGenome[v.g1chr]].startAngle) * (self.LINKsettings.LinkRadius)),
                  link_X2: (0 + Math.sin((v.g2start/2+v.g2end/2) * end_k + d[self.initGenome[v.g2chr]].startAngle) * (self.LINKsettings.LinkRadius)),
                  link_Y2: (0 - Math.cos((v.g2start/2+v.g2end/2) * end_k + d[self.initGenome[v.g2chr]].startAngle) * (self.LINKsettings.LinkRadius))
                };
              });
            }
        for(var linki=0; linki<self.LINK.length; linki++){
            self.update_LINKsettings(self.LINKConfig[linki]);

            var link_objects = BioCircosLINK(chord.groups())

            if(self.LINKsettings.displayLinkAxis==true){
                svg.append("g")
                    .attr("class", "LINKAxis")
                    .selectAll("circle")
                    .data(["0"])
                    .enter().append("circle")
                    .attr("id", "LINKAxis")
                    .attr("cx", 0)
                    .attr("cy", 0)
                    .attr("fill", "none")
                    .attr("stroke",self.LINKsettings.LinkAxisColor)
                    .attr("stroke-width",self.LINKsettings.LinkAxisWidth)
                    .attr("r", function(d) { return self.LINKsettings.LinkRadius+self.LINKsettings.LinkAxisPad; });
            }

            var Link_svg = svg.append("g")
                .attr("class", "BioCircosLINK")
              .selectAll("path")
                .data(link_objects)
                .enter().append("path")
                .attr("d", function(d) { return "M"+d.link_X1+","+d.link_Y1+" Q0,0 "+d.link_X2+","+d.link_Y2+""; })
                .attr("class", "BioCircosLINK")
                .attr("fill","none")
                .attr("stroke",self.LINKsettings.LinkFillColor)
                .attr("stroke-width",self.LINKsettings.LinkWidth);

            if(self.LINKsettings.displayLinkLabel==true){
            svg.append("g")
                  .attr("class", "BioCircosLINKLabel")
              .selectAll("text")
                .data(link_objects)
                .enter().append("text")
                .attr("transform", function(d) {
                  return "rotate(" + (d.link_angle2 * 180 / Math.PI - 90) + ")"
                      + "translate(" + (self.LINKsettings.LinkRadius+self.LINKsettings.LinkAxisPad) + ",0)";
                })
                .attr("class", "BioCircosLINKLabel")
                .attr("id", function(d) { return d.link_pair; })
                .attr("x", self.LINKsettings.LinkLabelPad)
                .attr("dy", ".35em")
                .attr("fill", self.LINKsettings.LinkLabelColor)
                .style("font-size",self.LINKsettings.LinkLabelSize)
                .text(function(d) { return d.link_pair; });
            }
            self.init_LINKsettings();

            function draglinkmove(d) {
                d3.select(this)
                  .attr("x", d3.event.x )
                  .attr("y", d3.event.y );
            }
			
            var draglinklabel = d3.behavior.drag()
                      .on("drag", draglinkmove);

            if(self.settings.LINKLabelDragEvent==true){
                svg.selectAll("text.BioCircosLINKLabel").call(draglinklabel);
            }

        }

        if(self.settings.LINKMouseEvent==true){
            var LINKMouseOnTooltip = d3.select("body")
                .append("div")
                .attr("class","BioCircosLINKTooltip")
                .attr("id","BioCircosLINKTooltip")
                .style("opacity",0);

            var LINKMouseOn = svg.selectAll("path.BioCircosLINK");

            if(self.settings.LINKMouseOverDisplay==true){
                LINKMouseOn.on("mouseover",function(d){
                      LINKMouseOnTooltip.html(self.settings.LINKMouseOverTooltipsHtml01+d.link_pair+self.settings.LINKMouseOverTooltipsHtml02)
                       .style("left", (d3.event.pageX) + "px")
                       .style("top", (d3.event.pageY + 20) + "px")
                       .style("position", self.settings.LINKMouseOverTooltipsPosition)
                       .style("background-color", self.settings.LINKMouseOverTooltipsBackgroundColor)
                       .style("border-style", self.settings.LINKMouseOverTooltipsBorderStyle)
                       .style("border-width", self.settings.LINKMouseOverTooltipsBorderWidth)
                       .style("padding", self.settings.LINKMouseOverTooltipsPadding)
                       .style("border-radius", self.settings.LINKMouseOverTooltipsBorderRadius)
                       .style("opacity", self.settings.LINKMouseOverTooltipsOpacity)
                    d3.select(this)
                       .style("opacity", function(d,i) { if(self.settings.LINKMouseOverOpacity=="none"){return "";}else{return self.settings.LINKMouseOverOpacity;} })
                       .style("stroke", function(d,i) { if(self.settings.LINKMouseOverStrokeColor=="none"){return "";}else{return self.settings.LINKMouseOverStrokeColor;} })
                       .style("stroke-width", function(d,i) { if(self.settings.LINKMouseOverStrokeWidth=="none"){return "";}else{return self.settings.LINKMouseOverStrokeWidth;} });
                })
            }
            if(self.settings.LINKMouseClickDisplay==true){
                LINKMouseOn.on("click",function(d){
                    d3.select(this)
                       .style("opacity", function(d,i) { if(self.settings.LINKMouseClickOpacity=="none"){return "";}else{return self.settings.LINKMouseClickOpacity;} })
                       .style("stroke", function(d,i) { if(self.settings.LINKMouseClickStrokeColor=="none"){return "";}else{return self.settings.LINKMouseClickStrokeColor;} })
                       .style("stroke-width", function(d,i) { if(self.settings.LINKMouseClickStrokeWidth=="none"){return "";}else{return self.settings.LINKMouseClickStrokeWidth;} });
                })
            }
            if(self.settings.LINKMouseDownDisplay==true){
               LINKMouseOn.on("mousedown",function(d){
                   d3.select(this)
                       .style("opacity", function(d,i) { if(self.settings.LINKMouseDownOpacity=="none"){return "";}else{return self.settings.LINKMouseDownOpacity;} })
                       .style("stroke", function(d,i) { if(self.settings.LINKMouseDownStrokeColor=="none"){return "";}else{return self.settings.LINKMouseDownStrokeColor;} })
                       .style("stroke-width", function(d,i) { if(self.settings.LINKMouseDownStrokeWidth=="none"){return "";}else{return self.settings.LINKMouseDownStrokeWidth;} });
               })
            }
            if(self.settings.LINKMouseEnterDisplay==true){
               LINKMouseOn.on("mouseenter",function(d){
                   d3.select(this)
                       .style("opacity", function(d,i) { if(self.settings.LINKMouseEnterOpacity=="none"){return "";}else{return self.settings.LINKMouseEnterOpacity;} })
                       .style("stroke", function(d,i) { if(self.settings.LINKMouseEnterStrokeColor=="none"){return "";}else{return self.settings.LINKMouseEnterStrokeColor;} })
                       .style("stroke-width", function(d,i) { if(self.settings.LINKMouseEnterStrokeWidth=="none"){return "";}else{return self.settings.LINKMouseEnterStrokeWidth;} });
               })
            }
            if(self.settings.LINKMouseLeaveDisplay==true){
               LINKMouseOn.on("mouseleave",function(d){
                   LINKMouseOnTooltip.style("opacity",0.0);
                   d3.select(this)
                       .style("opacity", function(d,i) { if(self.settings.LINKMouseLeaveOpacity=="none"){return "";}else{return self.settings.LINKMouseLeaveOpacity;} })
                       .style("stroke", function(d,i) { if(self.settings.LINKMouseLeaveStrokeColor=="none"){return "";}else{return self.settings.LINKMouseLeaveStrokeColor;} })
                       .style("stroke-width", function(d,i) { if(self.settings.LINKMouseLeaveStrokeWidth=="none"){return "";}else{return self.settings.LINKMouseLeaveStrokeWidth;} });
               })
            }
            if(self.settings.LINKMouseUpDisplay==true){
               LINKMouseOn.on("mouseup",function(d){
                   d3.select(this)
                       .style("opacity", function(d,i) { if(self.settings.LINKMouseUpOpacity=="none"){return "";}else{return self.settings.LINKMouseUpOpacity;} })
                       .style("stroke", function(d,i) { if(self.settings.LINKMouseUpStrokeColor=="none"){return "";}else{return self.settings.LINKMouseUpStrokeColor;} })
                       .style("stroke-width", function(d,i) { if(self.settings.LINKMouseUpStrokeWidth=="none"){return "";}else{return self.settings.LINKMouseUpStrokeWidth;} });
               })
            }
            if(self.settings.LINKMouseMoveDisplay==true){
               LINKMouseOn.on("mousemove",function(d){
                   d3.select(this)
                       .style("opacity", function(d,i) { if(self.settings.LINKMouseMoveOpacity=="none"){return "";}else{return self.settings.LINKMouseMoveOpacity;} })
                       .style("stroke", function(d,i) { if(self.settings.LINKMouseMoveStrokeColor=="none"){return "";}else{return self.settings.LINKMouseMoveStrokeColor;} })
                       .style("stroke-width", function(d,i) { if(self.settings.LINKMouseMoveStrokeWidth=="none"){return "";}else{return self.settings.LINKMouseMoveStrokeWidth;} });
                   LINKMouseOnTooltip.style("left", (d3.event.pageX) + "px")
                   .style("top", (d3.event.pageY + 20) + "px");
               })
            }
            if(self.settings.LINKMouseOutDisplay==true){
               LINKMouseOn.on("mouseout",function(d){
                   LINKMouseOnTooltip.style("opacity",0.0);
                   d3.select(this)
                       .transition()
                       .duration(self.settings.LINKMouseOutAnimationTime)
                       .style("opacity", function(d,i) { if(self.settings.LINKMouseOutOpacity=="none"){return "";}else{return self.settings.LINKMouseOutOpacity;} })
                       .style("stroke", function(d,i) { if(self.settings.LINKMouseOutStrokeColor=="none"){return "";}else{return self.settings.LINKMouseOutStrokeColor;} })
                       .style("stroke-width", function(d,i) { if(self.settings.LINKMouseOutStrokeWidth=="none"){return "";}else{return self.settings.LINKMouseOutStrokeWidth;} });
               });
            }

        }
    }

    }
}(jQuery));
```
