## Supplementary Material to Figure 4 and Figure 6 for "BTRR complex deficiency is a driver for genomic instability in Bloom syndrome": d3.js

```
d3 = function() {
  var d3 = {
    version: "3.3.10"
  };
  if (!Date.now) Date.now = function() {
    return +new Date();
  };
  var d3_arraySlice = [].slice, d3_array = function(list) {
    return d3_arraySlice.call(list);
  };
  var d3_document = document, d3_documentElement = d3_document.documentElement, d3_window = window;
  try {
    d3_array(d3_documentElement.childNodes)[0].nodeType;
  } catch (e) {
    d3_array = function(list) {
      var i = list.length, array = new Array(i);
      while (i--) array[i] = list[i];
      return array;
    };
  }
  try {
    d3_document.createElement("div").style.setProperty("opacity", 0, "");
  } catch (error) {
    var d3_element_prototype = d3_window.Element.prototype, d3_element_setAttribute = d3_element_prototype.setAttribute, d3_element_setAttributeNS = d3_element_prototype.setAttributeNS, d3_style_prototype = d3_window.CSSStyleDeclaration.prototype, d3_style_setProperty = d3_style_prototype.setProperty;
    d3_element_prototype.setAttribute = function(name, value) {
      d3_element_setAttribute.call(this, name, value + "");
    };
    d3_element_prototype.setAttributeNS = function(space, local, value) {
      d3_element_setAttributeNS.call(this, space, local, value + "");
    };
    d3_style_prototype.setProperty = function(name, value, priority) {
      d3_style_setProperty.call(this, name, value + "", priority);
    };
  }
  d3.ascending = function(a, b) {
    return a < b ? -1 : a > b ? 1 : a >= b ? 0 : NaN;
  };
  d3.descending = function(a, b) {
    return b < a ? -1 : b > a ? 1 : b >= a ? 0 : NaN;
  };
  d3.min = function(array, f) {
    var i = -1, n = array.length, a, b;
    if (arguments.length === 1) {
      while (++i < n && !((a = array[i]) != null && a <= a)) a = undefined;
      while (++i < n) if ((b = array[i]) != null && a > b) a = b;
    } else {
      while (++i < n && !((a = f.call(array, array[i], i)) != null && a <= a)) a = undefined;
      while (++i < n) if ((b = f.call(array, array[i], i)) != null && a > b) a = b;
    }
    return a;
  };
  d3.max = function(array, f) {
    var i = -1, n = array.length, a, b;
    if (arguments.length === 1) {
      while (++i < n && !((a = array[i]) != null && a <= a)) a = undefined;
      while (++i < n) if ((b = array[i]) != null && b > a) a = b;
    } else {
      while (++i < n && !((a = f.call(array, array[i], i)) != null && a <= a)) a = undefined;
      while (++i < n) if ((b = f.call(array, array[i], i)) != null && b > a) a = b;
    }
    return a;
  };
  d3.extent = function(array, f) {
    var i = -1, n = array.length, a, b, c;
    if (arguments.length === 1) {
      while (++i < n && !((a = c = array[i]) != null && a <= a)) a = c = undefined;
      while (++i < n) if ((b = array[i]) != null) {
        if (a > b) a = b;
        if (c < b) c = b;
      }
    } else {
      while (++i < n && !((a = c = f.call(array, array[i], i)) != null && a <= a)) a = undefined;
      while (++i < n) if ((b = f.call(array, array[i], i)) != null) {
        if (a > b) a = b;
        if (c < b) c = b;
      }
    }
    return [ a, c ];
  };
  d3.sum = function(array, f) {
    var s = 0, n = array.length, a, i = -1;
    if (arguments.length === 1) {
      while (++i < n) if (!isNaN(a = +array[i])) s += a;
    } else {
      while (++i < n) if (!isNaN(a = +f.call(array, array[i], i))) s += a;
    }
    return s;
  };
  function d3_number(x) {
    return x != null && !isNaN(x);
  }
  d3.mean = function(array, f) {
    var n = array.length, a, m = 0, i = -1, j = 0;
    if (arguments.length === 1) {
      while (++i < n) if (d3_number(a = array[i])) m += (a - m) / ++j;
    } else {
      while (++i < n) if (d3_number(a = f.call(array, array[i], i))) m += (a - m) / ++j;
    }
    return j ? m : undefined;
  };
  d3.quantile = function(values, p) {
    var H = (values.length - 1) * p + 1, h = Math.floor(H), v = +values[h - 1], e = H - h;
    return e ? v + e * (values[h] - v) : v;
  };
  d3.median = function(array, f) {
    if (arguments.length > 1) array = array.map(f);
    array = array.filter(d3_number);
    return array.length ? d3.quantile(array.sort(d3.ascending), .5) : undefined;
  };
  d3.bisector = function(f) {
    return {
      left: function(a, x, lo, hi) {
        if (arguments.length < 3) lo = 0;
        if (arguments.length < 4) hi = a.length;
        while (lo < hi) {
          var mid = lo + hi >>> 1;
          if (f.call(a, a[mid], mid) < x) lo = mid + 1; else hi = mid;
        }
        return lo;
      },
      right: function(a, x, lo, hi) {
        if (arguments.length < 3) lo = 0;
        if (arguments.length < 4) hi = a.length;
        while (lo < hi) {
          var mid = lo + hi >>> 1;
          if (x < f.call(a, a[mid], mid)) hi = mid; else lo = mid + 1;
        }
        return lo;
      }
    };
  };
  var d3_bisector = d3.bisector(function(d) {
    return d;
  });
  d3.bisectLeft = d3_bisector.left;
  d3.bisect = d3.bisectRight = d3_bisector.right;
  d3.shuffle = function(array) {
    var m = array.length, t, i;
    while (m) {
      i = Math.random() * m-- | 0;
      t = array[m], array[m] = array[i], array[i] = t;
    }
    return array;
  };
  d3.permute = function(array, indexes) {
    var i = indexes.length, permutes = new Array(i);
    while (i--) permutes[i] = array[indexes[i]];
    return permutes;
  };
  d3.pairs = function(array) {
    var i = 0, n = array.length - 1, p0, p1 = array[0], pairs = new Array(n < 0 ? 0 : n);
    while (i < n) pairs[i] = [ p0 = p1, p1 = array[++i] ];
    return pairs;
  };
  d3.zip = function() {
    if (!(n = arguments.length)) return [];
    for (var i = -1, m = d3.min(arguments, d3_zipLength), zips = new Array(m); ++i < m; ) {
      for (var j = -1, n, zip = zips[i] = new Array(n); ++j < n; ) {
        zip[j] = arguments[j][i];
      }
    }
    return zips;
  };
  function d3_zipLength(d) {
    return d.length;
  }
  d3.transpose = function(matrix) {
    return d3.zip.apply(d3, matrix);
  };
  d3.keys = function(map) {
    var keys = [];
    for (var key in map) keys.push(key);
    return keys;
  };
  d3.values = function(map) {
    var values = [];
    for (var key in map) values.push(map[key]);
    return values;
  };
  d3.entries = function(map) {
    var entries = [];
    for (var key in map) entries.push({
      key: key,
      value: map[key]
    });
    return entries;
  };
  d3.merge = function(arrays) {
    var n = arrays.length, m, i = -1, j = 0, merged, array;
    while (++i < n) j += arrays[i].length;
    merged = new Array(j);
    while (--n >= 0) {
      array = arrays[n];
      m = array.length;
      while (--m >= 0) {
        merged[--j] = array[m];
      }
    }
    return merged;
  };
  var abs = Math.abs;
  d3.range = function(start, stop, step) {
    if (arguments.length < 3) {
      step = 1;
      if (arguments.length < 2) {
        stop = start;
        start = 0;
      }
    }
    if ((stop - start) / step === Infinity) throw new Error("infinite range");
    var range = [], k = d3_range_integerScale(abs(step)), i = -1, j;
    start *= k, stop *= k, step *= k;
    if (step < 0) while ((j = start + step * ++i) > stop) range.push(j / k); else while ((j = start + step * ++i) < stop) range.push(j / k);
    return range;
  };
  function d3_range_integerScale(x) {
    var k = 1;
    while (x * k % 1) k *= 10;
    return k;
  }
  function d3_class(ctor, properties) {
    try {
      for (var key in properties) {
        Object.defineProperty(ctor.prototype, key, {
          value: properties[key],
          enumerable: false
        });
      }
    } catch (e) {
      ctor.prototype = properties;
    }
  }
  d3.map = function(object) {
    var map = new d3_Map();
    if (object instanceof d3_Map) object.forEach(function(key, value) {
      map.set(key, value);
    }); else for (var key in object) map.set(key, object[key]);
    return map;
  };
  function d3_Map() {}
  d3_class(d3_Map, {
    has: function(key) {
      return d3_map_prefix + key in this;
    },
    get: function(key) {
      return this[d3_map_prefix + key];
    },
    set: function(key, value) {
      return this[d3_map_prefix + key] = value;
    },
    remove: function(key) {
      key = d3_map_prefix + key;
      return key in this && delete this[key];
    },
    keys: function() {
      var keys = [];
      this.forEach(function(key) {
        keys.push(key);
      });
      return keys;
    },
    values: function() {
      var values = [];
      this.forEach(function(key, value) {
        values.push(value);
      });
      return values;
    },
    entries: function() {
      var entries = [];
      this.forEach(function(key, value) {
        entries.push({
          key: key,
          value: value
        });
      });
      return entries;
    },
    forEach: function(f) {
      for (var key in this) {
        if (key.charCodeAt(0) === d3_map_prefixCode) {
          f.call(this, key.substring(1), this[key]);
        }
      }
    }
  });
  var d3_map_prefix = "\x00", d3_map_prefixCode = d3_map_prefix.charCodeAt(0);
  d3.nest = function() {
    var nest = {}, keys = [], sortKeys = [], sortValues, rollup;
    function map(mapType, array, depth) {
      if (depth >= keys.length) return rollup ? rollup.call(nest, array) : sortValues ? array.sort(sortValues) : array;
      var i = -1, n = array.length, key = keys[depth++], keyValue, object, setter, valuesByKey = new d3_Map(), values;
      while (++i < n) {
        if (values = valuesByKey.get(keyValue = key(object = array[i]))) {
          values.push(object);
        } else {
          valuesByKey.set(keyValue, [ object ]);
        }
      }
      if (mapType) {
        object = mapType();
        setter = function(keyValue, values) {
          object.set(keyValue, map(mapType, values, depth));
        };
      } else {
        object = {};
        setter = function(keyValue, values) {
          object[keyValue] = map(mapType, values, depth);
        };
      }
      valuesByKey.forEach(setter);
      return object;
    }
    function entries(map, depth) {
      if (depth >= keys.length) return map;
      var array = [], sortKey = sortKeys[depth++];
      map.forEach(function(key, keyMap) {
        array.push({
          key: key,
          values: entries(keyMap, depth)
        });
      });
      return sortKey ? array.sort(function(a, b) {
        return sortKey(a.key, b.key);
      }) : array;
    }
    nest.map = function(array, mapType) {
      return map(mapType, array, 0);
    };
    nest.entries = function(array) {
      return entries(map(d3.map, array, 0), 0);
    };
    nest.key = function(d) {
      keys.push(d);
      return nest;
    };
    nest.sortKeys = function(order) {
      sortKeys[keys.length - 1] = order;
      return nest;
    };
    nest.sortValues = function(order) {
      sortValues = order;
      return nest;
    };
    nest.rollup = function(f) {
      rollup = f;
      return nest;
    };
    return nest;
  };
  d3.set = function(array) {
    var set = new d3_Set();
    if (array) for (var i = 0, n = array.length; i < n; ++i) set.add(array[i]);
    return set;
  };
  function d3_Set() {}
  d3_class(d3_Set, {
    has: function(value) {
      return d3_map_prefix + value in this;
    },
    add: function(value) {
      this[d3_map_prefix + value] = true;
      return value;
    },
    remove: function(value) {
      value = d3_map_prefix + value;
      return value in this && delete this[value];
    },
    values: function() {
      var values = [];
      this.forEach(function(value) {
        values.push(value);
      });
      return values;
    },
    forEach: function(f) {
      for (var value in this) {
        if (value.charCodeAt(0) === d3_map_prefixCode) {
          f.call(this, value.substring(1));
        }
      }
    }
  });
  d3.behavior = {};
  d3.rebind = function(target, source) {
    var i = 1, n = arguments.length, method;
    while (++i < n) target[method = arguments[i]] = d3_rebind(target, source, source[method]);
    return target;
  };
  function d3_rebind(target, source, method) {
    return function() {
      var value = method.apply(source, arguments);
      return value === source ? target : value;
    };
  }
  function d3_vendorSymbol(object, name) {
    if (name in object) return name;
    name = name.charAt(0).toUpperCase() + name.substring(1);
    for (var i = 0, n = d3_vendorPrefixes.length; i < n; ++i) {
      var prefixName = d3_vendorPrefixes[i] + name;
      if (prefixName in object) return prefixName;
    }
  }
  var d3_vendorPrefixes = [ "webkit", "ms", "moz", "Moz", "o", "O" ];
  function d3_noop() {}
  d3.dispatch = function() {
    var dispatch = new d3_dispatch(), i = -1, n = arguments.length;
    while (++i < n) dispatch[arguments[i]] = d3_dispatch_event(dispatch);
    return dispatch;
  };
  function d3_dispatch() {}
  d3_dispatch.prototype.on = function(type, listener) {
    var i = type.indexOf("."), name = "";
    if (i >= 0) {
      name = type.substring(i + 1);
      type = type.substring(0, i);
    }
    if (type) return arguments.length < 2 ? this[type].on(name) : this[type].on(name, listener);
    if (arguments.length === 2) {
      if (listener == null) for (type in this) {
        if (this.hasOwnProperty(type)) this[type].on(name, null);
      }
      return this;
    }
  };
  function d3_dispatch_event(dispatch) {
    var listeners = [], listenerByName = new d3_Map();
    function event() {
      var z = listeners, i = -1, n = z.length, l;
      while (++i < n) if (l = z[i].on) l.apply(this, arguments);
      return dispatch;
    }
    event.on = function(name, listener) {
      var l = listenerByName.get(name), i;
      if (arguments.length < 2) return l && l.on;
      if (l) {
        l.on = null;
        listeners = listeners.slice(0, i = listeners.indexOf(l)).concat(listeners.slice(i + 1));
        listenerByName.remove(name);
      }
      if (listener) listeners.push(listenerByName.set(name, {
        on: listener
      }));
      return dispatch;
    };
    return event;
  }
  d3.event = null;
  function d3_eventPreventDefault() {
    d3.event.preventDefault();
  }
  function d3_eventSource() {
    var e = d3.event, s;
    while (s = e.sourceEvent) e = s;
    return e;
  }
  function d3_eventDispatch(target) {
    var dispatch = new d3_dispatch(), i = 0, n = arguments.length;
    while (++i < n) dispatch[arguments[i]] = d3_dispatch_event(dispatch);
    dispatch.of = function(thiz, argumentz) {
      return function(e1) {
        try {
          var e0 = e1.sourceEvent = d3.event;
          e1.target = target;
          d3.event = e1;
          dispatch[e1.type].apply(thiz, argumentz);
        } finally {
          d3.event = e0;
        }
      };
    };
    return dispatch;
  }
  d3.requote = function(s) {
    return s.replace(d3_requote_re, "\\$&");
  };
  var d3_requote_re = /[\\\^\$\*\+\?\|\[\]\(\)\.\{\}]/g;
  var d3_subclass = {}.__proto__ ? function(object, prototype) {
    object.__proto__ = prototype;
  } : function(object, prototype) {
    for (var property in prototype) object[property] = prototype[property];
  };
  function d3_selection(groups) {
    d3_subclass(groups, d3_selectionPrototype);
    return groups;
  }
  var d3_select = function(s, n) {
    return n.querySelector(s);
  }, d3_selectAll = function(s, n) {
    return n.querySelectorAll(s);
  }, d3_selectMatcher = d3_documentElement[d3_vendorSymbol(d3_documentElement, "matchesSelector")], d3_selectMatches = function(n, s) {
    return d3_selectMatcher.call(n, s);
  };
  if (typeof Sizzle === "function") {
    d3_select = function(s, n) {
      return Sizzle(s, n)[0] || null;
    };
    d3_selectAll = function(s, n) {
      return Sizzle.uniqueSort(Sizzle(s, n));
    };
    d3_selectMatches = Sizzle.matchesSelector;
  }
  d3.selection = function() {
    return d3_selectionRoot;
  };
  var d3_selectionPrototype = d3.selection.prototype = [];
  d3_selectionPrototype.select = function(selector) {
    var subgroups = [], subgroup, subnode, group, node;
    selector = d3_selection_selector(selector);
    for (var j = -1, m = this.length; ++j < m; ) {
      subgroups.push(subgroup = []);
      subgroup.parentNode = (group = this[j]).parentNode;
      for (var i = -1, n = group.length; ++i < n; ) {
        if (node = group[i]) {
          subgroup.push(subnode = selector.call(node, node.__data__, i, j));
          if (subnode && "__data__" in node) subnode.__data__ = node.__data__;
        } else {
          subgroup.push(null);
        }
      }
    }
    return d3_selection(subgroups);
  };
  function d3_selection_selector(selector) {
    return typeof selector === "function" ? selector : function() {
      return d3_select(selector, this);
    };
  }
  d3_selectionPrototype.selectAll = function(selector) {
    var subgroups = [], subgroup, node;
    selector = d3_selection_selectorAll(selector);
    for (var j = -1, m = this.length; ++j < m; ) {
      for (var group = this[j], i = -1, n = group.length; ++i < n; ) {
        if (node = group[i]) {
          subgroups.push(subgroup = d3_array(selector.call(node, node.__data__, i, j)));
          subgroup.parentNode = node;
        }
      }
    }
    return d3_selection(subgroups);
  };
  function d3_selection_selectorAll(selector) {
    return typeof selector === "function" ? selector : function() {
      return d3_selectAll(selector, this);
    };
  }
  var d3_nsPrefix = {
    svg: "http://www.w3.org/2000/svg",
    xhtml: "http://www.w3.org/1999/xhtml",
    xlink: "http://www.w3.org/1999/xlink",
    xml: "http://www.w3.org/XML/1998/namespace",
    xmlns: "http://www.w3.org/2000/xmlns/"
  };
  d3.ns = {
    prefix: d3_nsPrefix,
    qualify: function(name) {
      var i = name.indexOf(":"), prefix = name;
      if (i >= 0) {
        prefix = name.substring(0, i);
        name = name.substring(i + 1);
      }
      return d3_nsPrefix.hasOwnProperty(prefix) ? {
        space: d3_nsPrefix[prefix],
        local: name
      } : name;
    }
  };
  d3_selectionPrototype.attr = function(name, value) {
    if (arguments.length < 2) {
      if (typeof name === "string") {
        var node = this.node();
        name = d3.ns.qualify(name);
        return name.local ? node.getAttributeNS(name.space, name.local) : node.getAttribute(name);
      }
      for (value in name) this.each(d3_selection_attr(value, name[value]));
      return this;
    }
    return this.each(d3_selection_attr(name, value));
  };
  function d3_selection_attr(name, value) {
    name = d3.ns.qualify(name);
    function attrNull() {
      this.removeAttribute(name);
    }
    function attrNullNS() {
      this.removeAttributeNS(name.space, name.local);
    }
    function attrConstant() {
      this.setAttribute(name, value);
    }
    function attrConstantNS() {
      this.setAttributeNS(name.space, name.local, value);
    }
    function attrFunction() {
      var x = value.apply(this, arguments);
      if (x == null) this.removeAttribute(name); else this.setAttribute(name, x);
    }
    function attrFunctionNS() {
      var x = value.apply(this, arguments);
      if (x == null) this.removeAttributeNS(name.space, name.local); else this.setAttributeNS(name.space, name.local, x);
    }
    return value == null ? name.local ? attrNullNS : attrNull : typeof value === "function" ? name.local ? attrFunctionNS : attrFunction : name.local ? attrConstantNS : attrConstant;
  }
  function d3_collapse(s) {
    return s.trim().replace(/\s+/g, " ");
  }
  d3_selectionPrototype.classed = function(name, value) {
    if (arguments.length < 2) {
      if (typeof name === "string") {
        var node = this.node(), n = (name = name.trim().split(/^|\s+/g)).length, i = -1;
        if (value = node.classList) {
          while (++i < n) if (!value.contains(name[i])) return false;
        } else {
          value = node.getAttribute("class");
          while (++i < n) if (!d3_selection_classedRe(name[i]).test(value)) return false;
        }
        return true;
      }
      for (value in name) this.each(d3_selection_classed(value, name[value]));
      return this;
    }
    return this.each(d3_selection_classed(name, value));
  };
  function d3_selection_classedRe(name) {
    return new RegExp("(?:^|\\s+)" + d3.requote(name) + "(?:\\s+|$)", "g");
  }
  function d3_selection_classed(name, value) {
    name = name.trim().split(/\s+/).map(d3_selection_classedName);
    var n = name.length;
    function classedConstant() {
      var i = -1;
      while (++i < n) name[i](this, value);
    }
    function classedFunction() {
      var i = -1, x = value.apply(this, arguments);
      while (++i < n) name[i](this, x);
    }
    return typeof value === "function" ? classedFunction : classedConstant;
  }
  function d3_selection_classedName(name) {
    var re = d3_selection_classedRe(name);
    return function(node, value) {
      if (c = node.classList) return value ? c.add(name) : c.remove(name);
      var c = node.getAttribute("class") || "";
      if (value) {
        re.lastIndex = 0;
        if (!re.test(c)) node.setAttribute("class", d3_collapse(c + " " + name));
      } else {
        node.setAttribute("class", d3_collapse(c.replace(re, " ")));
      }
    };
  }
  d3_selectionPrototype.style = function(name, value, priority) {
    var n = arguments.length;
    if (n < 3) {
      if (typeof name !== "string") {
        if (n < 2) value = "";
        for (priority in name) this.each(d3_selection_style(priority, name[priority], value));
        return this;
      }
      if (n < 2) return d3_window.getComputedStyle(this.node(), null).getPropertyValue(name);
      priority = "";
    }
    return this.each(d3_selection_style(name, value, priority));
  };
  function d3_selection_style(name, value, priority) {
    function styleNull() {
      this.style.removeProperty(name);
    }
    function styleConstant() {
      this.style.setProperty(name, value, priority);
    }
    function styleFunction() {
      var x = value.apply(this, arguments);
      if (x == null) this.style.removeProperty(name); else this.style.setProperty(name, x, priority);
    }
    return value == null ? styleNull : typeof value === "function" ? styleFunction : styleConstant;
  }
  d3_selectionPrototype.property = function(name, value) {
    if (arguments.length < 2) {
      if (typeof name === "string") return this.node()[name];
      for (value in name) this.each(d3_selection_property(value, name[value]));
      return this;
    }
    return this.each(d3_selection_property(name, value));
  };
  function d3_selection_property(name, value) {
    function propertyNull() {
      delete this[name];
    }
    function propertyConstant() {
      this[name] = value;
    }
    function propertyFunction() {
      var x = value.apply(this, arguments);
      if (x == null) delete this[name]; else this[name] = x;
    }
    return value == null ? propertyNull : typeof value === "function" ? propertyFunction : propertyConstant;
  }
  d3_selectionPrototype.text = function(value) {
    return arguments.length ? this.each(typeof value === "function" ? function() {
      var v = value.apply(this, arguments);
      this.textContent = v == null ? "" : v;
    } : value == null ? function() {
      this.textContent = "";
    } : function() {
      this.textContent = value;
    }) : this.node().textContent;
  };
  d3_selectionPrototype.html = function(value) {
    return arguments.length ? this.each(typeof value === "function" ? function() {
      var v = value.apply(this, arguments);
      this.innerHTML = v == null ? "" : v;
    } : value == null ? function() {
      this.innerHTML = "";
    } : function() {
      this.innerHTML = value;
    }) : this.node().innerHTML;
  };
  d3_selectionPrototype.append = function(name) {
    name = d3_selection_creator(name);
    return this.select(function() {
      return this.appendChild(name.apply(this, arguments));
    });
  };
  function d3_selection_creator(name) {
    return typeof name === "function" ? name : (name = d3.ns.qualify(name)).local ? function() {
      return this.ownerDocument.createElementNS(name.space, name.local);
    } : function() {
      return this.ownerDocument.createElementNS(this.namespaceURI, name);
    };
  }
  d3_selectionPrototype.insert = function(name, before) {
    name = d3_selection_creator(name);
    before = d3_selection_selector(before);
    return this.select(function() {
      return this.insertBefore(name.apply(this, arguments), before.apply(this, arguments) || null);
    });
  };
  d3_selectionPrototype.remove = function() {
    return this.each(function() {
      var parent = this.parentNode;
      if (parent) parent.removeChild(this);
    });
  };
  d3_selectionPrototype.data = function(value, key) {
    var i = -1, n = this.length, group, node;
    if (!arguments.length) {
      value = new Array(n = (group = this[0]).length);
      while (++i < n) {
        if (node = group[i]) {
          value[i] = node.__data__;
        }
      }
      return value;
    }
    function bind(group, groupData) {
      var i, n = group.length, m = groupData.length, n0 = Math.min(n, m), updateNodes = new Array(m), enterNodes = new Array(m), exitNodes = new Array(n), node, nodeData;
      if (key) {
        var nodeByKeyValue = new d3_Map(), dataByKeyValue = new d3_Map(), keyValues = [], keyValue;
        for (i = -1; ++i < n; ) {
          keyValue = key.call(node = group[i], node.__data__, i);
          if (nodeByKeyValue.has(keyValue)) {
            exitNodes[i] = node;
          } else {
            nodeByKeyValue.set(keyValue, node);
          }
          keyValues.push(keyValue);
        }
        for (i = -1; ++i < m; ) {
          keyValue = key.call(groupData, nodeData = groupData[i], i);
          if (node = nodeByKeyValue.get(keyValue)) {
            updateNodes[i] = node;
            node.__data__ = nodeData;
          } else if (!dataByKeyValue.has(keyValue)) {
            enterNodes[i] = d3_selection_dataNode(nodeData);
          }
          dataByKeyValue.set(keyValue, nodeData);
          nodeByKeyValue.remove(keyValue);
        }
        for (i = -1; ++i < n; ) {
          if (nodeByKeyValue.has(keyValues[i])) {
            exitNodes[i] = group[i];
          }
        }
      } else {
        for (i = -1; ++i < n0; ) {
          node = group[i];
          nodeData = groupData[i];
          if (node) {
            node.__data__ = nodeData;
            updateNodes[i] = node;
          } else {
            enterNodes[i] = d3_selection_dataNode(nodeData);
          }
        }
        for (;i < m; ++i) {
          enterNodes[i] = d3_selection_dataNode(groupData[i]);
        }
        for (;i < n; ++i) {
          exitNodes[i] = group[i];
        }
      }
      enterNodes.update = updateNodes;
      enterNodes.parentNode = updateNodes.parentNode = exitNodes.parentNode = group.parentNode;
      enter.push(enterNodes);
      update.push(updateNodes);
      exit.push(exitNodes);
    }
    var enter = d3_selection_enter([]), update = d3_selection([]), exit = d3_selection([]);
    if (typeof value === "function") {
      while (++i < n) {
        bind(group = this[i], value.call(group, group.parentNode.__data__, i));
      }
    } else {
      while (++i < n) {
        bind(group = this[i], value);
      }
    }
    update.enter = function() {
      return enter;
    };
    update.exit = function() {
      return exit;
    };
    return update;
  };
  function d3_selection_dataNode(data) {
    return {
      __data__: data
    };
  }
  d3_selectionPrototype.datum = function(value) {
    return arguments.length ? this.property("__data__", value) : this.property("__data__");
  };
  d3_selectionPrototype.filter = function(filter) {
    var subgroups = [], subgroup, group, node;
    if (typeof filter !== "function") filter = d3_selection_filter(filter);
    for (var j = 0, m = this.length; j < m; j++) {
      subgroups.push(subgroup = []);
      subgroup.parentNode = (group = this[j]).parentNode;
      for (var i = 0, n = group.length; i < n; i++) {
        if ((node = group[i]) && filter.call(node, node.__data__, i, j)) {
          subgroup.push(node);
        }
      }
    }
    return d3_selection(subgroups);
  };
  function d3_selection_filter(selector) {
    return function() {
      return d3_selectMatches(this, selector);
    };
  }
  d3_selectionPrototype.order = function() {
    for (var j = -1, m = this.length; ++j < m; ) {
      for (var group = this[j], i = group.length - 1, next = group[i], node; --i >= 0; ) {
        if (node = group[i]) {
          if (next && next !== node.nextSibling) next.parentNode.insertBefore(node, next);
          next = node;
        }
      }
    }
    return this;
  };
  d3_selectionPrototype.sort = function(comparator) {
    comparator = d3_selection_sortComparator.apply(this, arguments);
    for (var j = -1, m = this.length; ++j < m; ) this[j].sort(comparator);
    return this.order();
  };
  function d3_selection_sortComparator(comparator) {
    if (!arguments.length) comparator = d3.ascending;
    return function(a, b) {
      return a && b ? comparator(a.__data__, b.__data__) : !a - !b;
    };
  }
  d3_selectionPrototype.each = function(callback) {
    return d3_selection_each(this, function(node, i, j) {
      callback.call(node, node.__data__, i, j);
    });
  };
  function d3_selection_each(groups, callback) {
    for (var j = 0, m = groups.length; j < m; j++) {
      for (var group = groups[j], i = 0, n = group.length, node; i < n; i++) {
        if (node = group[i]) callback(node, i, j);
      }
    }
    return groups;
  }
  d3_selectionPrototype.call = function(callback) {
    var args = d3_array(arguments);
    callback.apply(args[0] = this, args);
    return this;
  };
  d3_selectionPrototype.empty = function() {
    return !this.node();
  };
  d3_selectionPrototype.node = function() {
    for (var j = 0, m = this.length; j < m; j++) {
      for (var group = this[j], i = 0, n = group.length; i < n; i++) {
        var node = group[i];
        if (node) return node;
      }
    }
    return null;
  };
  d3_selectionPrototype.size = function() {
    var n = 0;
    this.each(function() {
      ++n;
    });
    return n;
  };
  function d3_selection_enter(selection) {
    d3_subclass(selection, d3_selection_enterPrototype);
    return selection;
  }
  var d3_selection_enterPrototype = [];
  d3.selection.enter = d3_selection_enter;
  d3.selection.enter.prototype = d3_selection_enterPrototype;
  d3_selection_enterPrototype.append = d3_selectionPrototype.append;
  d3_selection_enterPrototype.empty = d3_selectionPrototype.empty;
  d3_selection_enterPrototype.node = d3_selectionPrototype.node;
  d3_selection_enterPrototype.call = d3_selectionPrototype.call;
  d3_selection_enterPrototype.size = d3_selectionPrototype.size;
  d3_selection_enterPrototype.select = function(selector) {
    var subgroups = [], subgroup, subnode, upgroup, group, node;
    for (var j = -1, m = this.length; ++j < m; ) {
      upgroup = (group = this[j]).update;
      subgroups.push(subgroup = []);
      subgroup.parentNode = group.parentNode;
      for (var i = -1, n = group.length; ++i < n; ) {
        if (node = group[i]) {
          subgroup.push(upgroup[i] = subnode = selector.call(group.parentNode, node.__data__, i, j));
          subnode.__data__ = node.__data__;
        } else {
          subgroup.push(null);
        }
      }
    }
    return d3_selection(subgroups);
  };
  d3_selection_enterPrototype.insert = function(name, before) {
    if (arguments.length < 2) before = d3_selection_enterInsertBefore(this);
    return d3_selectionPrototype.insert.call(this, name, before);
  };
  function d3_selection_enterInsertBefore(enter) {
    var i0, j0;
    return function(d, i, j) {
      var group = enter[j].update, n = group.length, node;
      if (j != j0) j0 = j, i0 = 0;
      if (i >= i0) i0 = i + 1;
      while (!(node = group[i0]) && ++i0 < n) ;
      return node;
    };
  }
  d3_selectionPrototype.transition = function() {
    var id = d3_transitionInheritId || ++d3_transitionId, subgroups = [], subgroup, node, transition = d3_transitionInherit || {
      time: Date.now(),
      ease: d3_ease_cubicInOut,
      delay: 0,
      duration: 250
    };
    for (var j = -1, m = this.length; ++j < m; ) {
      subgroups.push(subgroup = []);
      for (var group = this[j], i = -1, n = group.length; ++i < n; ) {
        if (node = group[i]) d3_transitionNode(node, i, id, transition);
        subgroup.push(node);
      }
    }
    return d3_transition(subgroups, id);
  };
  d3_selectionPrototype.interrupt = function() {
    return this.each(d3_selection_interrupt);
  };
  function d3_selection_interrupt() {
    var lock = this.__transition__;
    if (lock) ++lock.active;
  }
  d3.select = function(node) {
    var group = [ typeof node === "string" ? d3_select(node, d3_document) : node ];
    group.parentNode = d3_documentElement;
    return d3_selection([ group ]);
  };
  d3.selectAll = function(nodes) {
    var group = d3_array(typeof nodes === "string" ? d3_selectAll(nodes, d3_document) : nodes);
    group.parentNode = d3_documentElement;
    return d3_selection([ group ]);
  };
  var d3_selectionRoot = d3.select(d3_documentElement);
  d3_selectionPrototype.on = function(type, listener, capture) {
    var n = arguments.length;
    if (n < 3) {
      if (typeof type !== "string") {
        if (n < 2) listener = false;
        for (capture in type) this.each(d3_selection_on(capture, type[capture], listener));
        return this;
      }
      if (n < 2) return (n = this.node()["__on" + type]) && n._;
      capture = false;
    }
    return this.each(d3_selection_on(type, listener, capture));
  };
  function d3_selection_on(type, listener, capture) {
    var name = "__on" + type, i = type.indexOf("."), wrap = d3_selection_onListener;
    if (i > 0) type = type.substring(0, i);
    var filter = d3_selection_onFilters.get(type);
    if (filter) type = filter, wrap = d3_selection_onFilter;
    function onRemove() {
      var l = this[name];
      if (l) {
        this.removeEventListener(type, l, l.$);
        delete this[name];
      }
    }
    function onAdd() {
      var l = wrap(listener, d3_array(arguments));
      onRemove.call(this);
      this.addEventListener(type, this[name] = l, l.$ = capture);
      l._ = listener;
    }
    function removeAll() {
      var re = new RegExp("^__on([^.]+)" + d3.requote(type) + "$"), match;
      for (var name in this) {
        if (match = name.match(re)) {
          var l = this[name];
          this.removeEventListener(match[1], l, l.$);
          delete this[name];
        }
      }
    }
    return i ? listener ? onAdd : onRemove : listener ? d3_noop : removeAll;
  }
  var d3_selection_onFilters = d3.map({
    mouseenter: "mouseover",
    mouseleave: "mouseout"
  });
  d3_selection_onFilters.forEach(function(k) {
    if ("on" + k in d3_document) d3_selection_onFilters.remove(k);
  });
  function d3_selection_onListener(listener, argumentz) {
    return function(e) {
      var o = d3.event;
      d3.event = e;
      argumentz[0] = this.__data__;
      try {
        listener.apply(this, argumentz);
      } finally {
        d3.event = o;
      }
    };
  }
  function d3_selection_onFilter(listener, argumentz) {
    var l = d3_selection_onListener(listener, argumentz);
    return function(e) {
      var target = this, related = e.relatedTarget;
      if (!related || related !== target && !(related.compareDocumentPosition(target) & 8)) {
        l.call(target, e);
      }
    };
  }
  var d3_event_dragSelect = "onselectstart" in d3_document ? null : d3_vendorSymbol(d3_documentElement.style, "userSelect"), d3_event_dragId = 0;
  function d3_event_dragSuppress() {
    var name = ".dragsuppress-" + ++d3_event_dragId, click = "click" + name, w = d3.select(d3_window).on("touchmove" + name, d3_eventPreventDefault).on("dragstart" + name, d3_eventPreventDefault).on("selectstart" + name, d3_eventPreventDefault);
    if (d3_event_dragSelect) {
      var style = d3_documentElement.style, select = style[d3_event_dragSelect];
      style[d3_event_dragSelect] = "none";
    }
    return function(suppressClick) {
      w.on(name, null);
      if (d3_event_dragSelect) style[d3_event_dragSelect] = select;
      if (suppressClick) {
        function off() {
          w.on(click, null);
        }
        w.on(click, function() {
          d3_eventPreventDefault();
          off();
        }, true);
        setTimeout(off, 0);
      }
    };
  }
  d3.mouse = function(container) {
    return d3_mousePoint(container, d3_eventSource());
  };
  var d3_mouse_bug44083 = /WebKit/.test(d3_window.navigator.userAgent) ? -1 : 0;
  function d3_mousePoint(container, e) {
    if (e.changedTouches) e = e.changedTouches[0];
    var svg = container.ownerSVGElement || container;
    if (svg.createSVGPoint) {
      var point = svg.createSVGPoint();
      if (d3_mouse_bug44083 < 0 && (d3_window.scrollX || d3_window.scrollY)) {
        svg = d3.select("body").append("svg").style({
          position: "absolute",
          top: 0,
          left: 0,
          margin: 0,
          padding: 0,
          border: "none"
        }, "important");
        var ctm = svg[0][0].getScreenCTM();
        d3_mouse_bug44083 = !(ctm.f || ctm.e);
        svg.remove();
      }
      if (d3_mouse_bug44083) point.x = e.pageX, point.y = e.pageY; else point.x = e.clientX, 
      point.y = e.clientY;
      point = point.matrixTransform(container.getScreenCTM().inverse());
      return [ point.x, point.y ];
    }
    var rect = container.getBoundingClientRect();
    return [ e.clientX - rect.left - container.clientLeft, e.clientY - rect.top - container.clientTop ];
  }
  d3.touches = function(container, touches) {
    if (arguments.length < 2) touches = d3_eventSource().touches;
    return touches ? d3_array(touches).map(function(touch) {
      var point = d3_mousePoint(container, touch);
      point.identifier = touch.identifier;
      return point;
    }) : [];
  };
  d3.behavior.drag = function() {
    var event = d3_eventDispatch(drag, "drag", "dragstart", "dragend"), origin = null, mousedown = dragstart(d3_noop, d3.mouse, "mousemove", "mouseup"), touchstart = dragstart(touchid, touchposition, "touchmove", "touchend");
    function drag() {
      this.on("mousedown.drag", mousedown).on("touchstart.drag", touchstart);
    }
    function touchid() {
      return d3.event.changedTouches[0].identifier;
    }
    function touchposition(parent, id) {
      return d3.touches(parent).filter(function(p) {
        return p.identifier === id;
      })[0];
    }
    function dragstart(id, position, move, end) {
      return function() {
        var target = this, parent = target.parentNode, event_ = event.of(target, arguments), eventTarget = d3.event.target, eventId = id(), drag = eventId == null ? "drag" : "drag-" + eventId, origin_ = position(parent, eventId), dragged = 0, offset, w = d3.select(d3_window).on(move + "." + drag, moved).on(end + "." + drag, ended), dragRestore = d3_event_dragSuppress();
        if (origin) {
          offset = origin.apply(target, arguments);
          offset = [ offset.x - origin_[0], offset.y - origin_[1] ];
        } else {
          offset = [ 0, 0 ];
        }
        event_({
          type: "dragstart"
        });
        function moved() {
          var p = position(parent, eventId), dx = p[0] - origin_[0], dy = p[1] - origin_[1];
          dragged |= dx | dy;
          origin_ = p;
          event_({
            type: "drag",
            x: p[0] + offset[0],
            y: p[1] + offset[1],
            dx: dx,
            dy: dy
          });
        }
        function ended() {
          w.on(move + "." + drag, null).on(end + "." + drag, null);
          dragRestore(dragged && d3.event.target === eventTarget);
          event_({
            type: "dragend"
          });
        }
      };
    }
    drag.origin = function(x) {
      if (!arguments.length) return origin;
      origin = x;
      return drag;
    };
    return d3.rebind(drag, event, "on");
  };
  var π = Math.PI, τ = 2 * π, halfπ = π / 2, ε = 1e-6, ε2 = ε * ε, d3_radians = π / 180, d3_degrees = 180 / π;
  function d3_sgn(x) {
    return x > 0 ? 1 : x < 0 ? -1 : 0;
  }
  function d3_acos(x) {
    return x > 1 ? 0 : x < -1 ? π : Math.acos(x);
  }
  function d3_asin(x) {
    return x > 1 ? halfπ : x < -1 ? -halfπ : Math.asin(x);
  }
  function d3_sinh(x) {
    return ((x = Math.exp(x)) - 1 / x) / 2;
  }
  function d3_cosh(x) {
    return ((x = Math.exp(x)) + 1 / x) / 2;
  }
  function d3_tanh(x) {
    return ((x = Math.exp(2 * x)) - 1) / (x + 1);
  }
  function d3_haversin(x) {
    return (x = Math.sin(x / 2)) * x;
  }
  var ρ = Math.SQRT2, ρ2 = 2, ρ4 = 4;
  d3.interpolateZoom = function(p0, p1) {
    var ux0 = p0[0], uy0 = p0[1], w0 = p0[2], ux1 = p1[0], uy1 = p1[1], w1 = p1[2];
    var dx = ux1 - ux0, dy = uy1 - uy0, d2 = dx * dx + dy * dy, d1 = Math.sqrt(d2), b0 = (w1 * w1 - w0 * w0 + ρ4 * d2) / (2 * w0 * ρ2 * d1), b1 = (w1 * w1 - w0 * w0 - ρ4 * d2) / (2 * w1 * ρ2 * d1), r0 = Math.log(Math.sqrt(b0 * b0 + 1) - b0), r1 = Math.log(Math.sqrt(b1 * b1 + 1) - b1), dr = r1 - r0, S = (dr || Math.log(w1 / w0)) / ρ;
    function interpolate(t) {
      var s = t * S;
      if (dr) {
        var coshr0 = d3_cosh(r0), u = w0 / (ρ2 * d1) * (coshr0 * d3_tanh(ρ * s + r0) - d3_sinh(r0));
        return [ ux0 + u * dx, uy0 + u * dy, w0 * coshr0 / d3_cosh(ρ * s + r0) ];
      }
      return [ ux0 + t * dx, uy0 + t * dy, w0 * Math.exp(ρ * s) ];
    }
    interpolate.duration = S * 1e3;
    return interpolate;
  };
  d3.behavior.zoom = function() {
    var view = {
      x: 0,
      y: 0,
      k: 1
    }, translate0, center, size = [ 960, 500 ], scaleExtent = d3_behavior_zoomInfinity, mousedown = "mousedown.zoom", mousemove = "mousemove.zoom", mouseup = "mouseup.zoom", mousewheelTimer, touchstart = "touchstart.zoom", touchtime, event = d3_eventDispatch(zoom, "zoomstart", "zoom", "zoomend"), x0, x1, y0, y1;
    function zoom(g) {
      g.on(mousedown, mousedowned).on(d3_behavior_zoomWheel + ".zoom", mousewheeled).on(mousemove, mousewheelreset).on("dblclick.zoom", dblclicked).on(touchstart, touchstarted);
    }
    zoom.event = function(g) {
      g.each(function() {
        var event_ = event.of(this, arguments), view1 = view;
        if (d3_transitionInheritId) {
          d3.select(this).transition().each("start.zoom", function() {
            view = this.__chart__ || {
              x: 0,
              y: 0,
              k: 1
            };
            zoomstarted(event_);
          }).tween("zoom:zoom", function() {
            var dx = size[0], dy = size[1], cx = dx / 2, cy = dy / 2, i = d3.interpolateZoom([ (cx - view.x) / view.k, (cy - view.y) / view.k, dx / view.k ], [ (cx - view1.x) / view1.k, (cy - view1.y) / view1.k, dx / view1.k ]);
            return function(t) {
              var l = i(t), k = dx / l[2];
              this.__chart__ = view = {
                x: cx - l[0] * k,
                y: cy - l[1] * k,
                k: k
              };
              zoomed(event_);
            };
          }).each("end.zoom", function() {
            zoomended(event_);
          });
        } else {
          this.__chart__ = view;
          zoomstarted(event_);
          zoomed(event_);
          zoomended(event_);
        }
      });
    };
    zoom.translate = function(_) {
      if (!arguments.length) return [ view.x, view.y ];
      view = {
        x: +_[0],
        y: +_[1],
        k: view.k
      };
      rescale();
      return zoom;
    };
    zoom.scale = function(_) {
      if (!arguments.length) return view.k;
      view = {
        x: view.x,
        y: view.y,
        k: +_
      };
      rescale();
      return zoom;
    };
    zoom.scaleExtent = function(_) {
      if (!arguments.length) return scaleExtent;
      scaleExtent = _ == null ? d3_behavior_zoomInfinity : [ +_[0], +_[1] ];
      return zoom;
    };
    zoom.center = function(_) {
      if (!arguments.length) return center;
      center = _ && [ +_[0], +_[1] ];
      return zoom;
    };
    zoom.size = function(_) {
      if (!arguments.length) return size;
      size = _ && [ +_[0], +_[1] ];
      return zoom;
    };
    zoom.x = function(z) {
      if (!arguments.length) return x1;
      x1 = z;
      x0 = z.copy();
      view = {
        x: 0,
        y: 0,
        k: 1
      };
      return zoom;
    };
    zoom.y = function(z) {
      if (!arguments.length) return y1;
      y1 = z;
      y0 = z.copy();
      view = {
        x: 0,
        y: 0,
        k: 1
      };
      return zoom;
    };
    function location(p) {
      return [ (p[0] - view.x) / view.k, (p[1] - view.y) / view.k ];
    }
    function point(l) {
      return [ l[0] * view.k + view.x, l[1] * view.k + view.y ];
    }
    function scaleTo(s) {
      view.k = Math.max(scaleExtent[0], Math.min(scaleExtent[1], s));
    }
    function translateTo(p, l) {
      l = point(l);
      view.x += p[0] - l[0];
      view.y += p[1] - l[1];
    }
    function rescale() {
      if (x1) x1.domain(x0.range().map(function(x) {
        return (x - view.x) / view.k;
      }).map(x0.invert));
      if (y1) y1.domain(y0.range().map(function(y) {
        return (y - view.y) / view.k;
      }).map(y0.invert));
    }
    function zoomstarted(event) {
      event({
        type: "zoomstart"
      });
    }
    function zoomed(event) {
      rescale();
      event({
        type: "zoom",
        scale: view.k,
        translate: [ view.x, view.y ]
      });
    }
    function zoomended(event) {
      event({
        type: "zoomend"
      });
    }
    function mousedowned() {
      var target = this, event_ = event.of(target, arguments), eventTarget = d3.event.target, dragged = 0, w = d3.select(d3_window).on(mousemove, moved).on(mouseup, ended), l = location(d3.mouse(target)), dragRestore = d3_event_dragSuppress();
      d3_selection_interrupt.call(target);
      zoomstarted(event_);
      function moved() {
        dragged = 1;
        translateTo(d3.mouse(target), l);
        zoomed(event_);
      }
      function ended() {
        w.on(mousemove, d3_window === target ? mousewheelreset : null).on(mouseup, null);
        dragRestore(dragged && d3.event.target === eventTarget);
        zoomended(event_);
      }
    }
    function touchstarted() {
      var target = this, event_ = event.of(target, arguments), locations0 = {}, distance0 = 0, scale0, eventId = d3.event.changedTouches[0].identifier, touchmove = "touchmove.zoom-" + eventId, touchend = "touchend.zoom-" + eventId, w = d3.select(d3_window).on(touchmove, moved).on(touchend, ended), t = d3.select(target).on(mousedown, null).on(touchstart, started), dragRestore = d3_event_dragSuppress();
      d3_selection_interrupt.call(target);
      started();
      zoomstarted(event_);
      function relocate() {
        var touches = d3.touches(target);
        scale0 = view.k;
        touches.forEach(function(t) {
          if (t.identifier in locations0) locations0[t.identifier] = location(t);
        });
        return touches;
      }
      function started() {
        var changed = d3.event.changedTouches;
        for (var i = 0, n = changed.length; i < n; ++i) {
          locations0[changed[i].identifier] = null;
        }
        var touches = relocate(), now = Date.now();
        if (touches.length === 1) {
          if (now - touchtime < 500) {
            var p = touches[0], l = locations0[p.identifier];
            scaleTo(view.k * 2);
            translateTo(p, l);
            d3_eventPreventDefault();
            zoomed(event_);
          }
          touchtime = now;
        } else if (touches.length > 1) {
          var p = touches[0], q = touches[1], dx = p[0] - q[0], dy = p[1] - q[1];
          distance0 = dx * dx + dy * dy;
        }
      }
      function moved() {
        var touches = d3.touches(target), p0, l0, p1, l1;
        for (var i = 0, n = touches.length; i < n; ++i, l1 = null) {
          p1 = touches[i];
          if (l1 = locations0[p1.identifier]) {
            if (l0) break;
            p0 = p1, l0 = l1;
          }
        }
        if (l1) {
          var distance1 = (distance1 = p1[0] - p0[0]) * distance1 + (distance1 = p1[1] - p0[1]) * distance1, scale1 = distance0 && Math.sqrt(distance1 / distance0);
          p0 = [ (p0[0] + p1[0]) / 2, (p0[1] + p1[1]) / 2 ];
          l0 = [ (l0[0] + l1[0]) / 2, (l0[1] + l1[1]) / 2 ];
          scaleTo(scale1 * scale0);
        }
        touchtime = null;
        translateTo(p0, l0);
        zoomed(event_);
      }
      function ended() {
        if (d3.event.touches.length) {
          var changed = d3.event.changedTouches;
          for (var i = 0, n = changed.length; i < n; ++i) {
            delete locations0[changed[i].identifier];
          }
          for (var identifier in locations0) {
            return void relocate();
          }
        }
        w.on(touchmove, null).on(touchend, null);
        t.on(mousedown, mousedowned).on(touchstart, touchstarted);
        dragRestore();
        zoomended(event_);
      }
    }
    function mousewheeled() {
      var event_ = event.of(this, arguments);
      if (mousewheelTimer) clearTimeout(mousewheelTimer); else d3_selection_interrupt.call(this), 
      zoomstarted(event_);
      mousewheelTimer = setTimeout(function() {
        mousewheelTimer = null;
        zoomended(event_);
      }, 50);
      d3_eventPreventDefault();
      var point = center || d3.mouse(this);
      if (!translate0) translate0 = location(point);
      scaleTo(Math.pow(2, d3_behavior_zoomDelta() * .002) * view.k);
      translateTo(point, translate0);
      zoomed(event_);
    }
    function mousewheelreset() {
      translate0 = null;
    }
    function dblclicked() {
      var event_ = event.of(this, arguments), p = d3.mouse(this), l = location(p), k = Math.log(view.k) / Math.LN2;
      zoomstarted(event_);
      scaleTo(Math.pow(2, d3.event.shiftKey ? Math.ceil(k) - 1 : Math.floor(k) + 1));
      translateTo(p, l);
      zoomed(event_);
      zoomended(event_);
    }
    return d3.rebind(zoom, event, "on");
  };
  var d3_behavior_zoomInfinity = [ 0, Infinity ];
  var d3_behavior_zoomDelta, d3_behavior_zoomWheel = "onwheel" in d3_document ? (d3_behavior_zoomDelta = function() {
    return -d3.event.deltaY * (d3.event.deltaMode ? 120 : 1);
  }, "wheel") : "onmousewheel" in d3_document ? (d3_behavior_zoomDelta = function() {
    return d3.event.wheelDelta;
  }, "mousewheel") : (d3_behavior_zoomDelta = function() {
    return -d3.event.detail;
  }, "MozMousePixelScroll");
  function d3_Color() {}
  d3_Color.prototype.toString = function() {
    return this.rgb() + "";
  };
  d3.hsl = function(h, s, l) {
    return arguments.length === 1 ? h instanceof d3_Hsl ? d3_hsl(h.h, h.s, h.l) : d3_rgb_parse("" + h, d3_rgb_hsl, d3_hsl) : d3_hsl(+h, +s, +l);
  };
  function d3_hsl(h, s, l) {
    return new d3_Hsl(h, s, l);
  }
  function d3_Hsl(h, s, l) {
    this.h = h;
    this.s = s;
    this.l = l;
  }
  var d3_hslPrototype = d3_Hsl.prototype = new d3_Color();
  d3_hslPrototype.brighter = function(k) {
    k = Math.pow(.7, arguments.length ? k : 1);
    return d3_hsl(this.h, this.s, this.l / k);
  };
  d3_hslPrototype.darker = function(k) {
    k = Math.pow(.7, arguments.length ? k : 1);
    return d3_hsl(this.h, this.s, k * this.l);
  };
  d3_hslPrototype.rgb = function() {
    return d3_hsl_rgb(this.h, this.s, this.l);
  };
  function d3_hsl_rgb(h, s, l) {
    var m1, m2;
    h = isNaN(h) ? 0 : (h %= 360) < 0 ? h + 360 : h;
    s = isNaN(s) ? 0 : s < 0 ? 0 : s > 1 ? 1 : s;
    l = l < 0 ? 0 : l > 1 ? 1 : l;
    m2 = l <= .5 ? l * (1 + s) : l + s - l * s;
    m1 = 2 * l - m2;
    function v(h) {
      if (h > 360) h -= 360; else if (h < 0) h += 360;
      if (h < 60) return m1 + (m2 - m1) * h / 60;
      if (h < 180) return m2;
      if (h < 240) return m1 + (m2 - m1) * (240 - h) / 60;
      return m1;
    }
    function vv(h) {
      return Math.round(v(h) * 255);
    }
    return d3_rgb(vv(h + 120), vv(h), vv(h - 120));
  }
  d3.hcl = function(h, c, l) {
    return arguments.length === 1 ? h instanceof d3_Hcl ? d3_hcl(h.h, h.c, h.l) : h instanceof d3_Lab ? d3_lab_hcl(h.l, h.a, h.b) : d3_lab_hcl((h = d3_rgb_lab((h = d3.rgb(h)).r, h.g, h.b)).l, h.a, h.b) : d3_hcl(+h, +c, +l);
  };
  function d3_hcl(h, c, l) {
    return new d3_Hcl(h, c, l);
  }
  function d3_Hcl(h, c, l) {
    this.h = h;
    this.c = c;
    this.l = l;
  }
  var d3_hclPrototype = d3_Hcl.prototype = new d3_Color();
  d3_hclPrototype.brighter = function(k) {
    return d3_hcl(this.h, this.c, Math.min(100, this.l + d3_lab_K * (arguments.length ? k : 1)));
  };
  d3_hclPrototype.darker = function(k) {
    return d3_hcl(this.h, this.c, Math.max(0, this.l - d3_lab_K * (arguments.length ? k : 1)));
  };
  d3_hclPrototype.rgb = function() {
    return d3_hcl_lab(this.h, this.c, this.l).rgb();
  };
  function d3_hcl_lab(h, c, l) {
    if (isNaN(h)) h = 0;
    if (isNaN(c)) c = 0;
    return d3_lab(l, Math.cos(h *= d3_radians) * c, Math.sin(h) * c);
  }
  d3.lab = function(l, a, b) {
    return arguments.length === 1 ? l instanceof d3_Lab ? d3_lab(l.l, l.a, l.b) : l instanceof d3_Hcl ? d3_hcl_lab(l.l, l.c, l.h) : d3_rgb_lab((l = d3.rgb(l)).r, l.g, l.b) : d3_lab(+l, +a, +b);
  };
  function d3_lab(l, a, b) {
    return new d3_Lab(l, a, b);
  }
  function d3_Lab(l, a, b) {
    this.l = l;
    this.a = a;
    this.b = b;
  }
  var d3_lab_K = 18;
  var d3_lab_X = .95047, d3_lab_Y = 1, d3_lab_Z = 1.08883;
  var d3_labPrototype = d3_Lab.prototype = new d3_Color();
  d3_labPrototype.brighter = function(k) {
    return d3_lab(Math.min(100, this.l + d3_lab_K * (arguments.length ? k : 1)), this.a, this.b);
  };
  d3_labPrototype.darker = function(k) {
    return d3_lab(Math.max(0, this.l - d3_lab_K * (arguments.length ? k : 1)), this.a, this.b);
  };
  d3_labPrototype.rgb = function() {
    return d3_lab_rgb(this.l, this.a, this.b);
  };
  function d3_lab_rgb(l, a, b) {
    var y = (l + 16) / 116, x = y + a / 500, z = y - b / 200;
    x = d3_lab_xyz(x) * d3_lab_X;
    y = d3_lab_xyz(y) * d3_lab_Y;
    z = d3_lab_xyz(z) * d3_lab_Z;
    return d3_rgb(d3_xyz_rgb(3.2404542 * x - 1.5371385 * y - .4985314 * z), d3_xyz_rgb(-.969266 * x + 1.8760108 * y + .041556 * z), d3_xyz_rgb(.0556434 * x - .2040259 * y + 1.0572252 * z));
  }
  function d3_lab_hcl(l, a, b) {
    return l > 0 ? d3_hcl(Math.atan2(b, a) * d3_degrees, Math.sqrt(a * a + b * b), l) : d3_hcl(NaN, NaN, l);
  }
  function d3_lab_xyz(x) {
    return x > .206893034 ? x * x * x : (x - 4 / 29) / 7.787037;
  }
  function d3_xyz_lab(x) {
    return x > .008856 ? Math.pow(x, 1 / 3) : 7.787037 * x + 4 / 29;
  }
  function d3_xyz_rgb(r) {
    return Math.round(255 * (r <= .00304 ? 12.92 * r : 1.055 * Math.pow(r, 1 / 2.4) - .055));
  }
  d3.rgb = function(r, g, b) {
    return arguments.length === 1 ? r instanceof d3_Rgb ? d3_rgb(r.r, r.g, r.b) : d3_rgb_parse("" + r, d3_rgb, d3_hsl_rgb) : d3_rgb(~~r, ~~g, ~~b);
  };
  function d3_rgbNumber(value) {
    return d3_rgb(value >> 16, value >> 8 & 255, value & 255);
  }
  function d3_rgbString(value) {
    return d3_rgbNumber(value) + "";
  }
  function d3_rgb(r, g, b) {
    return new d3_Rgb(r, g, b);
  }
  function d3_Rgb(r, g, b) {
    this.r = r;
    this.g = g;
    this.b = b;
  }
  var d3_rgbPrototype = d3_Rgb.prototype = new d3_Color();
  d3_rgbPrototype.brighter = function(k) {
    k = Math.pow(.7, arguments.length ? k : 1);
    var r = this.r, g = this.g, b = this.b, i = 30;
    if (!r && !g && !b) return d3_rgb(i, i, i);
    if (r && r < i) r = i;
    if (g && g < i) g = i;
    if (b && b < i) b = i;
    return d3_rgb(Math.min(255, ~~(r / k)), Math.min(255, ~~(g / k)), Math.min(255, ~~(b / k)));
  };
  d3_rgbPrototype.darker = function(k) {
    k = Math.pow(.7, arguments.length ? k : 1);
    return d3_rgb(~~(k * this.r), ~~(k * this.g), ~~(k * this.b));
  };
  d3_rgbPrototype.hsl = function() {
    return d3_rgb_hsl(this.r, this.g, this.b);
  };
  d3_rgbPrototype.toString = function() {
    return "#" + d3_rgb_hex(this.r) + d3_rgb_hex(this.g) + d3_rgb_hex(this.b);
  };
  function d3_rgb_hex(v) {
    return v < 16 ? "0" + Math.max(0, v).toString(16) : Math.min(255, v).toString(16);
  }
  function d3_rgb_parse(format, rgb, hsl) {
    var r = 0, g = 0, b = 0, m1, m2, name;
    m1 = /([a-z]+)\((.*)\)/i.exec(format);
    if (m1) {
      m2 = m1[2].split(",");
      switch (m1[1]) {
       case "hsl":
        {
          return hsl(parseFloat(m2[0]), parseFloat(m2[1]) / 100, parseFloat(m2[2]) / 100);
        }

       case "rgb":
        {
          return rgb(d3_rgb_parseNumber(m2[0]), d3_rgb_parseNumber(m2[1]), d3_rgb_parseNumber(m2[2]));
        }
      }
    }
    if (name = d3_rgb_names.get(format)) return rgb(name.r, name.g, name.b);
    if (format != null && format.charAt(0) === "#") {
      if (format.length === 4) {
        r = format.charAt(1);
        r += r;
        g = format.charAt(2);
        g += g;
        b = format.charAt(3);
        b += b;
      } else if (format.length === 7) {
        r = format.substring(1, 3);
        g = format.substring(3, 5);
        b = format.substring(5, 7);
      }
      r = parseInt(r, 16);
      g = parseInt(g, 16);
      b = parseInt(b, 16);
    }
    return rgb(r, g, b);
  }
  function d3_rgb_hsl(r, g, b) {
    var min = Math.min(r /= 255, g /= 255, b /= 255), max = Math.max(r, g, b), d = max - min, h, s, l = (max + min) / 2;
    if (d) {
      s = l < .5 ? d / (max + min) : d / (2 - max - min);
      if (r == max) h = (g - b) / d + (g < b ? 6 : 0); else if (g == max) h = (b - r) / d + 2; else h = (r - g) / d + 4;
      h *= 60;
    } else {
      h = NaN;
      s = l > 0 && l < 1 ? 0 : h;
    }
    return d3_hsl(h, s, l);
  }
  function d3_rgb_lab(r, g, b) {
    r = d3_rgb_xyz(r);
    g = d3_rgb_xyz(g);
    b = d3_rgb_xyz(b);
    var x = d3_xyz_lab((.4124564 * r + .3575761 * g + .1804375 * b) / d3_lab_X), y = d3_xyz_lab((.2126729 * r + .7151522 * g + .072175 * b) / d3_lab_Y), z = d3_xyz_lab((.0193339 * r + .119192 * g + .9503041 * b) / d3_lab_Z);
    return d3_lab(116 * y - 16, 500 * (x - y), 200 * (y - z));
  }
  function d3_rgb_xyz(r) {
    return (r /= 255) <= .04045 ? r / 12.92 : Math.pow((r + .055) / 1.055, 2.4);
  }
  function d3_rgb_parseNumber(c) {
    var f = parseFloat(c);
    return c.charAt(c.length - 1) === "%" ? Math.round(f * 2.55) : f;
  }
  var d3_rgb_names = d3.map({
    aliceblue: 15792383,
    antiquewhite: 16444375,
    aqua: 65535,
    aquamarine: 8388564,
    azure: 15794175,
    beige: 16119260,
    bisque: 16770244,
    black: 0,
    blanchedalmond: 16772045,
    blue: 255,
    blueviolet: 9055202,
    brown: 10824234,
    burlywood: 14596231,
    cadetblue: 6266528,
    chartreuse: 8388352,
    chocolate: 13789470,
    coral: 16744272,
    cornflowerblue: 6591981,
    cornsilk: 16775388,
    crimson: 14423100,
    cyan: 65535,
    darkblue: 139,
    darkcyan: 35723,
    darkgoldenrod: 12092939,
    darkgray: 11119017,
    darkgreen: 25600,
    darkgrey: 11119017,
    darkkhaki: 12433259,
    darkmagenta: 9109643,
    darkolivegreen: 5597999,
    darkorange: 16747520,
    darkorchid: 10040012,
    darkred: 9109504,
    darksalmon: 15308410,
    darkseagreen: 9419919,
    darkslateblue: 4734347,
    darkslategray: 3100495,
    darkslategrey: 3100495,
    darkturquoise: 52945,
    darkviolet: 9699539,
    deeppink: 16716947,
    deepskyblue: 49151,
    dimgray: 6908265,
    dimgrey: 6908265,
    dodgerblue: 2003199,
    firebrick: 11674146,
    floralwhite: 16775920,
    forestgreen: 2263842,
    fuchsia: 16711935,
    gainsboro: 14474460,
    ghostwhite: 16316671,
    gold: 16766720,
    goldenrod: 14329120,
    gray: 8421504,
    green: 32768,
    greenyellow: 11403055,
    grey: 8421504,
    honeydew: 15794160,
    hotpink: 16738740,
    indianred: 13458524,
    indigo: 4915330,
    ivory: 16777200,
    khaki: 15787660,
    lavender: 15132410,
    lavenderblush: 16773365,
    lawngreen: 8190976,
    lemonchiffon: 16775885,
    lightblue: 11393254,
    lightcoral: 15761536,
    lightcyan: 14745599,
    lightgoldenrodyellow: 16448210,
    lightgray: 13882323,
    lightgreen: 9498256,
    lightgrey: 13882323,
    lightpink: 16758465,
    lightsalmon: 16752762,
    lightseagreen: 2142890,
    lightskyblue: 8900346,
    lightslategray: 7833753,
    lightslategrey: 7833753,
    lightsteelblue: 11584734,
    lightyellow: 16777184,
    lime: 65280,
    limegreen: 3329330,
    linen: 16445670,
    magenta: 16711935,
    maroon: 8388608,
    mediumaquamarine: 6737322,
    mediumblue: 205,
    mediumorchid: 12211667,
    mediumpurple: 9662683,
    mediumseagreen: 3978097,
    mediumslateblue: 8087790,
    mediumspringgreen: 64154,
    mediumturquoise: 4772300,
    mediumvioletred: 13047173,
    midnightblue: 1644912,
    mintcream: 16121850,
    mistyrose: 16770273,
    moccasin: 16770229,
    navajowhite: 16768685,
    navy: 128,
    oldlace: 16643558,
    olive: 8421376,
    olivedrab: 7048739,
    orange: 16753920,
    orangered: 16729344,
    orchid: 14315734,
    palegoldenrod: 15657130,
    palegreen: 10025880,
    paleturquoise: 11529966,
    palevioletred: 14381203,
    papayawhip: 16773077,
    peachpuff: 16767673,
    peru: 13468991,
    pink: 16761035,
    plum: 14524637,
    powderblue: 11591910,
    purple: 8388736,
    red: 16711680,
    rosybrown: 12357519,
    royalblue: 4286945,
    saddlebrown: 9127187,
    salmon: 16416882,
    sandybrown: 16032864,
    seagreen: 3050327,
    seashell: 16774638,
    sienna: 10506797,
    silver: 12632256,
    skyblue: 8900331,
    slateblue: 6970061,
    slategray: 7372944,
    slategrey: 7372944,
    snow: 16775930,
    springgreen: 65407,
    steelblue: 4620980,
    tan: 13808780,
    teal: 32896,
    thistle: 14204888,
    tomato: 16737095,
    turquoise: 4251856,
    violet: 15631086,
    wheat: 16113331,
    white: 16777215,
    whitesmoke: 16119285,
    yellow: 16776960,
    yellowgreen: 10145074
  });
  d3_rgb_names.forEach(function(key, value) {
    d3_rgb_names.set(key, d3_rgbNumber(value));
  });
  function d3_functor(v) {
    return typeof v === "function" ? v : function() {
      return v;
    };
  }
  d3.functor = d3_functor;
  function d3_identity(d) {
    return d;
  }
  d3.xhr = d3_xhrType(d3_identity);
  function d3_xhrType(response) {
    return function(url, mimeType, callback) {
      if (arguments.length === 2 && typeof mimeType === "function") callback = mimeType, 
      mimeType = null;
      return d3_xhr(url, mimeType, response, callback);
    };
  }
  function d3_xhr(url, mimeType, response, callback) {
    var xhr = {}, dispatch = d3.dispatch("beforesend", "progress", "load", "error"), headers = {}, request = new XMLHttpRequest(), responseType = null;
    if (d3_window.XDomainRequest && !("withCredentials" in request) && /^(http(s)?:)?\/\//.test(url)) request = new XDomainRequest();
    "onload" in request ? request.onload = request.onerror = respond : request.onreadystatechange = function() {
      request.readyState > 3 && respond();
    };
    function respond() {
      var status = request.status, result;
      if (!status && request.responseText || status >= 200 && status < 300 || status === 304) {
        try {
          result = response.call(xhr, request);
        } catch (e) {
          dispatch.error.call(xhr, e);
          return;
        }
        dispatch.load.call(xhr, result);
      } else {
        dispatch.error.call(xhr, request);
      }
    }
    request.onprogress = function(event) {
      var o = d3.event;
      d3.event = event;
      try {
        dispatch.progress.call(xhr, request);
      } finally {
        d3.event = o;
      }
    };
    xhr.header = function(name, value) {
      name = (name + "").toLowerCase();
      if (arguments.length < 2) return headers[name];
      if (value == null) delete headers[name]; else headers[name] = value + "";
      return xhr;
    };
    xhr.mimeType = function(value) {
      if (!arguments.length) return mimeType;
      mimeType = value == null ? null : value + "";
      return xhr;
    };
    xhr.responseType = function(value) {
      if (!arguments.length) return responseType;
      responseType = value;
      return xhr;
    };
    xhr.response = function(value) {
      response = value;
      return xhr;
    };
    [ "get", "post" ].forEach(function(method) {
      xhr[method] = function() {
        return xhr.send.apply(xhr, [ method ].concat(d3_array(arguments)));
      };
    });
    xhr.send = function(method, data, callback) {
      if (arguments.length === 2 && typeof data === "function") callback = data, data = null;
      request.open(method, url, true);
      if (mimeType != null && !("accept" in headers)) headers["accept"] = mimeType + ",*/*";
      if (request.setRequestHeader) for (var name in headers) request.setRequestHeader(name, headers[name]);
      if (mimeType != null && request.overrideMimeType) request.overrideMimeType(mimeType);
      if (responseType != null) request.responseType = responseType;
      if (callback != null) xhr.on("error", callback).on("load", function(request) {
        callback(null, request);
      });
      dispatch.beforesend.call(xhr, request);
      request.send(data == null ? null : data);
      return xhr;
    };
    xhr.abort = function() {
      request.abort();
      return xhr;
    };
    d3.rebind(xhr, dispatch, "on");
    return callback == null ? xhr : xhr.get(d3_xhr_fixCallback(callback));
  }
  function d3_xhr_fixCallback(callback) {
    return callback.length === 1 ? function(error, request) {
      callback(error == null ? request : null);
    } : callback;
  }
  d3.dsv = function(delimiter, mimeType) {
    var reFormat = new RegExp('["' + delimiter + "\n]"), delimiterCode = delimiter.charCodeAt(0);
    function dsv(url, row, callback) {
      if (arguments.length < 3) callback = row, row = null;
      var xhr = d3.xhr(url, mimeType, callback);
      xhr.row = function(_) {
        return arguments.length ? xhr.response((row = _) == null ? response : typedResponse(_)) : row;
      };
      return xhr.row(row);
    }
    function response(request) {
      return dsv.parse(request.responseText);
    }
    function typedResponse(f) {
      return function(request) {
        return dsv.parse(request.responseText, f);
      };
    }
    dsv.parse = function(text, f) {
      var o;
      return dsv.parseRows(text, function(row, i) {
        if (o) return o(row, i - 1);
        var a = new Function("d", "return {" + row.map(function(name, i) {
          return JSON.stringify(name) + ": d[" + i + "]";
        }).join(",") + "}");
        o = f ? function(row, i) {
          return f(a(row), i);
        } : a;
      });
    };
    dsv.parseRows = function(text, f) {
      var EOL = {}, EOF = {}, rows = [], N = text.length, I = 0, n = 0, t, eol;
      function token() {
        if (I >= N) return EOF;
        if (eol) return eol = false, EOL;
        var j = I;
        if (text.charCodeAt(j) === 34) {
          var i = j;
          while (i++ < N) {
            if (text.charCodeAt(i) === 34) {
              if (text.charCodeAt(i + 1) !== 34) break;
              ++i;
            }
          }
          I = i + 2;
          var c = text.charCodeAt(i + 1);
          if (c === 13) {
            eol = true;
            if (text.charCodeAt(i + 2) === 10) ++I;
          } else if (c === 10) {
            eol = true;
          }
          return text.substring(j + 1, i).replace(/""/g, '"');
        }
        while (I < N) {
          var c = text.charCodeAt(I++), k = 1;
          if (c === 10) eol = true; else if (c === 13) {
            eol = true;
            if (text.charCodeAt(I) === 10) ++I, ++k;
          } else if (c !== delimiterCode) continue;
          return text.substring(j, I - k);
        }
        return text.substring(j);
      }
      while ((t = token()) !== EOF) {
        var a = [];
        while (t !== EOL && t !== EOF) {
          a.push(t);
          t = token();
        }
        if (f && !(a = f(a, n++))) continue;
        rows.push(a);
      }
      return rows;
    };
    dsv.format = function(rows) {
      if (Array.isArray(rows[0])) return dsv.formatRows(rows);
      var fieldSet = new d3_Set(), fields = [];
      rows.forEach(function(row) {
        for (var field in row) {
          if (!fieldSet.has(field)) {
            fields.push(fieldSet.add(field));
          }
        }
      });
      return [ fields.map(formatValue).join(delimiter) ].concat(rows.map(function(row) {
        return fields.map(function(field) {
          return formatValue(row[field]);
        }).join(delimiter);
      })).join("\n");
    };
    dsv.formatRows = function(rows) {
      return rows.map(formatRow).join("\n");
    };
    function formatRow(row) {
      return row.map(formatValue).join(delimiter);
    }
    function formatValue(text) {
      return reFormat.test(text) ? '"' + text.replace(/\"/g, '""') + '"' : text;
    }
    return dsv;
  };
  d3.csv = d3.dsv(",", "text/csv");
  d3.tsv = d3.dsv("	", "text/tab-separated-values");
  var d3_timer_queueHead, d3_timer_queueTail, d3_timer_interval, d3_timer_timeout, d3_timer_active, d3_timer_frame = d3_window[d3_vendorSymbol(d3_window, "requestAnimationFrame")] || function(callback) {
    setTimeout(callback, 17);
  };
  d3.timer = function(callback, delay, then) {
    var n = arguments.length;
    if (n < 2) delay = 0;
    if (n < 3) then = Date.now();
    var time = then + delay, timer = {
      c: callback,
      t: time,
      f: false,
      n: null
    };
    if (d3_timer_queueTail) d3_timer_queueTail.n = timer; else d3_timer_queueHead = timer;
    d3_timer_queueTail = timer;
    if (!d3_timer_interval) {
      d3_timer_timeout = clearTimeout(d3_timer_timeout);
      d3_timer_interval = 1;
      d3_timer_frame(d3_timer_step);
    }
  };
  function d3_timer_step() {
    var now = d3_timer_mark(), delay = d3_timer_sweep() - now;
    if (delay > 24) {
      if (isFinite(delay)) {
        clearTimeout(d3_timer_timeout);
        d3_timer_timeout = setTimeout(d3_timer_step, delay);
      }
      d3_timer_interval = 0;
    } else {
      d3_timer_interval = 1;
      d3_timer_frame(d3_timer_step);
    }
  }
  d3.timer.flush = function() {
    d3_timer_mark();
    d3_timer_sweep();
  };
  function d3_timer_mark() {
    var now = Date.now();
    d3_timer_active = d3_timer_queueHead;
    while (d3_timer_active) {
      if (now >= d3_timer_active.t) d3_timer_active.f = d3_timer_active.c(now - d3_timer_active.t);
      d3_timer_active = d3_timer_active.n;
    }
    return now;
  }
  function d3_timer_sweep() {
    var t0, t1 = d3_timer_queueHead, time = Infinity;
    while (t1) {
      if (t1.f) {
        t1 = t0 ? t0.n = t1.n : d3_timer_queueHead = t1.n;
      } else {
        if (t1.t < time) time = t1.t;
        t1 = (t0 = t1).n;
      }
    }
    d3_timer_queueTail = t0;
    return time;
  }
  var d3_format_decimalPoint = ".", d3_format_thousandsSeparator = ",", d3_format_grouping = [ 3, 3 ], d3_format_currencySymbol = "$";
  var d3_formatPrefixes = [ "y", "z", "a", "f", "p", "n", "µ", "m", "", "k", "M", "G", "T", "P", "E", "Z", "Y" ].map(d3_formatPrefix);
  d3.formatPrefix = function(value, precision) {
    var i = 0;
    if (value) {
      if (value < 0) value *= -1;
      if (precision) value = d3.round(value, d3_format_precision(value, precision));
      i = 1 + Math.floor(1e-12 + Math.log(value) / Math.LN10);
      i = Math.max(-24, Math.min(24, Math.floor((i <= 0 ? i + 1 : i - 1) / 3) * 3));
    }
    return d3_formatPrefixes[8 + i / 3];
  };
  function d3_formatPrefix(d, i) {
    var k = Math.pow(10, abs(8 - i) * 3);
    return {
      scale: i > 8 ? function(d) {
        return d / k;
      } : function(d) {
        return d * k;
      },
      symbol: d
    };
  }
  d3.round = function(x, n) {
    return n ? Math.round(x * (n = Math.pow(10, n))) / n : Math.round(x);
  };
  d3.format = function(specifier) {
    var match = d3_format_re.exec(specifier), fill = match[1] || " ", align = match[2] || ">", sign = match[3] || "", symbol = match[4] || "", zfill = match[5], width = +match[6], comma = match[7], precision = match[8], type = match[9], scale = 1, suffix = "", integer = false;
    if (precision) precision = +precision.substring(1);
    if (zfill || fill === "0" && align === "=") {
      zfill = fill = "0";
      align = "=";
      if (comma) width -= Math.floor((width - 1) / 4);
    }
    switch (type) {
     case "n":
      comma = true;
      type = "g";
      break;

     case "%":
      scale = 100;
      suffix = "%";
      type = "f";
      break;

     case "p":
      scale = 100;
      suffix = "%";
      type = "r";
      break;

     case "b":
     case "o":
     case "x":
     case "X":
      if (symbol === "#") symbol = "0" + type.toLowerCase();

     case "c":
     case "d":
      integer = true;
      precision = 0;
      break;

     case "s":
      scale = -1;
      type = "r";
      break;
    }
    if (symbol === "#") symbol = ""; else if (symbol === "$") symbol = d3_format_currencySymbol;
    if (type == "r" && !precision) type = "g";
    if (precision != null) {
      if (type == "g") precision = Math.max(1, Math.min(21, precision)); else if (type == "e" || type == "f") precision = Math.max(0, Math.min(20, precision));
    }
    type = d3_format_types.get(type) || d3_format_typeDefault;
    var zcomma = zfill && comma;
    return function(value) {
      if (integer && value % 1) return "";
      var negative = value < 0 || value === 0 && 1 / value < 0 ? (value = -value, "-") : sign;
      if (scale < 0) {
        var prefix = d3.formatPrefix(value, precision);
        value = prefix.scale(value);
        suffix = prefix.symbol;
      } else {
        value *= scale;
      }
      value = type(value, precision);
      var i = value.lastIndexOf("."), before = i < 0 ? value : value.substring(0, i), after = i < 0 ? "" : d3_format_decimalPoint + value.substring(i + 1);
      if (!zfill && comma) before = d3_format_group(before);
      var length = symbol.length + before.length + after.length + (zcomma ? 0 : negative.length), padding = length < width ? new Array(length = width - length + 1).join(fill) : "";
      if (zcomma) before = d3_format_group(padding + before);
      negative += symbol;
      value = before + after;
      return (align === "<" ? negative + value + padding : align === ">" ? padding + negative + value : align === "^" ? padding.substring(0, length >>= 1) + negative + value + padding.substring(length) : negative + (zcomma ? value : padding + value)) + suffix;
    };
  };
  var d3_format_re = /(?:([^{])?([<>=^]))?([+\- ])?([$#])?(0)?(\d+)?(,)?(\.-?\d+)?([a-z%])?/i;
  var d3_format_types = d3.map({
    b: function(x) {
      return x.toString(2);
    },
    c: function(x) {
      return String.fromCharCode(x);
    },
    o: function(x) {
      return x.toString(8);
    },
    x: function(x) {
      return x.toString(16);
    },
    X: function(x) {
      return x.toString(16).toUpperCase();
    },
    g: function(x, p) {
      return x.toPrecision(p);
    },
    e: function(x, p) {
      return x.toExponential(p);
    },
    f: function(x, p) {
      return x.toFixed(p);
    },
    r: function(x, p) {
      return (x = d3.round(x, d3_format_precision(x, p))).toFixed(Math.max(0, Math.min(20, d3_format_precision(x * (1 + 1e-15), p))));
    }
  });
  function d3_format_precision(x, p) {
    return p - (x ? Math.ceil(Math.log(x) / Math.LN10) : 1);
  }
  function d3_format_typeDefault(x) {
    return x + "";
  }
  var d3_format_group = d3_identity;
  if (d3_format_grouping) {
    var d3_format_groupingLength = d3_format_grouping.length;
    d3_format_group = function(value) {
      var i = value.length, t = [], j = 0, g = d3_format_grouping[0];
      while (i > 0 && g > 0) {
        t.push(value.substring(i -= g, i + g));
        g = d3_format_grouping[j = (j + 1) % d3_format_groupingLength];
      }
      return t.reverse().join(d3_format_thousandsSeparator);
    };
  }
  d3.geo = {};
  function d3_adder() {}
  d3_adder.prototype = {
    s: 0,
    t: 0,
    add: function(y) {
      d3_adderSum(y, this.t, d3_adderTemp);
      d3_adderSum(d3_adderTemp.s, this.s, this);
      if (this.s) this.t += d3_adderTemp.t; else this.s = d3_adderTemp.t;
    },
    reset: function() {
      this.s = this.t = 0;
    },
    valueOf: function() {
      return this.s;
    }
  };
  var d3_adderTemp = new d3_adder();
  function d3_adderSum(a, b, o) {
    var x = o.s = a + b, bv = x - a, av = x - bv;
    o.t = a - av + (b - bv);
  }
  d3.geo.stream = function(object, listener) {
    if (object && d3_geo_streamObjectType.hasOwnProperty(object.type)) {
      d3_geo_streamObjectType[object.type](object, listener);
    } else {
      d3_geo_streamGeometry(object, listener);
    }
  };
  function d3_geo_streamGeometry(geometry, listener) {
    if (geometry && d3_geo_streamGeometryType.hasOwnProperty(geometry.type)) {
      d3_geo_streamGeometryType[geometry.type](geometry, listener);
    }
  }
  var d3_geo_streamObjectType = {
    Feature: function(feature, listener) {
      d3_geo_streamGeometry(feature.geometry, listener);
    },
    FeatureCollection: function(object, listener) {
      var features = object.features, i = -1, n = features.length;
      while (++i < n) d3_geo_streamGeometry(features[i].geometry, listener);
    }
  };
  var d3_geo_streamGeometryType = {
    Sphere: function(object, listener) {
      listener.sphere();
    },
    Point: function(object, listener) {
      object = object.coordinates;
      listener.point(object[0], object[1], object[2]);
    },
    MultiPoint: function(object, listener) {
      var coordinates = object.coordinates, i = -1, n = coordinates.length;
      while (++i < n) object = coordinates[i], listener.point(object[0], object[1], object[2]);
    },
    LineString: function(object, listener) {
      d3_geo_streamLine(object.coordinates, listener, 0);
    },
    MultiLineString: function(object, listener) {
      var coordinates = object.coordinates, i = -1, n = coordinates.length;
      while (++i < n) d3_geo_streamLine(coordinates[i], listener, 0);
    },
    Polygon: function(object, listener) {
      d3_geo_streamPolygon(object.coordinates, listener);
    },
    MultiPolygon: function(object, listener) {
      var coordinates = object.coordinates, i = -1, n = coordinates.length;
      while (++i < n) d3_geo_streamPolygon(coordinates[i], listener);
    },
    GeometryCollection: function(object, listener) {
      var geometries = object.geometries, i = -1, n = geometries.length;
      while (++i < n) d3_geo_streamGeometry(geometries[i], listener);
    }
  };
  function d3_geo_streamLine(coordinates, listener, closed) {
    var i = -1, n = coordinates.length - closed, coordinate;
    listener.lineStart();
    while (++i < n) coordinate = coordinates[i], listener.point(coordinate[0], coordinate[1], coordinate[2]);
    listener.lineEnd();
  }
  function d3_geo_streamPolygon(coordinates, listener) {
    var i = -1, n = coordinates.length;
    listener.polygonStart();
    while (++i < n) d3_geo_streamLine(coordinates[i], listener, 1);
    listener.polygonEnd();
  }
  d3.geo.area = function(object) {
    d3_geo_areaSum = 0;
    d3.geo.stream(object, d3_geo_area);
    return d3_geo_areaSum;
  };
  var d3_geo_areaSum, d3_geo_areaRingSum = new d3_adder();
  var d3_geo_area = {
    sphere: function() {
      d3_geo_areaSum += 4 * π;
    },
    point: d3_noop,
    lineStart: d3_noop,
    lineEnd: d3_noop,
    polygonStart: function() {
      d3_geo_areaRingSum.reset();
      d3_geo_area.lineStart = d3_geo_areaRingStart;
    },
    polygonEnd: function() {
      var area = 2 * d3_geo_areaRingSum;
      d3_geo_areaSum += area < 0 ? 4 * π + area : area;
      d3_geo_area.lineStart = d3_geo_area.lineEnd = d3_geo_area.point = d3_noop;
    }
  };
  function d3_geo_areaRingStart() {
    var λ00, φ00, λ0, cosφ0, sinφ0;
    d3_geo_area.point = function(λ, φ) {
      d3_geo_area.point = nextPoint;
      λ0 = (λ00 = λ) * d3_radians, cosφ0 = Math.cos(φ = (φ00 = φ) * d3_radians / 2 + π / 4), 
      sinφ0 = Math.sin(φ);
    };
    function nextPoint(λ, φ) {
      λ *= d3_radians;
      φ = φ * d3_radians / 2 + π / 4;
      var dλ = λ - λ0, cosφ = Math.cos(φ), sinφ = Math.sin(φ), k = sinφ0 * sinφ, u = cosφ0 * cosφ + k * Math.cos(dλ), v = k * Math.sin(dλ);
      d3_geo_areaRingSum.add(Math.atan2(v, u));
      λ0 = λ, cosφ0 = cosφ, sinφ0 = sinφ;
    }
    d3_geo_area.lineEnd = function() {
      nextPoint(λ00, φ00);
    };
  }
  function d3_geo_cartesian(spherical) {
    var λ = spherical[0], φ = spherical[1], cosφ = Math.cos(φ);
    return [ cosφ * Math.cos(λ), cosφ * Math.sin(λ), Math.sin(φ) ];
  }
  function d3_geo_cartesianDot(a, b) {
    return a[0] * b[0] + a[1] * b[1] + a[2] * b[2];
  }
  function d3_geo_cartesianCross(a, b) {
    return [ a[1] * b[2] - a[2] * b[1], a[2] * b[0] - a[0] * b[2], a[0] * b[1] - a[1] * b[0] ];
  }
  function d3_geo_cartesianAdd(a, b) {
    a[0] += b[0];
    a[1] += b[1];
    a[2] += b[2];
  }
  function d3_geo_cartesianScale(vector, k) {
    return [ vector[0] * k, vector[1] * k, vector[2] * k ];
  }
  function d3_geo_cartesianNormalize(d) {
    var l = Math.sqrt(d[0] * d[0] + d[1] * d[1] + d[2] * d[2]);
    d[0] /= l;
    d[1] /= l;
    d[2] /= l;
  }
  function d3_geo_spherical(cartesian) {
    return [ Math.atan2(cartesian[1], cartesian[0]), d3_asin(cartesian[2]) ];
  }
  function d3_geo_sphericalEqual(a, b) {
    return abs(a[0] - b[0]) < ε && abs(a[1] - b[1]) < ε;
  }
  d3.geo.bounds = function() {
    var λ0, φ0, λ1, φ1, λ_, λ__, φ__, p0, dλSum, ranges, range;
    var bound = {
      point: point,
      lineStart: lineStart,
      lineEnd: lineEnd,
      polygonStart: function() {
        bound.point = ringPoint;
        bound.lineStart = ringStart;
        bound.lineEnd = ringEnd;
        dλSum = 0;
        d3_geo_area.polygonStart();
      },
      polygonEnd: function() {
        d3_geo_area.polygonEnd();
        bound.point = point;
        bound.lineStart = lineStart;
        bound.lineEnd = lineEnd;
        if (d3_geo_areaRingSum < 0) λ0 = -(λ1 = 180), φ0 = -(φ1 = 90); else if (dλSum > ε) φ1 = 90; else if (dλSum < -ε) φ0 = -90;
        range[0] = λ0, range[1] = λ1;
      }
    };
    function point(λ, φ) {
      ranges.push(range = [ λ0 = λ, λ1 = λ ]);
      if (φ < φ0) φ0 = φ;
      if (φ > φ1) φ1 = φ;
    }
    function linePoint(λ, φ) {
      var p = d3_geo_cartesian([ λ * d3_radians, φ * d3_radians ]);
      if (p0) {
        var normal = d3_geo_cartesianCross(p0, p), equatorial = [ normal[1], -normal[0], 0 ], inflection = d3_geo_cartesianCross(equatorial, normal);
        d3_geo_cartesianNormalize(inflection);
        inflection = d3_geo_spherical(inflection);
        var dλ = λ - λ_, s = dλ > 0 ? 1 : -1, λi = inflection[0] * d3_degrees * s, antimeridian = abs(dλ) > 180;
        if (antimeridian ^ (s * λ_ < λi && λi < s * λ)) {
          var φi = inflection[1] * d3_degrees;
          if (φi > φ1) φ1 = φi;
        } else if (λi = (λi + 360) % 360 - 180, antimeridian ^ (s * λ_ < λi && λi < s * λ)) {
          var φi = -inflection[1] * d3_degrees;
          if (φi < φ0) φ0 = φi;
        } else {
          if (φ < φ0) φ0 = φ;
          if (φ > φ1) φ1 = φ;
        }
        if (antimeridian) {
          if (λ < λ_) {
            if (angle(λ0, λ) > angle(λ0, λ1)) λ1 = λ;
          } else {
            if (angle(λ, λ1) > angle(λ0, λ1)) λ0 = λ;
          }
        } else {
          if (λ1 >= λ0) {
            if (λ < λ0) λ0 = λ;
            if (λ > λ1) λ1 = λ;
          } else {
            if (λ > λ_) {
              if (angle(λ0, λ) > angle(λ0, λ1)) λ1 = λ;
            } else {
              if (angle(λ, λ1) > angle(λ0, λ1)) λ0 = λ;
            }
          }
        }
      } else {
        point(λ, φ);
      }
      p0 = p, λ_ = λ;
    }
    function lineStart() {
      bound.point = linePoint;
    }
    function lineEnd() {
      range[0] = λ0, range[1] = λ1;
      bound.point = point;
      p0 = null;
    }
    function ringPoint(λ, φ) {
      if (p0) {
        var dλ = λ - λ_;
        dλSum += abs(dλ) > 180 ? dλ + (dλ > 0 ? 360 : -360) : dλ;
      } else λ__ = λ, φ__ = φ;
      d3_geo_area.point(λ, φ);
      linePoint(λ, φ);
    }
    function ringStart() {
      d3_geo_area.lineStart();
    }
    function ringEnd() {
      ringPoint(λ__, φ__);
      d3_geo_area.lineEnd();
      if (abs(dλSum) > ε) λ0 = -(λ1 = 180);
      range[0] = λ0, range[1] = λ1;
      p0 = null;
    }
    function angle(λ0, λ1) {
      return (λ1 -= λ0) < 0 ? λ1 + 360 : λ1;
    }
    function compareRanges(a, b) {
      return a[0] - b[0];
    }
    function withinRange(x, range) {
      return range[0] <= range[1] ? range[0] <= x && x <= range[1] : x < range[0] || range[1] < x;
    }
    return function(feature) {
      φ1 = λ1 = -(λ0 = φ0 = Infinity);
      ranges = [];
      d3.geo.stream(feature, bound);
      var n = ranges.length;
      if (n) {
        ranges.sort(compareRanges);
        for (var i = 1, a = ranges[0], b, merged = [ a ]; i < n; ++i) {
          b = ranges[i];
          if (withinRange(b[0], a) || withinRange(b[1], a)) {
            if (angle(a[0], b[1]) > angle(a[0], a[1])) a[1] = b[1];
            if (angle(b[0], a[1]) > angle(a[0], a[1])) a[0] = b[0];
          } else {
            merged.push(a = b);
          }
        }
        var best = -Infinity, dλ;
        for (var n = merged.length - 1, i = 0, a = merged[n], b; i <= n; a = b, ++i) {
          b = merged[i];
          if ((dλ = angle(a[1], b[0])) > best) best = dλ, λ0 = b[0], λ1 = a[1];
        }
      }
      ranges = range = null;
      return λ0 === Infinity || φ0 === Infinity ? [ [ NaN, NaN ], [ NaN, NaN ] ] : [ [ λ0, φ0 ], [ λ1, φ1 ] ];
    };
  }();
  d3.geo.centroid = function(object) {
    d3_geo_centroidW0 = d3_geo_centroidW1 = d3_geo_centroidX0 = d3_geo_centroidY0 = d3_geo_centroidZ0 = d3_geo_centroidX1 = d3_geo_centroidY1 = d3_geo_centroidZ1 = d3_geo_centroidX2 = d3_geo_centroidY2 = d3_geo_centroidZ2 = 0;
    d3.geo.stream(object, d3_geo_centroid);
    var x = d3_geo_centroidX2, y = d3_geo_centroidY2, z = d3_geo_centroidZ2, m = x * x + y * y + z * z;
    if (m < ε2) {
      x = d3_geo_centroidX1, y = d3_geo_centroidY1, z = d3_geo_centroidZ1;
      if (d3_geo_centroidW1 < ε) x = d3_geo_centroidX0, y = d3_geo_centroidY0, z = d3_geo_centroidZ0;
      m = x * x + y * y + z * z;
      if (m < ε2) return [ NaN, NaN ];
    }
    return [ Math.atan2(y, x) * d3_degrees, d3_asin(z / Math.sqrt(m)) * d3_degrees ];
  };
  var d3_geo_centroidW0, d3_geo_centroidW1, d3_geo_centroidX0, d3_geo_centroidY0, d3_geo_centroidZ0, d3_geo_centroidX1, d3_geo_centroidY1, d3_geo_centroidZ1, d3_geo_centroidX2, d3_geo_centroidY2, d3_geo_centroidZ2;
  var d3_geo_centroid = {
    sphere: d3_noop,
    point: d3_geo_centroidPoint,
    lineStart: d3_geo_centroidLineStart,
    lineEnd: d3_geo_centroidLineEnd,
    polygonStart: function() {
      d3_geo_centroid.lineStart = d3_geo_centroidRingStart;
    },
    polygonEnd: function() {
      d3_geo_centroid.lineStart = d3_geo_centroidLineStart;
    }
  };
  function d3_geo_centroidPoint(λ, φ) {
    λ *= d3_radians;
    var cosφ = Math.cos(φ *= d3_radians);
    d3_geo_centroidPointXYZ(cosφ * Math.cos(λ), cosφ * Math.sin(λ), Math.sin(φ));
  }
  function d3_geo_centroidPointXYZ(x, y, z) {
    ++d3_geo_centroidW0;
    d3_geo_centroidX0 += (x - d3_geo_centroidX0) / d3_geo_centroidW0;
    d3_geo_centroidY0 += (y - d3_geo_centroidY0) / d3_geo_centroidW0;
    d3_geo_centroidZ0 += (z - d3_geo_centroidZ0) / d3_geo_centroidW0;
  }
  function d3_geo_centroidLineStart() {
    var x0, y0, z0;
    d3_geo_centroid.point = function(λ, φ) {
      λ *= d3_radians;
      var cosφ = Math.cos(φ *= d3_radians);
      x0 = cosφ * Math.cos(λ);
      y0 = cosφ * Math.sin(λ);
      z0 = Math.sin(φ);
      d3_geo_centroid.point = nextPoint;
      d3_geo_centroidPointXYZ(x0, y0, z0);
    };
    function nextPoint(λ, φ) {
      λ *= d3_radians;
      var cosφ = Math.cos(φ *= d3_radians), x = cosφ * Math.cos(λ), y = cosφ * Math.sin(λ), z = Math.sin(φ), w = Math.atan2(Math.sqrt((w = y0 * z - z0 * y) * w + (w = z0 * x - x0 * z) * w + (w = x0 * y - y0 * x) * w), x0 * x + y0 * y + z0 * z);
      d3_geo_centroidW1 += w;
      d3_geo_centroidX1 += w * (x0 + (x0 = x));
      d3_geo_centroidY1 += w * (y0 + (y0 = y));
      d3_geo_centroidZ1 += w * (z0 + (z0 = z));
      d3_geo_centroidPointXYZ(x0, y0, z0);
    }
  }
  function d3_geo_centroidLineEnd() {
    d3_geo_centroid.point = d3_geo_centroidPoint;
  }
  function d3_geo_centroidRingStart() {
    var λ00, φ00, x0, y0, z0;
    d3_geo_centroid.point = function(λ, φ) {
      λ00 = λ, φ00 = φ;
      d3_geo_centroid.point = nextPoint;
      λ *= d3_radians;
      var cosφ = Math.cos(φ *= d3_radians);
      x0 = cosφ * Math.cos(λ);
      y0 = cosφ * Math.sin(λ);
      z0 = Math.sin(φ);
      d3_geo_centroidPointXYZ(x0, y0, z0);
    };
    d3_geo_centroid.lineEnd = function() {
      nextPoint(λ00, φ00);
      d3_geo_centroid.lineEnd = d3_geo_centroidLineEnd;
      d3_geo_centroid.point = d3_geo_centroidPoint;
    };
    function nextPoint(λ, φ) {
      λ *= d3_radians;
      var cosφ = Math.cos(φ *= d3_radians), x = cosφ * Math.cos(λ), y = cosφ * Math.sin(λ), z = Math.sin(φ), cx = y0 * z - z0 * y, cy = z0 * x - x0 * z, cz = x0 * y - y0 * x, m = Math.sqrt(cx * cx + cy * cy + cz * cz), u = x0 * x + y0 * y + z0 * z, v = m && -d3_acos(u) / m, w = Math.atan2(m, u);
      d3_geo_centroidX2 += v * cx;
      d3_geo_centroidY2 += v * cy;
      d3_geo_centroidZ2 += v * cz;
      d3_geo_centroidW1 += w;
      d3_geo_centroidX1 += w * (x0 + (x0 = x));
      d3_geo_centroidY1 += w * (y0 + (y0 = y));
      d3_geo_centroidZ1 += w * (z0 + (z0 = z));
      d3_geo_centroidPointXYZ(x0, y0, z0);
    }
  }
  function d3_true() {
    return true;
  }
  function d3_geo_clipPolygon(segments, compare, clipStartInside, interpolate, listener) {
    var subject = [], clip = [];
    segments.forEach(function(segment) {
      if ((n = segment.length - 1) <= 0) return;
      var n, p0 = segment[0], p1 = segment[n];
      if (d3_geo_sphericalEqual(p0, p1)) {
        listener.lineStart();
        for (var i = 0; i < n; ++i) listener.point((p0 = segment[i])[0], p0[1]);
        listener.lineEnd();
        return;
      }
      var a = new d3_geo_clipPolygonIntersection(p0, segment, null, true), b = new d3_geo_clipPolygonIntersection(p0, null, a, false);
      a.o = b;
      subject.push(a);
      clip.push(b);
      a = new d3_geo_clipPolygonIntersection(p1, segment, null, false);
      b = new d3_geo_clipPolygonIntersection(p1, null, a, true);
      a.o = b;
      subject.push(a);
      clip.push(b);
    });
    clip.sort(compare);
    d3_geo_clipPolygonLinkCircular(subject);
    d3_geo_clipPolygonLinkCircular(clip);
    if (!subject.length) return;
    for (var i = 0, entry = clipStartInside, n = clip.length; i < n; ++i) {
      clip[i].e = entry = !entry;
    }
    var start = subject[0], points, point;
    while (1) {
      var current = start, isSubject = true;
      while (current.v) if ((current = current.n) === start) return;
      points = current.z;
      listener.lineStart();
      do {
        current.v = current.o.v = true;
        if (current.e) {
          if (isSubject) {
            for (var i = 0, n = points.length; i < n; ++i) listener.point((point = points[i])[0], point[1]);
          } else {
            interpolate(current.x, current.n.x, 1, listener);
          }
          current = current.n;
        } else {
          if (isSubject) {
            points = current.p.z;
            for (var i = points.length - 1; i >= 0; --i) listener.point((point = points[i])[0], point[1]);
          } else {
            interpolate(current.x, current.p.x, -1, listener);
          }
          current = current.p;
        }
        current = current.o;
        points = current.z;
        isSubject = !isSubject;
      } while (!current.v);
      listener.lineEnd();
    }
  }
  function d3_geo_clipPolygonLinkCircular(array) {
    if (!(n = array.length)) return;
    var n, i = 0, a = array[0], b;
    while (++i < n) {
      a.n = b = array[i];
      b.p = a;
      a = b;
    }
    a.n = b = array[0];
    b.p = a;
  }
  function d3_geo_clipPolygonIntersection(point, points, other, entry) {
    this.x = point;
    this.z = points;
    this.o = other;
    this.e = entry;
    this.v = false;
    this.n = this.p = null;
  }
  function d3_geo_clip(pointVisible, clipLine, interpolate, clipStart) {
    return function(rotate, listener) {
      var line = clipLine(listener), rotatedClipStart = rotate.invert(clipStart[0], clipStart[1]);
      var clip = {
        point: point,
        lineStart: lineStart,
        lineEnd: lineEnd,
        polygonStart: function() {
          clip.point = pointRing;
          clip.lineStart = ringStart;
          clip.lineEnd = ringEnd;
          segments = [];
          polygon = [];
          listener.polygonStart();
        },
        polygonEnd: function() {
          clip.point = point;
          clip.lineStart = lineStart;
          clip.lineEnd = lineEnd;
          segments = d3.merge(segments);
          var clipStartInside = d3_geo_pointInPolygon(rotatedClipStart, polygon);
          if (segments.length) {
            d3_geo_clipPolygon(segments, d3_geo_clipSort, clipStartInside, interpolate, listener);
          } else if (clipStartInside) {
            listener.lineStart();
            interpolate(null, null, 1, listener);
            listener.lineEnd();
          }
          listener.polygonEnd();
          segments = polygon = null;
        },
        sphere: function() {
          listener.polygonStart();
          listener.lineStart();
          interpolate(null, null, 1, listener);
          listener.lineEnd();
          listener.polygonEnd();
        }
      };
      function point(λ, φ) {
        var point = rotate(λ, φ);
        if (pointVisible(λ = point[0], φ = point[1])) listener.point(λ, φ);
      }
      function pointLine(λ, φ) {
        var point = rotate(λ, φ);
        line.point(point[0], point[1]);
      }
      function lineStart() {
        clip.point = pointLine;
        line.lineStart();
      }
      function lineEnd() {
        clip.point = point;
        line.lineEnd();
      }
      var segments;
      var buffer = d3_geo_clipBufferListener(), ringListener = clipLine(buffer), polygon, ring;
      function pointRing(λ, φ) {
        ring.push([ λ, φ ]);
        var point = rotate(λ, φ);
        ringListener.point(point[0], point[1]);
      }
      function ringStart() {
        ringListener.lineStart();
        ring = [];
      }
      function ringEnd() {
        pointRing(ring[0][0], ring[0][1]);
        ringListener.lineEnd();
        var clean = ringListener.clean(), ringSegments = buffer.buffer(), segment, n = ringSegments.length;
        ring.pop();
        polygon.push(ring);
        ring = null;
        if (!n) return;
        if (clean & 1) {
          segment = ringSegments[0];
          var n = segment.length - 1, i = -1, point;
          listener.lineStart();
          while (++i < n) listener.point((point = segment[i])[0], point[1]);
          listener.lineEnd();
          return;
        }
        if (n > 1 && clean & 2) ringSegments.push(ringSegments.pop().concat(ringSegments.shift()));
        segments.push(ringSegments.filter(d3_geo_clipSegmentLength1));
      }
      return clip;
    };
  }
  function d3_geo_clipSegmentLength1(segment) {
    return segment.length > 1;
  }
  function d3_geo_clipBufferListener() {
    var lines = [], line;
    return {
      lineStart: function() {
        lines.push(line = []);
      },
      point: function(λ, φ) {
        line.push([ λ, φ ]);
      },
      lineEnd: d3_noop,
      buffer: function() {
        var buffer = lines;
        lines = [];
        line = null;
        return buffer;
      },
      rejoin: function() {
        if (lines.length > 1) lines.push(lines.pop().concat(lines.shift()));
      }
    };
  }
  function d3_geo_clipSort(a, b) {
    return ((a = a.x)[0] < 0 ? a[1] - halfπ - ε : halfπ - a[1]) - ((b = b.x)[0] < 0 ? b[1] - halfπ - ε : halfπ - b[1]);
  }
  function d3_geo_pointInPolygon(point, polygon) {
    var meridian = point[0], parallel = point[1], meridianNormal = [ Math.sin(meridian), -Math.cos(meridian), 0 ], polarAngle = 0, winding = 0;
    d3_geo_areaRingSum.reset();
    for (var i = 0, n = polygon.length; i < n; ++i) {
      var ring = polygon[i], m = ring.length;
      if (!m) continue;
      var point0 = ring[0], λ0 = point0[0], φ0 = point0[1] / 2 + π / 4, sinφ0 = Math.sin(φ0), cosφ0 = Math.cos(φ0), j = 1;
      while (true) {
        if (j === m) j = 0;
        point = ring[j];
        var λ = point[0], φ = point[1] / 2 + π / 4, sinφ = Math.sin(φ), cosφ = Math.cos(φ), dλ = λ - λ0, antimeridian = abs(dλ) > π, k = sinφ0 * sinφ;
        d3_geo_areaRingSum.add(Math.atan2(k * Math.sin(dλ), cosφ0 * cosφ + k * Math.cos(dλ)));
        polarAngle += antimeridian ? dλ + (dλ >= 0 ? τ : -τ) : dλ;
        if (antimeridian ^ λ0 >= meridian ^ λ >= meridian) {
          var arc = d3_geo_cartesianCross(d3_geo_cartesian(point0), d3_geo_cartesian(point));
          d3_geo_cartesianNormalize(arc);
          var intersection = d3_geo_cartesianCross(meridianNormal, arc);
          d3_geo_cartesianNormalize(intersection);
          var φarc = (antimeridian ^ dλ >= 0 ? -1 : 1) * d3_asin(intersection[2]);
          if (parallel > φarc || parallel === φarc && (arc[0] || arc[1])) {
            winding += antimeridian ^ dλ >= 0 ? 1 : -1;
          }
        }
        if (!j++) break;
        λ0 = λ, sinφ0 = sinφ, cosφ0 = cosφ, point0 = point;
      }
    }
    return (polarAngle < -ε || polarAngle < ε && d3_geo_areaRingSum < 0) ^ winding & 1;
  }
  var d3_geo_clipAntimeridian = d3_geo_clip(d3_true, d3_geo_clipAntimeridianLine, d3_geo_clipAntimeridianInterpolate, [ -π, -π / 2 ]);
  function d3_geo_clipAntimeridianLine(listener) {
    var λ0 = NaN, φ0 = NaN, sλ0 = NaN, clean;
    return {
      lineStart: function() {
        listener.lineStart();
        clean = 1;
      },
      point: function(λ1, φ1) {
        var sλ1 = λ1 > 0 ? π : -π, dλ = abs(λ1 - λ0);
        if (abs(dλ - π) < ε) {
          listener.point(λ0, φ0 = (φ0 + φ1) / 2 > 0 ? halfπ : -halfπ);
          listener.point(sλ0, φ0);
          listener.lineEnd();
          listener.lineStart();
          listener.point(sλ1, φ0);
          listener.point(λ1, φ0);
          clean = 0;
        } else if (sλ0 !== sλ1 && dλ >= π) {
          if (abs(λ0 - sλ0) < ε) λ0 -= sλ0 * ε;
          if (abs(λ1 - sλ1) < ε) λ1 -= sλ1 * ε;
          φ0 = d3_geo_clipAntimeridianIntersect(λ0, φ0, λ1, φ1);
          listener.point(sλ0, φ0);
          listener.lineEnd();
          listener.lineStart();
          listener.point(sλ1, φ0);
          clean = 0;
        }
        listener.point(λ0 = λ1, φ0 = φ1);
        sλ0 = sλ1;
      },
      lineEnd: function() {
        listener.lineEnd();
        λ0 = φ0 = NaN;
      },
      clean: function() {
        return 2 - clean;
      }
    };
  }
  function d3_geo_clipAntimeridianIntersect(λ0, φ0, λ1, φ1) {
    var cosφ0, cosφ1, sinλ0_λ1 = Math.sin(λ0 - λ1);
    return abs(sinλ0_λ1) > ε ? Math.atan((Math.sin(φ0) * (cosφ1 = Math.cos(φ1)) * Math.sin(λ1) - Math.sin(φ1) * (cosφ0 = Math.cos(φ0)) * Math.sin(λ0)) / (cosφ0 * cosφ1 * sinλ0_λ1)) : (φ0 + φ1) / 2;
  }
  function d3_geo_clipAntimeridianInterpolate(from, to, direction, listener) {
    var φ;
    if (from == null) {
      φ = direction * halfπ;
      listener.point(-π, φ);
      listener.point(0, φ);
      listener.point(π, φ);
      listener.point(π, 0);
      listener.point(π, -φ);
      listener.point(0, -φ);
      listener.point(-π, -φ);
      listener.point(-π, 0);
      listener.point(-π, φ);
    } else if (abs(from[0] - to[0]) > ε) {
      var s = from[0] < to[0] ? π : -π;
      φ = direction * s / 2;
      listener.point(-s, φ);
      listener.point(0, φ);
      listener.point(s, φ);
    } else {
      listener.point(to[0], to[1]);
    }
  }
  function d3_geo_clipCircle(radius) {
    var cr = Math.cos(radius), smallRadius = cr > 0, notHemisphere = abs(cr) > ε, interpolate = d3_geo_circleInterpolate(radius, 6 * d3_radians);
    return d3_geo_clip(visible, clipLine, interpolate, smallRadius ? [ 0, -radius ] : [ -π, radius - π ]);
    function visible(λ, φ) {
      return Math.cos(λ) * Math.cos(φ) > cr;
    }
    function clipLine(listener) {
      var point0, c0, v0, v00, clean;
      return {
        lineStart: function() {
          v00 = v0 = false;
          clean = 1;
        },
        point: function(λ, φ) {
          var point1 = [ λ, φ ], point2, v = visible(λ, φ), c = smallRadius ? v ? 0 : code(λ, φ) : v ? code(λ + (λ < 0 ? π : -π), φ) : 0;
          if (!point0 && (v00 = v0 = v)) listener.lineStart();
          if (v !== v0) {
            point2 = intersect(point0, point1);
            if (d3_geo_sphericalEqual(point0, point2) || d3_geo_sphericalEqual(point1, point2)) {
              point1[0] += ε;
              point1[1] += ε;
              v = visible(point1[0], point1[1]);
            }
          }
          if (v !== v0) {
            clean = 0;
            if (v) {
              listener.lineStart();
              point2 = intersect(point1, point0);
              listener.point(point2[0], point2[1]);
            } else {
              point2 = intersect(point0, point1);
              listener.point(point2[0], point2[1]);
              listener.lineEnd();
            }
            point0 = point2;
          } else if (notHemisphere && point0 && smallRadius ^ v) {
            var t;
            if (!(c & c0) && (t = intersect(point1, point0, true))) {
              clean = 0;
              if (smallRadius) {
                listener.lineStart();
                listener.point(t[0][0], t[0][1]);
                listener.point(t[1][0], t[1][1]);
                listener.lineEnd();
              } else {
                listener.point(t[1][0], t[1][1]);
                listener.lineEnd();
                listener.lineStart();
                listener.point(t[0][0], t[0][1]);
              }
            }
          }
          if (v && (!point0 || !d3_geo_sphericalEqual(point0, point1))) {
            listener.point(point1[0], point1[1]);
          }
          point0 = point1, v0 = v, c0 = c;
        },
        lineEnd: function() {
          if (v0) listener.lineEnd();
          point0 = null;
        },
        clean: function() {
          return clean | (v00 && v0) << 1;
        }
      };
    }
    function intersect(a, b, two) {
      var pa = d3_geo_cartesian(a), pb = d3_geo_cartesian(b);
      var n1 = [ 1, 0, 0 ], n2 = d3_geo_cartesianCross(pa, pb), n2n2 = d3_geo_cartesianDot(n2, n2), n1n2 = n2[0], determinant = n2n2 - n1n2 * n1n2;
      if (!determinant) return !two && a;
      var c1 = cr * n2n2 / determinant, c2 = -cr * n1n2 / determinant, n1xn2 = d3_geo_cartesianCross(n1, n2), A = d3_geo_cartesianScale(n1, c1), B = d3_geo_cartesianScale(n2, c2);
      d3_geo_cartesianAdd(A, B);
      var u = n1xn2, w = d3_geo_cartesianDot(A, u), uu = d3_geo_cartesianDot(u, u), t2 = w * w - uu * (d3_geo_cartesianDot(A, A) - 1);
      if (t2 < 0) return;
      var t = Math.sqrt(t2), q = d3_geo_cartesianScale(u, (-w - t) / uu);
      d3_geo_cartesianAdd(q, A);
      q = d3_geo_spherical(q);
      if (!two) return q;
      var λ0 = a[0], λ1 = b[0], φ0 = a[1], φ1 = b[1], z;
      if (λ1 < λ0) z = λ0, λ0 = λ1, λ1 = z;
      var δλ = λ1 - λ0, polar = abs(δλ - π) < ε, meridian = polar || δλ < ε;
      if (!polar && φ1 < φ0) z = φ0, φ0 = φ1, φ1 = z;
      if (meridian ? polar ? φ0 + φ1 > 0 ^ q[1] < (abs(q[0] - λ0) < ε ? φ0 : φ1) : φ0 <= q[1] && q[1] <= φ1 : δλ > π ^ (λ0 <= q[0] && q[0] <= λ1)) {
        var q1 = d3_geo_cartesianScale(u, (-w + t) / uu);
        d3_geo_cartesianAdd(q1, A);
        return [ q, d3_geo_spherical(q1) ];
      }
    }
    function code(λ, φ) {
      var r = smallRadius ? radius : π - radius, code = 0;
      if (λ < -r) code |= 1; else if (λ > r) code |= 2;
      if (φ < -r) code |= 4; else if (φ > r) code |= 8;
      return code;
    }
  }
  function d3_geom_clipLine(x0, y0, x1, y1) {
    return function(line) {
      var a = line.a, b = line.b, ax = a.x, ay = a.y, bx = b.x, by = b.y, t0 = 0, t1 = 1, dx = bx - ax, dy = by - ay, r;
      r = x0 - ax;
      if (!dx && r > 0) return;
      r /= dx;
      if (dx < 0) {
        if (r < t0) return;
        if (r < t1) t1 = r;
      } else if (dx > 0) {
        if (r > t1) return;
        if (r > t0) t0 = r;
      }
      r = x1 - ax;
      if (!dx && r < 0) return;
      r /= dx;
      if (dx < 0) {
        if (r > t1) return;
        if (r > t0) t0 = r;
      } else if (dx > 0) {
        if (r < t0) return;
        if (r < t1) t1 = r;
      }
      r = y0 - ay;
      if (!dy && r > 0) return;
      r /= dy;
      if (dy < 0) {
        if (r < t0) return;
        if (r < t1) t1 = r;
      } else if (dy > 0) {
        if (r > t1) return;
        if (r > t0) t0 = r;
      }
      r = y1 - ay;
      if (!dy && r < 0) return;
      r /= dy;
      if (dy < 0) {
        if (r > t1) return;
        if (r > t0) t0 = r;
      } else if (dy > 0) {
        if (r < t0) return;
        if (r < t1) t1 = r;
      }
      if (t0 > 0) line.a = {
        x: ax + t0 * dx,
        y: ay + t0 * dy
      };
      if (t1 < 1) line.b = {
        x: ax + t1 * dx,
        y: ay + t1 * dy
      };
      return line;
    };
  }
  var d3_geo_clipExtentMAX = 1e9;
  d3.geo.clipExtent = function() {
    var x0, y0, x1, y1, stream, clip, clipExtent = {
      stream: function(output) {
        if (stream) stream.valid = false;
        stream = clip(output);
        stream.valid = true;
        return stream;
      },
      extent: function(_) {
        if (!arguments.length) return [ [ x0, y0 ], [ x1, y1 ] ];
        clip = d3_geo_clipExtent(x0 = +_[0][0], y0 = +_[0][1], x1 = +_[1][0], y1 = +_[1][1]);
        if (stream) stream.valid = false, stream = null;
        return clipExtent;
      }
    };
    return clipExtent.extent([ [ 0, 0 ], [ 960, 500 ] ]);
  };
  function d3_geo_clipExtent(x0, y0, x1, y1) {
    return function(listener) {
      var listener_ = listener, bufferListener = d3_geo_clipBufferListener(), clipLine = d3_geom_clipLine(x0, y0, x1, y1), segments, polygon, ring;
      var clip = {
        point: point,
        lineStart: lineStart,
        lineEnd: lineEnd,
        polygonStart: function() {
          listener = bufferListener;
          segments = [];
          polygon = [];
          clean = true;
        },
        polygonEnd: function() {
          listener = listener_;
          segments = d3.merge(segments);
          var clipStartInside = insidePolygon([ x0, y1 ]), inside = clean && clipStartInside, visible = segments.length;
          if (inside || visible) {
            listener.polygonStart();
            if (inside) {
              listener.lineStart();
              interpolate(null, null, 1, listener);
              listener.lineEnd();
            }
            if (visible) {
              d3_geo_clipPolygon(segments, compare, clipStartInside, interpolate, listener);
            }
            listener.polygonEnd();
          }
          segments = polygon = ring = null;
        }
      };
      function insidePolygon(p) {
        var wn = 0, n = polygon.length, y = p[1];
        for (var i = 0; i < n; ++i) {
          for (var j = 1, v = polygon[i], m = v.length, a = v[0], b; j < m; ++j) {
            b = v[j];
            if (a[1] <= y) {
              if (b[1] > y && isLeft(a, b, p) > 0) ++wn;
            } else {
              if (b[1] <= y && isLeft(a, b, p) < 0) --wn;
            }
            a = b;
          }
        }
        return wn !== 0;
      }
      function isLeft(a, b, c) {
        return (b[0] - a[0]) * (c[1] - a[1]) - (c[0] - a[0]) * (b[1] - a[1]);
      }
      function interpolate(from, to, direction, listener) {
        var a = 0, a1 = 0;
        if (from == null || (a = corner(from, direction)) !== (a1 = corner(to, direction)) || comparePoints(from, to) < 0 ^ direction > 0) {
          do {
            listener.point(a === 0 || a === 3 ? x0 : x1, a > 1 ? y1 : y0);
          } while ((a = (a + direction + 4) % 4) !== a1);
        } else {
          listener.point(to[0], to[1]);
        }
      }
      function pointVisible(x, y) {
        return x0 <= x && x <= x1 && y0 <= y && y <= y1;
      }
      function point(x, y) {
        if (pointVisible(x, y)) listener.point(x, y);
      }
      var x__, y__, v__, x_, y_, v_, first, clean;
      function lineStart() {
        clip.point = linePoint;
        if (polygon) polygon.push(ring = []);
        first = true;
        v_ = false;
        x_ = y_ = NaN;
      }
      function lineEnd() {
        if (segments) {
          linePoint(x__, y__);
          if (v__ && v_) bufferListener.rejoin();
          segments.push(bufferListener.buffer());
        }
        clip.point = point;
        if (v_) listener.lineEnd();
      }
      function linePoint(x, y) {
        x = Math.max(-d3_geo_clipExtentMAX, Math.min(d3_geo_clipExtentMAX, x));
        y = Math.max(-d3_geo_clipExtentMAX, Math.min(d3_geo_clipExtentMAX, y));
        var v = pointVisible(x, y);
        if (polygon) ring.push([ x, y ]);
        if (first) {
          x__ = x, y__ = y, v__ = v;
          first = false;
          if (v) {
            listener.lineStart();
            listener.point(x, y);
          }
        } else {
          if (v && v_) listener.point(x, y); else {
            var l = {
              a: {
                x: x_,
                y: y_
              },
              b: {
                x: x,
                y: y
              }
            };
            if (clipLine(l)) {
              if (!v_) {
                listener.lineStart();
                listener.point(l.a.x, l.a.y);
              }
              listener.point(l.b.x, l.b.y);
              if (!v) listener.lineEnd();
              clean = false;
            } else if (v) {
              listener.lineStart();
              listener.point(x, y);
              clean = false;
            }
          }
        }
        x_ = x, y_ = y, v_ = v;
      }
      return clip;
    };
    function corner(p, direction) {
      return abs(p[0] - x0) < ε ? direction > 0 ? 0 : 3 : abs(p[0] - x1) < ε ? direction > 0 ? 2 : 1 : abs(p[1] - y0) < ε ? direction > 0 ? 1 : 0 : direction > 0 ? 3 : 2;
    }
    function compare(a, b) {
      return comparePoints(a.x, b.x);
    }
    function comparePoints(a, b) {
      var ca = corner(a, 1), cb = corner(b, 1);
      return ca !== cb ? ca - cb : ca === 0 ? b[1] - a[1] : ca === 1 ? a[0] - b[0] : ca === 2 ? a[1] - b[1] : b[0] - a[0];
    }
  }
  function d3_geo_compose(a, b) {
    function compose(x, y) {
      return x = a(x, y), b(x[0], x[1]);
    }
    if (a.invert && b.invert) compose.invert = function(x, y) {
      return x = b.invert(x, y), x && a.invert(x[0], x[1]);
    };
    return compose;
  }
  function d3_geo_conic(projectAt) {
    var φ0 = 0, φ1 = π / 3, m = d3_geo_projectionMutator(projectAt), p = m(φ0, φ1);
    p.parallels = function(_) {
      if (!arguments.length) return [ φ0 / π * 180, φ1 / π * 180 ];
      return m(φ0 = _[0] * π / 180, φ1 = _[1] * π / 180);
    };
    return p;
  }
  function d3_geo_conicEqualArea(φ0, φ1) {
    var sinφ0 = Math.sin(φ0), n = (sinφ0 + Math.sin(φ1)) / 2, C = 1 + sinφ0 * (2 * n - sinφ0), ρ0 = Math.sqrt(C) / n;
    function forward(λ, φ) {
      var ρ = Math.sqrt(C - 2 * n * Math.sin(φ)) / n;
      return [ ρ * Math.sin(λ *= n), ρ0 - ρ * Math.cos(λ) ];
    }
    forward.invert = function(x, y) {
      var ρ0_y = ρ0 - y;
      return [ Math.atan2(x, ρ0_y) / n, d3_asin((C - (x * x + ρ0_y * ρ0_y) * n * n) / (2 * n)) ];
    };
    return forward;
  }
  (d3.geo.conicEqualArea = function() {
    return d3_geo_conic(d3_geo_conicEqualArea);
  }).raw = d3_geo_conicEqualArea;
  d3.geo.albers = function() {
    return d3.geo.conicEqualArea().rotate([ 96, 0 ]).center([ -.6, 38.7 ]).parallels([ 29.5, 45.5 ]).scale(1070);
  };
  d3.geo.albersUsa = function() {
    var lower48 = d3.geo.albers();
    var alaska = d3.geo.conicEqualArea().rotate([ 154, 0 ]).center([ -2, 58.5 ]).parallels([ 55, 65 ]);
    var hawaii = d3.geo.conicEqualArea().rotate([ 157, 0 ]).center([ -3, 19.9 ]).parallels([ 8, 18 ]);
    var point, pointStream = {
      point: function(x, y) {
        point = [ x, y ];
      }
    }, lower48Point, alaskaPoint, hawaiiPoint;
    function albersUsa(coordinates) {
      var x = coordinates[0], y = coordinates[1];
      point = null;
      (lower48Point(x, y), point) || (alaskaPoint(x, y), point) || hawaiiPoint(x, y);
      return point;
    }
    albersUsa.invert = function(coordinates) {
      var k = lower48.scale(), t = lower48.translate(), x = (coordinates[0] - t[0]) / k, y = (coordinates[1] - t[1]) / k;
      return (y >= .12 && y < .234 && x >= -.425 && x < -.214 ? alaska : y >= .166 && y < .234 && x >= -.214 && x < -.115 ? hawaii : lower48).invert(coordinates);
    };
    albersUsa.stream = function(stream) {
      var lower48Stream = lower48.stream(stream), alaskaStream = alaska.stream(stream), hawaiiStream = hawaii.stream(stream);
      return {
        point: function(x, y) {
          lower48Stream.point(x, y);
          alaskaStream.point(x, y);
          hawaiiStream.point(x, y);
        },
        sphere: function() {
          lower48Stream.sphere();
          alaskaStream.sphere();
          hawaiiStream.sphere();
        },
        lineStart: function() {
          lower48Stream.lineStart();
          alaskaStream.lineStart();
          hawaiiStream.lineStart();
        },
        lineEnd: function() {
          lower48Stream.lineEnd();
          alaskaStream.lineEnd();
          hawaiiStream.lineEnd();
        },
        polygonStart: function() {
          lower48Stream.polygonStart();
          alaskaStream.polygonStart();
          hawaiiStream.polygonStart();
        },
        polygonEnd: function() {
          lower48Stream.polygonEnd();
          alaskaStream.polygonEnd();
          hawaiiStream.polygonEnd();
        }
      };
    };
    albersUsa.precision = function(_) {
      if (!arguments.length) return lower48.precision();
      lower48.precision(_);
      alaska.precision(_);
      hawaii.precision(_);
      return albersUsa;
    };
    albersUsa.scale = function(_) {
      if (!arguments.length) return lower48.scale();
      lower48.scale(_);
      alaska.scale(_ * .35);
      hawaii.scale(_);
      return albersUsa.translate(lower48.translate());
    };
    albersUsa.translate = function(_) {
      if (!arguments.length) return lower48.translate();
      var k = lower48.scale(), x = +_[0], y = +_[1];
      lower48Point = lower48.translate(_).clipExtent([ [ x - .455 * k, y - .238 * k ], [ x + .455 * k, y + .238 * k ] ]).stream(pointStream).point;
      alaskaPoint = alaska.translate([ x - .307 * k, y + .201 * k ]).clipExtent([ [ x - .425 * k + ε, y + .12 * k + ε ], [ x - .214 * k - ε, y + .234 * k - ε ] ]).stream(pointStream).point;
      hawaiiPoint = hawaii.translate([ x - .205 * k, y + .212 * k ]).clipExtent([ [ x - .214 * k + ε, y + .166 * k + ε ], [ x - .115 * k - ε, y + .234 * k - ε ] ]).stream(pointStream).point;
      return albersUsa;
    };
    return albersUsa.scale(1070);
  };
  var d3_geo_pathAreaSum, d3_geo_pathAreaPolygon, d3_geo_pathArea = {
    point: d3_noop,
    lineStart: d3_noop,
    lineEnd: d3_noop,
    polygonStart: function() {
      d3_geo_pathAreaPolygon = 0;
      d3_geo_pathArea.lineStart = d3_geo_pathAreaRingStart;
    },
    polygonEnd: function() {
      d3_geo_pathArea.lineStart = d3_geo_pathArea.lineEnd = d3_geo_pathArea.point = d3_noop;
      d3_geo_pathAreaSum += abs(d3_geo_pathAreaPolygon / 2);
    }
  };
  function d3_geo_pathAreaRingStart() {
    var x00, y00, x0, y0;
    d3_geo_pathArea.point = function(x, y) {
      d3_geo_pathArea.point = nextPoint;
      x00 = x0 = x, y00 = y0 = y;
    };
    function nextPoint(x, y) {
      d3_geo_pathAreaPolygon += y0 * x - x0 * y;
      x0 = x, y0 = y;
    }
    d3_geo_pathArea.lineEnd = function() {
      nextPoint(x00, y00);
    };
  }
  var d3_geo_pathBoundsX0, d3_geo_pathBoundsY0, d3_geo_pathBoundsX1, d3_geo_pathBoundsY1;
  var d3_geo_pathBounds = {
    point: d3_geo_pathBoundsPoint,
    lineStart: d3_noop,
    lineEnd: d3_noop,
    polygonStart: d3_noop,
    polygonEnd: d3_noop
  };
  function d3_geo_pathBoundsPoint(x, y) {
    if (x < d3_geo_pathBoundsX0) d3_geo_pathBoundsX0 = x;
    if (x > d3_geo_pathBoundsX1) d3_geo_pathBoundsX1 = x;
    if (y < d3_geo_pathBoundsY0) d3_geo_pathBoundsY0 = y;
    if (y > d3_geo_pathBoundsY1) d3_geo_pathBoundsY1 = y;
  }
  function d3_geo_pathBuffer() {
    var pointCircle = d3_geo_pathBufferCircle(4.5), buffer = [];
    var stream = {
      point: point,
      lineStart: function() {
        stream.point = pointLineStart;
      },
      lineEnd: lineEnd,
      polygonStart: function() {
        stream.lineEnd = lineEndPolygon;
      },
      polygonEnd: function() {
        stream.lineEnd = lineEnd;
        stream.point = point;
      },
      pointRadius: function(_) {
        pointCircle = d3_geo_pathBufferCircle(_);
        return stream;
      },
      result: function() {
        if (buffer.length) {
          var result = buffer.join("");
          buffer = [];
          return result;
        }
      }
    };
    function point(x, y) {
      buffer.push("M", x, ",", y, pointCircle);
    }
    function pointLineStart(x, y) {
      buffer.push("M", x, ",", y);
      stream.point = pointLine;
    }
    function pointLine(x, y) {
      buffer.push("L", x, ",", y);
    }
    function lineEnd() {
      stream.point = point;
    }
    function lineEndPolygon() {
      buffer.push("Z");
    }
    return stream;
  }
  function d3_geo_pathBufferCircle(radius) {
    return "m0," + radius + "a" + radius + "," + radius + " 0 1,1 0," + -2 * radius + "a" + radius + "," + radius + " 0 1,1 0," + 2 * radius + "z";
  }
  var d3_geo_pathCentroid = {
    point: d3_geo_pathCentroidPoint,
    lineStart: d3_geo_pathCentroidLineStart,
    lineEnd: d3_geo_pathCentroidLineEnd,
    polygonStart: function() {
      d3_geo_pathCentroid.lineStart = d3_geo_pathCentroidRingStart;
    },
    polygonEnd: function() {
      d3_geo_pathCentroid.point = d3_geo_pathCentroidPoint;
      d3_geo_pathCentroid.lineStart = d3_geo_pathCentroidLineStart;
      d3_geo_pathCentroid.lineEnd = d3_geo_pathCentroidLineEnd;
    }
  };
  function d3_geo_pathCentroidPoint(x, y) {
    d3_geo_centroidX0 += x;
    d3_geo_centroidY0 += y;
    ++d3_geo_centroidZ0;
  }
  function d3_geo_pathCentroidLineStart() {
    var x0, y0;
    d3_geo_pathCentroid.point = function(x, y) {
      d3_geo_pathCentroid.point = nextPoint;
      d3_geo_pathCentroidPoint(x0 = x, y0 = y);
    };
    function nextPoint(x, y) {
      var dx = x - x0, dy = y - y0, z = Math.sqrt(dx * dx + dy * dy);
      d3_geo_centroidX1 += z * (x0 + x) / 2;
      d3_geo_centroidY1 += z * (y0 + y) / 2;
      d3_geo_centroidZ1 += z;
      d3_geo_pathCentroidPoint(x0 = x, y0 = y);
    }
  }
  function d3_geo_pathCentroidLineEnd() {
    d3_geo_pathCentroid.point = d3_geo_pathCentroidPoint;
  }
  function d3_geo_pathCentroidRingStart() {
    var x00, y00, x0, y0;
    d3_geo_pathCentroid.point = function(x, y) {
      d3_geo_pathCentroid.point = nextPoint;
      d3_geo_pathCentroidPoint(x00 = x0 = x, y00 = y0 = y);
    };
    function nextPoint(x, y) {
      var dx = x - x0, dy = y - y0, z = Math.sqrt(dx * dx + dy * dy);
      d3_geo_centroidX1 += z * (x0 + x) / 2;
      d3_geo_centroidY1 += z * (y0 + y) / 2;
      d3_geo_centroidZ1 += z;
      z = y0 * x - x0 * y;
      d3_geo_centroidX2 += z * (x0 + x);
      d3_geo_centroidY2 += z * (y0 + y);
      d3_geo_centroidZ2 += z * 3;
      d3_geo_pathCentroidPoint(x0 = x, y0 = y);
    }
    d3_geo_pathCentroid.lineEnd = function() {
      nextPoint(x00, y00);
    };
  }
  function d3_geo_pathContext(context) {
    var pointRadius = 4.5;
    var stream = {
      point: point,
      lineStart: function() {
        stream.point = pointLineStart;
      },
      lineEnd: lineEnd,
      polygonStart: function() {
        stream.lineEnd = lineEndPolygon;
      },
      polygonEnd: function() {
        stream.lineEnd = lineEnd;
        stream.point = point;
      },
      pointRadius: function(_) {
        pointRadius = _;
        return stream;
      },
      result: d3_noop
    };
    function point(x, y) {
      context.moveTo(x, y);
      context.arc(x, y, pointRadius, 0, τ);
    }
    function pointLineStart(x, y) {
      context.moveTo(x, y);
      stream.point = pointLine;
    }
    function pointLine(x, y) {
      context.lineTo(x, y);
    }
    function lineEnd() {
      stream.point = point;
    }
    function lineEndPolygon() {
      context.closePath();
    }
    return stream;
  }
  function d3_geo_resample(project) {
    var δ2 = .5, cosMinDistance = Math.cos(30 * d3_radians), maxDepth = 16;
    function resample(stream) {
      return (maxDepth ? resampleRecursive : resampleNone)(stream);
    }
    function resampleNone(stream) {
      return d3_geo_transformPoint(stream, function(x, y) {
        x = project(x, y);
        stream.point(x[0], x[1]);
      });
    }
    function resampleRecursive(stream) {
      var λ00, φ00, x00, y00, a00, b00, c00, λ0, x0, y0, a0, b0, c0;
      var resample = {
        point: point,
        lineStart: lineStart,
        lineEnd: lineEnd,
        polygonStart: function() {
          stream.polygonStart();
          resample.lineStart = ringStart;
        },
        polygonEnd: function() {
          stream.polygonEnd();
          resample.lineStart = lineStart;
        }
      };
      function point(x, y) {
        x = project(x, y);
        stream.point(x[0], x[1]);
      }
      function lineStart() {
        x0 = NaN;
        resample.point = linePoint;
        stream.lineStart();
      }
      function linePoint(λ, φ) {
        var c = d3_geo_cartesian([ λ, φ ]), p = project(λ, φ);
        resampleLineTo(x0, y0, λ0, a0, b0, c0, x0 = p[0], y0 = p[1], λ0 = λ, a0 = c[0], b0 = c[1], c0 = c[2], maxDepth, stream);
        stream.point(x0, y0);
      }
      function lineEnd() {
        resample.point = point;
        stream.lineEnd();
      }
      function ringStart() {
        lineStart();
        resample.point = ringPoint;
        resample.lineEnd = ringEnd;
      }
      function ringPoint(λ, φ) {
        linePoint(λ00 = λ, φ00 = φ), x00 = x0, y00 = y0, a00 = a0, b00 = b0, c00 = c0;
        resample.point = linePoint;
      }
      function ringEnd() {
        resampleLineTo(x0, y0, λ0, a0, b0, c0, x00, y00, λ00, a00, b00, c00, maxDepth, stream);
        resample.lineEnd = lineEnd;
        lineEnd();
      }
      return resample;
    }
    function resampleLineTo(x0, y0, λ0, a0, b0, c0, x1, y1, λ1, a1, b1, c1, depth, stream) {
      var dx = x1 - x0, dy = y1 - y0, d2 = dx * dx + dy * dy;
      if (d2 > 4 * δ2 && depth--) {
        var a = a0 + a1, b = b0 + b1, c = c0 + c1, m = Math.sqrt(a * a + b * b + c * c), φ2 = Math.asin(c /= m), λ2 = abs(abs(c) - 1) < ε || abs(λ0 - λ1) < ε ? (λ0 + λ1) / 2 : Math.atan2(b, a), p = project(λ2, φ2), x2 = p[0], y2 = p[1], dx2 = x2 - x0, dy2 = y2 - y0, dz = dy * dx2 - dx * dy2;
        if (dz * dz / d2 > δ2 || abs((dx * dx2 + dy * dy2) / d2 - .5) > .3 || a0 * a1 + b0 * b1 + c0 * c1 < cosMinDistance) {
          resampleLineTo(x0, y0, λ0, a0, b0, c0, x2, y2, λ2, a /= m, b /= m, c, depth, stream);
          stream.point(x2, y2);
          resampleLineTo(x2, y2, λ2, a, b, c, x1, y1, λ1, a1, b1, c1, depth, stream);
        }
      }
    }
    resample.precision = function(_) {
      if (!arguments.length) return Math.sqrt(δ2);
      maxDepth = (δ2 = _ * _) > 0 && 16;
      return resample;
    };
    return resample;
  }
  d3.geo.path = function() {
    var pointRadius = 4.5, projection, context, projectStream, contextStream, cacheStream;
    function path(object) {
      if (object) {
        if (typeof pointRadius === "function") contextStream.pointRadius(+pointRadius.apply(this, arguments));
        if (!cacheStream || !cacheStream.valid) cacheStream = projectStream(contextStream);
        d3.geo.stream(object, cacheStream);
      }
      return contextStream.result();
    }
    path.area = function(object) {
      d3_geo_pathAreaSum = 0;
      d3.geo.stream(object, projectStream(d3_geo_pathArea));
      return d3_geo_pathAreaSum;
    };
    path.centroid = function(object) {
      d3_geo_centroidX0 = d3_geo_centroidY0 = d3_geo_centroidZ0 = d3_geo_centroidX1 = d3_geo_centroidY1 = d3_geo_centroidZ1 = d3_geo_centroidX2 = d3_geo_centroidY2 = d3_geo_centroidZ2 = 0;
      d3.geo.stream(object, projectStream(d3_geo_pathCentroid));
      return d3_geo_centroidZ2 ? [ d3_geo_centroidX2 / d3_geo_centroidZ2, d3_geo_centroidY2 / d3_geo_centroidZ2 ] : d3_geo_centroidZ1 ? [ d3_geo_centroidX1 / d3_geo_centroidZ1, d3_geo_centroidY1 / d3_geo_centroidZ1 ] : d3_geo_centroidZ0 ? [ d3_geo_centroidX0 / d3_geo_centroidZ0, d3_geo_centroidY0 / d3_geo_centroidZ0 ] : [ NaN, NaN ];
    };
    path.bounds = function(object) {
      d3_geo_pathBoundsX1 = d3_geo_pathBoundsY1 = -(d3_geo_pathBoundsX0 = d3_geo_pathBoundsY0 = Infinity);
      d3.geo.stream(object, projectStream(d3_geo_pathBounds));
      return [ [ d3_geo_pathBoundsX0, d3_geo_pathBoundsY0 ], [ d3_geo_pathBoundsX1, d3_geo_pathBoundsY1 ] ];
    };
    path.projection = function(_) {
      if (!arguments.length) return projection;
      projectStream = (projection = _) ? _.stream || d3_geo_pathProjectStream(_) : d3_identity;
      return reset();
    };
    path.context = function(_) {
      if (!arguments.length) return context;
      contextStream = (context = _) == null ? new d3_geo_pathBuffer() : new d3_geo_pathContext(_);
      if (typeof pointRadius !== "function") contextStream.pointRadius(pointRadius);
      return reset();
    };
    path.pointRadius = function(_) {
      if (!arguments.length) return pointRadius;
      pointRadius = typeof _ === "function" ? _ : (contextStream.pointRadius(+_), +_);
      return path;
    };
    function reset() {
      cacheStream = null;
      return path;
    }
    return path.projection(d3.geo.albersUsa()).context(null);
  };
  function d3_geo_pathProjectStream(project) {
    var resample = d3_geo_resample(function(x, y) {
      return project([ x * d3_degrees, y * d3_degrees ]);
    });
    return function(stream) {
      return d3_geo_projectionRadians(resample(stream));
    };
  }
  d3.geo.transform = function(methods) {
    return {
      stream: function(stream) {
        var transform = new d3_geo_transform(stream);
        for (var k in methods) transform[k] = methods[k];
        return transform;
      }
    };
  };
  function d3_geo_transform(stream) {
    this.stream = stream;
  }
  d3_geo_transform.prototype = {
    point: function(x, y) {
      this.stream.point(x, y);
    },
    sphere: function() {
      this.stream.sphere();
    },
    lineStart: function() {
      this.stream.lineStart();
    },
    lineEnd: function() {
      this.stream.lineEnd();
    },
    polygonStart: function() {
      this.stream.polygonStart();
    },
    polygonEnd: function() {
      this.stream.polygonEnd();
    }
  };
  function d3_geo_transformPoint(stream, point) {
    return {
      point: point,
      sphere: function() {
        stream.sphere();
      },
      lineStart: function() {
        stream.lineStart();
      },
      lineEnd: function() {
        stream.lineEnd();
      },
      polygonStart: function() {
        stream.polygonStart();
      },
      polygonEnd: function() {
        stream.polygonEnd();
      }
    };
  }
  d3.geo.projection = d3_geo_projection;
  d3.geo.projectionMutator = d3_geo_projectionMutator;
  function d3_geo_projection(project) {
    return d3_geo_projectionMutator(function() {
      return project;
    })();
  }
  function d3_geo_projectionMutator(projectAt) {
    var project, rotate, projectRotate, projectResample = d3_geo_resample(function(x, y) {
      x = project(x, y);
      return [ x[0] * k + δx, δy - x[1] * k ];
    }), k = 150, x = 480, y = 250, λ = 0, φ = 0, δλ = 0, δφ = 0, δγ = 0, δx, δy, preclip = d3_geo_clipAntimeridian, postclip = d3_identity, clipAngle = null, clipExtent = null, stream;
    function projection(point) {
      point = projectRotate(point[0] * d3_radians, point[1] * d3_radians);
      return [ point[0] * k + δx, δy - point[1] * k ];
    }
    function invert(point) {
      point = projectRotate.invert((point[0] - δx) / k, (δy - point[1]) / k);
      return point && [ point[0] * d3_degrees, point[1] * d3_degrees ];
    }
    projection.stream = function(output) {
      if (stream) stream.valid = false;
      stream = d3_geo_projectionRadians(preclip(rotate, projectResample(postclip(output))));
      stream.valid = true;
      return stream;
    };
    projection.clipAngle = function(_) {
      if (!arguments.length) return clipAngle;
      preclip = _ == null ? (clipAngle = _, d3_geo_clipAntimeridian) : d3_geo_clipCircle((clipAngle = +_) * d3_radians);
      return invalidate();
    };
    projection.clipExtent = function(_) {
      if (!arguments.length) return clipExtent;
      clipExtent = _;
      postclip = _ ? d3_geo_clipExtent(_[0][0], _[0][1], _[1][0], _[1][1]) : d3_identity;
      return invalidate();
    };
    projection.scale = function(_) {
      if (!arguments.length) return k;
      k = +_;
      return reset();
    };
    projection.translate = function(_) {
      if (!arguments.length) return [ x, y ];
      x = +_[0];
      y = +_[1];
      return reset();
    };
    projection.center = function(_) {
      if (!arguments.length) return [ λ * d3_degrees, φ * d3_degrees ];
      λ = _[0] % 360 * d3_radians;
      φ = _[1] % 360 * d3_radians;
      return reset();
    };
    projection.rotate = function(_) {
      if (!arguments.length) return [ δλ * d3_degrees, δφ * d3_degrees, δγ * d3_degrees ];
      δλ = _[0] % 360 * d3_radians;
      δφ = _[1] % 360 * d3_radians;
      δγ = _.length > 2 ? _[2] % 360 * d3_radians : 0;
      return reset();
    };
    d3.rebind(projection, projectResample, "precision");
    function reset() {
      projectRotate = d3_geo_compose(rotate = d3_geo_rotation(δλ, δφ, δγ), project);
      var center = project(λ, φ);
      δx = x - center[0] * k;
      δy = y + center[1] * k;
      return invalidate();
    }
    function invalidate() {
      if (stream) stream.valid = false, stream = null;
      return projection;
    }
    return function() {
      project = projectAt.apply(this, arguments);
      projection.invert = project.invert && invert;
      return reset();
    };
  }
  function d3_geo_projectionRadians(stream) {
    return d3_geo_transformPoint(stream, function(x, y) {
      stream.point(x * d3_radians, y * d3_radians);
    });
  }
  function d3_geo_equirectangular(λ, φ) {
    return [ λ, φ ];
  }
  (d3.geo.equirectangular = function() {
    return d3_geo_projection(d3_geo_equirectangular);
  }).raw = d3_geo_equirectangular.invert = d3_geo_equirectangular;
  d3.geo.rotation = function(rotate) {
    rotate = d3_geo_rotation(rotate[0] % 360 * d3_radians, rotate[1] * d3_radians, rotate.length > 2 ? rotate[2] * d3_radians : 0);
    function forward(coordinates) {
      coordinates = rotate(coordinates[0] * d3_radians, coordinates[1] * d3_radians);
      return coordinates[0] *= d3_degrees, coordinates[1] *= d3_degrees, coordinates;
    }
    forward.invert = function(coordinates) {
      coordinates = rotate.invert(coordinates[0] * d3_radians, coordinates[1] * d3_radians);
      return coordinates[0] *= d3_degrees, coordinates[1] *= d3_degrees, coordinates;
    };
    return forward;
  };
  function d3_geo_identityRotation(λ, φ) {
    return [ λ > π ? λ - τ : λ < -π ? λ + τ : λ, φ ];
  }
  d3_geo_identityRotation.invert = d3_geo_equirectangular;
  function d3_geo_rotation(δλ, δφ, δγ) {
    return δλ ? δφ || δγ ? d3_geo_compose(d3_geo_rotationλ(δλ), d3_geo_rotationφγ(δφ, δγ)) : d3_geo_rotationλ(δλ) : δφ || δγ ? d3_geo_rotationφγ(δφ, δγ) : d3_geo_identityRotation;
  }
  function d3_geo_forwardRotationλ(δλ) {
    return function(λ, φ) {
      return λ += δλ, [ λ > π ? λ - τ : λ < -π ? λ + τ : λ, φ ];
    };
  }
  function d3_geo_rotationλ(δλ) {
    var rotation = d3_geo_forwardRotationλ(δλ);
    rotation.invert = d3_geo_forwardRotationλ(-δλ);
    return rotation;
  }
  function d3_geo_rotationφγ(δφ, δγ) {
    var cosδφ = Math.cos(δφ), sinδφ = Math.sin(δφ), cosδγ = Math.cos(δγ), sinδγ = Math.sin(δγ);
    function rotation(λ, φ) {
      var cosφ = Math.cos(φ), x = Math.cos(λ) * cosφ, y = Math.sin(λ) * cosφ, z = Math.sin(φ), k = z * cosδφ + x * sinδφ;
      return [ Math.atan2(y * cosδγ - k * sinδγ, x * cosδφ - z * sinδφ), d3_asin(k * cosδγ + y * sinδγ) ];
    }
    rotation.invert = function(λ, φ) {
      var cosφ = Math.cos(φ), x = Math.cos(λ) * cosφ, y = Math.sin(λ) * cosφ, z = Math.sin(φ), k = z * cosδγ - y * sinδγ;
      return [ Math.atan2(y * cosδγ + z * sinδγ, x * cosδφ + k * sinδφ), d3_asin(k * cosδφ - x * sinδφ) ];
    };
    return rotation;
  }
  d3.geo.circle = function() {
    var origin = [ 0, 0 ], angle, precision = 6, interpolate;
    function circle() {
      var center = typeof origin === "function" ? origin.apply(this, arguments) : origin, rotate = d3_geo_rotation(-center[0] * d3_radians, -center[1] * d3_radians, 0).invert, ring = [];
      interpolate(null, null, 1, {
        point: function(x, y) {
          ring.push(x = rotate(x, y));
          x[0] *= d3_degrees, x[1] *= d3_degrees;
        }
      });
      return {
        type: "Polygon",
        coordinates: [ ring ]
      };
    }
    circle.origin = function(x) {
      if (!arguments.length) return origin;
      origin = x;
      return circle;
    };
    circle.angle = function(x) {
      if (!arguments.length) return angle;
      interpolate = d3_geo_circleInterpolate((angle = +x) * d3_radians, precision * d3_radians);
      return circle;
    };
    circle.precision = function(_) {
      if (!arguments.length) return precision;
      interpolate = d3_geo_circleInterpolate(angle * d3_radians, (precision = +_) * d3_radians);
      return circle;
    };
    return circle.angle(90);
  };
  function d3_geo_circleInterpolate(radius, precision) {
    var cr = Math.cos(radius), sr = Math.sin(radius);
    return function(from, to, direction, listener) {
      var step = direction * precision;
      if (from != null) {
        from = d3_geo_circleAngle(cr, from);
        to = d3_geo_circleAngle(cr, to);
        if (direction > 0 ? from < to : from > to) from += direction * τ;
      } else {
        from = radius + direction * τ;
        to = radius - .5 * step;
      }
      for (var point, t = from; direction > 0 ? t > to : t < to; t -= step) {
        listener.point((point = d3_geo_spherical([ cr, -sr * Math.cos(t), -sr * Math.sin(t) ]))[0], point[1]);
      }
    };
  }
  function d3_geo_circleAngle(cr, point) {
    var a = d3_geo_cartesian(point);
    a[0] -= cr;
    d3_geo_cartesianNormalize(a);
    var angle = d3_acos(-a[1]);
    return ((-a[2] < 0 ? -angle : angle) + 2 * Math.PI - ε) % (2 * Math.PI);
  }
  d3.geo.distance = function(a, b) {
    var Δλ = (b[0] - a[0]) * d3_radians, φ0 = a[1] * d3_radians, φ1 = b[1] * d3_radians, sinΔλ = Math.sin(Δλ), cosΔλ = Math.cos(Δλ), sinφ0 = Math.sin(φ0), cosφ0 = Math.cos(φ0), sinφ1 = Math.sin(φ1), cosφ1 = Math.cos(φ1), t;
    return Math.atan2(Math.sqrt((t = cosφ1 * sinΔλ) * t + (t = cosφ0 * sinφ1 - sinφ0 * cosφ1 * cosΔλ) * t), sinφ0 * sinφ1 + cosφ0 * cosφ1 * cosΔλ);
  };
  d3.geo.graticule = function() {
    var x1, x0, X1, X0, y1, y0, Y1, Y0, dx = 10, dy = dx, DX = 90, DY = 360, x, y, X, Y, precision = 2.5;
    function graticule() {
      return {
        type: "MultiLineString",
        coordinates: lines()
      };
    }
    function lines() {
      return d3.range(Math.ceil(X0 / DX) * DX, X1, DX).map(X).concat(d3.range(Math.ceil(Y0 / DY) * DY, Y1, DY).map(Y)).concat(d3.range(Math.ceil(x0 / dx) * dx, x1, dx).filter(function(x) {
        return abs(x % DX) > ε;
      }).map(x)).concat(d3.range(Math.ceil(y0 / dy) * dy, y1, dy).filter(function(y) {
        return abs(y % DY) > ε;
      }).map(y));
    }
    graticule.lines = function() {
      return lines().map(function(coordinates) {
        return {
          type: "LineString",
          coordinates: coordinates
        };
      });
    };
    graticule.outline = function() {
      return {
        type: "Polygon",
        coordinates: [ X(X0).concat(Y(Y1).slice(1), X(X1).reverse().slice(1), Y(Y0).reverse().slice(1)) ]
      };
    };
    graticule.extent = function(_) {
      if (!arguments.length) return graticule.minorExtent();
      return graticule.majorExtent(_).minorExtent(_);
    };
    graticule.majorExtent = function(_) {
      if (!arguments.length) return [ [ X0, Y0 ], [ X1, Y1 ] ];
      X0 = +_[0][0], X1 = +_[1][0];
      Y0 = +_[0][1], Y1 = +_[1][1];
      if (X0 > X1) _ = X0, X0 = X1, X1 = _;
      if (Y0 > Y1) _ = Y0, Y0 = Y1, Y1 = _;
      return graticule.precision(precision);
    };
    graticule.minorExtent = function(_) {
      if (!arguments.length) return [ [ x0, y0 ], [ x1, y1 ] ];
      x0 = +_[0][0], x1 = +_[1][0];
      y0 = +_[0][1], y1 = +_[1][1];
      if (x0 > x1) _ = x0, x0 = x1, x1 = _;
      if (y0 > y1) _ = y0, y0 = y1, y1 = _;
      return graticule.precision(precision);
    };
    graticule.step = function(_) {
      if (!arguments.length) return graticule.minorStep();
      return graticule.majorStep(_).minorStep(_);
    };
    graticule.majorStep = function(_) {
      if (!arguments.length) return [ DX, DY ];
      DX = +_[0], DY = +_[1];
      return graticule;
    };
    graticule.minorStep = function(_) {
      if (!arguments.length) return [ dx, dy ];
      dx = +_[0], dy = +_[1];
      return graticule;
    };
    graticule.precision = function(_) {
      if (!arguments.length) return precision;
      precision = +_;
      x = d3_geo_graticuleX(y0, y1, 90);
      y = d3_geo_graticuleY(x0, x1, precision);
      X = d3_geo_graticuleX(Y0, Y1, 90);
      Y = d3_geo_graticuleY(X0, X1, precision);
      return graticule;
    };
    return graticule.majorExtent([ [ -180, -90 + ε ], [ 180, 90 - ε ] ]).minorExtent([ [ -180, -80 - ε ], [ 180, 80 + ε ] ]);
  };
  function d3_geo_graticuleX(y0, y1, dy) {
    var y = d3.range(y0, y1 - ε, dy).concat(y1);
    return function(x) {
      return y.map(function(y) {
        return [ x, y ];
      });
    };
  }
  function d3_geo_graticuleY(x0, x1, dx) {
    var x = d3.range(x0, x1 - ε, dx).concat(x1);
    return function(y) {
      return x.map(function(x) {
        return [ x, y ];
      });
    };
  }
  function d3_source(d) {
    return d.source;
  }
  function d3_target(d) {
    return d.target;
  }
  d3.geo.greatArc = function() {
    var source = d3_source, source_, target = d3_target, target_;
    function greatArc() {
      return {
        type: "LineString",
        coordinates: [ source_ || source.apply(this, arguments), target_ || target.apply(this, arguments) ]
      };
    }
    greatArc.distance = function() {
      return d3.geo.distance(source_ || source.apply(this, arguments), target_ || target.apply(this, arguments));
    };
    greatArc.source = function(_) {
      if (!arguments.length) return source;
      source = _, source_ = typeof _ === "function" ? null : _;
      return greatArc;
    };
    greatArc.target = function(_) {
      if (!arguments.length) return target;
      target = _, target_ = typeof _ === "function" ? null : _;
      return greatArc;
    };
    greatArc.precision = function() {
      return arguments.length ? greatArc : 0;
    };
    return greatArc;
  };
  d3.geo.interpolate = function(source, target) {
    return d3_geo_interpolate(source[0] * d3_radians, source[1] * d3_radians, target[0] * d3_radians, target[1] * d3_radians);
  };
  function d3_geo_interpolate(x0, y0, x1, y1) {
    var cy0 = Math.cos(y0), sy0 = Math.sin(y0), cy1 = Math.cos(y1), sy1 = Math.sin(y1), kx0 = cy0 * Math.cos(x0), ky0 = cy0 * Math.sin(x0), kx1 = cy1 * Math.cos(x1), ky1 = cy1 * Math.sin(x1), d = 2 * Math.asin(Math.sqrt(d3_haversin(y1 - y0) + cy0 * cy1 * d3_haversin(x1 - x0))), k = 1 / Math.sin(d);
    var interpolate = d ? function(t) {
      var B = Math.sin(t *= d) * k, A = Math.sin(d - t) * k, x = A * kx0 + B * kx1, y = A * ky0 + B * ky1, z = A * sy0 + B * sy1;
      return [ Math.atan2(y, x) * d3_degrees, Math.atan2(z, Math.sqrt(x * x + y * y)) * d3_degrees ];
    } : function() {
      return [ x0 * d3_degrees, y0 * d3_degrees ];
    };
    interpolate.distance = d;
    return interpolate;
  }
  d3.geo.length = function(object) {
    d3_geo_lengthSum = 0;
    d3.geo.stream(object, d3_geo_length);
    return d3_geo_lengthSum;
  };
  var d3_geo_lengthSum;
  var d3_geo_length = {
    sphere: d3_noop,
    point: d3_noop,
    lineStart: d3_geo_lengthLineStart,
    lineEnd: d3_noop,
    polygonStart: d3_noop,
    polygonEnd: d3_noop
  };
  function d3_geo_lengthLineStart() {
    var λ0, sinφ0, cosφ0;
    d3_geo_length.point = function(λ, φ) {
      λ0 = λ * d3_radians, sinφ0 = Math.sin(φ *= d3_radians), cosφ0 = Math.cos(φ);
      d3_geo_length.point = nextPoint;
    };
    d3_geo_length.lineEnd = function() {
      d3_geo_length.point = d3_geo_length.lineEnd = d3_noop;
    };
    function nextPoint(λ, φ) {
      var sinφ = Math.sin(φ *= d3_radians), cosφ = Math.cos(φ), t = abs((λ *= d3_radians) - λ0), cosΔλ = Math.cos(t);
      d3_geo_lengthSum += Math.atan2(Math.sqrt((t = cosφ * Math.sin(t)) * t + (t = cosφ0 * sinφ - sinφ0 * cosφ * cosΔλ) * t), sinφ0 * sinφ + cosφ0 * cosφ * cosΔλ);
      λ0 = λ, sinφ0 = sinφ, cosφ0 = cosφ;
    }
  }
  function d3_geo_azimuthal(scale, angle) {
    function azimuthal(λ, φ) {
      var cosλ = Math.cos(λ), cosφ = Math.cos(φ), k = scale(cosλ * cosφ);
      return [ k * cosφ * Math.sin(λ), k * Math.sin(φ) ];
    }
    azimuthal.invert = function(x, y) {
      var ρ = Math.sqrt(x * x + y * y), c = angle(ρ), sinc = Math.sin(c), cosc = Math.cos(c);
      return [ Math.atan2(x * sinc, ρ * cosc), Math.asin(ρ && y * sinc / ρ) ];
    };
    return azimuthal;
  }
  var d3_geo_azimuthalEqualArea = d3_geo_azimuthal(function(cosλcosφ) {
    return Math.sqrt(2 / (1 + cosλcosφ));
  }, function(ρ) {
    return 2 * Math.asin(ρ / 2);
  });
  (d3.geo.azimuthalEqualArea = function() {
    return d3_geo_projection(d3_geo_azimuthalEqualArea);
  }).raw = d3_geo_azimuthalEqualArea;
  var d3_geo_azimuthalEquidistant = d3_geo_azimuthal(function(cosλcosφ) {
    var c = Math.acos(cosλcosφ);
    return c && c / Math.sin(c);
  }, d3_identity);
  (d3.geo.azimuthalEquidistant = function() {
    return d3_geo_projection(d3_geo_azimuthalEquidistant);
  }).raw = d3_geo_azimuthalEquidistant;
  function d3_geo_conicConformal(φ0, φ1) {
    var cosφ0 = Math.cos(φ0), t = function(φ) {
      return Math.tan(π / 4 + φ / 2);
    }, n = φ0 === φ1 ? Math.sin(φ0) : Math.log(cosφ0 / Math.cos(φ1)) / Math.log(t(φ1) / t(φ0)), F = cosφ0 * Math.pow(t(φ0), n) / n;
    if (!n) return d3_geo_mercator;
    function forward(λ, φ) {
      var ρ = abs(abs(φ) - halfπ) < ε ? 0 : F / Math.pow(t(φ), n);
      return [ ρ * Math.sin(n * λ), F - ρ * Math.cos(n * λ) ];
    }
    forward.invert = function(x, y) {
      var ρ0_y = F - y, ρ = d3_sgn(n) * Math.sqrt(x * x + ρ0_y * ρ0_y);
      return [ Math.atan2(x, ρ0_y) / n, 2 * Math.atan(Math.pow(F / ρ, 1 / n)) - halfπ ];
    };
    return forward;
  }
  (d3.geo.conicConformal = function() {
    return d3_geo_conic(d3_geo_conicConformal);
  }).raw = d3_geo_conicConformal;
  function d3_geo_conicEquidistant(φ0, φ1) {
    var cosφ0 = Math.cos(φ0), n = φ0 === φ1 ? Math.sin(φ0) : (cosφ0 - Math.cos(φ1)) / (φ1 - φ0), G = cosφ0 / n + φ0;
    if (abs(n) < ε) return d3_geo_equirectangular;
    function forward(λ, φ) {
      var ρ = G - φ;
      return [ ρ * Math.sin(n * λ), G - ρ * Math.cos(n * λ) ];
    }
    forward.invert = function(x, y) {
      var ρ0_y = G - y;
      return [ Math.atan2(x, ρ0_y) / n, G - d3_sgn(n) * Math.sqrt(x * x + ρ0_y * ρ0_y) ];
    };
    return forward;
  }
  (d3.geo.conicEquidistant = function() {
    return d3_geo_conic(d3_geo_conicEquidistant);
  }).raw = d3_geo_conicEquidistant;
  var d3_geo_gnomonic = d3_geo_azimuthal(function(cosλcosφ) {
    return 1 / cosλcosφ;
  }, Math.atan);
  (d3.geo.gnomonic = function() {
    return d3_geo_projection(d3_geo_gnomonic);
  }).raw = d3_geo_gnomonic;
  function d3_geo_mercator(λ, φ) {
    return [ λ, Math.log(Math.tan(π / 4 + φ / 2)) ];
  }
  d3_geo_mercator.invert = function(x, y) {
    return [ x, 2 * Math.atan(Math.exp(y)) - halfπ ];
  };
  function d3_geo_mercatorProjection(project) {
    var m = d3_geo_projection(project), scale = m.scale, translate = m.translate, clipExtent = m.clipExtent, clipAuto;
    m.scale = function() {
      var v = scale.apply(m, arguments);
      return v === m ? clipAuto ? m.clipExtent(null) : m : v;
    };
    m.translate = function() {
      var v = translate.apply(m, arguments);
      return v === m ? clipAuto ? m.clipExtent(null) : m : v;
    };
    m.clipExtent = function(_) {
      var v = clipExtent.apply(m, arguments);
      if (v === m) {
        if (clipAuto = _ == null) {
          var k = π * scale(), t = translate();
          clipExtent([ [ t[0] - k, t[1] - k ], [ t[0] + k, t[1] + k ] ]);
        }
      } else if (clipAuto) {
        v = null;
      }
      return v;
    };
    return m.clipExtent(null);
  }
  (d3.geo.mercator = function() {
    return d3_geo_mercatorProjection(d3_geo_mercator);
  }).raw = d3_geo_mercator;
  var d3_geo_orthographic = d3_geo_azimuthal(function() {
    return 1;
  }, Math.asin);
  (d3.geo.orthographic = function() {
    return d3_geo_projection(d3_geo_orthographic);
  }).raw = d3_geo_orthographic;
  var d3_geo_stereographic = d3_geo_azimuthal(function(cosλcosφ) {
    return 1 / (1 + cosλcosφ);
  }, function(ρ) {
    return 2 * Math.atan(ρ);
  });
  (d3.geo.stereographic = function() {
    return d3_geo_projection(d3_geo_stereographic);
  }).raw = d3_geo_stereographic;
  function d3_geo_transverseMercator(λ, φ) {
    var B = Math.cos(φ) * Math.sin(λ);
    return [ Math.log((1 + B) / (1 - B)) / 2, Math.atan2(Math.tan(φ), Math.cos(λ)) ];
  }
  d3_geo_transverseMercator.invert = function(x, y) {
    return [ Math.atan2(d3_sinh(x), Math.cos(y)), d3_asin(Math.sin(y) / d3_cosh(x)) ];
  };
  (d3.geo.transverseMercator = function() {
    return d3_geo_mercatorProjection(d3_geo_transverseMercator);
  }).raw = d3_geo_transverseMercator;
  d3.geom = {};
  function d3_geom_pointX(d) {
    return d[0];
  }
  function d3_geom_pointY(d) {
    return d[1];
  }
  d3.geom.hull = function(vertices) {
    var x = d3_geom_pointX, y = d3_geom_pointY;
    if (arguments.length) return hull(vertices);
    function hull(data) {
      if (data.length < 3) return [];
      var fx = d3_functor(x), fy = d3_functor(y), n = data.length, vertices, plen = n - 1, points = [], stack = [], d, i, j, h = 0, x1, y1, x2, y2, u, v, a, sp;
      if (fx === d3_geom_pointX && y === d3_geom_pointY) vertices = data; else for (i = 0, 
      vertices = []; i < n; ++i) {
        vertices.push([ +fx.call(this, d = data[i], i), +fy.call(this, d, i) ]);
      }
      for (i = 1; i < n; ++i) {
        if (vertices[i][1] < vertices[h][1] || vertices[i][1] == vertices[h][1] && vertices[i][0] < vertices[h][0]) h = i;
      }
      for (i = 0; i < n; ++i) {
        if (i === h) continue;
        y1 = vertices[i][1] - vertices[h][1];
        x1 = vertices[i][0] - vertices[h][0];
        points.push({
          angle: Math.atan2(y1, x1),
          index: i
        });
      }
      points.sort(function(a, b) {
        return a.angle - b.angle;
      });
      a = points[0].angle;
      v = points[0].index;
      u = 0;
      for (i = 1; i < plen; ++i) {
        j = points[i].index;
        if (a == points[i].angle) {
          x1 = vertices[v][0] - vertices[h][0];
          y1 = vertices[v][1] - vertices[h][1];
          x2 = vertices[j][0] - vertices[h][0];
          y2 = vertices[j][1] - vertices[h][1];
          if (x1 * x1 + y1 * y1 >= x2 * x2 + y2 * y2) {
            points[i].index = -1;
            continue;
          } else {
            points[u].index = -1;
          }
        }
        a = points[i].angle;
        u = i;
        v = j;
      }
      stack.push(h);
      for (i = 0, j = 0; i < 2; ++j) {
        if (points[j].index > -1) {
          stack.push(points[j].index);
          i++;
        }
      }
      sp = stack.length;
      for (;j < plen; ++j) {
        if (points[j].index < 0) continue;
        while (!d3_geom_hullCCW(stack[sp - 2], stack[sp - 1], points[j].index, vertices)) {
          --sp;
        }
        stack[sp++] = points[j].index;
      }
      var poly = [];
      for (i = sp - 1; i >= 0; --i) poly.push(data[stack[i]]);
      return poly;
    }
    hull.x = function(_) {
      return arguments.length ? (x = _, hull) : x;
    };
    hull.y = function(_) {
      return arguments.length ? (y = _, hull) : y;
    };
    return hull;
  };
  function d3_geom_hullCCW(i1, i2, i3, v) {
    var t, a, b, c, d, e, f;
    t = v[i1];
    a = t[0];
    b = t[1];
    t = v[i2];
    c = t[0];
    d = t[1];
    t = v[i3];
    e = t[0];
    f = t[1];
    return (f - b) * (c - a) - (d - b) * (e - a) > 0;
  }
  d3.geom.polygon = function(coordinates) {
    d3_subclass(coordinates, d3_geom_polygonPrototype);
    return coordinates;
  };
  var d3_geom_polygonPrototype = d3.geom.polygon.prototype = [];
  d3_geom_polygonPrototype.area = function() {
    var i = -1, n = this.length, a, b = this[n - 1], area = 0;
    while (++i < n) {
      a = b;
      b = this[i];
      area += a[1] * b[0] - a[0] * b[1];
    }
    return area * .5;
  };
  d3_geom_polygonPrototype.centroid = function(k) {
    var i = -1, n = this.length, x = 0, y = 0, a, b = this[n - 1], c;
    if (!arguments.length) k = -1 / (6 * this.area());
    while (++i < n) {
      a = b;
      b = this[i];
      c = a[0] * b[1] - b[0] * a[1];
      x += (a[0] + b[0]) * c;
      y += (a[1] + b[1]) * c;
    }
    return [ x * k, y * k ];
  };
  d3_geom_polygonPrototype.clip = function(subject) {
    var input, closed = d3_geom_polygonClosed(subject), i = -1, n = this.length - d3_geom_polygonClosed(this), j, m, a = this[n - 1], b, c, d;
    while (++i < n) {
      input = subject.slice();
      subject.length = 0;
      b = this[i];
      c = input[(m = input.length - closed) - 1];
      j = -1;
      while (++j < m) {
        d = input[j];
        if (d3_geom_polygonInside(d, a, b)) {
          if (!d3_geom_polygonInside(c, a, b)) {
            subject.push(d3_geom_polygonIntersect(c, d, a, b));
          }
          subject.push(d);
        } else if (d3_geom_polygonInside(c, a, b)) {
          subject.push(d3_geom_polygonIntersect(c, d, a, b));
        }
        c = d;
      }
      if (closed) subject.push(subject[0]);
      a = b;
    }
    return subject;
  };
  function d3_geom_polygonInside(p, a, b) {
    return (b[0] - a[0]) * (p[1] - a[1]) < (b[1] - a[1]) * (p[0] - a[0]);
  }
  function d3_geom_polygonIntersect(c, d, a, b) {
    var x1 = c[0], x3 = a[0], x21 = d[0] - x1, x43 = b[0] - x3, y1 = c[1], y3 = a[1], y21 = d[1] - y1, y43 = b[1] - y3, ua = (x43 * (y1 - y3) - y43 * (x1 - x3)) / (y43 * x21 - x43 * y21);
    return [ x1 + ua * x21, y1 + ua * y21 ];
  }
  function d3_geom_polygonClosed(coordinates) {
    var a = coordinates[0], b = coordinates[coordinates.length - 1];
    return !(a[0] - b[0] || a[1] - b[1]);
  }
  var d3_geom_voronoiEdges, d3_geom_voronoiCells, d3_geom_voronoiBeaches, d3_geom_voronoiBeachPool = [], d3_geom_voronoiFirstCircle, d3_geom_voronoiCircles, d3_geom_voronoiCirclePool = [];
  function d3_geom_voronoiBeach() {
    d3_geom_voronoiRedBlackNode(this);
    this.edge = this.site = this.circle = null;
  }
  function d3_geom_voronoiCreateBeach(site) {
    var beach = d3_geom_voronoiBeachPool.pop() || new d3_geom_voronoiBeach();
    beach.site = site;
    return beach;
  }
  function d3_geom_voronoiDetachBeach(beach) {
    d3_geom_voronoiDetachCircle(beach);
    d3_geom_voronoiBeaches.remove(beach);
    d3_geom_voronoiBeachPool.push(beach);
    d3_geom_voronoiRedBlackNode(beach);
  }
  function d3_geom_voronoiRemoveBeach(beach) {
    var circle = beach.circle, x = circle.x, y = circle.cy, vertex = {
      x: x,
      y: y
    }, previous = beach.P, next = beach.N, disappearing = [ beach ];
    d3_geom_voronoiDetachBeach(beach);
    var lArc = previous;
    while (lArc.circle && abs(x - lArc.circle.x) < ε && abs(y - lArc.circle.cy) < ε) {
      previous = lArc.P;
      disappearing.unshift(lArc);
      d3_geom_voronoiDetachBeach(lArc);
      lArc = previous;
    }
    disappearing.unshift(lArc);
    d3_geom_voronoiDetachCircle(lArc);
    var rArc = next;
    while (rArc.circle && abs(x - rArc.circle.x) < ε && abs(y - rArc.circle.cy) < ε) {
      next = rArc.N;
      disappearing.push(rArc);
      d3_geom_voronoiDetachBeach(rArc);
      rArc = next;
    }
    disappearing.push(rArc);
    d3_geom_voronoiDetachCircle(rArc);
    var nArcs = disappearing.length, iArc;
    for (iArc = 1; iArc < nArcs; ++iArc) {
      rArc = disappearing[iArc];
      lArc = disappearing[iArc - 1];
      d3_geom_voronoiSetEdgeEnd(rArc.edge, lArc.site, rArc.site, vertex);
    }
    lArc = disappearing[0];
    rArc = disappearing[nArcs - 1];
    rArc.edge = d3_geom_voronoiCreateEdge(lArc.site, rArc.site, null, vertex);
    d3_geom_voronoiAttachCircle(lArc);
    d3_geom_voronoiAttachCircle(rArc);
  }
  function d3_geom_voronoiAddBeach(site) {
    var x = site.x, directrix = site.y, lArc, rArc, dxl, dxr, node = d3_geom_voronoiBeaches._;
    while (node) {
      dxl = d3_geom_voronoiLeftBreakPoint(node, directrix) - x;
      if (dxl > ε) node = node.L; else {
        dxr = x - d3_geom_voronoiRightBreakPoint(node, directrix);
        if (dxr > ε) {
          if (!node.R) {
            lArc = node;
            break;
          }
          node = node.R;
        } else {
          if (dxl > -ε) {
            lArc = node.P;
            rArc = node;
          } else if (dxr > -ε) {
            lArc = node;
            rArc = node.N;
          } else {
            lArc = rArc = node;
          }
          break;
        }
      }
    }
    var newArc = d3_geom_voronoiCreateBeach(site);
    d3_geom_voronoiBeaches.insert(lArc, newArc);
    if (!lArc && !rArc) return;
    if (lArc === rArc) {
      d3_geom_voronoiDetachCircle(lArc);
      rArc = d3_geom_voronoiCreateBeach(lArc.site);
      d3_geom_voronoiBeaches.insert(newArc, rArc);
      newArc.edge = rArc.edge = d3_geom_voronoiCreateEdge(lArc.site, newArc.site);
      d3_geom_voronoiAttachCircle(lArc);
      d3_geom_voronoiAttachCircle(rArc);
      return;
    }
    if (!rArc) {
      newArc.edge = d3_geom_voronoiCreateEdge(lArc.site, newArc.site);
      return;
    }
    d3_geom_voronoiDetachCircle(lArc);
    d3_geom_voronoiDetachCircle(rArc);
    var lSite = lArc.site, ax = lSite.x, ay = lSite.y, bx = site.x - ax, by = site.y - ay, rSite = rArc.site, cx = rSite.x - ax, cy = rSite.y - ay, d = 2 * (bx * cy - by * cx), hb = bx * bx + by * by, hc = cx * cx + cy * cy, vertex = {
      x: (cy * hb - by * hc) / d + ax,
      y: (bx * hc - cx * hb) / d + ay
    };
    d3_geom_voronoiSetEdgeEnd(rArc.edge, lSite, rSite, vertex);
    newArc.edge = d3_geom_voronoiCreateEdge(lSite, site, null, vertex);
    rArc.edge = d3_geom_voronoiCreateEdge(site, rSite, null, vertex);
    d3_geom_voronoiAttachCircle(lArc);
    d3_geom_voronoiAttachCircle(rArc);
  }
  function d3_geom_voronoiLeftBreakPoint(arc, directrix) {
    var site = arc.site, rfocx = site.x, rfocy = site.y, pby2 = rfocy - directrix;
    if (!pby2) return rfocx;
    var lArc = arc.P;
    if (!lArc) return -Infinity;
    site = lArc.site;
    var lfocx = site.x, lfocy = site.y, plby2 = lfocy - directrix;
    if (!plby2) return lfocx;
    var hl = lfocx - rfocx, aby2 = 1 / pby2 - 1 / plby2, b = hl / plby2;
    if (aby2) return (-b + Math.sqrt(b * b - 2 * aby2 * (hl * hl / (-2 * plby2) - lfocy + plby2 / 2 + rfocy - pby2 / 2))) / aby2 + rfocx;
    return (rfocx + lfocx) / 2;
  }
  function d3_geom_voronoiRightBreakPoint(arc, directrix) {
    var rArc = arc.N;
    if (rArc) return d3_geom_voronoiLeftBreakPoint(rArc, directrix);
    var site = arc.site;
    return site.y === directrix ? site.x : Infinity;
  }
  function d3_geom_voronoiCell(site) {
    this.site = site;
    this.edges = [];
  }
  d3_geom_voronoiCell.prototype.prepare = function() {
    var halfEdges = this.edges, iHalfEdge = halfEdges.length, edge;
    while (iHalfEdge--) {
      edge = halfEdges[iHalfEdge].edge;
      if (!edge.b || !edge.a) halfEdges.splice(iHalfEdge, 1);
    }
    halfEdges.sort(d3_geom_voronoiHalfEdgeOrder);
    return halfEdges.length;
  };
  function d3_geom_voronoiCloseCells(extent) {
    var x0 = extent[0][0], x1 = extent[1][0], y0 = extent[0][1], y1 = extent[1][1], x2, y2, x3, y3, cells = d3_geom_voronoiCells, iCell = cells.length, cell, iHalfEdge, halfEdges, nHalfEdges, start, end;
    while (iCell--) {
      cell = cells[iCell];
      if (!cell || !cell.prepare()) continue;
      halfEdges = cell.edges;
      nHalfEdges = halfEdges.length;
      iHalfEdge = 0;
      while (iHalfEdge < nHalfEdges) {
        end = halfEdges[iHalfEdge].end(), x3 = end.x, y3 = end.y;
        start = halfEdges[++iHalfEdge % nHalfEdges].start(), x2 = start.x, y2 = start.y;
        if (abs(x3 - x2) > ε || abs(y3 - y2) > ε) {
          halfEdges.splice(iHalfEdge, 0, new d3_geom_voronoiHalfEdge(d3_geom_voronoiCreateBorderEdge(cell.site, end, abs(x3 - x0) < ε && y1 - y3 > ε ? {
            x: x0,
            y: abs(x2 - x0) < ε ? y2 : y1
          } : abs(y3 - y1) < ε && x1 - x3 > ε ? {
            x: abs(y2 - y1) < ε ? x2 : x1,
            y: y1
          } : abs(x3 - x1) < ε && y3 - y0 > ε ? {
            x: x1,
            y: abs(x2 - x1) < ε ? y2 : y0
          } : abs(y3 - y0) < ε && x3 - x0 > ε ? {
            x: abs(y2 - y0) < ε ? x2 : x0,
            y: y0
          } : null), cell.site, null));
          ++nHalfEdges;
        }
      }
    }
  }
  function d3_geom_voronoiHalfEdgeOrder(a, b) {
    return b.angle - a.angle;
  }
  function d3_geom_voronoiCircle() {
    d3_geom_voronoiRedBlackNode(this);
    this.x = this.y = this.arc = this.site = this.cy = null;
  }
  function d3_geom_voronoiAttachCircle(arc) {
    var lArc = arc.P, rArc = arc.N;
    if (!lArc || !rArc) return;
    var lSite = lArc.site, cSite = arc.site, rSite = rArc.site;
    if (lSite === rSite) return;
    var bx = cSite.x, by = cSite.y, ax = lSite.x - bx, ay = lSite.y - by, cx = rSite.x - bx, cy = rSite.y - by;
    var d = 2 * (ax * cy - ay * cx);
    if (d >= -ε2) return;
    var ha = ax * ax + ay * ay, hc = cx * cx + cy * cy, x = (cy * ha - ay * hc) / d, y = (ax * hc - cx * ha) / d, cy = y + by;
    var circle = d3_geom_voronoiCirclePool.pop() || new d3_geom_voronoiCircle();
    circle.arc = arc;
    circle.site = cSite;
    circle.x = x + bx;
    circle.y = cy + Math.sqrt(x * x + y * y);
    circle.cy = cy;
    arc.circle = circle;
    var before = null, node = d3_geom_voronoiCircles._;
    while (node) {
      if (circle.y < node.y || circle.y === node.y && circle.x <= node.x) {
        if (node.L) node = node.L; else {
          before = node.P;
          break;
        }
      } else {
        if (node.R) node = node.R; else {
          before = node;
          break;
        }
      }
    }
    d3_geom_voronoiCircles.insert(before, circle);
    if (!before) d3_geom_voronoiFirstCircle = circle;
  }
  function d3_geom_voronoiDetachCircle(arc) {
    var circle = arc.circle;
    if (circle) {
      if (!circle.P) d3_geom_voronoiFirstCircle = circle.N;
      d3_geom_voronoiCircles.remove(circle);
      d3_geom_voronoiCirclePool.push(circle);
      d3_geom_voronoiRedBlackNode(circle);
      arc.circle = null;
    }
  }
  function d3_geom_voronoiClipEdges(extent) {
    var edges = d3_geom_voronoiEdges, clip = d3_geom_clipLine(extent[0][0], extent[0][1], extent[1][0], extent[1][1]), i = edges.length, e;
    while (i--) {
      e = edges[i];
      if (!d3_geom_voronoiConnectEdge(e, extent) || !clip(e) || abs(e.a.x - e.b.x) < ε && abs(e.a.y - e.b.y) < ε) {
        e.a = e.b = null;
        edges.splice(i, 1);
      }
    }
  }
  function d3_geom_voronoiConnectEdge(edge, extent) {
    var vb = edge.b;
    if (vb) return true;
    var va = edge.a, x0 = extent[0][0], x1 = extent[1][0], y0 = extent[0][1], y1 = extent[1][1], lSite = edge.l, rSite = edge.r, lx = lSite.x, ly = lSite.y, rx = rSite.x, ry = rSite.y, fx = (lx + rx) / 2, fy = (ly + ry) / 2, fm, fb;
    if (ry === ly) {
      if (fx < x0 || fx >= x1) return;
      if (lx > rx) {
        if (!va) va = {
          x: fx,
          y: y0
        }; else if (va.y >= y1) return;
        vb = {
          x: fx,
          y: y1
        };
      } else {
        if (!va) va = {
          x: fx,
          y: y1
        }; else if (va.y < y0) return;
        vb = {
          x: fx,
          y: y0
        };
      }
    } else {
      fm = (lx - rx) / (ry - ly);
      fb = fy - fm * fx;
      if (fm < -1 || fm > 1) {
        if (lx > rx) {
          if (!va) va = {
            x: (y0 - fb) / fm,
            y: y0
          }; else if (va.y >= y1) return;
          vb = {
            x: (y1 - fb) / fm,
            y: y1
          };
        } else {
          if (!va) va = {
            x: (y1 - fb) / fm,
            y: y1
          }; else if (va.y < y0) return;
          vb = {
            x: (y0 - fb) / fm,
            y: y0
          };
        }
      } else {
        if (ly < ry) {
          if (!va) va = {
            x: x0,
            y: fm * x0 + fb
          }; else if (va.x >= x1) return;
          vb = {
            x: x1,
            y: fm * x1 + fb
          };
        } else {
          if (!va) va = {
            x: x1,
            y: fm * x1 + fb
          }; else if (va.x < x0) return;
          vb = {
            x: x0,
            y: fm * x0 + fb
          };
        }
      }
    }
    edge.a = va;
    edge.b = vb;
    return true;
  }
  function d3_geom_voronoiEdge(lSite, rSite) {
    this.l = lSite;
    this.r = rSite;
    this.a = this.b = null;
  }
  function d3_geom_voronoiCreateEdge(lSite, rSite, va, vb) {
    var edge = new d3_geom_voronoiEdge(lSite, rSite);
    d3_geom_voronoiEdges.push(edge);
    if (va) d3_geom_voronoiSetEdgeEnd(edge, lSite, rSite, va);
    if (vb) d3_geom_voronoiSetEdgeEnd(edge, rSite, lSite, vb);
    d3_geom_voronoiCells[lSite.i].edges.push(new d3_geom_voronoiHalfEdge(edge, lSite, rSite));
    d3_geom_voronoiCells[rSite.i].edges.push(new d3_geom_voronoiHalfEdge(edge, rSite, lSite));
    return edge;
  }
  function d3_geom_voronoiCreateBorderEdge(lSite, va, vb) {
    var edge = new d3_geom_voronoiEdge(lSite, null);
    edge.a = va;
    edge.b = vb;
    d3_geom_voronoiEdges.push(edge);
    return edge;
  }
  function d3_geom_voronoiSetEdgeEnd(edge, lSite, rSite, vertex) {
    if (!edge.a && !edge.b) {
      edge.a = vertex;
      edge.l = lSite;
      edge.r = rSite;
    } else if (edge.l === rSite) {
      edge.b = vertex;
    } else {
      edge.a = vertex;
    }
  }
  function d3_geom_voronoiHalfEdge(edge, lSite, rSite) {
    var va = edge.a, vb = edge.b;
    this.edge = edge;
    this.site = lSite;
    this.angle = rSite ? Math.atan2(rSite.y - lSite.y, rSite.x - lSite.x) : edge.l === lSite ? Math.atan2(vb.x - va.x, va.y - vb.y) : Math.atan2(va.x - vb.x, vb.y - va.y);
  }
  d3_geom_voronoiHalfEdge.prototype = {
    start: function() {
      return this.edge.l === this.site ? this.edge.a : this.edge.b;
    },
    end: function() {
      return this.edge.l === this.site ? this.edge.b : this.edge.a;
    }
  };
  function d3_geom_voronoiRedBlackTree() {
    this._ = null;
  }
  function d3_geom_voronoiRedBlackNode(node) {
    node.U = node.C = node.L = node.R = node.P = node.N = null;
  }
  d3_geom_voronoiRedBlackTree.prototype = {
    insert: function(after, node) {
      var parent, grandpa, uncle;
      if (after) {
        node.P = after;
        node.N = after.N;
        if (after.N) after.N.P = node;
        after.N = node;
        if (after.R) {
          after = after.R;
          while (after.L) after = after.L;
          after.L = node;
        } else {
          after.R = node;
        }
        parent = after;
      } else if (this._) {
        after = d3_geom_voronoiRedBlackFirst(this._);
        node.P = null;
        node.N = after;
        after.P = after.L = node;
        parent = after;
      } else {
        node.P = node.N = null;
        this._ = node;
        parent = null;
      }
      node.L = node.R = null;
      node.U = parent;
      node.C = true;
      after = node;
      while (parent && parent.C) {
        grandpa = parent.U;
        if (parent === grandpa.L) {
          uncle = grandpa.R;
          if (uncle && uncle.C) {
            parent.C = uncle.C = false;
            grandpa.C = true;
            after = grandpa;
          } else {
            if (after === parent.R) {
              d3_geom_voronoiRedBlackRotateLeft(this, parent);
              after = parent;
              parent = after.U;
            }
            parent.C = false;
            grandpa.C = true;
            d3_geom_voronoiRedBlackRotateRight(this, grandpa);
          }
        } else {
          uncle = grandpa.L;
          if (uncle && uncle.C) {
            parent.C = uncle.C = false;
            grandpa.C = true;
            after = grandpa;
          } else {
            if (after === parent.L) {
              d3_geom_voronoiRedBlackRotateRight(this, parent);
              after = parent;
              parent = after.U;
            }
            parent.C = false;
            grandpa.C = true;
            d3_geom_voronoiRedBlackRotateLeft(this, grandpa);
          }
        }
        parent = after.U;
      }
      this._.C = false;
    },
    remove: function(node) {
      if (node.N) node.N.P = node.P;
      if (node.P) node.P.N = node.N;
      node.N = node.P = null;
      var parent = node.U, sibling, left = node.L, right = node.R, next, red;
      if (!left) next = right; else if (!right) next = left; else next = d3_geom_voronoiRedBlackFirst(right);
      if (parent) {
        if (parent.L === node) parent.L = next; else parent.R = next;
      } else {
        this._ = next;
      }
      if (left && right) {
        red = next.C;
        next.C = node.C;
        next.L = left;
        left.U = next;
        if (next !== right) {
          parent = next.U;
          next.U = node.U;
          node = next.R;
          parent.L = node;
          next.R = right;
          right.U = next;
        } else {
          next.U = parent;
          parent = next;
          node = next.R;
        }
      } else {
        red = node.C;
        node = next;
      }
      if (node) node.U = parent;
      if (red) return;
      if (node && node.C) {
        node.C = false;
        return;
      }
      do {
        if (node === this._) break;
        if (node === parent.L) {
          sibling = parent.R;
          if (sibling.C) {
            sibling.C = false;
            parent.C = true;
            d3_geom_voronoiRedBlackRotateLeft(this, parent);
            sibling = parent.R;
          }
          if (sibling.L && sibling.L.C || sibling.R && sibling.R.C) {
            if (!sibling.R || !sibling.R.C) {
              sibling.L.C = false;
              sibling.C = true;
              d3_geom_voronoiRedBlackRotateRight(this, sibling);
              sibling = parent.R;
            }
            sibling.C = parent.C;
            parent.C = sibling.R.C = false;
            d3_geom_voronoiRedBlackRotateLeft(this, parent);
            node = this._;
            break;
          }
        } else {
          sibling = parent.L;
          if (sibling.C) {
            sibling.C = false;
            parent.C = true;
            d3_geom_voronoiRedBlackRotateRight(this, parent);
            sibling = parent.L;
          }
          if (sibling.L && sibling.L.C || sibling.R && sibling.R.C) {
            if (!sibling.L || !sibling.L.C) {
              sibling.R.C = false;
              sibling.C = true;
              d3_geom_voronoiRedBlackRotateLeft(this, sibling);
              sibling = parent.L;
            }
            sibling.C = parent.C;
            parent.C = sibling.L.C = false;
            d3_geom_voronoiRedBlackRotateRight(this, parent);
            node = this._;
            break;
          }
        }
        sibling.C = true;
        node = parent;
        parent = parent.U;
      } while (!node.C);
      if (node) node.C = false;
    }
  };
  function d3_geom_voronoiRedBlackRotateLeft(tree, node) {
    var p = node, q = node.R, parent = p.U;
    if (parent) {
      if (parent.L === p) parent.L = q; else parent.R = q;
    } else {
      tree._ = q;
    }
    q.U = parent;
    p.U = q;
    p.R = q.L;
    if (p.R) p.R.U = p;
    q.L = p;
  }
  function d3_geom_voronoiRedBlackRotateRight(tree, node) {
    var p = node, q = node.L, parent = p.U;
    if (parent) {
      if (parent.L === p) parent.L = q; else parent.R = q;
    } else {
      tree._ = q;
    }
    q.U = parent;
    p.U = q;
    p.L = q.R;
    if (p.L) p.L.U = p;
    q.R = p;
  }
  function d3_geom_voronoiRedBlackFirst(node) {
    while (node.L) node = node.L;
    return node;
  }
  function d3_geom_voronoi(sites, bbox) {
    var site = sites.sort(d3_geom_voronoiVertexOrder).pop(), x0, y0, circle;
    d3_geom_voronoiEdges = [];
    d3_geom_voronoiCells = new Array(sites.length);
    d3_geom_voronoiBeaches = new d3_geom_voronoiRedBlackTree();
    d3_geom_voronoiCircles = new d3_geom_voronoiRedBlackTree();
    while (true) {
      circle = d3_geom_voronoiFirstCircle;
      if (site && (!circle || site.y < circle.y || site.y === circle.y && site.x < circle.x)) {
        if (site.x !== x0 || site.y !== y0) {
          d3_geom_voronoiCells[site.i] = new d3_geom_voronoiCell(site);
          d3_geom_voronoiAddBeach(site);
          x0 = site.x, y0 = site.y;
        }
        site = sites.pop();
      } else if (circle) {
        d3_geom_voronoiRemoveBeach(circle.arc);
      } else {
        break;
      }
    }
    if (bbox) d3_geom_voronoiClipEdges(bbox), d3_geom_voronoiCloseCells(bbox);
    var diagram = {
      cells: d3_geom_voronoiCells,
      edges: d3_geom_voronoiEdges
    };
    d3_geom_voronoiBeaches = d3_geom_voronoiCircles = d3_geom_voronoiEdges = d3_geom_voronoiCells = null;
    return diagram;
  }
  function d3_geom_voronoiVertexOrder(a, b) {
    return b.y - a.y || b.x - a.x;
  }
  d3.geom.voronoi = function(points) {
    var x = d3_geom_pointX, y = d3_geom_pointY, fx = x, fy = y, clipExtent = d3_geom_voronoiClipExtent;
    if (points) return voronoi(points);
    function voronoi(data) {
      var polygons = new Array(data.length), x0 = clipExtent[0][0], y0 = clipExtent[0][1], x1 = clipExtent[1][0], y1 = clipExtent[1][1];
      d3_geom_voronoi(sites(data), clipExtent).cells.forEach(function(cell, i) {
        var edges = cell.edges, site = cell.site, polygon = polygons[i] = edges.length ? edges.map(function(e) {
          var s = e.start();
          return [ s.x, s.y ];
        }) : site.x >= x0 && site.x <= x1 && site.y >= y0 && site.y <= y1 ? [ [ x0, y1 ], [ x1, y1 ], [ x1, y0 ], [ x0, y0 ] ] : [];
        polygon.point = data[i];
      });
      return polygons;
    }
    function sites(data) {
      return data.map(function(d, i) {
        return {
          x: Math.round(fx(d, i) / ε) * ε,
          y: Math.round(fy(d, i) / ε) * ε,
          i: i
        };
      });
    }
    voronoi.links = function(data) {
      return d3_geom_voronoi(sites(data)).edges.filter(function(edge) {
        return edge.l && edge.r;
      }).map(function(edge) {
        return {
          source: data[edge.l.i],
          target: data[edge.r.i]
        };
      });
    };
    voronoi.triangles = function(data) {
      var triangles = [];
      d3_geom_voronoi(sites(data)).cells.forEach(function(cell, i) {
        var site = cell.site, edges = cell.edges.sort(d3_geom_voronoiHalfEdgeOrder), j = -1, m = edges.length, e0, s0, e1 = edges[m - 1].edge, s1 = e1.l === site ? e1.r : e1.l;
        while (++j < m) {
          e0 = e1;
          s0 = s1;
          e1 = edges[j].edge;
          s1 = e1.l === site ? e1.r : e1.l;
          if (i < s0.i && i < s1.i && d3_geom_voronoiTriangleArea(site, s0, s1) < 0) {
            triangles.push([ data[i], data[s0.i], data[s1.i] ]);
          }
        }
      });
      return triangles;
    };
    voronoi.x = function(_) {
      return arguments.length ? (fx = d3_functor(x = _), voronoi) : x;
    };
    voronoi.y = function(_) {
      return arguments.length ? (fy = d3_functor(y = _), voronoi) : y;
    };
    voronoi.clipExtent = function(_) {
      if (!arguments.length) return clipExtent === d3_geom_voronoiClipExtent ? null : clipExtent;
      clipExtent = _ == null ? d3_geom_voronoiClipExtent : _;
      return voronoi;
    };
    voronoi.size = function(_) {
      if (!arguments.length) return clipExtent === d3_geom_voronoiClipExtent ? null : clipExtent && clipExtent[1];
      return voronoi.clipExtent(_ && [ [ 0, 0 ], _ ]);
    };
    return voronoi;
  };
  var d3_geom_voronoiClipExtent = [ [ -1e6, -1e6 ], [ 1e6, 1e6 ] ];
  function d3_geom_voronoiTriangleArea(a, b, c) {
    return (a.x - c.x) * (b.y - a.y) - (a.x - b.x) * (c.y - a.y);
  }
  d3.geom.delaunay = function(vertices) {
    return d3.geom.voronoi().triangles(vertices);
  };
  d3.geom.quadtree = function(points, x1, y1, x2, y2) {
    var x = d3_geom_pointX, y = d3_geom_pointY, compat;
    if (compat = arguments.length) {
      x = d3_geom_quadtreeCompatX;
      y = d3_geom_quadtreeCompatY;
      if (compat === 3) {
        y2 = y1;
        x2 = x1;
        y1 = x1 = 0;
      }
      return quadtree(points);
    }
    function quadtree(data) {
      var d, fx = d3_functor(x), fy = d3_functor(y), xs, ys, i, n, x1_, y1_, x2_, y2_;
      if (x1 != null) {
        x1_ = x1, y1_ = y1, x2_ = x2, y2_ = y2;
      } else {
        x2_ = y2_ = -(x1_ = y1_ = Infinity);
        xs = [], ys = [];
        n = data.length;
        if (compat) for (i = 0; i < n; ++i) {
          d = data[i];
          if (d.x < x1_) x1_ = d.x;
          if (d.y < y1_) y1_ = d.y;
          if (d.x > x2_) x2_ = d.x;
          if (d.y > y2_) y2_ = d.y;
          xs.push(d.x);
          ys.push(d.y);
        } else for (i = 0; i < n; ++i) {
          var x_ = +fx(d = data[i], i), y_ = +fy(d, i);
          if (x_ < x1_) x1_ = x_;
          if (y_ < y1_) y1_ = y_;
          if (x_ > x2_) x2_ = x_;
          if (y_ > y2_) y2_ = y_;
          xs.push(x_);
          ys.push(y_);
        }
      }
      var dx = x2_ - x1_, dy = y2_ - y1_;
      if (dx > dy) y2_ = y1_ + dx; else x2_ = x1_ + dy;
      function insert(n, d, x, y, x1, y1, x2, y2) {
        if (isNaN(x) || isNaN(y)) return;
        if (n.leaf) {
          var nx = n.x, ny = n.y;
          if (nx != null) {
            if (abs(nx - x) + abs(ny - y) < .01) {
              insertChild(n, d, x, y, x1, y1, x2, y2);
            } else {
              var nPoint = n.point;
              n.x = n.y = n.point = null;
              insertChild(n, nPoint, nx, ny, x1, y1, x2, y2);
              insertChild(n, d, x, y, x1, y1, x2, y2);
            }
          } else {
            n.x = x, n.y = y, n.point = d;
          }
        } else {
          insertChild(n, d, x, y, x1, y1, x2, y2);
        }
      }
      function insertChild(n, d, x, y, x1, y1, x2, y2) {
        var sx = (x1 + x2) * .5, sy = (y1 + y2) * .5, right = x >= sx, bottom = y >= sy, i = (bottom << 1) + right;
        n.leaf = false;
        n = n.nodes[i] || (n.nodes[i] = d3_geom_quadtreeNode());
        if (right) x1 = sx; else x2 = sx;
        if (bottom) y1 = sy; else y2 = sy;
        insert(n, d, x, y, x1, y1, x2, y2);
      }
      var root = d3_geom_quadtreeNode();
      root.add = function(d) {
        insert(root, d, +fx(d, ++i), +fy(d, i), x1_, y1_, x2_, y2_);
      };
      root.visit = function(f) {
        d3_geom_quadtreeVisit(f, root, x1_, y1_, x2_, y2_);
      };
      i = -1;
      if (x1 == null) {
        while (++i < n) {
          insert(root, data[i], xs[i], ys[i], x1_, y1_, x2_, y2_);
        }
        --i;
      } else data.forEach(root.add);
      xs = ys = data = d = null;
      return root;
    }
    quadtree.x = function(_) {
      return arguments.length ? (x = _, quadtree) : x;
    };
    quadtree.y = function(_) {
      return arguments.length ? (y = _, quadtree) : y;
    };
    quadtree.extent = function(_) {
      if (!arguments.length) return x1 == null ? null : [ [ x1, y1 ], [ x2, y2 ] ];
      if (_ == null) x1 = y1 = x2 = y2 = null; else x1 = +_[0][0], y1 = +_[0][1], x2 = +_[1][0], 
      y2 = +_[1][1];
      return quadtree;
    };
    quadtree.size = function(_) {
      if (!arguments.length) return x1 == null ? null : [ x2 - x1, y2 - y1 ];
      if (_ == null) x1 = y1 = x2 = y2 = null; else x1 = y1 = 0, x2 = +_[0], y2 = +_[1];
      return quadtree;
    };
    return quadtree;
  };
  function d3_geom_quadtreeCompatX(d) {
    return d.x;
  }
  function d3_geom_quadtreeCompatY(d) {
    return d.y;
  }
  function d3_geom_quadtreeNode() {
    return {
      leaf: true,
      nodes: [],
      point: null,
      x: null,
      y: null
    };
  }
  function d3_geom_quadtreeVisit(f, node, x1, y1, x2, y2) {
    if (!f(node, x1, y1, x2, y2)) {
      var sx = (x1 + x2) * .5, sy = (y1 + y2) * .5, children = node.nodes;
      if (children[0]) d3_geom_quadtreeVisit(f, children[0], x1, y1, sx, sy);
      if (children[1]) d3_geom_quadtreeVisit(f, children[1], sx, y1, x2, sy);
      if (children[2]) d3_geom_quadtreeVisit(f, children[2], x1, sy, sx, y2);
      if (children[3]) d3_geom_quadtreeVisit(f, children[3], sx, sy, x2, y2);
    }
  }
  d3.interpolateRgb = d3_interpolateRgb;
  function d3_interpolateRgb(a, b) {
    a = d3.rgb(a);
    b = d3.rgb(b);
    var ar = a.r, ag = a.g, ab = a.b, br = b.r - ar, bg = b.g - ag, bb = b.b - ab;
    return function(t) {
      return "#" + d3_rgb_hex(Math.round(ar + br * t)) + d3_rgb_hex(Math.round(ag + bg * t)) + d3_rgb_hex(Math.round(ab + bb * t));
    };
  }
  d3.interpolateObject = d3_interpolateObject;
  function d3_interpolateObject(a, b) {
    var i = {}, c = {}, k;
    for (k in a) {
      if (k in b) {
        i[k] = d3_interpolate(a[k], b[k]);
      } else {
        c[k] = a[k];
      }
    }
    for (k in b) {
      if (!(k in a)) {
        c[k] = b[k];
      }
    }
    return function(t) {
      for (k in i) c[k] = i[k](t);
      return c;
    };
  }
  d3.interpolateNumber = d3_interpolateNumber;
  function d3_interpolateNumber(a, b) {
    b -= a = +a;
    return function(t) {
      return a + b * t;
    };
  }
  d3.interpolateString = d3_interpolateString;
  function d3_interpolateString(a, b) {
    var m, i, j, s0 = 0, s1 = 0, s = [], q = [], n, o;
    a = a + "", b = b + "";
    d3_interpolate_number.lastIndex = 0;
    for (i = 0; m = d3_interpolate_number.exec(b); ++i) {
      if (m.index) s.push(b.substring(s0, s1 = m.index));
      q.push({
        i: s.length,
        x: m[0]
      });
      s.push(null);
      s0 = d3_interpolate_number.lastIndex;
    }
    if (s0 < b.length) s.push(b.substring(s0));
    for (i = 0, n = q.length; (m = d3_interpolate_number.exec(a)) && i < n; ++i) {
      o = q[i];
      if (o.x == m[0]) {
        if (o.i) {
          if (s[o.i + 1] == null) {
            s[o.i - 1] += o.x;
            s.splice(o.i, 1);
            for (j = i + 1; j < n; ++j) q[j].i--;
          } else {
            s[o.i - 1] += o.x + s[o.i + 1];
            s.splice(o.i, 2);
            for (j = i + 1; j < n; ++j) q[j].i -= 2;
          }
        } else {
          if (s[o.i + 1] == null) {
            s[o.i] = o.x;
          } else {
            s[o.i] = o.x + s[o.i + 1];
            s.splice(o.i + 1, 1);
            for (j = i + 1; j < n; ++j) q[j].i--;
          }
        }
        q.splice(i, 1);
        n--;
        i--;
      } else {
        o.x = d3_interpolateNumber(parseFloat(m[0]), parseFloat(o.x));
      }
    }
    while (i < n) {
      o = q.pop();
      if (s[o.i + 1] == null) {
        s[o.i] = o.x;
      } else {
        s[o.i] = o.x + s[o.i + 1];
        s.splice(o.i + 1, 1);
      }
      n--;
    }
    if (s.length === 1) {
      return s[0] == null ? (o = q[0].x, function(t) {
        return o(t) + "";
      }) : function() {
        return b;
      };
    }
    return function(t) {
      for (i = 0; i < n; ++i) s[(o = q[i]).i] = o.x(t);
      return s.join("");
    };
  }
  var d3_interpolate_number = /[-+]?(?:\d+\.?\d*|\.?\d+)(?:[eE][-+]?\d+)?/g;
  d3.interpolate = d3_interpolate;
  function d3_interpolate(a, b) {
    var i = d3.interpolators.length, f;
    while (--i >= 0 && !(f = d3.interpolators[i](a, b))) ;
    return f;
  }
  d3.interpolators = [ function(a, b) {
    var t = typeof b;
    return (t === "string" ? d3_rgb_names.has(b) || /^(#|rgb\(|hsl\()/.test(b) ? d3_interpolateRgb : d3_interpolateString : b instanceof d3_Color ? d3_interpolateRgb : t === "object" ? Array.isArray(b) ? d3_interpolateArray : d3_interpolateObject : d3_interpolateNumber)(a, b);
  } ];
  d3.interpolateArray = d3_interpolateArray;
  function d3_interpolateArray(a, b) {
    var x = [], c = [], na = a.length, nb = b.length, n0 = Math.min(a.length, b.length), i;
    for (i = 0; i < n0; ++i) x.push(d3_interpolate(a[i], b[i]));
    for (;i < na; ++i) c[i] = a[i];
    for (;i < nb; ++i) c[i] = b[i];
    return function(t) {
      for (i = 0; i < n0; ++i) c[i] = x[i](t);
      return c;
    };
  }
  var d3_ease_default = function() {
    return d3_identity;
  };
  var d3_ease = d3.map({
    linear: d3_ease_default,
    poly: d3_ease_poly,
    quad: function() {
      return d3_ease_quad;
    },
    cubic: function() {
      return d3_ease_cubic;
    },
    sin: function() {
      return d3_ease_sin;
    },
    exp: function() {
      return d3_ease_exp;
    },
    circle: function() {
      return d3_ease_circle;
    },
    elastic: d3_ease_elastic,
    back: d3_ease_back,
    bounce: function() {
      return d3_ease_bounce;
    }
  });
  var d3_ease_mode = d3.map({
    "in": d3_identity,
    out: d3_ease_reverse,
    "in-out": d3_ease_reflect,
    "out-in": function(f) {
      return d3_ease_reflect(d3_ease_reverse(f));
    }
  });
  d3.ease = function(name) {
    var i = name.indexOf("-"), t = i >= 0 ? name.substring(0, i) : name, m = i >= 0 ? name.substring(i + 1) : "in";
    t = d3_ease.get(t) || d3_ease_default;
    m = d3_ease_mode.get(m) || d3_identity;
    return d3_ease_clamp(m(t.apply(null, d3_arraySlice.call(arguments, 1))));
  };
  function d3_ease_clamp(f) {
    return function(t) {
      return t <= 0 ? 0 : t >= 1 ? 1 : f(t);
    };
  }
  function d3_ease_reverse(f) {
    return function(t) {
      return 1 - f(1 - t);
    };
  }
  function d3_ease_reflect(f) {
    return function(t) {
      return .5 * (t < .5 ? f(2 * t) : 2 - f(2 - 2 * t));
    };
  }
  function d3_ease_quad(t) {
    return t * t;
  }
  function d3_ease_cubic(t) {
    return t * t * t;
  }
  function d3_ease_cubicInOut(t) {
    if (t <= 0) return 0;
    if (t >= 1) return 1;
    var t2 = t * t, t3 = t2 * t;
    return 4 * (t < .5 ? t3 : 3 * (t - t2) + t3 - .75);
  }
  function d3_ease_poly(e) {
    return function(t) {
      return Math.pow(t, e);
    };
  }
  function d3_ease_sin(t) {
    return 1 - Math.cos(t * halfπ);
  }
  function d3_ease_exp(t) {
    return Math.pow(2, 10 * (t - 1));
  }
  function d3_ease_circle(t) {
    return 1 - Math.sqrt(1 - t * t);
  }
  function d3_ease_elastic(a, p) {
    var s;
    if (arguments.length < 2) p = .45;
    if (arguments.length) s = p / τ * Math.asin(1 / a); else a = 1, s = p / 4;
    return function(t) {
      return 1 + a * Math.pow(2, -10 * t) * Math.sin((t - s) * τ / p);
    };
  }
  function d3_ease_back(s) {
    if (!s) s = 1.70158;
    return function(t) {
      return t * t * ((s + 1) * t - s);
    };
  }
  function d3_ease_bounce(t) {
    return t < 1 / 2.75 ? 7.5625 * t * t : t < 2 / 2.75 ? 7.5625 * (t -= 1.5 / 2.75) * t + .75 : t < 2.5 / 2.75 ? 7.5625 * (t -= 2.25 / 2.75) * t + .9375 : 7.5625 * (t -= 2.625 / 2.75) * t + .984375;
  }
  d3.interpolateHcl = d3_interpolateHcl;
  function d3_interpolateHcl(a, b) {
    a = d3.hcl(a);
    b = d3.hcl(b);
    var ah = a.h, ac = a.c, al = a.l, bh = b.h - ah, bc = b.c - ac, bl = b.l - al;
    if (isNaN(bc)) bc = 0, ac = isNaN(ac) ? b.c : ac;
    if (isNaN(bh)) bh = 0, ah = isNaN(ah) ? b.h : ah; else if (bh > 180) bh -= 360; else if (bh < -180) bh += 360;
    return function(t) {
      return d3_hcl_lab(ah + bh * t, ac + bc * t, al + bl * t) + "";
    };
  }
  d3.interpolateHsl = d3_interpolateHsl;
  function d3_interpolateHsl(a, b) {
    a = d3.hsl(a);
    b = d3.hsl(b);
    var ah = a.h, as = a.s, al = a.l, bh = b.h - ah, bs = b.s - as, bl = b.l - al;
    if (isNaN(bs)) bs = 0, as = isNaN(as) ? b.s : as;
    if (isNaN(bh)) bh = 0, ah = isNaN(ah) ? b.h : ah; else if (bh > 180) bh -= 360; else if (bh < -180) bh += 360;
    return function(t) {
      return d3_hsl_rgb(ah + bh * t, as + bs * t, al + bl * t) + "";
    };
  }
  d3.interpolateLab = d3_interpolateLab;
  function d3_interpolateLab(a, b) {
    a = d3.lab(a);
    b = d3.lab(b);
    var al = a.l, aa = a.a, ab = a.b, bl = b.l - al, ba = b.a - aa, bb = b.b - ab;
    return function(t) {
      return d3_lab_rgb(al + bl * t, aa + ba * t, ab + bb * t) + "";
    };
  }
  d3.interpolateRound = d3_interpolateRound;
  function d3_interpolateRound(a, b) {
    b -= a;
    return function(t) {
      return Math.round(a + b * t);
    };
  }
  d3.transform = function(string) {
    var g = d3_document.createElementNS(d3.ns.prefix.svg, "g");
    return (d3.transform = function(string) {
      if (string != null) {
        g.setAttribute("transform", string);
        var t = g.transform.baseVal.consolidate();
      }
      return new d3_transform(t ? t.matrix : d3_transformIdentity);
    })(string);
  };
  function d3_transform(m) {
    var r0 = [ m.a, m.b ], r1 = [ m.c, m.d ], kx = d3_transformNormalize(r0), kz = d3_transformDot(r0, r1), ky = d3_transformNormalize(d3_transformCombine(r1, r0, -kz)) || 0;
    if (r0[0] * r1[1] < r1[0] * r0[1]) {
      r0[0] *= -1;
      r0[1] *= -1;
      kx *= -1;
      kz *= -1;
    }
    this.rotate = (kx ? Math.atan2(r0[1], r0[0]) : Math.atan2(-r1[0], r1[1])) * d3_degrees;
    this.translate = [ m.e, m.f ];
    this.scale = [ kx, ky ];
    this.skew = ky ? Math.atan2(kz, ky) * d3_degrees : 0;
  }
  d3_transform.prototype.toString = function() {
    return "translate(" + this.translate + ")rotate(" + this.rotate + ")skewX(" + this.skew + ")scale(" + this.scale + ")";
  };
  function d3_transformDot(a, b) {
    return a[0] * b[0] + a[1] * b[1];
  }
  function d3_transformNormalize(a) {
    var k = Math.sqrt(d3_transformDot(a, a));
    if (k) {
      a[0] /= k;
      a[1] /= k;
    }
    return k;
  }
  function d3_transformCombine(a, b, k) {
    a[0] += k * b[0];
    a[1] += k * b[1];
    return a;
  }
  var d3_transformIdentity = {
    a: 1,
    b: 0,
    c: 0,
    d: 1,
    e: 0,
    f: 0
  };
  d3.interpolateTransform = d3_interpolateTransform;
  function d3_interpolateTransform(a, b) {
    var s = [], q = [], n, A = d3.transform(a), B = d3.transform(b), ta = A.translate, tb = B.translate, ra = A.rotate, rb = B.rotate, wa = A.skew, wb = B.skew, ka = A.scale, kb = B.scale;
    if (ta[0] != tb[0] || ta[1] != tb[1]) {
      s.push("translate(", null, ",", null, ")");
      q.push({
        i: 1,
        x: d3_interpolateNumber(ta[0], tb[0])
      }, {
        i: 3,
        x: d3_interpolateNumber(ta[1], tb[1])
      });
    } else if (tb[0] || tb[1]) {
      s.push("translate(" + tb + ")");
    } else {
      s.push("");
    }
    if (ra != rb) {
      if (ra - rb > 180) rb += 360; else if (rb - ra > 180) ra += 360;
      q.push({
        i: s.push(s.pop() + "rotate(", null, ")") - 2,
        x: d3_interpolateNumber(ra, rb)
      });
    } else if (rb) {
      s.push(s.pop() + "rotate(" + rb + ")");
    }
    if (wa != wb) {
      q.push({
        i: s.push(s.pop() + "skewX(", null, ")") - 2,
        x: d3_interpolateNumber(wa, wb)
      });
    } else if (wb) {
      s.push(s.pop() + "skewX(" + wb + ")");
    }
    if (ka[0] != kb[0] || ka[1] != kb[1]) {
      n = s.push(s.pop() + "scale(", null, ",", null, ")");
      q.push({
        i: n - 4,
        x: d3_interpolateNumber(ka[0], kb[0])
      }, {
        i: n - 2,
        x: d3_interpolateNumber(ka[1], kb[1])
      });
    } else if (kb[0] != 1 || kb[1] != 1) {
      s.push(s.pop() + "scale(" + kb + ")");
    }
    n = q.length;
    return function(t) {
      var i = -1, o;
      while (++i < n) s[(o = q[i]).i] = o.x(t);
      return s.join("");
    };
  }
  function d3_uninterpolateNumber(a, b) {
    b = b - (a = +a) ? 1 / (b - a) : 0;
    return function(x) {
      return (x - a) * b;
    };
  }
  function d3_uninterpolateClamp(a, b) {
    b = b - (a = +a) ? 1 / (b - a) : 0;
    return function(x) {
      return Math.max(0, Math.min(1, (x - a) * b));
    };
  }
  d3.layout = {};
  d3.layout.bundle = function() {
    return function(links) {
      var paths = [], i = -1, n = links.length;
      while (++i < n) paths.push(d3_layout_bundlePath(links[i]));
      return paths;
    };
  };
  function d3_layout_bundlePath(link) {
    var start = link.source, end = link.target, lca = d3_layout_bundleLeastCommonAncestor(start, end), points = [ start ];
    while (start !== lca) {
      start = start.parent;
      points.push(start);
    }
    var k = points.length;
    while (end !== lca) {
      points.splice(k, 0, end);
      end = end.parent;
    }
    return points;
  }
  function d3_layout_bundleAncestors(node) {
    var ancestors = [], parent = node.parent;
    while (parent != null) {
      ancestors.push(node);
      node = parent;
      parent = parent.parent;
    }
    ancestors.push(node);
    return ancestors;
  }
  function d3_layout_bundleLeastCommonAncestor(a, b) {
    if (a === b) return a;
    var aNodes = d3_layout_bundleAncestors(a), bNodes = d3_layout_bundleAncestors(b), aNode = aNodes.pop(), bNode = bNodes.pop(), sharedNode = null;
    while (aNode === bNode) {
      sharedNode = aNode;
      aNode = aNodes.pop();
      bNode = bNodes.pop();
    }
    return sharedNode;
  }
  d3.layout.chord = function() {
    var chord = {}, chords, groups, matrix, n, padding = 0, sortGroups, sortSubgroups, sortChords;
    function relayout() {
      var subgroups = {}, groupSums = [], groupIndex = d3.range(n), subgroupIndex = [], k, x, x0, i, j;
      chords = [];
      groups = [];
      k = 0, i = -1;
      while (++i < n) {
        x = 0, j = -1;
        while (++j < n) {
          x += matrix[i][j];
        }
        groupSums.push(x);
        subgroupIndex.push(d3.range(n));
        k += x;
      }
      if (sortGroups) {
        groupIndex.sort(function(a, b) {
          return sortGroups(groupSums[a], groupSums[b]);
        });
      }
      if (sortSubgroups) {
        subgroupIndex.forEach(function(d, i) {
          d.sort(function(a, b) {
            return sortSubgroups(matrix[i][a], matrix[i][b]);
          });
        });
      }
      k = (τ - padding * n) / k;
      x = 0, i = -1;
      while (++i < n) {
        x0 = x, j = -1;
        while (++j < n) {
          var di = groupIndex[i], dj = subgroupIndex[di][j], v = matrix[di][dj], a0 = x, a1 = x += v * k;
          subgroups[di + "-" + dj] = {
            index: di,
            subindex: dj,
            startAngle: a0,
            endAngle: a1,
            value: v
          };
        }
        groups[di] = {
          index: di,
          startAngle: x0,
          endAngle: x,
          value: (x - x0) / k
        };
        x += padding;
      }
      i = -1;
      while (++i < n) {
        j = i - 1;
        while (++j < n) {
          var source = subgroups[i + "-" + j], target = subgroups[j + "-" + i];
          if (source.value || target.value) {
            chords.push(source.value < target.value ? {
              source: target,
              target: source
            } : {
              source: source,
              target: target
            });
          }
        }
      }
      if (sortChords) resort();
    }
    function resort() {
      chords.sort(function(a, b) {
        return sortChords((a.source.value + a.target.value) / 2, (b.source.value + b.target.value) / 2);
      });
    }
    chord.matrix = function(x) {
      if (!arguments.length) return matrix;
      n = (matrix = x) && matrix.length;
      chords = groups = null;
      return chord;
    };
    chord.padding = function(x) {
      if (!arguments.length) return padding;
      padding = x;
      chords = groups = null;
      return chord;
    };
    chord.sortGroups = function(x) {
      if (!arguments.length) return sortGroups;
      sortGroups = x;
      chords = groups = null;
      return chord;
    };
    chord.sortSubgroups = function(x) {
      if (!arguments.length) return sortSubgroups;
      sortSubgroups = x;
      chords = null;
      return chord;
    };
    chord.sortChords = function(x) {
      if (!arguments.length) return sortChords;
      sortChords = x;
      if (chords) resort();
      return chord;
    };
    chord.chords = function() {
      if (!chords) relayout();
      return chords;
    };
    chord.groups = function() {
      if (!groups) relayout();
      return groups;
    };
    return chord;
  };
  d3.layout.force = function() {
    var force = {}, event = d3.dispatch("start", "tick", "end"), size = [ 1, 1 ], drag, alpha, friction = .9, linkDistance = d3_layout_forceLinkDistance, linkStrength = d3_layout_forceLinkStrength, charge = -30, gravity = .1, theta = .8, nodes = [], links = [], distances, strengths, charges;
    function repulse(node) {
      return function(quad, x1, _, x2) {
        if (quad.point !== node) {
          var dx = quad.cx - node.x, dy = quad.cy - node.y, dn = 1 / Math.sqrt(dx * dx + dy * dy);
          if ((x2 - x1) * dn < theta) {
            var k = quad.charge * dn * dn;
            node.px -= dx * k;
            node.py -= dy * k;
            return true;
          }
          if (quad.point && isFinite(dn)) {
            var k = quad.pointCharge * dn * dn;
            node.px -= dx * k;
            node.py -= dy * k;
          }
        }
        return !quad.charge;
      };
    }
    force.tick = function() {
      if ((alpha *= .99) < .005) {
        event.end({
          type: "end",
          alpha: alpha = 0
        });
        return true;
      }
      var n = nodes.length, m = links.length, q, i, o, s, t, l, k, x, y;
      for (i = 0; i < m; ++i) {
        o = links[i];
        s = o.source;
        t = o.target;
        x = t.x - s.x;
        y = t.y - s.y;
        if (l = x * x + y * y) {
          l = alpha * strengths[i] * ((l = Math.sqrt(l)) - distances[i]) / l;
          x *= l;
          y *= l;
          t.x -= x * (k = s.weight / (t.weight + s.weight));
          t.y -= y * k;
          s.x += x * (k = 1 - k);
          s.y += y * k;
        }
      }
      if (k = alpha * gravity) {
        x = size[0] / 2;
        y = size[1] / 2;
        i = -1;
        if (k) while (++i < n) {
          o = nodes[i];
          o.x += (x - o.x) * k;
          o.y += (y - o.y) * k;
        }
      }
      if (charge) {
        d3_layout_forceAccumulate(q = d3.geom.quadtree(nodes), alpha, charges);
        i = -1;
        while (++i < n) {
          if (!(o = nodes[i]).fixed) {
            q.visit(repulse(o));
          }
        }
      }
      i = -1;
      while (++i < n) {
        o = nodes[i];
        if (o.fixed) {
          o.x = o.px;
          o.y = o.py;
        } else {
          o.x -= (o.px - (o.px = o.x)) * friction;
          o.y -= (o.py - (o.py = o.y)) * friction;
        }
      }
      event.tick({
        type: "tick",
        alpha: alpha
      });
    };
    force.nodes = function(x) {
      if (!arguments.length) return nodes;
      nodes = x;
      return force;
    };
    force.links = function(x) {
      if (!arguments.length) return links;
      links = x;
      return force;
    };
    force.size = function(x) {
      if (!arguments.length) return size;
      size = x;
      return force;
    };
    force.linkDistance = function(x) {
      if (!arguments.length) return linkDistance;
      linkDistance = typeof x === "function" ? x : +x;
      return force;
    };
    force.distance = force.linkDistance;
    force.linkStrength = function(x) {
      if (!arguments.length) return linkStrength;
      linkStrength = typeof x === "function" ? x : +x;
      return force;
    };
    force.friction = function(x) {
      if (!arguments.length) return friction;
      friction = +x;
      return force;
    };
    force.charge = function(x) {
      if (!arguments.length) return charge;
      charge = typeof x === "function" ? x : +x;
      return force;
    };
    force.gravity = function(x) {
      if (!arguments.length) return gravity;
      gravity = +x;
      return force;
    };
    force.theta = function(x) {
      if (!arguments.length) return theta;
      theta = +x;
      return force;
    };
    force.alpha = function(x) {
      if (!arguments.length) return alpha;
      x = +x;
      if (alpha) {
        if (x > 0) alpha = x; else alpha = 0;
      } else if (x > 0) {
        event.start({
          type: "start",
          alpha: alpha = x
        });
        d3.timer(force.tick);
      }
      return force;
    };
    force.start = function() {
      var i, n = nodes.length, m = links.length, w = size[0], h = size[1], neighbors, o;
      for (i = 0; i < n; ++i) {
        (o = nodes[i]).index = i;
        o.weight = 0;
      }
      for (i = 0; i < m; ++i) {
        o = links[i];
        if (typeof o.source == "number") o.source = nodes[o.source];
        if (typeof o.target == "number") o.target = nodes[o.target];
        ++o.source.weight;
        ++o.target.weight;
      }
      for (i = 0; i < n; ++i) {
        o = nodes[i];
        if (isNaN(o.x)) o.x = position("x", w);
        if (isNaN(o.y)) o.y = position("y", h);
        if (isNaN(o.px)) o.px = o.x;
        if (isNaN(o.py)) o.py = o.y;
      }
      distances = [];
      if (typeof linkDistance === "function") for (i = 0; i < m; ++i) distances[i] = +linkDistance.call(this, links[i], i); else for (i = 0; i < m; ++i) distances[i] = linkDistance;
      strengths = [];
      if (typeof linkStrength === "function") for (i = 0; i < m; ++i) strengths[i] = +linkStrength.call(this, links[i], i); else for (i = 0; i < m; ++i) strengths[i] = linkStrength;
      charges = [];
      if (typeof charge === "function") for (i = 0; i < n; ++i) charges[i] = +charge.call(this, nodes[i], i); else for (i = 0; i < n; ++i) charges[i] = charge;
      function position(dimension, size) {
        if (!neighbors) {
          neighbors = new Array(n);
          for (j = 0; j < n; ++j) {
            neighbors[j] = [];
          }
          for (j = 0; j < m; ++j) {
            var o = links[j];
            neighbors[o.source.index].push(o.target);
            neighbors[o.target.index].push(o.source);
          }
        }
        var candidates = neighbors[i], j = -1, m = candidates.length, x;
        while (++j < m) if (!isNaN(x = candidates[j][dimension])) return x;
        return Math.random() * size;
      }
      return force.resume();
    };
    force.resume = function() {
      return force.alpha(.1);
    };
    force.stop = function() {
      return force.alpha(0);
    };
    force.drag = function() {
      if (!drag) drag = d3.behavior.drag().origin(d3_identity).on("dragstart.force", d3_layout_forceDragstart).on("drag.force", dragmove).on("dragend.force", d3_layout_forceDragend);
      if (!arguments.length) return drag;
      this.on("mouseover.force", d3_layout_forceMouseover).on("mouseout.force", d3_layout_forceMouseout).call(drag);
    };
    function dragmove(d) {
      d.px = d3.event.x, d.py = d3.event.y;
      force.resume();
    }
    return d3.rebind(force, event, "on");
  };
  function d3_layout_forceDragstart(d) {
    d.fixed |= 2;
  }
  function d3_layout_forceDragend(d) {
    d.fixed &= ~6;
  }
  function d3_layout_forceMouseover(d) {
    d.fixed |= 4;
    d.px = d.x, d.py = d.y;
  }
  function d3_layout_forceMouseout(d) {
    d.fixed &= ~4;
  }
  function d3_layout_forceAccumulate(quad, alpha, charges) {
    var cx = 0, cy = 0;
    quad.charge = 0;
    if (!quad.leaf) {
      var nodes = quad.nodes, n = nodes.length, i = -1, c;
      while (++i < n) {
        c = nodes[i];
        if (c == null) continue;
        d3_layout_forceAccumulate(c, alpha, charges);
        quad.charge += c.charge;
        cx += c.charge * c.cx;
        cy += c.charge * c.cy;
      }
    }
    if (quad.point) {
      if (!quad.leaf) {
        quad.point.x += Math.random() - .5;
        quad.point.y += Math.random() - .5;
      }
      var k = alpha * charges[quad.point.index];
      quad.charge += quad.pointCharge = k;
      cx += k * quad.point.x;
      cy += k * quad.point.y;
    }
    quad.cx = cx / quad.charge;
    quad.cy = cy / quad.charge;
  }
  var d3_layout_forceLinkDistance = 20, d3_layout_forceLinkStrength = 1;
  d3.layout.hierarchy = function() {
    var sort = d3_layout_hierarchySort, children = d3_layout_hierarchyChildren, value = d3_layout_hierarchyValue;
    function recurse(node, depth, nodes) {
      var childs = children.call(hierarchy, node, depth);
      node.depth = depth;
      nodes.push(node);
      if (childs && (n = childs.length)) {
        var i = -1, n, c = node.children = new Array(n), v = 0, j = depth + 1, d;
        while (++i < n) {
          d = c[i] = recurse(childs[i], j, nodes);
          d.parent = node;
          v += d.value;
        }
        if (sort) c.sort(sort);
        if (value) node.value = v;
      } else {
        delete node.children;
        if (value) {
          node.value = +value.call(hierarchy, node, depth) || 0;
        }
      }
      return node;
    }
    function revalue(node, depth) {
      var children = node.children, v = 0;
      if (children && (n = children.length)) {
        var i = -1, n, j = depth + 1;
        while (++i < n) v += revalue(children[i], j);
      } else if (value) {
        v = +value.call(hierarchy, node, depth) || 0;
      }
      if (value) node.value = v;
      return v;
    }
    function hierarchy(d) {
      var nodes = [];
      recurse(d, 0, nodes);
      return nodes;
    }
    hierarchy.sort = function(x) {
      if (!arguments.length) return sort;
      sort = x;
      return hierarchy;
    };
    hierarchy.children = function(x) {
      if (!arguments.length) return children;
      children = x;
      return hierarchy;
    };
    hierarchy.value = function(x) {
      if (!arguments.length) return value;
      value = x;
      return hierarchy;
    };
    hierarchy.revalue = function(root) {
      revalue(root, 0);
      return root;
    };
    return hierarchy;
  };
  function d3_layout_hierarchyRebind(object, hierarchy) {
    d3.rebind(object, hierarchy, "sort", "children", "value");
    object.nodes = object;
    object.links = d3_layout_hierarchyLinks;
    return object;
  }
  function d3_layout_hierarchyChildren(d) {
    return d.children;
  }
  function d3_layout_hierarchyValue(d) {
    return d.value;
  }
  function d3_layout_hierarchySort(a, b) {
    return b.value - a.value;
  }
  function d3_layout_hierarchyLinks(nodes) {
    return d3.merge(nodes.map(function(parent) {
      return (parent.children || []).map(function(child) {
        return {
          source: parent,
          target: child
        };
      });
    }));
  }
  d3.layout.partition = function() {
    var hierarchy = d3.layout.hierarchy(), size = [ 1, 1 ];
    function position(node, x, dx, dy) {
      var children = node.children;
      node.x = x;
      node.y = node.depth * dy;
      node.dx = dx;
      node.dy = dy;
      if (children && (n = children.length)) {
        var i = -1, n, c, d;
        dx = node.value ? dx / node.value : 0;
        while (++i < n) {
          position(c = children[i], x, d = c.value * dx, dy);
          x += d;
        }
      }
    }
    function depth(node) {
      var children = node.children, d = 0;
      if (children && (n = children.length)) {
        var i = -1, n;
        while (++i < n) d = Math.max(d, depth(children[i]));
      }
      return 1 + d;
    }
    function partition(d, i) {
      var nodes = hierarchy.call(this, d, i);
      position(nodes[0], 0, size[0], size[1] / depth(nodes[0]));
      return nodes;
    }
    partition.size = function(x) {
      if (!arguments.length) return size;
      size = x;
      return partition;
    };
    return d3_layout_hierarchyRebind(partition, hierarchy);
  };
  d3.layout.pie = function() {
    var value = Number, sort = d3_layout_pieSortByValue, startAngle = 0, endAngle = τ;
    function pie(data) {
      var values = data.map(function(d, i) {
        return +value.call(pie, d, i);
      });
      var a = +(typeof startAngle === "function" ? startAngle.apply(this, arguments) : startAngle);
      var k = ((typeof endAngle === "function" ? endAngle.apply(this, arguments) : endAngle) - a) / d3.sum(values);
      var index = d3.range(data.length);
      if (sort != null) index.sort(sort === d3_layout_pieSortByValue ? function(i, j) {
        return values[j] - values[i];
      } : function(i, j) {
        return sort(data[i], data[j]);
      });
      var arcs = [];
      index.forEach(function(i) {
        var d;
        arcs[i] = {
          data: data[i],
          value: d = values[i],
          startAngle: a,
          endAngle: a += d * k
        };
      });
      return arcs;
    }
    pie.value = function(x) {
      if (!arguments.length) return value;
      value = x;
      return pie;
    };
    pie.sort = function(x) {
      if (!arguments.length) return sort;
      sort = x;
      return pie;
    };
    pie.startAngle = function(x) {
      if (!arguments.length) return startAngle;
      startAngle = x;
      return pie;
    };
    pie.endAngle = function(x) {
      if (!arguments.length) return endAngle;
      endAngle = x;
      return pie;
    };
    return pie;
  };
  var d3_layout_pieSortByValue = {};
  d3.layout.stack = function() {
    var values = d3_identity, order = d3_layout_stackOrderDefault, offset = d3_layout_stackOffsetZero, out = d3_layout_stackOut, x = d3_layout_stackX, y = d3_layout_stackY;
    function stack(data, index) {
      var series = data.map(function(d, i) {
        return values.call(stack, d, i);
      });
      var points = series.map(function(d) {
        return d.map(function(v, i) {
          return [ x.call(stack, v, i), y.call(stack, v, i) ];
        });
      });
      var orders = order.call(stack, points, index);
      series = d3.permute(series, orders);
      points = d3.permute(points, orders);
      var offsets = offset.call(stack, points, index);
      var n = series.length, m = series[0].length, i, j, o;
      for (j = 0; j < m; ++j) {
        out.call(stack, series[0][j], o = offsets[j], points[0][j][1]);
        for (i = 1; i < n; ++i) {
          out.call(stack, series[i][j], o += points[i - 1][j][1], points[i][j][1]);
        }
      }
      return data;
    }
    stack.values = function(x) {
      if (!arguments.length) return values;
      values = x;
      return stack;
    };
    stack.order = function(x) {
      if (!arguments.length) return order;
      order = typeof x === "function" ? x : d3_layout_stackOrders.get(x) || d3_layout_stackOrderDefault;
      return stack;
    };
    stack.offset = function(x) {
      if (!arguments.length) return offset;
      offset = typeof x === "function" ? x : d3_layout_stackOffsets.get(x) || d3_layout_stackOffsetZero;
      return stack;
    };
    stack.x = function(z) {
      if (!arguments.length) return x;
      x = z;
      return stack;
    };
    stack.y = function(z) {
      if (!arguments.length) return y;
      y = z;
      return stack;
    };
    stack.out = function(z) {
      if (!arguments.length) return out;
      out = z;
      return stack;
    };
    return stack;
  };
  function d3_layout_stackX(d) {
    return d.x;
  }
  function d3_layout_stackY(d) {
    return d.y;
  }
  function d3_layout_stackOut(d, y0, y) {
    d.y0 = y0;
    d.y = y;
  }
  var d3_layout_stackOrders = d3.map({
    "inside-out": function(data) {
      var n = data.length, i, j, max = data.map(d3_layout_stackMaxIndex), sums = data.map(d3_layout_stackReduceSum), index = d3.range(n).sort(function(a, b) {
        return max[a] - max[b];
      }), top = 0, bottom = 0, tops = [], bottoms = [];
      for (i = 0; i < n; ++i) {
        j = index[i];
        if (top < bottom) {
          top += sums[j];
          tops.push(j);
        } else {
          bottom += sums[j];
          bottoms.push(j);
        }
      }
      return bottoms.reverse().concat(tops);
    },
    reverse: function(data) {
      return d3.range(data.length).reverse();
    },
    "default": d3_layout_stackOrderDefault
  });
  var d3_layout_stackOffsets = d3.map({
    silhouette: function(data) {
      var n = data.length, m = data[0].length, sums = [], max = 0, i, j, o, y0 = [];
      for (j = 0; j < m; ++j) {
        for (i = 0, o = 0; i < n; i++) o += data[i][j][1];
        if (o > max) max = o;
        sums.push(o);
      }
      for (j = 0; j < m; ++j) {
        y0[j] = (max - sums[j]) / 2;
      }
      return y0;
    },
    wiggle: function(data) {
      var n = data.length, x = data[0], m = x.length, i, j, k, s1, s2, s3, dx, o, o0, y0 = [];
      y0[0] = o = o0 = 0;
      for (j = 1; j < m; ++j) {
        for (i = 0, s1 = 0; i < n; ++i) s1 += data[i][j][1];
        for (i = 0, s2 = 0, dx = x[j][0] - x[j - 1][0]; i < n; ++i) {
          for (k = 0, s3 = (data[i][j][1] - data[i][j - 1][1]) / (2 * dx); k < i; ++k) {
            s3 += (data[k][j][1] - data[k][j - 1][1]) / dx;
          }
          s2 += s3 * data[i][j][1];
        }
        y0[j] = o -= s1 ? s2 / s1 * dx : 0;
        if (o < o0) o0 = o;
      }
      for (j = 0; j < m; ++j) y0[j] -= o0;
      return y0;
    },
    expand: function(data) {
      var n = data.length, m = data[0].length, k = 1 / n, i, j, o, y0 = [];
      for (j = 0; j < m; ++j) {
        for (i = 0, o = 0; i < n; i++) o += data[i][j][1];
        if (o) for (i = 0; i < n; i++) data[i][j][1] /= o; else for (i = 0; i < n; i++) data[i][j][1] = k;
      }
      for (j = 0; j < m; ++j) y0[j] = 0;
      return y0;
    },
    zero: d3_layout_stackOffsetZero
  });
  function d3_layout_stackOrderDefault(data) {
    return d3.range(data.length);
  }
  function d3_layout_stackOffsetZero(data) {
    var j = -1, m = data[0].length, y0 = [];
    while (++j < m) y0[j] = 0;
    return y0;
  }
  function d3_layout_stackMaxIndex(array) {
    var i = 1, j = 0, v = array[0][1], k, n = array.length;
    for (;i < n; ++i) {
      if ((k = array[i][1]) > v) {
        j = i;
        v = k;
      }
    }
    return j;
  }
  function d3_layout_stackReduceSum(d) {
    return d.reduce(d3_layout_stackSum, 0);
  }
  function d3_layout_stackSum(p, d) {
    return p + d[1];
  }
  d3.layout.histogram = function() {
    var frequency = true, valuer = Number, ranger = d3_layout_histogramRange, binner = d3_layout_histogramBinSturges;
    function histogram(data, i) {
      var bins = [], values = data.map(valuer, this), range = ranger.call(this, values, i), thresholds = binner.call(this, range, values, i), bin, i = -1, n = values.length, m = thresholds.length - 1, k = frequency ? 1 : 1 / n, x;
      while (++i < m) {
        bin = bins[i] = [];
        bin.dx = thresholds[i + 1] - (bin.x = thresholds[i]);
        bin.y = 0;
      }
      if (m > 0) {
        i = -1;
        while (++i < n) {
          x = values[i];
          if (x >= range[0] && x <= range[1]) {
            bin = bins[d3.bisect(thresholds, x, 1, m) - 1];
            bin.y += k;
            bin.push(data[i]);
          }
        }
      }
      return bins;
    }
    histogram.value = function(x) {
      if (!arguments.length) return valuer;
      valuer = x;
      return histogram;
    };
    histogram.range = function(x) {
      if (!arguments.length) return ranger;
      ranger = d3_functor(x);
      return histogram;
    };
    histogram.bins = function(x) {
      if (!arguments.length) return binner;
      binner = typeof x === "number" ? function(range) {
        return d3_layout_histogramBinFixed(range, x);
      } : d3_functor(x);
      return histogram;
    };
    histogram.frequency = function(x) {
      if (!arguments.length) return frequency;
      frequency = !!x;
      return histogram;
    };
    return histogram;
  };
  function d3_layout_histogramBinSturges(range, values) {
    return d3_layout_histogramBinFixed(range, Math.ceil(Math.log(values.length) / Math.LN2 + 1));
  }
  function d3_layout_histogramBinFixed(range, n) {
    var x = -1, b = +range[0], m = (range[1] - b) / n, f = [];
    while (++x <= n) f[x] = m * x + b;
    return f;
  }
  function d3_layout_histogramRange(values) {
    return [ d3.min(values), d3.max(values) ];
  }
  d3.layout.tree = function() {
    var hierarchy = d3.layout.hierarchy().sort(null).value(null), separation = d3_layout_treeSeparation, size = [ 1, 1 ], nodeSize = false;
    function tree(d, i) {
      var nodes = hierarchy.call(this, d, i), root = nodes[0];
      function firstWalk(node, previousSibling) {
        var children = node.children, layout = node._tree;
        if (children && (n = children.length)) {
          var n, firstChild = children[0], previousChild, ancestor = firstChild, child, i = -1;
          while (++i < n) {
            child = children[i];
            firstWalk(child, previousChild);
            ancestor = apportion(child, previousChild, ancestor);
            previousChild = child;
          }
          d3_layout_treeShift(node);
          var midpoint = .5 * (firstChild._tree.prelim + child._tree.prelim);
          if (previousSibling) {
            layout.prelim = previousSibling._tree.prelim + separation(node, previousSibling);
            layout.mod = layout.prelim - midpoint;
          } else {
            layout.prelim = midpoint;
          }
        } else {
          if (previousSibling) {
            layout.prelim = previousSibling._tree.prelim + separation(node, previousSibling);
          }
        }
      }
      function secondWalk(node, x) {
        node.x = node._tree.prelim + x;
        var children = node.children;
        if (children && (n = children.length)) {
          var i = -1, n;
          x += node._tree.mod;
          while (++i < n) {
            secondWalk(children[i], x);
          }
        }
      }
      function apportion(node, previousSibling, ancestor) {
        if (previousSibling) {
          var vip = node, vop = node, vim = previousSibling, vom = node.parent.children[0], sip = vip._tree.mod, sop = vop._tree.mod, sim = vim._tree.mod, som = vom._tree.mod, shift;
          while (vim = d3_layout_treeRight(vim), vip = d3_layout_treeLeft(vip), vim && vip) {
            vom = d3_layout_treeLeft(vom);
            vop = d3_layout_treeRight(vop);
            vop._tree.ancestor = node;
            shift = vim._tree.prelim + sim - vip._tree.prelim - sip + separation(vim, vip);
            if (shift > 0) {
              d3_layout_treeMove(d3_layout_treeAncestor(vim, node, ancestor), node, shift);
              sip += shift;
              sop += shift;
            }
            sim += vim._tree.mod;
            sip += vip._tree.mod;
            som += vom._tree.mod;
            sop += vop._tree.mod;
          }
          if (vim && !d3_layout_treeRight(vop)) {
            vop._tree.thread = vim;
            vop._tree.mod += sim - sop;
          }
          if (vip && !d3_layout_treeLeft(vom)) {
            vom._tree.thread = vip;
            vom._tree.mod += sip - som;
            ancestor = node;
          }
        }
        return ancestor;
      }
      d3_layout_treeVisitAfter(root, function(node, previousSibling) {
        node._tree = {
          ancestor: node,
          prelim: 0,
          mod: 0,
          change: 0,
          shift: 0,
          number: previousSibling ? previousSibling._tree.number + 1 : 0
        };
      });
      firstWalk(root);
      secondWalk(root, -root._tree.prelim);
      var left = d3_layout_treeSearch(root, d3_layout_treeLeftmost), right = d3_layout_treeSearch(root, d3_layout_treeRightmost), deep = d3_layout_treeSearch(root, d3_layout_treeDeepest), x0 = left.x - separation(left, right) / 2, x1 = right.x + separation(right, left) / 2, y1 = deep.depth || 1;
      d3_layout_treeVisitAfter(root, nodeSize ? function(node) {
        node.x *= size[0];
        node.y = node.depth * size[1];
        delete node._tree;
      } : function(node) {
        node.x = (node.x - x0) / (x1 - x0) * size[0];
        node.y = node.depth / y1 * size[1];
        delete node._tree;
      });
      return nodes;
    }
    tree.separation = function(x) {
      if (!arguments.length) return separation;
      separation = x;
      return tree;
    };
    tree.size = function(x) {
      if (!arguments.length) return nodeSize ? null : size;
      nodeSize = (size = x) == null;
      return tree;
    };
    tree.nodeSize = function(x) {
      if (!arguments.length) return nodeSize ? size : null;
      nodeSize = (size = x) != null;
      return tree;
    };
    return d3_layout_hierarchyRebind(tree, hierarchy);
  };
  function d3_layout_treeSeparation(a, b) {
    return a.parent == b.parent ? 1 : 2;
  }
  function d3_layout_treeLeft(node) {
    var children = node.children;
    return children && children.length ? children[0] : node._tree.thread;
  }
  function d3_layout_treeRight(node) {
    var children = node.children, n;
    return children && (n = children.length) ? children[n - 1] : node._tree.thread;
  }
  function d3_layout_treeSearch(node, compare) {
    var children = node.children;
    if (children && (n = children.length)) {
      var child, n, i = -1;
      while (++i < n) {
        if (compare(child = d3_layout_treeSearch(children[i], compare), node) > 0) {
          node = child;
        }
      }
    }
    return node;
  }
  function d3_layout_treeRightmost(a, b) {
    return a.x - b.x;
  }
  function d3_layout_treeLeftmost(a, b) {
    return b.x - a.x;
  }
  function d3_layout_treeDeepest(a, b) {
    return a.depth - b.depth;
  }
  function d3_layout_treeVisitAfter(node, callback) {
    function visit(node, previousSibling) {
      var children = node.children;
      if (children && (n = children.length)) {
        var child, previousChild = null, i = -1, n;
        while (++i < n) {
          child = children[i];
          visit(child, previousChild);
          previousChild = child;
        }
      }
      callback(node, previousSibling);
    }
    visit(node, null);
  }
  function d3_layout_treeShift(node) {
    var shift = 0, change = 0, children = node.children, i = children.length, child;
    while (--i >= 0) {
      child = children[i]._tree;
      child.prelim += shift;
      child.mod += shift;
      shift += child.shift + (change += child.change);
    }
  }
  function d3_layout_treeMove(ancestor, node, shift) {
    ancestor = ancestor._tree;
    node = node._tree;
    var change = shift / (node.number - ancestor.number);
    ancestor.change += change;
    node.change -= change;
    node.shift += shift;
    node.prelim += shift;
    node.mod += shift;
  }
  function d3_layout_treeAncestor(vim, node, ancestor) {
    return vim._tree.ancestor.parent == node.parent ? vim._tree.ancestor : ancestor;
  }
  d3.layout.pack = function() {
    var hierarchy = d3.layout.hierarchy().sort(d3_layout_packSort), padding = 0, size = [ 1, 1 ], radius;
    function pack(d, i) {
      var nodes = hierarchy.call(this, d, i), root = nodes[0], w = size[0], h = size[1], r = radius == null ? Math.sqrt : typeof radius === "function" ? radius : function() {
        return radius;
      };
      root.x = root.y = 0;
      d3_layout_treeVisitAfter(root, function(d) {
        d.r = +r(d.value);
      });
      d3_layout_treeVisitAfter(root, d3_layout_packSiblings);
      if (padding) {
        var dr = padding * (radius ? 1 : Math.max(2 * root.r / w, 2 * root.r / h)) / 2;
        d3_layout_treeVisitAfter(root, function(d) {
          d.r += dr;
        });
        d3_layout_treeVisitAfter(root, d3_layout_packSiblings);
        d3_layout_treeVisitAfter(root, function(d) {
          d.r -= dr;
        });
      }
      d3_layout_packTransform(root, w / 2, h / 2, radius ? 1 : 1 / Math.max(2 * root.r / w, 2 * root.r / h));
      return nodes;
    }
    pack.size = function(_) {
      if (!arguments.length) return size;
      size = _;
      return pack;
    };
    pack.radius = function(_) {
      if (!arguments.length) return radius;
      radius = _ == null || typeof _ === "function" ? _ : +_;
      return pack;
    };
    pack.padding = function(_) {
      if (!arguments.length) return padding;
      padding = +_;
      return pack;
    };
    return d3_layout_hierarchyRebind(pack, hierarchy);
  };
  function d3_layout_packSort(a, b) {
    return a.value - b.value;
  }
  function d3_layout_packInsert(a, b) {
    var c = a._pack_next;
    a._pack_next = b;
    b._pack_prev = a;
    b._pack_next = c;
    c._pack_prev = b;
  }
  function d3_layout_packSplice(a, b) {
    a._pack_next = b;
    b._pack_prev = a;
  }
  function d3_layout_packIntersects(a, b) {
    var dx = b.x - a.x, dy = b.y - a.y, dr = a.r + b.r;
    return .999 * dr * dr > dx * dx + dy * dy;
  }
  function d3_layout_packSiblings(node) {
    if (!(nodes = node.children) || !(n = nodes.length)) return;
    var nodes, xMin = Infinity, xMax = -Infinity, yMin = Infinity, yMax = -Infinity, a, b, c, i, j, k, n;
    function bound(node) {
      xMin = Math.min(node.x - node.r, xMin);
      xMax = Math.max(node.x + node.r, xMax);
      yMin = Math.min(node.y - node.r, yMin);
      yMax = Math.max(node.y + node.r, yMax);
    }
    nodes.forEach(d3_layout_packLink);
    a = nodes[0];
    a.x = -a.r;
    a.y = 0;
    bound(a);
    if (n > 1) {
      b = nodes[1];
      b.x = b.r;
      b.y = 0;
      bound(b);
      if (n > 2) {
        c = nodes[2];
        d3_layout_packPlace(a, b, c);
        bound(c);
        d3_layout_packInsert(a, c);
        a._pack_prev = c;
        d3_layout_packInsert(c, b);
        b = a._pack_next;
        for (i = 3; i < n; i++) {
          d3_layout_packPlace(a, b, c = nodes[i]);
          var isect = 0, s1 = 1, s2 = 1;
          for (j = b._pack_next; j !== b; j = j._pack_next, s1++) {
            if (d3_layout_packIntersects(j, c)) {
              isect = 1;
              break;
            }
          }
          if (isect == 1) {
            for (k = a._pack_prev; k !== j._pack_prev; k = k._pack_prev, s2++) {
              if (d3_layout_packIntersects(k, c)) {
                break;
              }
            }
          }
          if (isect) {
            if (s1 < s2 || s1 == s2 && b.r < a.r) d3_layout_packSplice(a, b = j); else d3_layout_packSplice(a = k, b);
            i--;
          } else {
            d3_layout_packInsert(a, c);
            b = c;
            bound(c);
          }
        }
      }
    }
    var cx = (xMin + xMax) / 2, cy = (yMin + yMax) / 2, cr = 0;
    for (i = 0; i < n; i++) {
      c = nodes[i];
      c.x -= cx;
      c.y -= cy;
      cr = Math.max(cr, c.r + Math.sqrt(c.x * c.x + c.y * c.y));
    }
    node.r = cr;
    nodes.forEach(d3_layout_packUnlink);
  }
  function d3_layout_packLink(node) {
    node._pack_next = node._pack_prev = node;
  }
  function d3_layout_packUnlink(node) {
    delete node._pack_next;
    delete node._pack_prev;
  }
  function d3_layout_packTransform(node, x, y, k) {
    var children = node.children;
    node.x = x += k * node.x;
    node.y = y += k * node.y;
    node.r *= k;
    if (children) {
      var i = -1, n = children.length;
      while (++i < n) d3_layout_packTransform(children[i], x, y, k);
    }
  }
  function d3_layout_packPlace(a, b, c) {
    var db = a.r + c.r, dx = b.x - a.x, dy = b.y - a.y;
    if (db && (dx || dy)) {
      var da = b.r + c.r, dc = dx * dx + dy * dy;
      da *= da;
      db *= db;
      var x = .5 + (db - da) / (2 * dc), y = Math.sqrt(Math.max(0, 2 * da * (db + dc) - (db -= dc) * db - da * da)) / (2 * dc);
      c.x = a.x + x * dx + y * dy;
      c.y = a.y + x * dy - y * dx;
    } else {
      c.x = a.x + db;
      c.y = a.y;
    }
  }
  d3.layout.cluster = function() {
    var hierarchy = d3.layout.hierarchy().sort(null).value(null), separation = d3_layout_treeSeparation, size = [ 1, 1 ], nodeSize = false;
    function cluster(d, i) {
      var nodes = hierarchy.call(this, d, i), root = nodes[0], previousNode, x = 0;
      d3_layout_treeVisitAfter(root, function(node) {
        var children = node.children;
        if (children && children.length) {
          node.x = d3_layout_clusterX(children);
          node.y = d3_layout_clusterY(children);
        } else {
          node.x = previousNode ? x += separation(node, previousNode) : 0;
          node.y = 0;
          previousNode = node;
        }
      });
      var left = d3_layout_clusterLeft(root), right = d3_layout_clusterRight(root), x0 = left.x - separation(left, right) / 2, x1 = right.x + separation(right, left) / 2;
      d3_layout_treeVisitAfter(root, nodeSize ? function(node) {
        node.x = (node.x - root.x) * size[0];
        node.y = (root.y - node.y) * size[1];
      } : function(node) {
        node.x = (node.x - x0) / (x1 - x0) * size[0];
        node.y = (1 - (root.y ? node.y / root.y : 1)) * size[1];
      });
      return nodes;
    }
    cluster.separation = function(x) {
      if (!arguments.length) return separation;
      separation = x;
      return cluster;
    };
    cluster.size = function(x) {
      if (!arguments.length) return nodeSize ? null : size;
      nodeSize = (size = x) == null;
      return cluster;
    };
    cluster.nodeSize = function(x) {
      if (!arguments.length) return nodeSize ? size : null;
      nodeSize = (size = x) != null;
      return cluster;
    };
    return d3_layout_hierarchyRebind(cluster, hierarchy);
  };
  function d3_layout_clusterY(children) {
    return 1 + d3.max(children, function(child) {
      return child.y;
    });
  }
  function d3_layout_clusterX(children) {
    return children.reduce(function(x, child) {
      return x + child.x;
    }, 0) / children.length;
  }
  function d3_layout_clusterLeft(node) {
    var children = node.children;
    return children && children.length ? d3_layout_clusterLeft(children[0]) : node;
  }
  function d3_layout_clusterRight(node) {
    var children = node.children, n;
    return children && (n = children.length) ? d3_layout_clusterRight(children[n - 1]) : node;
  }
  d3.layout.treemap = function() {
    var hierarchy = d3.layout.hierarchy(), round = Math.round, size = [ 1, 1 ], padding = null, pad = d3_layout_treemapPadNull, sticky = false, stickies, mode = "squarify", ratio = .5 * (1 + Math.sqrt(5));
    function scale(children, k) {
      var i = -1, n = children.length, child, area;
      while (++i < n) {
        area = (child = children[i]).value * (k < 0 ? 0 : k);
        child.area = isNaN(area) || area <= 0 ? 0 : area;
      }
    }
    function squarify(node) {
      var children = node.children;
      if (children && children.length) {
        var rect = pad(node), row = [], remaining = children.slice(), child, best = Infinity, score, u = mode === "slice" ? rect.dx : mode === "dice" ? rect.dy : mode === "slice-dice" ? node.depth & 1 ? rect.dy : rect.dx : Math.min(rect.dx, rect.dy), n;
        scale(remaining, rect.dx * rect.dy / node.value);
        row.area = 0;
        while ((n = remaining.length) > 0) {
          row.push(child = remaining[n - 1]);
          row.area += child.area;
          if (mode !== "squarify" || (score = worst(row, u)) <= best) {
            remaining.pop();
            best = score;
          } else {
            row.area -= row.pop().area;
            position(row, u, rect, false);
            u = Math.min(rect.dx, rect.dy);
            row.length = row.area = 0;
            best = Infinity;
          }
        }
        if (row.length) {
          position(row, u, rect, true);
          row.length = row.area = 0;
        }
        children.forEach(squarify);
      }
    }
    function stickify(node) {
      var children = node.children;
      if (children && children.length) {
        var rect = pad(node), remaining = children.slice(), child, row = [];
        scale(remaining, rect.dx * rect.dy / node.value);
        row.area = 0;
        while (child = remaining.pop()) {
          row.push(child);
          row.area += child.area;
          if (child.z != null) {
            position(row, child.z ? rect.dx : rect.dy, rect, !remaining.length);
            row.length = row.area = 0;
          }
        }
        children.forEach(stickify);
      }
    }
    function worst(row, u) {
      var s = row.area, r, rmax = 0, rmin = Infinity, i = -1, n = row.length;
      while (++i < n) {
        if (!(r = row[i].area)) continue;
        if (r < rmin) rmin = r;
        if (r > rmax) rmax = r;
      }
      s *= s;
      u *= u;
      return s ? Math.max(u * rmax * ratio / s, s / (u * rmin * ratio)) : Infinity;
    }
    function position(row, u, rect, flush) {
      var i = -1, n = row.length, x = rect.x, y = rect.y, v = u ? round(row.area / u) : 0, o;
      if (u == rect.dx) {
        if (flush || v > rect.dy) v = rect.dy;
        while (++i < n) {
          o = row[i];
          o.x = x;
          o.y = y;
          o.dy = v;
          x += o.dx = Math.min(rect.x + rect.dx - x, v ? round(o.area / v) : 0);
        }
        o.z = true;
        o.dx += rect.x + rect.dx - x;
        rect.y += v;
        rect.dy -= v;
      } else {
        if (flush || v > rect.dx) v = rect.dx;
        while (++i < n) {
          o = row[i];
          o.x = x;
          o.y = y;
          o.dx = v;
          y += o.dy = Math.min(rect.y + rect.dy - y, v ? round(o.area / v) : 0);
        }
        o.z = false;
        o.dy += rect.y + rect.dy - y;
        rect.x += v;
        rect.dx -= v;
      }
    }
    function treemap(d) {
      var nodes = stickies || hierarchy(d), root = nodes[0];
      root.x = 0;
      root.y = 0;
      root.dx = size[0];
      root.dy = size[1];
      if (stickies) hierarchy.revalue(root);
      scale([ root ], root.dx * root.dy / root.value);
      (stickies ? stickify : squarify)(root);
      if (sticky) stickies = nodes;
      return nodes;
    }
    treemap.size = function(x) {
      if (!arguments.length) return size;
      size = x;
      return treemap;
    };
    treemap.padding = function(x) {
      if (!arguments.length) return padding;
      function padFunction(node) {
        var p = x.call(treemap, node, node.depth);
        return p == null ? d3_layout_treemapPadNull(node) : d3_layout_treemapPad(node, typeof p === "number" ? [ p, p, p, p ] : p);
      }
      function padConstant(node) {
        return d3_layout_treemapPad(node, x);
      }
      var type;
      pad = (padding = x) == null ? d3_layout_treemapPadNull : (type = typeof x) === "function" ? padFunction : type === "number" ? (x = [ x, x, x, x ], 
      padConstant) : padConstant;
      return treemap;
    };
    treemap.round = function(x) {
      if (!arguments.length) return round != Number;
      round = x ? Math.round : Number;
      return treemap;
    };
    treemap.sticky = function(x) {
      if (!arguments.length) return sticky;
      sticky = x;
      stickies = null;
      return treemap;
    };
    treemap.ratio = function(x) {
      if (!arguments.length) return ratio;
      ratio = x;
      return treemap;
    };
    treemap.mode = function(x) {
      if (!arguments.length) return mode;
      mode = x + "";
      return treemap;
    };
    return d3_layout_hierarchyRebind(treemap, hierarchy);
  };
  function d3_layout_treemapPadNull(node) {
    return {
      x: node.x,
      y: node.y,
      dx: node.dx,
      dy: node.dy
    };
  }
  function d3_layout_treemapPad(node, padding) {
    var x = node.x + padding[3], y = node.y + padding[0], dx = node.dx - padding[1] - padding[3], dy = node.dy - padding[0] - padding[2];
    if (dx < 0) {
      x += dx / 2;
      dx = 0;
    }
    if (dy < 0) {
      y += dy / 2;
      dy = 0;
    }
    return {
      x: x,
      y: y,
      dx: dx,
      dy: dy
    };
  }
  d3.random = {
    normal: function(µ, σ) {
      var n = arguments.length;
      if (n < 2) σ = 1;
      if (n < 1) µ = 0;
      return function() {
        var x, y, r;
        do {
          x = Math.random() * 2 - 1;
          y = Math.random() * 2 - 1;
          r = x * x + y * y;
        } while (!r || r > 1);
        return µ + σ * x * Math.sqrt(-2 * Math.log(r) / r);
      };
    },
    logNormal: function() {
      var random = d3.random.normal.apply(d3, arguments);
      return function() {
        return Math.exp(random());
      };
    },
    irwinHall: function(m) {
      return function() {
        for (var s = 0, j = 0; j < m; j++) s += Math.random();
        return s / m;
      };
    }
  };
  d3.scale = {};
  function d3_scaleExtent(domain) {
    var start = domain[0], stop = domain[domain.length - 1];
    return start < stop ? [ start, stop ] : [ stop, start ];
  }
  function d3_scaleRange(scale) {
    return scale.rangeExtent ? scale.rangeExtent() : d3_scaleExtent(scale.range());
  }
  function d3_scale_bilinear(domain, range, uninterpolate, interpolate) {
    var u = uninterpolate(domain[0], domain[1]), i = interpolate(range[0], range[1]);
    return function(x) {
      return i(u(x));
    };
  }
  function d3_scale_nice(domain, nice) {
    var i0 = 0, i1 = domain.length - 1, x0 = domain[i0], x1 = domain[i1], dx;
    if (x1 < x0) {
      dx = i0, i0 = i1, i1 = dx;
      dx = x0, x0 = x1, x1 = dx;
    }
    domain[i0] = nice.floor(x0);
    domain[i1] = nice.ceil(x1);
    return domain;
  }
  function d3_scale_niceStep(step) {
    return step ? {
      floor: function(x) {
        return Math.floor(x / step) * step;
      },
      ceil: function(x) {
        return Math.ceil(x / step) * step;
      }
    } : d3_scale_niceIdentity;
  }
  var d3_scale_niceIdentity = {
    floor: d3_identity,
    ceil: d3_identity
  };
  function d3_scale_polylinear(domain, range, uninterpolate, interpolate) {
    var u = [], i = [], j = 0, k = Math.min(domain.length, range.length) - 1;
    if (domain[k] < domain[0]) {
      domain = domain.slice().reverse();
      range = range.slice().reverse();
    }
    while (++j <= k) {
      u.push(uninterpolate(domain[j - 1], domain[j]));
      i.push(interpolate(range[j - 1], range[j]));
    }
    return function(x) {
      var j = d3.bisect(domain, x, 1, k) - 1;
      return i[j](u[j](x));
    };
  }
  d3.scale.linear = function() {
    return d3_scale_linear([ 0, 1 ], [ 0, 1 ], d3_interpolate, false);
  };
  function d3_scale_linear(domain, range, interpolate, clamp) {
    var output, input;
    function rescale() {
      var linear = Math.min(domain.length, range.length) > 2 ? d3_scale_polylinear : d3_scale_bilinear, uninterpolate = clamp ? d3_uninterpolateClamp : d3_uninterpolateNumber;
      output = linear(domain, range, uninterpolate, interpolate);
      input = linear(range, domain, uninterpolate, d3_interpolate);
      return scale;
    }
    function scale(x) {
      return output(x);
    }
    scale.invert = function(y) {
      return input(y);
    };
    scale.domain = function(x) {
      if (!arguments.length) return domain;
      domain = x.map(Number);
      return rescale();
    };
    scale.range = function(x) {
      if (!arguments.length) return range;
      range = x;
      return rescale();
    };
    scale.rangeRound = function(x) {
      return scale.range(x).interpolate(d3_interpolateRound);
    };
    scale.clamp = function(x) {
      if (!arguments.length) return clamp;
      clamp = x;
      return rescale();
    };
    scale.interpolate = function(x) {
      if (!arguments.length) return interpolate;
      interpolate = x;
      return rescale();
    };
    scale.ticks = function(m) {
      return d3_scale_linearTicks(domain, m);
    };
    scale.tickFormat = function(m, format) {
      return d3_scale_linearTickFormat(domain, m, format);
    };
    scale.nice = function(m) {
      d3_scale_linearNice(domain, m);
      return rescale();
    };
    scale.copy = function() {
      return d3_scale_linear(domain, range, interpolate, clamp);
    };
    return rescale();
  }
  function d3_scale_linearRebind(scale, linear) {
    return d3.rebind(scale, linear, "range", "rangeRound", "interpolate", "clamp");
  }
  function d3_scale_linearNice(domain, m) {
    return d3_scale_nice(domain, d3_scale_niceStep(d3_scale_linearTickRange(domain, m)[2]));
  }
  function d3_scale_linearTickRange(domain, m) {
    if (m == null) m = 10;
    var extent = d3_scaleExtent(domain), span = extent[1] - extent[0], step = Math.pow(10, Math.floor(Math.log(span / m) / Math.LN10)), err = m / span * step;
    if (err <= .15) step *= 10; else if (err <= .35) step *= 5; else if (err <= .75) step *= 2;
    extent[0] = Math.ceil(extent[0] / step) * step;
    extent[1] = Math.floor(extent[1] / step) * step + step * .5;
    extent[2] = step;
    return extent;
  }
  function d3_scale_linearTicks(domain, m) {
    return d3.range.apply(d3, d3_scale_linearTickRange(domain, m));
  }
  function d3_scale_linearTickFormat(domain, m, format) {
    var range = d3_scale_linearTickRange(domain, m);
    return d3.format(format ? format.replace(d3_format_re, function(a, b, c, d, e, f, g, h, i, j) {
      return [ b, c, d, e, f, g, h, i || "." + d3_scale_linearFormatPrecision(j, range), j ].join("");
    }) : ",." + d3_scale_linearPrecision(range[2]) + "f");
  }
  var d3_scale_linearFormatSignificant = {
    s: 1,
    g: 1,
    p: 1,
    r: 1,
    e: 1
  };
  function d3_scale_linearPrecision(value) {
    return -Math.floor(Math.log(value) / Math.LN10 + .01);
  }
  function d3_scale_linearFormatPrecision(type, range) {
    var p = d3_scale_linearPrecision(range[2]);
    return type in d3_scale_linearFormatSignificant ? Math.abs(p - d3_scale_linearPrecision(Math.max(Math.abs(range[0]), Math.abs(range[1])))) + +(type !== "e") : p - (type === "%") * 2;
  }
  d3.scale.log = function() {
    return d3_scale_log(d3.scale.linear().domain([ 0, 1 ]), 10, true, [ 1, 10 ]);
  };
  function d3_scale_log(linear, base, positive, domain) {
    function log(x) {
      return (positive ? Math.log(x < 0 ? 0 : x) : -Math.log(x > 0 ? 0 : -x)) / Math.log(base);
    }
    function pow(x) {
      return positive ? Math.pow(base, x) : -Math.pow(base, -x);
    }
    function scale(x) {
      return linear(log(x));
    }
    scale.invert = function(x) {
      return pow(linear.invert(x));
    };
    scale.domain = function(x) {
      if (!arguments.length) return domain;
      positive = x[0] >= 0;
      linear.domain((domain = x.map(Number)).map(log));
      return scale;
    };
    scale.base = function(_) {
      if (!arguments.length) return base;
      base = +_;
      linear.domain(domain.map(log));
      return scale;
    };
    scale.nice = function() {
      var niced = d3_scale_nice(domain.map(log), positive ? Math : d3_scale_logNiceNegative);
      linear.domain(niced);
      domain = niced.map(pow);
      return scale;
    };
    scale.ticks = function() {
      var extent = d3_scaleExtent(domain), ticks = [], u = extent[0], v = extent[1], i = Math.floor(log(u)), j = Math.ceil(log(v)), n = base % 1 ? 2 : base;
      if (isFinite(j - i)) {
        if (positive) {
          for (;i < j; i++) for (var k = 1; k < n; k++) ticks.push(pow(i) * k);
          ticks.push(pow(i));
        } else {
          ticks.push(pow(i));
          for (;i++ < j; ) for (var k = n - 1; k > 0; k--) ticks.push(pow(i) * k);
        }
        for (i = 0; ticks[i] < u; i++) {}
        for (j = ticks.length; ticks[j - 1] > v; j--) {}
        ticks = ticks.slice(i, j);
      }
      return ticks;
    };
    scale.tickFormat = function(n, format) {
      if (!arguments.length) return d3_scale_logFormat;
      if (arguments.length < 2) format = d3_scale_logFormat; else if (typeof format !== "function") format = d3.format(format);
      var k = Math.max(.1, n / scale.ticks().length), f = positive ? (e = 1e-12, Math.ceil) : (e = -1e-12, 
      Math.floor), e;
      return function(d) {
        return d / pow(f(log(d) + e)) <= k ? format(d) : "";
      };
    };
    scale.copy = function() {
      return d3_scale_log(linear.copy(), base, positive, domain);
    };
    return d3_scale_linearRebind(scale, linear);
  }
  var d3_scale_logFormat = d3.format(".0e"), d3_scale_logNiceNegative = {
    floor: function(x) {
      return -Math.ceil(-x);
    },
    ceil: function(x) {
      return -Math.floor(-x);
    }
  };
  d3.scale.pow = function() {
    return d3_scale_pow(d3.scale.linear(), 1, [ 0, 1 ]);
  };
  function d3_scale_pow(linear, exponent, domain) {
    var powp = d3_scale_powPow(exponent), powb = d3_scale_powPow(1 / exponent);
    function scale(x) {
      return linear(powp(x));
    }
    scale.invert = function(x) {
      return powb(linear.invert(x));
    };
    scale.domain = function(x) {
      if (!arguments.length) return domain;
      linear.domain((domain = x.map(Number)).map(powp));
      return scale;
    };
    scale.ticks = function(m) {
      return d3_scale_linearTicks(domain, m);
    };
    scale.tickFormat = function(m, format) {
      return d3_scale_linearTickFormat(domain, m, format);
    };
    scale.nice = function(m) {
      return scale.domain(d3_scale_linearNice(domain, m));
    };
    scale.exponent = function(x) {
      if (!arguments.length) return exponent;
      powp = d3_scale_powPow(exponent = x);
      powb = d3_scale_powPow(1 / exponent);
      linear.domain(domain.map(powp));
      return scale;
    };
    scale.copy = function() {
      return d3_scale_pow(linear.copy(), exponent, domain);
    };
    return d3_scale_linearRebind(scale, linear);
  }
  function d3_scale_powPow(e) {
    return function(x) {
      return x < 0 ? -Math.pow(-x, e) : Math.pow(x, e);
    };
  }
  d3.scale.sqrt = function() {
    return d3.scale.pow().exponent(.5);
  };
  d3.scale.ordinal = function() {
    return d3_scale_ordinal([], {
      t: "range",
      a: [ [] ]
    });
  };
  function d3_scale_ordinal(domain, ranger) {
    var index, range, rangeBand;
    function scale(x) {
      return range[((index.get(x) || ranger.t === "range" && index.set(x, domain.push(x))) - 1) % range.length];
    }
    function steps(start, step) {
      return d3.range(domain.length).map(function(i) {
        return start + step * i;
      });
    }
    scale.domain = function(x) {
      if (!arguments.length) return domain;
      domain = [];
      index = new d3_Map();
      var i = -1, n = x.length, xi;
      while (++i < n) if (!index.has(xi = x[i])) index.set(xi, domain.push(xi));
      return scale[ranger.t].apply(scale, ranger.a);
    };
    scale.range = function(x) {
      if (!arguments.length) return range;
      range = x;
      rangeBand = 0;
      ranger = {
        t: "range",
        a: arguments
      };
      return scale;
    };
    scale.rangePoints = function(x, padding) {
      if (arguments.length < 2) padding = 0;
      var start = x[0], stop = x[1], step = (stop - start) / (Math.max(1, domain.length - 1) + padding);
      range = steps(domain.length < 2 ? (start + stop) / 2 : start + step * padding / 2, step);
      rangeBand = 0;
      ranger = {
        t: "rangePoints",
        a: arguments
      };
      return scale;
    };
    scale.rangeBands = function(x, padding, outerPadding) {
      if (arguments.length < 2) padding = 0;
      if (arguments.length < 3) outerPadding = padding;
      var reverse = x[1] < x[0], start = x[reverse - 0], stop = x[1 - reverse], step = (stop - start) / (domain.length - padding + 2 * outerPadding);
      range = steps(start + step * outerPadding, step);
      if (reverse) range.reverse();
      rangeBand = step * (1 - padding);
      ranger = {
        t: "rangeBands",
        a: arguments
      };
      return scale;
    };
    scale.rangeRoundBands = function(x, padding, outerPadding) {
      if (arguments.length < 2) padding = 0;
      if (arguments.length < 3) outerPadding = padding;
      var reverse = x[1] < x[0], start = x[reverse - 0], stop = x[1 - reverse], step = Math.floor((stop - start) / (domain.length - padding + 2 * outerPadding)), error = stop - start - (domain.length - padding) * step;
      range = steps(start + Math.round(error / 2), step);
      if (reverse) range.reverse();
      rangeBand = Math.round(step * (1 - padding));
      ranger = {
        t: "rangeRoundBands",
        a: arguments
      };
      return scale;
    };
    scale.rangeBand = function() {
      return rangeBand;
    };
    scale.rangeExtent = function() {
      return d3_scaleExtent(ranger.a[0]);
    };
    scale.copy = function() {
      return d3_scale_ordinal(domain, ranger);
    };
    return scale.domain(domain);
  }
  d3.scale.category10 = function() {
    return d3.scale.ordinal().range(d3_category10);
  };
  d3.scale.category20 = function() {
    return d3.scale.ordinal().range(d3_category20);
  };
  d3.scale.category20b = function() {
    return d3.scale.ordinal().range(d3_category20b);
  };
  d3.scale.category20c = function() {
    return d3.scale.ordinal().range(d3_category20c);
  };
  var d3_category10 = [ 2062260, 16744206, 2924588, 14034728, 9725885, 9197131, 14907330, 8355711, 12369186, 1556175 ].map(d3_rgbString);
  var d3_category20 = [ 2062260, 11454440, 16744206, 16759672, 2924588, 10018698, 14034728, 16750742, 9725885, 12955861, 9197131, 12885140, 14907330, 16234194, 8355711, 13092807, 12369186, 14408589, 1556175, 10410725 ].map(d3_rgbString);
  var d3_category20b = [ 3750777, 5395619, 7040719, 10264286, 6519097, 9216594, 11915115, 13556636, 9202993, 12426809, 15186514, 15190932, 8666169, 11356490, 14049643, 15177372, 8077683, 10834324, 13528509, 14589654 ].map(d3_rgbString);
  var d3_category20c = [ 3244733, 7057110, 10406625, 13032431, 15095053, 16616764, 16625259, 16634018, 3253076, 7652470, 10607003, 13101504, 7695281, 10394312, 12369372, 14342891, 6513507, 9868950, 12434877, 14277081 ].map(d3_rgbString);
  d3.scale.quantile = function() {
    return d3_scale_quantile([], []);
  };
  function d3_scale_quantile(domain, range) {
    var thresholds;
    function rescale() {
      var k = 0, q = range.length;
      thresholds = [];
      while (++k < q) thresholds[k - 1] = d3.quantile(domain, k / q);
      return scale;
    }
    function scale(x) {
      if (!isNaN(x = +x)) return range[d3.bisect(thresholds, x)];
    }
    scale.domain = function(x) {
      if (!arguments.length) return domain;
      domain = x.filter(function(d) {
        return !isNaN(d);
      }).sort(d3.ascending);
      return rescale();
    };
    scale.range = function(x) {
      if (!arguments.length) return range;
      range = x;
      return rescale();
    };
    scale.quantiles = function() {
      return thresholds;
    };
    scale.invertExtent = function(y) {
      y = range.indexOf(y);
      return y < 0 ? [ NaN, NaN ] : [ y > 0 ? thresholds[y - 1] : domain[0], y < thresholds.length ? thresholds[y] : domain[domain.length - 1] ];
    };
    scale.copy = function() {
      return d3_scale_quantile(domain, range);
    };
    return rescale();
  }
  d3.scale.quantize = function() {
    return d3_scale_quantize(0, 1, [ 0, 1 ]);
  };
  function d3_scale_quantize(x0, x1, range) {
    var kx, i;
    function scale(x) {
      return range[Math.max(0, Math.min(i, Math.floor(kx * (x - x0))))];
    }
    function rescale() {
      kx = range.length / (x1 - x0);
      i = range.length - 1;
      return scale;
    }
    scale.domain = function(x) {
      if (!arguments.length) return [ x0, x1 ];
      x0 = +x[0];
      x1 = +x[x.length - 1];
      return rescale();
    };
    scale.range = function(x) {
      if (!arguments.length) return range;
      range = x;
      return rescale();
    };
    scale.invertExtent = function(y) {
      y = range.indexOf(y);
      y = y < 0 ? NaN : y / kx + x0;
      return [ y, y + 1 / kx ];
    };
    scale.copy = function() {
      return d3_scale_quantize(x0, x1, range);
    };
    return rescale();
  }
  d3.scale.threshold = function() {
    return d3_scale_threshold([ .5 ], [ 0, 1 ]);
  };
  function d3_scale_threshold(domain, range) {
    function scale(x) {
      if (x <= x) return range[d3.bisect(domain, x)];
    }
    scale.domain = function(_) {
      if (!arguments.length) return domain;
      domain = _;
      return scale;
    };
    scale.range = function(_) {
      if (!arguments.length) return range;
      range = _;
      return scale;
    };
    scale.invertExtent = function(y) {
      y = range.indexOf(y);
      return [ domain[y - 1], domain[y] ];
    };
    scale.copy = function() {
      return d3_scale_threshold(domain, range);
    };
    return scale;
  }
  d3.scale.identity = function() {
    return d3_scale_identity([ 0, 1 ]);
  };
  function d3_scale_identity(domain) {
    function identity(x) {
      return +x;
    }
    identity.invert = identity;
    identity.domain = identity.range = function(x) {
      if (!arguments.length) return domain;
      domain = x.map(identity);
      return identity;
    };
    identity.ticks = function(m) {
      return d3_scale_linearTicks(domain, m);
    };
    identity.tickFormat = function(m, format) {
      return d3_scale_linearTickFormat(domain, m, format);
    };
    identity.copy = function() {
      return d3_scale_identity(domain);
    };
    return identity;
  }
  d3.svg = {};
  d3.svg.arc = function() {
    var innerRadius = d3_svg_arcInnerRadius, outerRadius = d3_svg_arcOuterRadius, startAngle = d3_svg_arcStartAngle, endAngle = d3_svg_arcEndAngle;
    function arc() {
      var r0 = innerRadius.apply(this, arguments), r1 = outerRadius.apply(this, arguments), a0 = startAngle.apply(this, arguments) + d3_svg_arcOffset, a1 = endAngle.apply(this, arguments) + d3_svg_arcOffset, da = (a1 < a0 && (da = a0, 
      a0 = a1, a1 = da), a1 - a0), df = da < π ? "0" : "1", c0 = Math.cos(a0), s0 = Math.sin(a0), c1 = Math.cos(a1), s1 = Math.sin(a1);
      return da >= d3_svg_arcMax ? r0 ? "M0," + r1 + "A" + r1 + "," + r1 + " 0 1,1 0," + -r1 + "A" + r1 + "," + r1 + " 0 1,1 0," + r1 + "M0," + r0 + "A" + r0 + "," + r0 + " 0 1,0 0," + -r0 + "A" + r0 + "," + r0 + " 0 1,0 0," + r0 + "Z" : "M0," + r1 + "A" + r1 + "," + r1 + " 0 1,1 0," + -r1 + "A" + r1 + "," + r1 + " 0 1,1 0," + r1 + "Z" : r0 ? "M" + r1 * c0 + "," + r1 * s0 + "A" + r1 + "," + r1 + " 0 " + df + ",1 " + r1 * c1 + "," + r1 * s1 + "L" + r0 * c1 + "," + r0 * s1 + "A" + r0 + "," + r0 + " 0 " + df + ",0 " + r0 * c0 + "," + r0 * s0 + "Z" : "M" + r1 * c0 + "," + r1 * s0 + "A" + r1 + "," + r1 + " 0 " + df + ",1 " + r1 * c1 + "," + r1 * s1 + "L0,0" + "Z";
    }
    arc.innerRadius = function(v) {
      if (!arguments.length) return innerRadius;
      innerRadius = d3_functor(v);
      return arc;
    };
    arc.outerRadius = function(v) {
      if (!arguments.length) return outerRadius;
      outerRadius = d3_functor(v);
      return arc;
    };
    arc.startAngle = function(v) {
      if (!arguments.length) return startAngle;
      startAngle = d3_functor(v);
      return arc;
    };
    arc.endAngle = function(v) {
      if (!arguments.length) return endAngle;
      endAngle = d3_functor(v);
      return arc;
    };
    arc.centroid = function() {
      var r = (innerRadius.apply(this, arguments) + outerRadius.apply(this, arguments)) / 2, a = (startAngle.apply(this, arguments) + endAngle.apply(this, arguments)) / 2 + d3_svg_arcOffset;
      return [ Math.cos(a) * r, Math.sin(a) * r ];
    };
    return arc;
  };
  var d3_svg_arcOffset = -halfπ, d3_svg_arcMax = τ - ε;
  function d3_svg_arcInnerRadius(d) {
    return d.innerRadius;
  }
  function d3_svg_arcOuterRadius(d) {
    return d.outerRadius;
  }
  function d3_svg_arcStartAngle(d) {
    return d.startAngle;
  }
  function d3_svg_arcEndAngle(d) {
    return d.endAngle;
  }
  function d3_svg_line(projection) {
    var x = d3_geom_pointX, y = d3_geom_pointY, defined = d3_true, interpolate = d3_svg_lineLinear, interpolateKey = interpolate.key, tension = .7;
    function line(data) {
      var segments = [], points = [], i = -1, n = data.length, d, fx = d3_functor(x), fy = d3_functor(y);
      function segment() {
        segments.push("M", interpolate(projection(points), tension));
      }
      while (++i < n) {
        if (defined.call(this, d = data[i], i)) {
          points.push([ +fx.call(this, d, i), +fy.call(this, d, i) ]);
        } else if (points.length) {
          segment();
          points = [];
        }
      }
      if (points.length) segment();
      return segments.length ? segments.join("") : null;
    }
    line.x = function(_) {
      if (!arguments.length) return x;
      x = _;
      return line;
    };
    line.y = function(_) {
      if (!arguments.length) return y;
      y = _;
      return line;
    };
    line.defined = function(_) {
      if (!arguments.length) return defined;
      defined = _;
      return line;
    };
    line.interpolate = function(_) {
      if (!arguments.length) return interpolateKey;
      if (typeof _ === "function") interpolateKey = interpolate = _; else interpolateKey = (interpolate = d3_svg_lineInterpolators.get(_) || d3_svg_lineLinear).key;
      return line;
    };
    line.tension = function(_) {
      if (!arguments.length) return tension;
      tension = _;
      return line;
    };
    return line;
  }
  d3.svg.line = function() {
    return d3_svg_line(d3_identity);
  };
  var d3_svg_lineInterpolators = d3.map({
    linear: d3_svg_lineLinear,
    "linear-closed": d3_svg_lineLinearClosed,
    step: d3_svg_lineStep,
    "step-before": d3_svg_lineStepBefore,
    "step-after": d3_svg_lineStepAfter,
    basis: d3_svg_lineBasis,
    "basis-open": d3_svg_lineBasisOpen,
    "basis-closed": d3_svg_lineBasisClosed,
    bundle: d3_svg_lineBundle,
    cardinal: d3_svg_lineCardinal,
    "cardinal-open": d3_svg_lineCardinalOpen,
    "cardinal-closed": d3_svg_lineCardinalClosed,
    monotone: d3_svg_lineMonotone
  });
  d3_svg_lineInterpolators.forEach(function(key, value) {
    value.key = key;
    value.closed = /-closed$/.test(key);
  });
  function d3_svg_lineLinear(points) {
    return points.join("L");
  }
  function d3_svg_lineLinearClosed(points) {
    return d3_svg_lineLinear(points) + "Z";
  }
  function d3_svg_lineStep(points) {
    var i = 0, n = points.length, p = points[0], path = [ p[0], ",", p[1] ];
    while (++i < n) path.push("H", (p[0] + (p = points[i])[0]) / 2, "V", p[1]);
    if (n > 1) path.push("H", p[0]);
    return path.join("");
  }
  function d3_svg_lineStepBefore(points) {
    var i = 0, n = points.length, p = points[0], path = [ p[0], ",", p[1] ];
    while (++i < n) path.push("V", (p = points[i])[1], "H", p[0]);
    return path.join("");
  }
  function d3_svg_lineStepAfter(points) {
    var i = 0, n = points.length, p = points[0], path = [ p[0], ",", p[1] ];
    while (++i < n) path.push("H", (p = points[i])[0], "V", p[1]);
    return path.join("");
  }
  function d3_svg_lineCardinalOpen(points, tension) {
    return points.length < 4 ? d3_svg_lineLinear(points) : points[1] + d3_svg_lineHermite(points.slice(1, points.length - 1), d3_svg_lineCardinalTangents(points, tension));
  }
  function d3_svg_lineCardinalClosed(points, tension) {
    return points.length < 3 ? d3_svg_lineLinear(points) : points[0] + d3_svg_lineHermite((points.push(points[0]), 
    points), d3_svg_lineCardinalTangents([ points[points.length - 2] ].concat(points, [ points[1] ]), tension));
  }
  function d3_svg_lineCardinal(points, tension) {
    return points.length < 3 ? d3_svg_lineLinear(points) : points[0] + d3_svg_lineHermite(points, d3_svg_lineCardinalTangents(points, tension));
  }
  function d3_svg_lineHermite(points, tangents) {
    if (tangents.length < 1 || points.length != tangents.length && points.length != tangents.length + 2) {
      return d3_svg_lineLinear(points);
    }
    var quad = points.length != tangents.length, path = "", p0 = points[0], p = points[1], t0 = tangents[0], t = t0, pi = 1;
    if (quad) {
      path += "Q" + (p[0] - t0[0] * 2 / 3) + "," + (p[1] - t0[1] * 2 / 3) + "," + p[0] + "," + p[1];
      p0 = points[1];
      pi = 2;
    }
    if (tangents.length > 1) {
      t = tangents[1];
      p = points[pi];
      pi++;
      path += "C" + (p0[0] + t0[0]) + "," + (p0[1] + t0[1]) + "," + (p[0] - t[0]) + "," + (p[1] - t[1]) + "," + p[0] + "," + p[1];
      for (var i = 2; i < tangents.length; i++, pi++) {
        p = points[pi];
        t = tangents[i];
        path += "S" + (p[0] - t[0]) + "," + (p[1] - t[1]) + "," + p[0] + "," + p[1];
      }
    }
    if (quad) {
      var lp = points[pi];
      path += "Q" + (p[0] + t[0] * 2 / 3) + "," + (p[1] + t[1] * 2 / 3) + "," + lp[0] + "," + lp[1];
    }
    return path;
  }
  function d3_svg_lineCardinalTangents(points, tension) {
    var tangents = [], a = (1 - tension) / 2, p0, p1 = points[0], p2 = points[1], i = 1, n = points.length;
    while (++i < n) {
      p0 = p1;
      p1 = p2;
      p2 = points[i];
      tangents.push([ a * (p2[0] - p0[0]), a * (p2[1] - p0[1]) ]);
    }
    return tangents;
  }
  function d3_svg_lineBasis(points) {
    if (points.length < 3) return d3_svg_lineLinear(points);
    var i = 1, n = points.length, pi = points[0], x0 = pi[0], y0 = pi[1], px = [ x0, x0, x0, (pi = points[1])[0] ], py = [ y0, y0, y0, pi[1] ], path = [ x0, ",", y0, "L", d3_svg_lineDot4(d3_svg_lineBasisBezier3, px), ",", d3_svg_lineDot4(d3_svg_lineBasisBezier3, py) ];
    points.push(points[n - 1]);
    while (++i <= n) {
      pi = points[i];
      px.shift();
      px.push(pi[0]);
      py.shift();
      py.push(pi[1]);
      d3_svg_lineBasisBezier(path, px, py);
    }
    points.pop();
    path.push("L", pi);
    return path.join("");
  }
  function d3_svg_lineBasisOpen(points) {
    if (points.length < 4) return d3_svg_lineLinear(points);
    var path = [], i = -1, n = points.length, pi, px = [ 0 ], py = [ 0 ];
    while (++i < 3) {
      pi = points[i];
      px.push(pi[0]);
      py.push(pi[1]);
    }
    path.push(d3_svg_lineDot4(d3_svg_lineBasisBezier3, px) + "," + d3_svg_lineDot4(d3_svg_lineBasisBezier3, py));
    --i;
    while (++i < n) {
      pi = points[i];
      px.shift();
      px.push(pi[0]);
      py.shift();
      py.push(pi[1]);
      d3_svg_lineBasisBezier(path, px, py);
    }
    return path.join("");
  }
  function d3_svg_lineBasisClosed(points) {
    var path, i = -1, n = points.length, m = n + 4, pi, px = [], py = [];
    while (++i < 4) {
      pi = points[i % n];
      px.push(pi[0]);
      py.push(pi[1]);
    }
    path = [ d3_svg_lineDot4(d3_svg_lineBasisBezier3, px), ",", d3_svg_lineDot4(d3_svg_lineBasisBezier3, py) ];
    --i;
    while (++i < m) {
      pi = points[i % n];
      px.shift();
      px.push(pi[0]);
      py.shift();
      py.push(pi[1]);
      d3_svg_lineBasisBezier(path, px, py);
    }
    return path.join("");
  }
  function d3_svg_lineBundle(points, tension) {
    var n = points.length - 1;
    if (n) {
      var x0 = points[0][0], y0 = points[0][1], dx = points[n][0] - x0, dy = points[n][1] - y0, i = -1, p, t;
      while (++i <= n) {
        p = points[i];
        t = i / n;
        p[0] = tension * p[0] + (1 - tension) * (x0 + t * dx);
        p[1] = tension * p[1] + (1 - tension) * (y0 + t * dy);
      }
    }
    return d3_svg_lineBasis(points);
  }
  function d3_svg_lineDot4(a, b) {
    return a[0] * b[0] + a[1] * b[1] + a[2] * b[2] + a[3] * b[3];
  }
  var d3_svg_lineBasisBezier1 = [ 0, 2 / 3, 1 / 3, 0 ], d3_svg_lineBasisBezier2 = [ 0, 1 / 3, 2 / 3, 0 ], d3_svg_lineBasisBezier3 = [ 0, 1 / 6, 2 / 3, 1 / 6 ];
  function d3_svg_lineBasisBezier(path, x, y) {
    path.push("C", d3_svg_lineDot4(d3_svg_lineBasisBezier1, x), ",", d3_svg_lineDot4(d3_svg_lineBasisBezier1, y), ",", d3_svg_lineDot4(d3_svg_lineBasisBezier2, x), ",", d3_svg_lineDot4(d3_svg_lineBasisBezier2, y), ",", d3_svg_lineDot4(d3_svg_lineBasisBezier3, x), ",", d3_svg_lineDot4(d3_svg_lineBasisBezier3, y));
  }
  function d3_svg_lineSlope(p0, p1) {
    return (p1[1] - p0[1]) / (p1[0] - p0[0]);
  }
  function d3_svg_lineFiniteDifferences(points) {
    var i = 0, j = points.length - 1, m = [], p0 = points[0], p1 = points[1], d = m[0] = d3_svg_lineSlope(p0, p1);
    while (++i < j) {
      m[i] = (d + (d = d3_svg_lineSlope(p0 = p1, p1 = points[i + 1]))) / 2;
    }
    m[i] = d;
    return m;
  }
  function d3_svg_lineMonotoneTangents(points) {
    var tangents = [], d, a, b, s, m = d3_svg_lineFiniteDifferences(points), i = -1, j = points.length - 1;
    while (++i < j) {
      d = d3_svg_lineSlope(points[i], points[i + 1]);
      if (abs(d) < ε) {
        m[i] = m[i + 1] = 0;
      } else {
        a = m[i] / d;
        b = m[i + 1] / d;
        s = a * a + b * b;
        if (s > 9) {
          s = d * 3 / Math.sqrt(s);
          m[i] = s * a;
          m[i + 1] = s * b;
        }
      }
    }
    i = -1;
    while (++i <= j) {
      s = (points[Math.min(j, i + 1)][0] - points[Math.max(0, i - 1)][0]) / (6 * (1 + m[i] * m[i]));
      tangents.push([ s || 0, m[i] * s || 0 ]);
    }
    return tangents;
  }
  function d3_svg_lineMonotone(points) {
    return points.length < 3 ? d3_svg_lineLinear(points) : points[0] + d3_svg_lineHermite(points, d3_svg_lineMonotoneTangents(points));
  }
  d3.svg.line.radial = function() {
    var line = d3_svg_line(d3_svg_lineRadial);
    line.radius = line.x, delete line.x;
    line.angle = line.y, delete line.y;
    return line;
  };
  function d3_svg_lineRadial(points) {
    var point, i = -1, n = points.length, r, a;
    while (++i < n) {
      point = points[i];
      r = point[0];
      a = point[1] + d3_svg_arcOffset;
      point[0] = r * Math.cos(a);
      point[1] = r * Math.sin(a);
    }
    return points;
  }
  function d3_svg_area(projection) {
    var x0 = d3_geom_pointX, x1 = d3_geom_pointX, y0 = 0, y1 = d3_geom_pointY, defined = d3_true, interpolate = d3_svg_lineLinear, interpolateKey = interpolate.key, interpolateReverse = interpolate, L = "L", tension = .7;
    function area(data) {
      var segments = [], points0 = [], points1 = [], i = -1, n = data.length, d, fx0 = d3_functor(x0), fy0 = d3_functor(y0), fx1 = x0 === x1 ? function() {
        return x;
      } : d3_functor(x1), fy1 = y0 === y1 ? function() {
        return y;
      } : d3_functor(y1), x, y;
      function segment() {
        segments.push("M", interpolate(projection(points1), tension), L, interpolateReverse(projection(points0.reverse()), tension), "Z");
      }
      while (++i < n) {
        if (defined.call(this, d = data[i], i)) {
          points0.push([ x = +fx0.call(this, d, i), y = +fy0.call(this, d, i) ]);
          points1.push([ +fx1.call(this, d, i), +fy1.call(this, d, i) ]);
        } else if (points0.length) {
          segment();
          points0 = [];
          points1 = [];
        }
      }
      if (points0.length) segment();
      return segments.length ? segments.join("") : null;
    }
    area.x = function(_) {
      if (!arguments.length) return x1;
      x0 = x1 = _;
      return area;
    };
    area.x0 = function(_) {
      if (!arguments.length) return x0;
      x0 = _;
      return area;
    };
    area.x1 = function(_) {
      if (!arguments.length) return x1;
      x1 = _;
      return area;
    };
    area.y = function(_) {
      if (!arguments.length) return y1;
      y0 = y1 = _;
      return area;
    };
    area.y0 = function(_) {
      if (!arguments.length) return y0;
      y0 = _;
      return area;
    };
    area.y1 = function(_) {
      if (!arguments.length) return y1;
      y1 = _;
      return area;
    };
    area.defined = function(_) {
      if (!arguments.length) return defined;
      defined = _;
      return area;
    };
    area.interpolate = function(_) {
      if (!arguments.length) return interpolateKey;
      if (typeof _ === "function") interpolateKey = interpolate = _; else interpolateKey = (interpolate = d3_svg_lineInterpolators.get(_) || d3_svg_lineLinear).key;
      interpolateReverse = interpolate.reverse || interpolate;
      L = interpolate.closed ? "M" : "L";
      return area;
    };
    area.tension = function(_) {
      if (!arguments.length) return tension;
      tension = _;
      return area;
    };
    return area;
  }
  d3_svg_lineStepBefore.reverse = d3_svg_lineStepAfter;
  d3_svg_lineStepAfter.reverse = d3_svg_lineStepBefore;
  d3.svg.area = function() {
    return d3_svg_area(d3_identity);
  };
  d3.svg.area.radial = function() {
    var area = d3_svg_area(d3_svg_lineRadial);
    area.radius = area.x, delete area.x;
    area.innerRadius = area.x0, delete area.x0;
    area.outerRadius = area.x1, delete area.x1;
    area.angle = area.y, delete area.y;
    area.startAngle = area.y0, delete area.y0;
    area.endAngle = area.y1, delete area.y1;
    return area;
  };
  d3.svg.chord = function() {
    var source = d3_source, target = d3_target, radius = d3_svg_chordRadius, startAngle = d3_svg_arcStartAngle, endAngle = d3_svg_arcEndAngle;
    function chord(d, i) {
      var s = subgroup(this, source, d, i), t = subgroup(this, target, d, i);
      return "M" + s.p0 + arc(s.r, s.p1, s.a1 - s.a0) + (equals(s, t) ? curve(s.r, s.p1, s.r, s.p0) : curve(s.r, s.p1, t.r, t.p0) + arc(t.r, t.p1, t.a1 - t.a0) + curve(t.r, t.p1, s.r, s.p0)) + "Z";
    }
    function subgroup(self, f, d, i) {
      var subgroup = f.call(self, d, i), r = radius.call(self, subgroup, i), a0 = startAngle.call(self, subgroup, i) + d3_svg_arcOffset, a1 = endAngle.call(self, subgroup, i) + d3_svg_arcOffset;
      return {
        r: r,
        a0: a0,
        a1: a1,
        p0: [ r * Math.cos(a0), r * Math.sin(a0) ],
        p1: [ r * Math.cos(a1), r * Math.sin(a1) ]
      };
    }
    function equals(a, b) {
      return a.a0 == b.a0 && a.a1 == b.a1;
    }
    function arc(r, p, a) {
      return "A" + r + "," + r + " 0 " + +(a > π) + ",1 " + p;
    }
    function curve(r0, p0, r1, p1) {
      return "Q 0,0 " + p1;
    }
    chord.radius = function(v) {
      if (!arguments.length) return radius;
      radius = d3_functor(v);
      return chord;
    };
    chord.source = function(v) {
      if (!arguments.length) return source;
      source = d3_functor(v);
      return chord;
    };
    chord.target = function(v) {
      if (!arguments.length) return target;
      target = d3_functor(v);
      return chord;
    };
    chord.startAngle = function(v) {
      if (!arguments.length) return startAngle;
      startAngle = d3_functor(v);
      return chord;
    };
    chord.endAngle = function(v) {
      if (!arguments.length) return endAngle;
      endAngle = d3_functor(v);
      return chord;
    };
    return chord;
  };
  function d3_svg_chordRadius(d) {
    return d.radius;
  }
  d3.svg.diagonal = function() {
    var source = d3_source, target = d3_target, projection = d3_svg_diagonalProjection;
    function diagonal(d, i) {
      var p0 = source.call(this, d, i), p3 = target.call(this, d, i), m = (p0.y + p3.y) / 2, p = [ p0, {
        x: p0.x,
        y: m
      }, {
        x: p3.x,
        y: m
      }, p3 ];
      p = p.map(projection);
      return "M" + p[0] + "C" + p[1] + " " + p[2] + " " + p[3];
    }
    diagonal.source = function(x) {
      if (!arguments.length) return source;
      source = d3_functor(x);
      return diagonal;
    };
    diagonal.target = function(x) {
      if (!arguments.length) return target;
      target = d3_functor(x);
      return diagonal;
    };
    diagonal.projection = function(x) {
      if (!arguments.length) return projection;
      projection = x;
      return diagonal;
    };
    return diagonal;
  };
  function d3_svg_diagonalProjection(d) {
    return [ d.x, d.y ];
  }
  d3.svg.diagonal.radial = function() {
    var diagonal = d3.svg.diagonal(), projection = d3_svg_diagonalProjection, projection_ = diagonal.projection;
    diagonal.projection = function(x) {
      return arguments.length ? projection_(d3_svg_diagonalRadialProjection(projection = x)) : projection;
    };
    return diagonal;
  };
  function d3_svg_diagonalRadialProjection(projection) {
    return function() {
      var d = projection.apply(this, arguments), r = d[0], a = d[1] + d3_svg_arcOffset;
      return [ r * Math.cos(a), r * Math.sin(a) ];
    };
  }
  d3.svg.symbol = function() {
    var type = d3_svg_symbolType, size = d3_svg_symbolSize;
    function symbol(d, i) {
      return (d3_svg_symbols.get(type.call(this, d, i)) || d3_svg_symbolCircle)(size.call(this, d, i));
    }
    symbol.type = function(x) {
      if (!arguments.length) return type;
      type = d3_functor(x);
      return symbol;
    };
    symbol.size = function(x) {
      if (!arguments.length) return size;
      size = d3_functor(x);
      return symbol;
    };
    return symbol;
  };
  function d3_svg_symbolSize() {
    return 64;
  }
  function d3_svg_symbolType() {
    return "circle";
  }
  function d3_svg_symbolCircle(size) {
    var r = Math.sqrt(size / π);
    return "M0," + r + "A" + r + "," + r + " 0 1,1 0," + -r + "A" + r + "," + r + " 0 1,1 0," + r + "Z";
  }
  var d3_svg_symbols = d3.map({
    circle: d3_svg_symbolCircle,
    cross: function(size) {
      var r = Math.sqrt(size / 5) / 2;
      return "M" + -3 * r + "," + -r + "H" + -r + "V" + -3 * r + "H" + r + "V" + -r + "H" + 3 * r + "V" + r + "H" + r + "V" + 3 * r + "H" + -r + "V" + r + "H" + -3 * r + "Z";
    },
    diamond: function(size) {
      var ry = Math.sqrt(size / (2 * d3_svg_symbolTan30)), rx = ry * d3_svg_symbolTan30;
      return "M0," + -ry + "L" + rx + ",0" + " 0," + ry + " " + -rx + ",0" + "Z";
    },
    square: function(size) {
      var r = Math.sqrt(size) / 2;
      return "M" + -r + "," + -r + "L" + r + "," + -r + " " + r + "," + r + " " + -r + "," + r + "Z";
    },
    "triangle-down": function(size) {
      var rx = Math.sqrt(size / d3_svg_symbolSqrt3), ry = rx * d3_svg_symbolSqrt3 / 2;
      return "M0," + ry + "L" + rx + "," + -ry + " " + -rx + "," + -ry + "Z";
    },
    "triangle-up": function(size) {
      var rx = Math.sqrt(size / d3_svg_symbolSqrt3), ry = rx * d3_svg_symbolSqrt3 / 2;
      return "M0," + -ry + "L" + rx + "," + ry + " " + -rx + "," + ry + "Z";
    }
  });
  d3.svg.symbolTypes = d3_svg_symbols.keys();
  var d3_svg_symbolSqrt3 = Math.sqrt(3), d3_svg_symbolTan30 = Math.tan(30 * d3_radians);
  function d3_transition(groups, id) {
    d3_subclass(groups, d3_transitionPrototype);
    groups.id = id;
    return groups;
  }
  var d3_transitionPrototype = [], d3_transitionId = 0, d3_transitionInheritId, d3_transitionInherit;
  d3_transitionPrototype.call = d3_selectionPrototype.call;
  d3_transitionPrototype.empty = d3_selectionPrototype.empty;
  d3_transitionPrototype.node = d3_selectionPrototype.node;
  d3_transitionPrototype.size = d3_selectionPrototype.size;
  d3.transition = function(selection) {
    return arguments.length ? d3_transitionInheritId ? selection.transition() : selection : d3_selectionRoot.transition();
  };
  d3.transition.prototype = d3_transitionPrototype;
  d3_transitionPrototype.select = function(selector) {
    var id = this.id, subgroups = [], subgroup, subnode, node;
    selector = d3_selection_selector(selector);
    for (var j = -1, m = this.length; ++j < m; ) {
      subgroups.push(subgroup = []);
      for (var group = this[j], i = -1, n = group.length; ++i < n; ) {
        if ((node = group[i]) && (subnode = selector.call(node, node.__data__, i, j))) {
          if ("__data__" in node) subnode.__data__ = node.__data__;
          d3_transitionNode(subnode, i, id, node.__transition__[id]);
          subgroup.push(subnode);
        } else {
          subgroup.push(null);
        }
      }
    }
    return d3_transition(subgroups, id);
  };
  d3_transitionPrototype.selectAll = function(selector) {
    var id = this.id, subgroups = [], subgroup, subnodes, node, subnode, transition;
    selector = d3_selection_selectorAll(selector);
    for (var j = -1, m = this.length; ++j < m; ) {
      for (var group = this[j], i = -1, n = group.length; ++i < n; ) {
        if (node = group[i]) {
          transition = node.__transition__[id];
          subnodes = selector.call(node, node.__data__, i, j);
          subgroups.push(subgroup = []);
          for (var k = -1, o = subnodes.length; ++k < o; ) {
            if (subnode = subnodes[k]) d3_transitionNode(subnode, k, id, transition);
            subgroup.push(subnode);
          }
        }
      }
    }
    return d3_transition(subgroups, id);
  };
  d3_transitionPrototype.filter = function(filter) {
    var subgroups = [], subgroup, group, node;
    if (typeof filter !== "function") filter = d3_selection_filter(filter);
    for (var j = 0, m = this.length; j < m; j++) {
      subgroups.push(subgroup = []);
      for (var group = this[j], i = 0, n = group.length; i < n; i++) {
        if ((node = group[i]) && filter.call(node, node.__data__, i, j)) {
          subgroup.push(node);
        }
      }
    }
    return d3_transition(subgroups, this.id);
  };
  d3_transitionPrototype.tween = function(name, tween) {
    var id = this.id;
    if (arguments.length < 2) return this.node().__transition__[id].tween.get(name);
    return d3_selection_each(this, tween == null ? function(node) {
      node.__transition__[id].tween.remove(name);
    } : function(node) {
      node.__transition__[id].tween.set(name, tween);
    });
  };
  function d3_transition_tween(groups, name, value, tween) {
    var id = groups.id;
    return d3_selection_each(groups, typeof value === "function" ? function(node, i, j) {
      node.__transition__[id].tween.set(name, tween(value.call(node, node.__data__, i, j)));
    } : (value = tween(value), function(node) {
      node.__transition__[id].tween.set(name, value);
    }));
  }
  d3_transitionPrototype.attr = function(nameNS, value) {
    if (arguments.length < 2) {
      for (value in nameNS) this.attr(value, nameNS[value]);
      return this;
    }
    var interpolate = nameNS == "transform" ? d3_interpolateTransform : d3_interpolate, name = d3.ns.qualify(nameNS);
    function attrNull() {
      this.removeAttribute(name);
    }
    function attrNullNS() {
      this.removeAttributeNS(name.space, name.local);
    }
    function attrTween(b) {
      return b == null ? attrNull : (b += "", function() {
        var a = this.getAttribute(name), i;
        return a !== b && (i = interpolate(a, b), function(t) {
          this.setAttribute(name, i(t));
        });
      });
    }
    function attrTweenNS(b) {
      return b == null ? attrNullNS : (b += "", function() {
        var a = this.getAttributeNS(name.space, name.local), i;
        return a !== b && (i = interpolate(a, b), function(t) {
          this.setAttributeNS(name.space, name.local, i(t));
        });
      });
    }
    return d3_transition_tween(this, "attr." + nameNS, value, name.local ? attrTweenNS : attrTween);
  };
  d3_transitionPrototype.attrTween = function(nameNS, tween) {
    var name = d3.ns.qualify(nameNS);
    function attrTween(d, i) {
      var f = tween.call(this, d, i, this.getAttribute(name));
      return f && function(t) {
        this.setAttribute(name, f(t));
      };
    }
    function attrTweenNS(d, i) {
      var f = tween.call(this, d, i, this.getAttributeNS(name.space, name.local));
      return f && function(t) {
        this.setAttributeNS(name.space, name.local, f(t));
      };
    }
    return this.tween("attr." + nameNS, name.local ? attrTweenNS : attrTween);
  };
  d3_transitionPrototype.style = function(name, value, priority) {
    var n = arguments.length;
    if (n < 3) {
      if (typeof name !== "string") {
        if (n < 2) value = "";
        for (priority in name) this.style(priority, name[priority], value);
        return this;
      }
      priority = "";
    }
    function styleNull() {
      this.style.removeProperty(name);
    }
    function styleString(b) {
      return b == null ? styleNull : (b += "", function() {
        var a = d3_window.getComputedStyle(this, null).getPropertyValue(name), i;
        return a !== b && (i = d3_interpolate(a, b), function(t) {
          this.style.setProperty(name, i(t), priority);
        });
      });
    }
    return d3_transition_tween(this, "style." + name, value, styleString);
  };
  d3_transitionPrototype.styleTween = function(name, tween, priority) {
    if (arguments.length < 3) priority = "";
    function styleTween(d, i) {
      var f = tween.call(this, d, i, d3_window.getComputedStyle(this, null).getPropertyValue(name));
      return f && function(t) {
        this.style.setProperty(name, f(t), priority);
      };
    }
    return this.tween("style." + name, styleTween);
  };
  d3_transitionPrototype.text = function(value) {
    return d3_transition_tween(this, "text", value, d3_transition_text);
  };
  function d3_transition_text(b) {
    if (b == null) b = "";
    return function() {
      this.textContent = b;
    };
  }
  d3_transitionPrototype.remove = function() {
    return this.each("end.transition", function() {
      var p;
      if (this.__transition__.count < 2 && (p = this.parentNode)) p.removeChild(this);
    });
  };
  d3_transitionPrototype.ease = function(value) {
    var id = this.id;
    if (arguments.length < 1) return this.node().__transition__[id].ease;
    if (typeof value !== "function") value = d3.ease.apply(d3, arguments);
    return d3_selection_each(this, function(node) {
      node.__transition__[id].ease = value;
    });
  };
  d3_transitionPrototype.delay = function(value) {
    var id = this.id;
    return d3_selection_each(this, typeof value === "function" ? function(node, i, j) {
      node.__transition__[id].delay = +value.call(node, node.__data__, i, j);
    } : (value = +value, function(node) {
      node.__transition__[id].delay = value;
    }));
  };
  d3_transitionPrototype.duration = function(value) {
    var id = this.id;
    return d3_selection_each(this, typeof value === "function" ? function(node, i, j) {
      node.__transition__[id].duration = Math.max(1, value.call(node, node.__data__, i, j));
    } : (value = Math.max(1, value), function(node) {
      node.__transition__[id].duration = value;
    }));
  };
  d3_transitionPrototype.each = function(type, listener) {
    var id = this.id;
    if (arguments.length < 2) {
      var inherit = d3_transitionInherit, inheritId = d3_transitionInheritId;
      d3_transitionInheritId = id;
      d3_selection_each(this, function(node, i, j) {
        d3_transitionInherit = node.__transition__[id];
        type.call(node, node.__data__, i, j);
      });
      d3_transitionInherit = inherit;
      d3_transitionInheritId = inheritId;
    } else {
      d3_selection_each(this, function(node) {
        var transition = node.__transition__[id];
        (transition.event || (transition.event = d3.dispatch("start", "end"))).on(type, listener);
      });
    }
    return this;
  };
  d3_transitionPrototype.transition = function() {
    var id0 = this.id, id1 = ++d3_transitionId, subgroups = [], subgroup, group, node, transition;
    for (var j = 0, m = this.length; j < m; j++) {
      subgroups.push(subgroup = []);
      for (var group = this[j], i = 0, n = group.length; i < n; i++) {
        if (node = group[i]) {
          transition = Object.create(node.__transition__[id0]);
          transition.delay += transition.duration;
          d3_transitionNode(node, i, id1, transition);
        }
        subgroup.push(node);
      }
    }
    return d3_transition(subgroups, id1);
  };
  function d3_transitionNode(node, i, id, inherit) {
    var lock = node.__transition__ || (node.__transition__ = {
      active: 0,
      count: 0
    }), transition = lock[id];
    if (!transition) {
      var time = inherit.time;
      transition = lock[id] = {
        tween: new d3_Map(),
        time: time,
        ease: inherit.ease,
        delay: inherit.delay,
        duration: inherit.duration
      };
      ++lock.count;
      d3.timer(function(elapsed) {
        var d = node.__data__, ease = transition.ease, delay = transition.delay, duration = transition.duration, timer = d3_timer_active, tweened = [];
        timer.t = delay + time;
        if (delay <= elapsed) return start(elapsed - delay);
        timer.c = start;
        function start(elapsed) {
          if (lock.active > id) return stop();
          lock.active = id;
          transition.event && transition.event.start.call(node, d, i);
          transition.tween.forEach(function(key, value) {
            if (value = value.call(node, d, i)) {
              tweened.push(value);
            }
          });
          d3.timer(function() {
            timer.c = tick(elapsed || 1) ? d3_true : tick;
            return 1;
          }, 0, time);
        }
        function tick(elapsed) {
          if (lock.active !== id) return stop();
          var t = elapsed / duration, e = ease(t), n = tweened.length;
          while (n > 0) {
            tweened[--n].call(node, e);
          }
          if (t >= 1) {
            transition.event && transition.event.end.call(node, d, i);
            return stop();
          }
        }
        function stop() {
          if (--lock.count) delete lock[id]; else delete node.__transition__;
          return 1;
        }
      }, 0, time);
    }
  }
  d3.svg.axis = function() {
    var scale = d3.scale.linear(), orient = d3_svg_axisDefaultOrient, innerTickSize = 6, outerTickSize = 6, tickPadding = 3, tickArguments_ = [ 10 ], tickValues = null, tickFormat_;
    function axis(g) {
      g.each(function() {
        var g = d3.select(this);
        var scale0 = this.__chart__ || scale, scale1 = this.__chart__ = scale.copy();
        var ticks = tickValues == null ? scale1.ticks ? scale1.ticks.apply(scale1, tickArguments_) : scale1.domain() : tickValues, tickFormat = tickFormat_ == null ? scale1.tickFormat ? scale1.tickFormat.apply(scale1, tickArguments_) : d3_identity : tickFormat_, tick = g.selectAll(".tick").data(ticks, scale1), tickEnter = tick.enter().insert("g", ".domain").attr("class", "tick").style("opacity", ε), tickExit = d3.transition(tick.exit()).style("opacity", ε).remove(), tickUpdate = d3.transition(tick).style("opacity", 1), tickTransform;
        var range = d3_scaleRange(scale1), path = g.selectAll(".domain").data([ 0 ]), pathUpdate = (path.enter().append("path").attr("class", "domain"), 
        d3.transition(path));
        tickEnter.append("line");
        tickEnter.append("text");
        var lineEnter = tickEnter.select("line"), lineUpdate = tickUpdate.select("line"), text = tick.select("text").text(tickFormat), textEnter = tickEnter.select("text"), textUpdate = tickUpdate.select("text");
        switch (orient) {
         case "bottom":
          {
            tickTransform = d3_svg_axisX;
            lineEnter.attr("y2", innerTickSize);
            textEnter.attr("y", Math.max(innerTickSize, 0) + tickPadding);
            lineUpdate.attr("x2", 0).attr("y2", innerTickSize);
            textUpdate.attr("x", 0).attr("y", Math.max(innerTickSize, 0) + tickPadding);
            text.attr("dy", ".71em").style("text-anchor", "middle");
            pathUpdate.attr("d", "M" + range[0] + "," + outerTickSize + "V0H" + range[1] + "V" + outerTickSize);
            break;
          }

         case "top":
          {
            tickTransform = d3_svg_axisX;
            lineEnter.attr("y2", -innerTickSize);
            textEnter.attr("y", -(Math.max(innerTickSize, 0) + tickPadding));
            lineUpdate.attr("x2", 0).attr("y2", -innerTickSize);
            textUpdate.attr("x", 0).attr("y", -(Math.max(innerTickSize, 0) + tickPadding));
            text.attr("dy", "0em").style("text-anchor", "middle");
            pathUpdate.attr("d", "M" + range[0] + "," + -outerTickSize + "V0H" + range[1] + "V" + -outerTickSize);
            break;
          }

         case "left":
          {
            tickTransform = d3_svg_axisY;
            lineEnter.attr("x2", -innerTickSize);
            textEnter.attr("x", -(Math.max(innerTickSize, 0) + tickPadding));
            lineUpdate.attr("x2", -innerTickSize).attr("y2", 0);
            textUpdate.attr("x", -(Math.max(innerTickSize, 0) + tickPadding)).attr("y", 0);
            text.attr("dy", ".32em").style("text-anchor", "end");
            pathUpdate.attr("d", "M" + -outerTickSize + "," + range[0] + "H0V" + range[1] + "H" + -outerTickSize);
            break;
          }

         case "right":
          {
            tickTransform = d3_svg_axisY;
            lineEnter.attr("x2", innerTickSize);
            textEnter.attr("x", Math.max(innerTickSize, 0) + tickPadding);
            lineUpdate.attr("x2", innerTickSize).attr("y2", 0);
            textUpdate.attr("x", Math.max(innerTickSize, 0) + tickPadding).attr("y", 0);
            text.attr("dy", ".32em").style("text-anchor", "start");
            pathUpdate.attr("d", "M" + outerTickSize + "," + range[0] + "H0V" + range[1] + "H" + outerTickSize);
            break;
          }
        }
        if (scale1.rangeBand) {
          var x = scale1, dx = x.rangeBand() / 2;
          scale0 = scale1 = function(d) {
            return x(d) + dx;
          };
        } else if (scale0.rangeBand) {
          scale0 = scale1;
        } else {
          tickExit.call(tickTransform, scale1);
        }
        tickEnter.call(tickTransform, scale0);
        tickUpdate.call(tickTransform, scale1);
      });
    }
    axis.scale = function(x) {
      if (!arguments.length) return scale;
      scale = x;
      return axis;
    };
    axis.orient = function(x) {
      if (!arguments.length) return orient;
      orient = x in d3_svg_axisOrients ? x + "" : d3_svg_axisDefaultOrient;
      return axis;
    };
    axis.ticks = function() {
      if (!arguments.length) return tickArguments_;
      tickArguments_ = arguments;
      return axis;
    };
    axis.tickValues = function(x) {
      if (!arguments.length) return tickValues;
      tickValues = x;
      return axis;
    };
    axis.tickFormat = function(x) {
      if (!arguments.length) return tickFormat_;
      tickFormat_ = x;
      return axis;
    };
    axis.tickSize = function(x) {
      var n = arguments.length;
      if (!n) return innerTickSize;
      innerTickSize = +x;
      outerTickSize = +arguments[n - 1];
      return axis;
    };
    axis.innerTickSize = function(x) {
      if (!arguments.length) return innerTickSize;
      innerTickSize = +x;
      return axis;
    };
    axis.outerTickSize = function(x) {
      if (!arguments.length) return outerTickSize;
      outerTickSize = +x;
      return axis;
    };
    axis.tickPadding = function(x) {
      if (!arguments.length) return tickPadding;
      tickPadding = +x;
      return axis;
    };
    axis.tickSubdivide = function() {
      return arguments.length && axis;
    };
    return axis;
  };
  var d3_svg_axisDefaultOrient = "bottom", d3_svg_axisOrients = {
    top: 1,
    right: 1,
    bottom: 1,
    left: 1
  };
  function d3_svg_axisX(selection, x) {
    selection.attr("transform", function(d) {
      return "translate(" + x(d) + ",0)";
    });
  }
  function d3_svg_axisY(selection, y) {
    selection.attr("transform", function(d) {
      return "translate(0," + y(d) + ")";
    });
  }
  d3.svg.brush = function() {
    var event = d3_eventDispatch(brush, "brushstart", "brush", "brushend"), x = null, y = null, xExtent = [ 0, 0 ], yExtent = [ 0, 0 ], xExtentDomain, yExtentDomain, xClamp = true, yClamp = true, resizes = d3_svg_brushResizes[0];
    function brush(g) {
      g.each(function() {
        var g = d3.select(this).style("pointer-events", "all").style("-webkit-tap-highlight-color", "rgba(0,0,0,0)").on("mousedown.brush", brushstart).on("touchstart.brush", brushstart);
        var background = g.selectAll(".background").data([ 0 ]);
        background.enter().append("rect").attr("class", "background").style("visibility", "hidden").style("cursor", "crosshair");
        g.selectAll(".extent").data([ 0 ]).enter().append("rect").attr("class", "extent").style("cursor", "move");
        var resize = g.selectAll(".resize").data(resizes, d3_identity);
        resize.exit().remove();
        resize.enter().append("g").attr("class", function(d) {
          return "resize " + d;
        }).style("cursor", function(d) {
          return d3_svg_brushCursor[d];
        }).append("rect").attr("x", function(d) {
          return /[ew]$/.test(d) ? -3 : null;
        }).attr("y", function(d) {
          return /^[ns]/.test(d) ? -3 : null;
        }).attr("width", 6).attr("height", 6).style("visibility", "hidden");
        resize.style("display", brush.empty() ? "none" : null);
        var gUpdate = d3.transition(g), backgroundUpdate = d3.transition(background), range;
        if (x) {
          range = d3_scaleRange(x);
          backgroundUpdate.attr("x", range[0]).attr("width", range[1] - range[0]);
          redrawX(gUpdate);
        }
        if (y) {
          range = d3_scaleRange(y);
          backgroundUpdate.attr("y", range[0]).attr("height", range[1] - range[0]);
          redrawY(gUpdate);
        }
        redraw(gUpdate);
      });
    }
    brush.event = function(g) {
      g.each(function() {
        var event_ = event.of(this, arguments), extent1 = {
          x: xExtent,
          y: yExtent,
          i: xExtentDomain,
          j: yExtentDomain
        }, extent0 = this.__chart__ || extent1;
        this.__chart__ = extent1;
        if (d3_transitionInheritId) {
          d3.select(this).transition().each("start.brush", function() {
            xExtentDomain = extent0.i;
            yExtentDomain = extent0.j;
            xExtent = extent0.x;
            yExtent = extent0.y;
            event_({
              type: "brushstart"
            });
          }).tween("brush:brush", function() {
            var xi = d3_interpolateArray(xExtent, extent1.x), yi = d3_interpolateArray(yExtent, extent1.y);
            xExtentDomain = yExtentDomain = null;
            return function(t) {
              xExtent = extent1.x = xi(t);
              yExtent = extent1.y = yi(t);
              event_({
                type: "brush",
                mode: "resize"
              });
            };
          }).each("end.brush", function() {
            xExtentDomain = extent1.i;
            yExtentDomain = extent1.j;
            event_({
              type: "brush",
              mode: "resize"
            });
            event_({
              type: "brushend"
            });
          });
        } else {
          event_({
            type: "brushstart"
          });
          event_({
            type: "brush",
            mode: "resize"
          });
          event_({
            type: "brushend"
          });
        }
      });
    };
    function redraw(g) {
      g.selectAll(".resize").attr("transform", function(d) {
        return "translate(" + xExtent[+/e$/.test(d)] + "," + yExtent[+/^s/.test(d)] + ")";
      });
    }
    function redrawX(g) {
      g.select(".extent").attr("x", xExtent[0]);
      g.selectAll(".extent,.n>rect,.s>rect").attr("width", xExtent[1] - xExtent[0]);
    }
    function redrawY(g) {
      g.select(".extent").attr("y", yExtent[0]);
      g.selectAll(".extent,.e>rect,.w>rect").attr("height", yExtent[1] - yExtent[0]);
    }
    function brushstart() {
      var target = this, eventTarget = d3.select(d3.event.target), event_ = event.of(target, arguments), g = d3.select(target), resizing = eventTarget.datum(), resizingX = !/^(n|s)$/.test(resizing) && x, resizingY = !/^(e|w)$/.test(resizing) && y, dragging = eventTarget.classed("extent"), dragRestore = d3_event_dragSuppress(), center, origin = d3.mouse(target), offset;
      var w = d3.select(d3_window).on("keydown.brush", keydown).on("keyup.brush", keyup);
      if (d3.event.changedTouches) {
        w.on("touchmove.brush", brushmove).on("touchend.brush", brushend);
      } else {
        w.on("mousemove.brush", brushmove).on("mouseup.brush", brushend);
      }
      g.interrupt().selectAll("*").interrupt();
      if (dragging) {
        origin[0] = xExtent[0] - origin[0];
        origin[1] = yExtent[0] - origin[1];
      } else if (resizing) {
        var ex = +/w$/.test(resizing), ey = +/^n/.test(resizing);
        offset = [ xExtent[1 - ex] - origin[0], yExtent[1 - ey] - origin[1] ];
        origin[0] = xExtent[ex];
        origin[1] = yExtent[ey];
      } else if (d3.event.altKey) center = origin.slice();
      g.style("pointer-events", "none").selectAll(".resize").style("display", null);
      d3.select("body").style("cursor", eventTarget.style("cursor"));
      event_({
        type: "brushstart"
      });
      brushmove();
      function keydown() {
        if (d3.event.keyCode == 32) {
          if (!dragging) {
            center = null;
            origin[0] -= xExtent[1];
            origin[1] -= yExtent[1];
            dragging = 2;
          }
          d3_eventPreventDefault();
        }
      }
      function keyup() {
        if (d3.event.keyCode == 32 && dragging == 2) {
          origin[0] += xExtent[1];
          origin[1] += yExtent[1];
          dragging = 0;
          d3_eventPreventDefault();
        }
      }
      function brushmove() {
        var point = d3.mouse(target), moved = false;
        if (offset) {
          point[0] += offset[0];
          point[1] += offset[1];
        }
        if (!dragging) {
          if (d3.event.altKey) {
            if (!center) center = [ (xExtent[0] + xExtent[1]) / 2, (yExtent[0] + yExtent[1]) / 2 ];
            origin[0] = xExtent[+(point[0] < center[0])];
            origin[1] = yExtent[+(point[1] < center[1])];
          } else center = null;
        }
        if (resizingX && move1(point, x, 0)) {
          redrawX(g);
          moved = true;
        }
        if (resizingY && move1(point, y, 1)) {
          redrawY(g);
          moved = true;
        }
        if (moved) {
          redraw(g);
          event_({
            type: "brush",
            mode: dragging ? "move" : "resize"
          });
        }
      }
      function move1(point, scale, i) {
        var range = d3_scaleRange(scale), r0 = range[0], r1 = range[1], position = origin[i], extent = i ? yExtent : xExtent, size = extent[1] - extent[0], min, max;
        if (dragging) {
          r0 -= position;
          r1 -= size + position;
        }
        min = (i ? yClamp : xClamp) ? Math.max(r0, Math.min(r1, point[i])) : point[i];
        if (dragging) {
          max = (min += position) + size;
        } else {
          if (center) position = Math.max(r0, Math.min(r1, 2 * center[i] - min));
          if (position < min) {
            max = min;
            min = position;
          } else {
            max = position;
          }
        }
        if (extent[0] != min || extent[1] != max) {
          if (i) yExtentDomain = null; else xExtentDomain = null;
          extent[0] = min;
          extent[1] = max;
          return true;
        }
      }
      function brushend() {
        brushmove();
        g.style("pointer-events", "all").selectAll(".resize").style("display", brush.empty() ? "none" : null);
        d3.select("body").style("cursor", null);
        w.on("mousemove.brush", null).on("mouseup.brush", null).on("touchmove.brush", null).on("touchend.brush", null).on("keydown.brush", null).on("keyup.brush", null);
        dragRestore();
        event_({
          type: "brushend"
        });
      }
    }
    brush.x = function(z) {
      if (!arguments.length) return x;
      x = z;
      resizes = d3_svg_brushResizes[!x << 1 | !y];
      return brush;
    };
    brush.y = function(z) {
      if (!arguments.length) return y;
      y = z;
      resizes = d3_svg_brushResizes[!x << 1 | !y];
      return brush;
    };
    brush.clamp = function(z) {
      if (!arguments.length) return x && y ? [ xClamp, yClamp ] : x ? xClamp : y ? yClamp : null;
      if (x && y) xClamp = !!z[0], yClamp = !!z[1]; else if (x) xClamp = !!z; else if (y) yClamp = !!z;
      return brush;
    };
    brush.extent = function(z) {
      var x0, x1, y0, y1, t;
      if (!arguments.length) {
        if (x) {
          if (xExtentDomain) {
            x0 = xExtentDomain[0], x1 = xExtentDomain[1];
          } else {
            x0 = xExtent[0], x1 = xExtent[1];
            if (x.invert) x0 = x.invert(x0), x1 = x.invert(x1);
            if (x1 < x0) t = x0, x0 = x1, x1 = t;
          }
        }
        if (y) {
          if (yExtentDomain) {
            y0 = yExtentDomain[0], y1 = yExtentDomain[1];
          } else {
            y0 = yExtent[0], y1 = yExtent[1];
            if (y.invert) y0 = y.invert(y0), y1 = y.invert(y1);
            if (y1 < y0) t = y0, y0 = y1, y1 = t;
          }
        }
        return x && y ? [ [ x0, y0 ], [ x1, y1 ] ] : x ? [ x0, x1 ] : y && [ y0, y1 ];
      }
      if (x) {
        x0 = z[0], x1 = z[1];
        if (y) x0 = x0[0], x1 = x1[0];
        xExtentDomain = [ x0, x1 ];
        if (x.invert) x0 = x(x0), x1 = x(x1);
        if (x1 < x0) t = x0, x0 = x1, x1 = t;
        if (x0 != xExtent[0] || x1 != xExtent[1]) xExtent = [ x0, x1 ];
      }
      if (y) {
        y0 = z[0], y1 = z[1];
        if (x) y0 = y0[1], y1 = y1[1];
        yExtentDomain = [ y0, y1 ];
        if (y.invert) y0 = y(y0), y1 = y(y1);
        if (y1 < y0) t = y0, y0 = y1, y1 = t;
        if (y0 != yExtent[0] || y1 != yExtent[1]) yExtent = [ y0, y1 ];
      }
      return brush;
    };
    brush.clear = function() {
      if (!brush.empty()) {
        xExtent = [ 0, 0 ], yExtent = [ 0, 0 ];
        xExtentDomain = yExtentDomain = null;
      }
      return brush;
    };
    brush.empty = function() {
      return !!x && xExtent[0] == xExtent[1] || !!y && yExtent[0] == yExtent[1];
    };
    return d3.rebind(brush, event, "on");
  };
  var d3_svg_brushCursor = {
    n: "ns-resize",
    e: "ew-resize",
    s: "ns-resize",
    w: "ew-resize",
    nw: "nwse-resize",
    ne: "nesw-resize",
    se: "nwse-resize",
    sw: "nesw-resize"
  };
  var d3_svg_brushResizes = [ [ "n", "e", "s", "w", "nw", "ne", "se", "sw" ], [ "e", "w" ], [ "n", "s" ], [] ];
  var d3_time = d3.time = {}, d3_date = Date, d3_time_daySymbols = [ "Sunday", "Monday", "Tuesday", "Wednesday", "Thursday", "Friday", "Saturday" ];
  function d3_date_utc() {
    this._ = new Date(arguments.length > 1 ? Date.UTC.apply(this, arguments) : arguments[0]);
  }
  d3_date_utc.prototype = {
    getDate: function() {
      return this._.getUTCDate();
    },
    getDay: function() {
      return this._.getUTCDay();
    },
    getFullYear: function() {
      return this._.getUTCFullYear();
    },
    getHours: function() {
      return this._.getUTCHours();
    },
    getMilliseconds: function() {
      return this._.getUTCMilliseconds();
    },
    getMinutes: function() {
      return this._.getUTCMinutes();
    },
    getMonth: function() {
      return this._.getUTCMonth();
    },
    getSeconds: function() {
      return this._.getUTCSeconds();
    },
    getTime: function() {
      return this._.getTime();
    },
    getTimezoneOffset: function() {
      return 0;
    },
    valueOf: function() {
      return this._.valueOf();
    },
    setDate: function() {
      d3_time_prototype.setUTCDate.apply(this._, arguments);
    },
    setDay: function() {
      d3_time_prototype.setUTCDay.apply(this._, arguments);
    },
    setFullYear: function() {
      d3_time_prototype.setUTCFullYear.apply(this._, arguments);
    },
    setHours: function() {
      d3_time_prototype.setUTCHours.apply(this._, arguments);
    },
    setMilliseconds: function() {
      d3_time_prototype.setUTCMilliseconds.apply(this._, arguments);
    },
    setMinutes: function() {
      d3_time_prototype.setUTCMinutes.apply(this._, arguments);
    },
    setMonth: function() {
      d3_time_prototype.setUTCMonth.apply(this._, arguments);
    },
    setSeconds: function() {
      d3_time_prototype.setUTCSeconds.apply(this._, arguments);
    },
    setTime: function() {
      d3_time_prototype.setTime.apply(this._, arguments);
    }
  };
  var d3_time_prototype = Date.prototype;
  var d3_time_formatDateTime = "%a %b %e %X %Y", d3_time_formatDate = "%m/%d/%Y", d3_time_formatTime = "%H:%M:%S";
  var d3_time_days = [ "Sunday", "Monday", "Tuesday", "Wednesday", "Thursday", "Friday", "Saturday" ], d3_time_dayAbbreviations = [ "Sun", "Mon", "Tue", "Wed", "Thu", "Fri", "Sat" ], d3_time_months = [ "January", "February", "March", "April", "May", "June", "July", "August", "September", "October", "November", "December" ], d3_time_monthAbbreviations = [ "Jan", "Feb", "Mar", "Apr", "May", "Jun", "Jul", "Aug", "Sep", "Oct", "Nov", "Dec" ];
  function d3_time_interval(local, step, number) {
    function round(date) {
      var d0 = local(date), d1 = offset(d0, 1);
      return date - d0 < d1 - date ? d0 : d1;
    }
    function ceil(date) {
      step(date = local(new d3_date(date - 1)), 1);
      return date;
    }
    function offset(date, k) {
      step(date = new d3_date(+date), k);
      return date;
    }
    function range(t0, t1, dt) {
      var time = ceil(t0), times = [];
      if (dt > 1) {
        while (time < t1) {
          if (!(number(time) % dt)) times.push(new Date(+time));
          step(time, 1);
        }
      } else {
        while (time < t1) times.push(new Date(+time)), step(time, 1);
      }
      return times;
    }
    function range_utc(t0, t1, dt) {
      try {
        d3_date = d3_date_utc;
        var utc = new d3_date_utc();
        utc._ = t0;
        return range(utc, t1, dt);
      } finally {
        d3_date = Date;
      }
    }
    local.floor = local;
    local.round = round;
    local.ceil = ceil;
    local.offset = offset;
    local.range = range;
    var utc = local.utc = d3_time_interval_utc(local);
    utc.floor = utc;
    utc.round = d3_time_interval_utc(round);
    utc.ceil = d3_time_interval_utc(ceil);
    utc.offset = d3_time_interval_utc(offset);
    utc.range = range_utc;
    return local;
  }
  function d3_time_interval_utc(method) {
    return function(date, k) {
      try {
        d3_date = d3_date_utc;
        var utc = new d3_date_utc();
        utc._ = date;
        return method(utc, k)._;
      } finally {
        d3_date = Date;
      }
    };
  }
  d3_time.year = d3_time_interval(function(date) {
    date = d3_time.day(date);
    date.setMonth(0, 1);
    return date;
  }, function(date, offset) {
    date.setFullYear(date.getFullYear() + offset);
  }, function(date) {
    return date.getFullYear();
  });
  d3_time.years = d3_time.year.range;
  d3_time.years.utc = d3_time.year.utc.range;
  d3_time.day = d3_time_interval(function(date) {
    var day = new d3_date(2e3, 0);
    day.setFullYear(date.getFullYear(), date.getMonth(), date.getDate());
    return day;
  }, function(date, offset) {
    date.setDate(date.getDate() + offset);
  }, function(date) {
    return date.getDate() - 1;
  });
  d3_time.days = d3_time.day.range;
  d3_time.days.utc = d3_time.day.utc.range;
  d3_time.dayOfYear = function(date) {
    var year = d3_time.year(date);
    return Math.floor((date - year - (date.getTimezoneOffset() - year.getTimezoneOffset()) * 6e4) / 864e5);
  };
  d3_time_daySymbols.forEach(function(day, i) {
    day = day.toLowerCase();
    i = 7 - i;
    var interval = d3_time[day] = d3_time_interval(function(date) {
      (date = d3_time.day(date)).setDate(date.getDate() - (date.getDay() + i) % 7);
      return date;
    }, function(date, offset) {
      date.setDate(date.getDate() + Math.floor(offset) * 7);
    }, function(date) {
      var day = d3_time.year(date).getDay();
      return Math.floor((d3_time.dayOfYear(date) + (day + i) % 7) / 7) - (day !== i);
    });
    d3_time[day + "s"] = interval.range;
    d3_time[day + "s"].utc = interval.utc.range;
    d3_time[day + "OfYear"] = function(date) {
      var day = d3_time.year(date).getDay();
      return Math.floor((d3_time.dayOfYear(date) + (day + i) % 7) / 7);
    };
  });
  d3_time.week = d3_time.sunday;
  d3_time.weeks = d3_time.sunday.range;
  d3_time.weeks.utc = d3_time.sunday.utc.range;
  d3_time.weekOfYear = d3_time.sundayOfYear;
  d3_time.format = d3_time_format;
  function d3_time_format(template) {
    var n = template.length;
    function format(date) {
      var string = [], i = -1, j = 0, c, p, f;
      while (++i < n) {
        if (template.charCodeAt(i) === 37) {
          string.push(template.substring(j, i));
          if ((p = d3_time_formatPads[c = template.charAt(++i)]) != null) c = template.charAt(++i);
          if (f = d3_time_formats[c]) c = f(date, p == null ? c === "e" ? " " : "0" : p);
          string.push(c);
          j = i + 1;
        }
      }
      string.push(template.substring(j, i));
      return string.join("");
    }
    format.parse = function(string) {
      var d = {
        y: 1900,
        m: 0,
        d: 1,
        H: 0,
        M: 0,
        S: 0,
        L: 0,
        Z: null
      }, i = d3_time_parse(d, template, string, 0);
      if (i != string.length) return null;
      if ("p" in d) d.H = d.H % 12 + d.p * 12;
      var localZ = d.Z != null && d3_date !== d3_date_utc, date = new (localZ ? d3_date_utc : d3_date)();
      if ("j" in d) date.setFullYear(d.y, 0, d.j); else if ("w" in d && ("W" in d || "U" in d)) {
        date.setFullYear(d.y, 0, 1);
        date.setFullYear(d.y, 0, "W" in d ? (d.w + 6) % 7 + d.W * 7 - (date.getDay() + 5) % 7 : d.w + d.U * 7 - (date.getDay() + 6) % 7);
      } else date.setFullYear(d.y, d.m, d.d);
      date.setHours(d.H + Math.floor(d.Z / 100), d.M + d.Z % 100, d.S, d.L);
      return localZ ? date._ : date;
    };
    format.toString = function() {
      return template;
    };
    return format;
  }
  function d3_time_parse(date, template, string, j) {
    var c, p, t, i = 0, n = template.length, m = string.length;
    while (i < n) {
      if (j >= m) return -1;
      c = template.charCodeAt(i++);
      if (c === 37) {
        t = template.charAt(i++);
        p = d3_time_parsers[t in d3_time_formatPads ? template.charAt(i++) : t];
        if (!p || (j = p(date, string, j)) < 0) return -1;
      } else if (c != string.charCodeAt(j++)) {
        return -1;
      }
    }
    return j;
  }
  function d3_time_formatRe(names) {
    return new RegExp("^(?:" + names.map(d3.requote).join("|") + ")", "i");
  }
  function d3_time_formatLookup(names) {
    var map = new d3_Map(), i = -1, n = names.length;
    while (++i < n) map.set(names[i].toLowerCase(), i);
    return map;
  }
  function d3_time_formatPad(value, fill, width) {
    var sign = value < 0 ? "-" : "", string = (sign ? -value : value) + "", length = string.length;
    return sign + (length < width ? new Array(width - length + 1).join(fill) + string : string);
  }
  var d3_time_dayRe = d3_time_formatRe(d3_time_days), d3_time_dayLookup = d3_time_formatLookup(d3_time_days), d3_time_dayAbbrevRe = d3_time_formatRe(d3_time_dayAbbreviations), d3_time_dayAbbrevLookup = d3_time_formatLookup(d3_time_dayAbbreviations), d3_time_monthRe = d3_time_formatRe(d3_time_months), d3_time_monthLookup = d3_time_formatLookup(d3_time_months), d3_time_monthAbbrevRe = d3_time_formatRe(d3_time_monthAbbreviations), d3_time_monthAbbrevLookup = d3_time_formatLookup(d3_time_monthAbbreviations), d3_time_percentRe = /^%/;
  var d3_time_formatPads = {
    "-": "",
    _: " ",
    "0": "0"
  };
  var d3_time_formats = {
    a: function(d) {
      return d3_time_dayAbbreviations[d.getDay()];
    },
    A: function(d) {
      return d3_time_days[d.getDay()];
    },
    b: function(d) {
      return d3_time_monthAbbreviations[d.getMonth()];
    },
    B: function(d) {
      return d3_time_months[d.getMonth()];
    },
    c: d3_time_format(d3_time_formatDateTime),
    d: function(d, p) {
      return d3_time_formatPad(d.getDate(), p, 2);
    },
    e: function(d, p) {
      return d3_time_formatPad(d.getDate(), p, 2);
    },
    H: function(d, p) {
      return d3_time_formatPad(d.getHours(), p, 2);
    },
    I: function(d, p) {
      return d3_time_formatPad(d.getHours() % 12 || 12, p, 2);
    },
    j: function(d, p) {
      return d3_time_formatPad(1 + d3_time.dayOfYear(d), p, 3);
    },
    L: function(d, p) {
      return d3_time_formatPad(d.getMilliseconds(), p, 3);
    },
    m: function(d, p) {
      return d3_time_formatPad(d.getMonth() + 1, p, 2);
    },
    M: function(d, p) {
      return d3_time_formatPad(d.getMinutes(), p, 2);
    },
    p: function(d) {
      return d.getHours() >= 12 ? "PM" : "AM";
    },
    S: function(d, p) {
      return d3_time_formatPad(d.getSeconds(), p, 2);
    },
    U: function(d, p) {
      return d3_time_formatPad(d3_time.sundayOfYear(d), p, 2);
    },
    w: function(d) {
      return d.getDay();
    },
    W: function(d, p) {
      return d3_time_formatPad(d3_time.mondayOfYear(d), p, 2);
    },
    x: d3_time_format(d3_time_formatDate),
    X: d3_time_format(d3_time_formatTime),
    y: function(d, p) {
      return d3_time_formatPad(d.getFullYear() % 100, p, 2);
    },
    Y: function(d, p) {
      return d3_time_formatPad(d.getFullYear() % 1e4, p, 4);
    },
    Z: d3_time_zone,
    "%": function() {
      return "%";
    }
  };
  var d3_time_parsers = {
    a: d3_time_parseWeekdayAbbrev,
    A: d3_time_parseWeekday,
    b: d3_time_parseMonthAbbrev,
    B: d3_time_parseMonth,
    c: d3_time_parseLocaleFull,
    d: d3_time_parseDay,
    e: d3_time_parseDay,
    H: d3_time_parseHour24,
    I: d3_time_parseHour24,
    j: d3_time_parseDayOfYear,
    L: d3_time_parseMilliseconds,
    m: d3_time_parseMonthNumber,
    M: d3_time_parseMinutes,
    p: d3_time_parseAmPm,
    S: d3_time_parseSeconds,
    U: d3_time_parseWeekNumberSunday,
    w: d3_time_parseWeekdayNumber,
    W: d3_time_parseWeekNumberMonday,
    x: d3_time_parseLocaleDate,
    X: d3_time_parseLocaleTime,
    y: d3_time_parseYear,
    Y: d3_time_parseFullYear,
    Z: d3_time_parseZone,
    "%": d3_time_parseLiteralPercent
  };
  function d3_time_parseWeekdayAbbrev(date, string, i) {
    d3_time_dayAbbrevRe.lastIndex = 0;
    var n = d3_time_dayAbbrevRe.exec(string.substring(i));
    return n ? (date.w = d3_time_dayAbbrevLookup.get(n[0].toLowerCase()), i + n[0].length) : -1;
  }
  function d3_time_parseWeekday(date, string, i) {
    d3_time_dayRe.lastIndex = 0;
    var n = d3_time_dayRe.exec(string.substring(i));
    return n ? (date.w = d3_time_dayLookup.get(n[0].toLowerCase()), i + n[0].length) : -1;
  }
  function d3_time_parseWeekdayNumber(date, string, i) {
    d3_time_numberRe.lastIndex = 0;
    var n = d3_time_numberRe.exec(string.substring(i, i + 1));
    return n ? (date.w = +n[0], i + n[0].length) : -1;
  }
  function d3_time_parseWeekNumberSunday(date, string, i) {
    d3_time_numberRe.lastIndex = 0;
    var n = d3_time_numberRe.exec(string.substring(i));
    return n ? (date.U = +n[0], i + n[0].length) : -1;
  }
  function d3_time_parseWeekNumberMonday(date, string, i) {
    d3_time_numberRe.lastIndex = 0;
    var n = d3_time_numberRe.exec(string.substring(i));
    return n ? (date.W = +n[0], i + n[0].length) : -1;
  }
  function d3_time_parseMonthAbbrev(date, string, i) {
    d3_time_monthAbbrevRe.lastIndex = 0;
    var n = d3_time_monthAbbrevRe.exec(string.substring(i));
    return n ? (date.m = d3_time_monthAbbrevLookup.get(n[0].toLowerCase()), i + n[0].length) : -1;
  }
  function d3_time_parseMonth(date, string, i) {
    d3_time_monthRe.lastIndex = 0;
    var n = d3_time_monthRe.exec(string.substring(i));
    return n ? (date.m = d3_time_monthLookup.get(n[0].toLowerCase()), i + n[0].length) : -1;
  }
  function d3_time_parseLocaleFull(date, string, i) {
    return d3_time_parse(date, d3_time_formats.c.toString(), string, i);
  }
  function d3_time_parseLocaleDate(date, string, i) {
    return d3_time_parse(date, d3_time_formats.x.toString(), string, i);
  }
  function d3_time_parseLocaleTime(date, string, i) {
    return d3_time_parse(date, d3_time_formats.X.toString(), string, i);
  }
  function d3_time_parseFullYear(date, string, i) {
    d3_time_numberRe.lastIndex = 0;
    var n = d3_time_numberRe.exec(string.substring(i, i + 4));
    return n ? (date.y = +n[0], i + n[0].length) : -1;
  }
  function d3_time_parseYear(date, string, i) {
    d3_time_numberRe.lastIndex = 0;
    var n = d3_time_numberRe.exec(string.substring(i, i + 2));
    return n ? (date.y = d3_time_expandYear(+n[0]), i + n[0].length) : -1;
  }
  function d3_time_parseZone(date, string, i) {
    return /^[+-]\d{4}$/.test(string = string.substring(i, i + 5)) ? (date.Z = +string, 
    i + 5) : -1;
  }
  function d3_time_expandYear(d) {
    return d + (d > 68 ? 1900 : 2e3);
  }
  function d3_time_parseMonthNumber(date, string, i) {
    d3_time_numberRe.lastIndex = 0;
    var n = d3_time_numberRe.exec(string.substring(i, i + 2));
    return n ? (date.m = n[0] - 1, i + n[0].length) : -1;
  }
  function d3_time_parseDay(date, string, i) {
    d3_time_numberRe.lastIndex = 0;
    var n = d3_time_numberRe.exec(string.substring(i, i + 2));
    return n ? (date.d = +n[0], i + n[0].length) : -1;
  }
  function d3_time_parseDayOfYear(date, string, i) {
    d3_time_numberRe.lastIndex = 0;
    var n = d3_time_numberRe.exec(string.substring(i, i + 3));
    return n ? (date.j = +n[0], i + n[0].length) : -1;
  }
  function d3_time_parseHour24(date, string, i) {
    d3_time_numberRe.lastIndex = 0;
    var n = d3_time_numberRe.exec(string.substring(i, i + 2));
    return n ? (date.H = +n[0], i + n[0].length) : -1;
  }
  function d3_time_parseMinutes(date, string, i) {
    d3_time_numberRe.lastIndex = 0;
    var n = d3_time_numberRe.exec(string.substring(i, i + 2));
    return n ? (date.M = +n[0], i + n[0].length) : -1;
  }
  function d3_time_parseSeconds(date, string, i) {
    d3_time_numberRe.lastIndex = 0;
    var n = d3_time_numberRe.exec(string.substring(i, i + 2));
    return n ? (date.S = +n[0], i + n[0].length) : -1;
  }
  function d3_time_parseMilliseconds(date, string, i) {
    d3_time_numberRe.lastIndex = 0;
    var n = d3_time_numberRe.exec(string.substring(i, i + 3));
    return n ? (date.L = +n[0], i + n[0].length) : -1;
  }
  var d3_time_numberRe = /^\s*\d+/;
  function d3_time_parseAmPm(date, string, i) {
    var n = d3_time_amPmLookup.get(string.substring(i, i += 2).toLowerCase());
    return n == null ? -1 : (date.p = n, i);
  }
  var d3_time_amPmLookup = d3.map({
    am: 0,
    pm: 1
  });
  function d3_time_zone(d) {
    var z = d.getTimezoneOffset(), zs = z > 0 ? "-" : "+", zh = ~~(abs(z) / 60), zm = abs(z) % 60;
    return zs + d3_time_formatPad(zh, "0", 2) + d3_time_formatPad(zm, "0", 2);
  }
  function d3_time_parseLiteralPercent(date, string, i) {
    d3_time_percentRe.lastIndex = 0;
    var n = d3_time_percentRe.exec(string.substring(i, i + 1));
    return n ? i + n[0].length : -1;
  }
  d3_time_format.utc = d3_time_formatUtc;
  function d3_time_formatUtc(template) {
    var local = d3_time_format(template);
    function format(date) {
      try {
        d3_date = d3_date_utc;
        var utc = new d3_date();
        utc._ = date;
        return local(utc);
      } finally {
        d3_date = Date;
      }
    }
    format.parse = function(string) {
      try {
        d3_date = d3_date_utc;
        var date = local.parse(string);
        return date && date._;
      } finally {
        d3_date = Date;
      }
    };
    format.toString = local.toString;
    return format;
  }
  var d3_time_formatIso = d3_time_formatUtc("%Y-%m-%dT%H:%M:%S.%LZ");
  d3_time_format.iso = Date.prototype.toISOString && +new Date("2000-01-01T00:00:00.000Z") ? d3_time_formatIsoNative : d3_time_formatIso;
  function d3_time_formatIsoNative(date) {
    return date.toISOString();
  }
  d3_time_formatIsoNative.parse = function(string) {
    var date = new Date(string);
    return isNaN(date) ? null : date;
  };
  d3_time_formatIsoNative.toString = d3_time_formatIso.toString;
  d3_time.second = d3_time_interval(function(date) {
    return new d3_date(Math.floor(date / 1e3) * 1e3);
  }, function(date, offset) {
    date.setTime(date.getTime() + Math.floor(offset) * 1e3);
  }, function(date) {
    return date.getSeconds();
  });
  d3_time.seconds = d3_time.second.range;
  d3_time.seconds.utc = d3_time.second.utc.range;
  d3_time.minute = d3_time_interval(function(date) {
    return new d3_date(Math.floor(date / 6e4) * 6e4);
  }, function(date, offset) {
    date.setTime(date.getTime() + Math.floor(offset) * 6e4);
  }, function(date) {
    return date.getMinutes();
  });
  d3_time.minutes = d3_time.minute.range;
  d3_time.minutes.utc = d3_time.minute.utc.range;
  d3_time.hour = d3_time_interval(function(date) {
    var timezone = date.getTimezoneOffset() / 60;
    return new d3_date((Math.floor(date / 36e5 - timezone) + timezone) * 36e5);
  }, function(date, offset) {
    date.setTime(date.getTime() + Math.floor(offset) * 36e5);
  }, function(date) {
    return date.getHours();
  });
  d3_time.hours = d3_time.hour.range;
  d3_time.hours.utc = d3_time.hour.utc.range;
  d3_time.month = d3_time_interval(function(date) {
    date = d3_time.day(date);
    date.setDate(1);
    return date;
  }, function(date, offset) {
    date.setMonth(date.getMonth() + offset);
  }, function(date) {
    return date.getMonth();
  });
  d3_time.months = d3_time.month.range;
  d3_time.months.utc = d3_time.month.utc.range;
  function d3_time_scale(linear, methods, format) {
    function scale(x) {
      return linear(x);
    }
    scale.invert = function(x) {
      return d3_time_scaleDate(linear.invert(x));
    };
    scale.domain = function(x) {
      if (!arguments.length) return linear.domain().map(d3_time_scaleDate);
      linear.domain(x);
      return scale;
    };
    function tickMethod(extent, count) {
      var span = extent[1] - extent[0], target = span / count, i = d3.bisect(d3_time_scaleSteps, target);
      return i == d3_time_scaleSteps.length ? [ methods.year, d3_scale_linearTickRange(extent.map(function(d) {
        return d / 31536e6;
      }), count)[2] ] : !i ? [ d3_time_scaleMilliseconds, d3_scale_linearTickRange(extent, count)[2] ] : methods[target / d3_time_scaleSteps[i - 1] < d3_time_scaleSteps[i] / target ? i - 1 : i];
    }
    scale.nice = function(interval, skip) {
      var domain = scale.domain(), extent = d3_scaleExtent(domain), method = interval == null ? tickMethod(extent, 10) : typeof interval === "number" && tickMethod(extent, interval);
      if (method) interval = method[0], skip = method[1];
      function skipped(date) {
        return !isNaN(date) && !interval.range(date, d3_time_scaleDate(+date + 1), skip).length;
      }
      return scale.domain(d3_scale_nice(domain, skip > 1 ? {
        floor: function(date) {
          while (skipped(date = interval.floor(date))) date = d3_time_scaleDate(date - 1);
          return date;
        },
        ceil: function(date) {
          while (skipped(date = interval.ceil(date))) date = d3_time_scaleDate(+date + 1);
          return date;
        }
      } : interval));
    };
    scale.ticks = function(interval, skip) {
      var extent = d3_scaleExtent(scale.domain()), method = interval == null ? tickMethod(extent, 10) : typeof interval === "number" ? tickMethod(extent, interval) : !interval.range && [ {
        range: interval
      }, skip ];
      if (method) interval = method[0], skip = method[1];
      return interval.range(extent[0], d3_time_scaleDate(+extent[1] + 1), skip < 1 ? 1 : skip);
    };
    scale.tickFormat = function() {
      return format;
    };
    scale.copy = function() {
      return d3_time_scale(linear.copy(), methods, format);
    };
    return d3_scale_linearRebind(scale, linear);
  }
  function d3_time_scaleDate(t) {
    return new Date(t);
  }
  function d3_time_scaleFormat(formats) {
    return function(date) {
      var i = formats.length - 1, f = formats[i];
      while (!f[1](date)) f = formats[--i];
      return f[0](date);
    };
  }
  var d3_time_scaleSteps = [ 1e3, 5e3, 15e3, 3e4, 6e4, 3e5, 9e5, 18e5, 36e5, 108e5, 216e5, 432e5, 864e5, 1728e5, 6048e5, 2592e6, 7776e6, 31536e6 ];
  var d3_time_scaleLocalMethods = [ [ d3_time.second, 1 ], [ d3_time.second, 5 ], [ d3_time.second, 15 ], [ d3_time.second, 30 ], [ d3_time.minute, 1 ], [ d3_time.minute, 5 ], [ d3_time.minute, 15 ], [ d3_time.minute, 30 ], [ d3_time.hour, 1 ], [ d3_time.hour, 3 ], [ d3_time.hour, 6 ], [ d3_time.hour, 12 ], [ d3_time.day, 1 ], [ d3_time.day, 2 ], [ d3_time.week, 1 ], [ d3_time.month, 1 ], [ d3_time.month, 3 ], [ d3_time.year, 1 ] ];
  var d3_time_scaleLocalFormats = [ [ d3_time_format("%Y"), d3_true ], [ d3_time_format("%B"), function(d) {
    return d.getMonth();
  } ], [ d3_time_format("%b %d"), function(d) {
    return d.getDate() != 1;
  } ], [ d3_time_format("%a %d"), function(d) {
    return d.getDay() && d.getDate() != 1;
  } ], [ d3_time_format("%I %p"), function(d) {
    return d.getHours();
  } ], [ d3_time_format("%I:%M"), function(d) {
    return d.getMinutes();
  } ], [ d3_time_format(":%S"), function(d) {
    return d.getSeconds();
  } ], [ d3_time_format(".%L"), function(d) {
    return d.getMilliseconds();
  } ] ];
  var d3_time_scaleLocalFormat = d3_time_scaleFormat(d3_time_scaleLocalFormats);
  d3_time_scaleLocalMethods.year = d3_time.year;
  d3_time.scale = function() {
    return d3_time_scale(d3.scale.linear(), d3_time_scaleLocalMethods, d3_time_scaleLocalFormat);
  };
  var d3_time_scaleMilliseconds = {
    range: function(start, stop, step) {
      return d3.range(+start, +stop, step).map(d3_time_scaleDate);
    }
  };
  var d3_time_scaleUTCMethods = d3_time_scaleLocalMethods.map(function(m) {
    return [ m[0].utc, m[1] ];
  });
  var d3_time_scaleUTCFormats = [ [ d3_time_formatUtc("%Y"), d3_true ], [ d3_time_formatUtc("%B"), function(d) {
    return d.getUTCMonth();
  } ], [ d3_time_formatUtc("%b %d"), function(d) {
    return d.getUTCDate() != 1;
  } ], [ d3_time_formatUtc("%a %d"), function(d) {
    return d.getUTCDay() && d.getUTCDate() != 1;
  } ], [ d3_time_formatUtc("%I %p"), function(d) {
    return d.getUTCHours();
  } ], [ d3_time_formatUtc("%I:%M"), function(d) {
    return d.getUTCMinutes();
  } ], [ d3_time_formatUtc(":%S"), function(d) {
    return d.getUTCSeconds();
  } ], [ d3_time_formatUtc(".%L"), function(d) {
    return d.getUTCMilliseconds();
  } ] ];
  var d3_time_scaleUTCFormat = d3_time_scaleFormat(d3_time_scaleUTCFormats);
  d3_time_scaleUTCMethods.year = d3_time.year.utc;
  d3_time.scale.utc = function() {
    return d3_time_scale(d3.scale.linear(), d3_time_scaleUTCMethods, d3_time_scaleUTCFormat);
  };
  d3.text = d3_xhrType(function(request) {
    return request.responseText;
  });
  d3.json = function(url, callback) {
    return d3_xhr(url, "application/json", d3_json, callback);
  };
  function d3_json(request) {
    return JSON.parse(request.responseText);
  }
  d3.html = function(url, callback) {
    return d3_xhr(url, "text/html", d3_html, callback);
  };
  function d3_html(request) {
    var range = d3_document.createRange();
    range.selectNode(d3_document.body);
    return range.createContextualFragment(request.responseText);
  }
  d3.xml = d3_xhrType(function(request) {
    return request.responseXML;
  });
  return d3;
}();
```
