## Supplementary Material to Figure 4 and Figure 6 for "BTRR complex deficiency is a driver for genomic instability in Bloom syndrome": BioCircos.js

```
HTMLWidgets.widget({

  name: 'BioCircos',

  type: 'output',

  factory: function(el, width, height) {

    // Define shared variables for this instance

    function objToArray(objectLiteral) {
      var piece1 = Object.keys(objectLiteral);
      var piece2 = Object.values(objectLiteral);
      var result = [];
      for (var i = 0; i < piece1.length; i++) {
        result.push([piece1[i], piece2[i]])
      }
      return result;
    }

    return {

      renderValue: function(opts) {
        // Remove previous occurences of plots in the <div> if any
        d3.select("#"+el.id).selectAll("svg").remove()

        var BioCircosGenome = objToArray(opts.genome);

        var maxRadius = Math.min(height, width)/2; // Compute maximal radial space

        // Adapt the radii from relative to absolute values
        for (var i = 0; i < opts.tracklist.length; i++) { // For each track
          opts.tracklist[i][1].maxRadius *= 0.7*maxRadius
          opts.tracklist[i][1].minRadius *= 0.7*maxRadius
          opts.tracklist[i][1].BgouterRadius *= 0.7*maxRadius
          opts.tracklist[i][1].BginnerRadius *= 0.7*maxRadius
          opts.tracklist[i][1].outerRadius *= 0.7*maxRadius
          opts.tracklist[i][1].innerRadius *= 0.7*maxRadius
          opts.tracklist[i][1].x *= 0.7*maxRadius
          opts.tracklist[i][1].y *= 0.7*maxRadius
          opts.tracklist[i][1].LinkRadius *= 0.7*maxRadius 
        }

        BioCircos01 = new BioCircos(... opts.tracklist, BioCircosGenome,{
          // Main configuration
          target : el.id,
          svgWidth : width,
          svgHeight : height,
          chrPad : opts.chrPad,
          innerRadius: 0.7*maxRadius,
          outerRadius: 0.8*maxRadius,
          zoom : opts.zoom, // Allow zoom and translation

          // SNP interaction options
          SNPMouseEvent : opts.SNPMouseEvent,
          SNPMouseClickDisplay : opts.SNPMouseClickDisplay,
          SNPMouseClickColor : opts.SNPMouseClickColor,
          SNPMouseClickCircleSize : opts.SNPMouseClickCircleSize,
          SNPMouseClickCircleOpacity : opts.SNPMouseClickCircleOpacity,
          SNPMouseClickCircleStrokeColor : opts.SNPMouseClickCircleStrokeColor,
          SNPMouseClickCircleStrokeWidth : opts.SNPMouseClickCircleStrokeWidth,
          SNPMouseClickTextFromData : opts.SNPMouseClickTextFromData,
          SNPMouseClickTextOpacity : opts.SNPMouseClickTextOpacity,
          SNPMouseClickTextColor : opts.SNPMouseClickTextColor,
          SNPMouseClickTextSize : opts.SNPMouseClickTextSize,
          SNPMouseClickTextPostionX : opts.SNPMouseClickTextPostionX,
          SNPMouseClickTextPostionY : opts.SNPMouseClickTextPostionY,
          SNPMouseClickTextDrag : opts.SNPMouseClickTextDrag,
          SNPMouseDownDisplay : opts.SNPMouseDownDisplay,
          SNPMouseDownColor : opts.SNPMouseDownColor,
          SNPMouseDownCircleSize : opts.SNPMouseDownCircleSize,
          SNPMouseDownCircleOpacity : opts.SNPMouseDownCircleOpacity,
          SNPMouseDownCircleStrokeColor : opts.SNPMouseDownCircleStrokeColor,
          SNPMouseDownCircleStrokeWidth : opts.SNPMouseDownCircleStrokeWidth,
          SNPMouseEnterDisplay : opts.SNPMouseEnterDisplay,
          SNPMouseEnterColor : opts.SNPMouseEnterColor,
          SNPMouseEnterCircleSize : opts.SNPMouseEnterCircleSize,
          SNPMouseEnterCircleOpacity : opts.SNPMouseEnterCircleOpacity,
          SNPMouseEnterCircleStrokeColor : opts.SNPMouseEnterCircleStrokeColor,
          SNPMouseEnterCircleStrokeWidth : opts.SNPMouseEnterCircleStrokeWidth,
          SNPMouseLeaveDisplay : opts.SNPMouseLeaveDisplay,
          SNPMouseLeaveColor : opts.SNPMouseLeaveColor,
          SNPMouseLeaveCircleSize : opts.SNPMouseLeaveCircleSize,
          SNPMouseLeaveCircleOpacity : opts.SNPMouseLeaveCircleOpacity,
          SNPMouseLeaveCircleStrokeColor : opts.SNPMouseLeaveCircleStrokeColor,
          SNPMouseLeaveCircleStrokeWidth : opts.SNPMouseLeaveCircleStrokeWidth,
          SNPMouseMoveDisplay : opts.SNPMouseMoveDisplay,
          SNPMouseMoveColor : opts.SNPMouseMoveColor,
          SNPMouseMoveCircleSize : opts.SNPMouseMoveCircleSize,
          SNPMouseMoveCircleOpacity : opts.SNPMouseMoveCircleOpacity,
          SNPMouseMoveCircleStrokeColor : opts.SNPMouseMoveCircleStrokeColor,
          SNPMouseMoveCircleStrokeWidth : opts.SNPMouseMoveCircleStrokeWidth,
          SNPMouseOutDisplay : opts.SNPMouseOutDisplay,
          SNPMouseOutAnimationTime : opts.SNPMouseOutAnimationTime,
          SNPMouseOutColor : opts.SNPMouseOutColor,
          SNPMouseOutCircleSize : opts.SNPMouseOutCircleSize,
          SNPMouseOutCircleOpacity : opts.SNPMouseOutCircleOpacity,
          SNPMouseOutCircleStrokeColor : opts.SNPMouseOutCircleStrokeColor,
          SNPMouseOutCircleStrokeWidth : opts.SNPMouseOutCircleStrokeWidth,
          SNPMouseUpDisplay : opts.SNPMouseUpDisplay,
          SNPMouseUpColor : opts.SNPMouseUpColor,
          SNPMouseUpCircleSize : opts.SNPMouseUpCircleSize,
          SNPMouseUpCircleOpacity : opts.SNPMouseUpCircleOpacity,
          SNPMouseUpCircleStrokeColor : opts.SNPMouseUpCircleStrokeColor,
          SNPMouseUpCircleStrokeWidth : opts.SNPMouseUpCircleStrokeWidth,
          SNPMouseOverDisplay : opts.SNPMouseOverDisplay,
          SNPMouseOverColor : opts.SNPMouseOverColor,
          SNPMouseOverCircleSize : opts.SNPMouseOverCircleSize,
          SNPMouseOverCircleOpacity : opts.SNPMouseOverCircleOpacity,
          SNPMouseOverCircleStrokeColor : opts.SNPMouseOverCircleStrokeColor,
          SNPMouseOverCircleStrokeWidth : opts.SNPMouseOverCircleStrokeWidth,

          SNPMouseOverTooltipsHtml01 : opts.SNPMouseOverTooltipsHtml01,
          SNPMouseOverTooltipsHtml02 : opts.SNPMouseOverTooltipsHtml02,
          SNPMouseOverTooltipsHtml03 : opts.SNPMouseOverTooltipsHtml03,
          SNPMouseOverTooltipsHtml04 : opts.SNPMouseOverTooltipsHtml04,
          SNPMouseOverTooltipsHtml05 : opts.SNPMouseOverTooltipsHtml05,
          SNPMouseOverTooltipsBorderWidth : opts.SNPMouseOverTooltipsBorderWidth,

          // Arc interaction options
          ARCMouseEvent : opts.ARCMouseEvent,
          ARCMouseClickDisplay : opts.ARCMouseClickDisplay,
          ARCMouseClickColor : opts.ARCMouseClickColor,
          ARCMouseClickArcOpacity : opts.ARCMouseClickArcOpacity,
          ARCMouseClickArcStrokeColor : opts.ARCMouseClickArcStrokeColor,
          ARCMouseClickArcStrokeWidth : opts.ARCMouseClickArcStrokeWidth,
          ARCMouseClickTextFromData : opts.ARCMouseClickTextFromData,
          ARCMouseClickTextOpacity : opts.ARCMouseClickTextOpacity,
          ARCMouseClickTextColor : opts.ARCMouseClickTextColor,
          ARCMouseClickTextSize : opts.ARCMouseClickTextSize,
          ARCMouseClickTextPostionX : opts.ARCMouseClickTextPostionX,
          ARCMouseClickTextPostionY : opts.ARCMouseClickTextPostionY,
          ARCMouseClickTextDrag : opts.ARCMouseClickTextDrag,
          ARCMouseDownDisplay : opts.ARCMouseDownDisplay,
          ARCMouseDownColor : opts.ARCMouseDownColor,
          ARCMouseDownArcOpacity : opts.ARCMouseDownArcOpacity,
          ARCMouseDownArcStrokeColor : opts.ARCMouseDownArcStrokeColor,
          ARCMouseDownArcStrokeWidth : opts.ARCMouseDownArcStrokeWidth,
          ARCMouseEnterDisplay : opts.ARCMouseEnterDisplay,
          ARCMouseEnterColor : opts.ARCMouseEnterColor,
          ARCMouseEnterArcOpacity : opts.ARCMouseEnterArcOpacity,
          ARCMouseEnterArcStrokeColor : opts.ARCMouseEnterArcStrokeColor,
          ARCMouseEnterArcStrokeWidth : opts.ARCMouseEnterArcStrokeWidth,
          ARCMouseLeaveDisplay : opts.ARCMouseLeaveDisplay,
          ARCMouseLeaveColor : opts.ARCMouseLeaveColor,
          ARCMouseLeaveArcOpacity : opts.ARCMouseLeaveArcOpacity,
          ARCMouseLeaveArcStrokeColor : opts.ARCMouseLeaveArcStrokeColor,
          ARCMouseLeaveArcStrokeWidth : opts.ARCMouseLeaveArcStrokeWidth,
          ARCMouseMoveDisplay : opts.ARCMouseMoveDisplay,
          ARCMouseMoveColor : opts.ARCMouseMoveColor,
          ARCMouseMoveArcOpacity : opts.ARCMouseMoveArcOpacity,
          ARCMouseMoveArcStrokeColor : opts.ARCMouseMoveArcStrokeColor,
          ARCMouseMoveArcStrokeWidth : opts.ARCMouseMoveArcStrokeWidth,
          ARCMouseOutDisplay : opts.ARCMouseOutDisplay,
          ARCMouseOutAnimationTime : opts.ARCMouseOutAnimationTime,
          ARCMouseOutColor : opts.ARCMouseOutColor,
          ARCMouseOutArcOpacity : opts.ARCMouseOutArcOpacity,
          ARCMouseOutArcStrokeColor : opts.ARCMouseOutArcStrokeColor,
          ARCMouseOutArcStrokeWidth : opts.ARCMouseOutArcStrokeWidth,
          ARCMouseUpDisplay : opts.ARCMouseUpDisplay,
          ARCMouseUpColor : opts.ARCMouseUpColor,
          ARCMouseUpArcOpacity : opts.ARCMouseUpArcOpacity,
          ARCMouseUpArcStrokeColor : opts.ARCMouseUpArcStrokeColor,
          ARCMouseUpArcStrokeWidth : opts.ARCMouseUpArcStrokeWidth,
          ARCMouseOverDisplay : opts.ARCMouseOverDisplay,
          ARCMouseOverColor : opts.ARCMouseOverColor,
          ARCMouseOverArcOpacity : opts.ARCMouseOverArcOpacity,
          ARCMouseOverArcStrokeColor : opts.ARCMouseOverArcStrokeColor,
          ARCMouseOverArcStrokeWidth : opts.ARCMouseOverArcStrokeWidth,

          ARCMouseOverTooltipsHtml01 : opts.ARCMouseOverTooltipsHtml01,
          ARCMouseOverTooltipsHtml02 : opts.ARCMouseOverTooltipsHtml02,
          ARCMouseOverTooltipsHtml03 : opts.ARCMouseOverTooltipsHtml03,
          ARCMouseOverTooltipsHtml04 : opts.ARCMouseOverTooltipsHtml04,
          ARCMouseOverTooltipsHtml05 : opts.ARCMouseOverTooltipsHtml05,
          ARCMouseOverTooltipsBorderWidth : opts.ARCMouseOverTooltipsBorderWidth,

          // CNV options
          CNVMouseEvent : opts.CNVMouseEvent,
          CNVMouseClickDisplay : opts.CNVMouseClickDisplay,
          CNVMouseClickColor : opts.CNVMouseClickColor,
          CNVMouseClickArcOpacity : opts.CNVMouseClickArcOpacity,
          CNVMouseClickArcStrokeColor : opts.CNVMouseClickArcStrokeColor,
          CNVMouseClickArcStrokeWidth : opts.CNVMouseClickArcStrokeWidth,
          CNVMouseClickTextFromData : opts.CNVMouseClickTextFromData,
          CNVMouseClickTextOpacity : opts.CNVMouseClickTextOpacity,
          CNVMouseClickTextColor : opts.CNVMouseClickTextColor,
          CNVMouseClickTextSize : opts.CNVMouseClickTextSize,
          CNVMouseClickTextPostionX : opts.CNVMouseClickTextPostionX,
          CNVMouseClickTextPostionY : opts.CNVMouseClickTextPostionY,
          CNVMouseClickTextDrag : opts.CNVMouseClickTextDrag,
          CNVMouseDownDisplay : opts.CNVMouseDownDisplay,
          CNVMouseDownColor : opts.CNVMouseDownColor,
          CNVMouseDownArcOpacity : opts.CNVMouseDownArcOpacity,
          CNVMouseDownArcStrokeColor : opts.CNVMouseDownArcStrokeColor,
          CNVMouseDownArcStrokeWidth : opts.CNVMouseDownArcStrokeWidth,
          CNVMouseEnterDisplay : opts.CNVMouseEnterDisplay,
          CNVMouseEnterColor : opts.CNVMouseEnterColor,
          CNVMouseEnterArcOpacity : opts.CNVMouseEnterArcOpacity,
          CNVMouseEnterArcStrokeColor : opts.CNVMouseEnterArcStrokeColor,
          CNVMouseEnterArcStrokeWidth : opts.CNVMouseEnterArcStrokeWidth,
          CNVMouseLeaveDisplay : opts.CNVMouseLeaveDisplay,
          CNVMouseLeaveColor : opts.CNVMouseLeaveColor,
          CNVMouseLeaveArcOpacity : opts.CNVMouseLeaveArcOpacity,
          CNVMouseLeaveArcStrokeColor : opts.CNVMouseLeaveArcStrokeColor,
          CNVMouseLeaveArcStrokeWidth : opts.CNVMouseLeaveArcStrokeWidth,
          CNVMouseMoveDisplay : opts.CNVMouseMoveDisplay,
          CNVMouseMoveColor : opts.CNVMouseMoveColor,
          CNVMouseMoveArcOpacity : opts.CNVMouseMoveArcOpacity,
          CNVMouseMoveArcStrokeColor : opts.CNVMouseMoveArcStrokeColor,
          CNVMouseMoveArcStrokeWidth : opts.CNVMouseMoveArcStrokeWidth,
          CNVMouseOutDisplay : opts.CNVMouseOutDisplay,
          CNVMouseOutAnimationTime : opts.CNVMouseOutAnimationTime,
          CNVMouseOutColor : opts.CNVMouseOutColor,
          CNVMouseOutArcOpacity : opts.CNVMouseOutArcOpacity,
          CNVMouseOutArcStrokeColor : opts.CNVMouseOutArcStrokeColor,
          CNVMouseOutArcStrokeWidth : opts.CNVMouseOutArcStrokeWidth,
          CNVMouseUpDisplay : opts.CNVMouseUpDisplay,
          CNVMouseUpColor : opts.CNVMouseUpColor,
          CNVMouseUpArcOpacity : opts.CNVMouseUpArcOpacity,
          CNVMouseUpArcStrokeColor : opts.CNVMouseUpArcStrokeColor,
          CNVMouseUpArcStrokeWidth : opts.CNVMouseUpArcStrokeWidth,
          CNVMouseOverDisplay : opts.CNVMouseOverDisplay,
          CNVMouseOverColor : opts.CNVMouseOverColor,
          CNVMouseOverArcOpacity : opts.CNVMouseOverArcOpacity,
          CNVMouseOverArcStrokeColor : opts.CNVMouseOverArcStrokeColor,
          CNVMouseOverArcStrokeWidth : opts.CNVMouseOverArcStrokeWidth,
          CNVMouseOverTooltipsHtml01 : opts.CNVMouseOverTooltipsHtml01,
          CNVMouseOverTooltipsHtml02 : opts.CNVMouseOverTooltipsHtml02,
          CNVMouseOverTooltipsHtml03 : opts.CNVMouseOverTooltipsHtml03,
          CNVMouseOverTooltipsHtml04 : opts.CNVMouseOverTooltipsHtml04,
          CNVMouseOverTooltipsHtml05 : opts.CNVMouseOverTooltipsHtml05,
          CNVMouseOverTooltipsPosition : opts.CNVMouseOverTooltipsPosition,
          CNVMouseOverTooltipsBackgroundColor : opts.CNVMouseOverTooltipsBackgroundColor,
          CNVMouseOverTooltipsBorderStyle : opts.CNVMouseOverTooltipsBorderStyle,
          CNVMouseOverTooltipsBorderWidth : opts.CNVMouseOverTooltipsBorderWidth,
          CNVMouseOverTooltipsPadding : opts.CNVMouseOverTooltipsPadding,
          CNVMouseOverTooltipsBorderRadius : opts.CNVMouseOverTooltipsBorderRadius,
          CNVMouseOverTooltipsOpacity : opts.CNVMouseOverTooltipsOpacity,

          // Line interaction options
          LINEMouseEvent : opts.LINEMouseEvent,
          LINEMouseClickDisplay : opts.LINEMouseClickDisplay,
          LINEMouseClickLineOpacity : opts.LINEMouseClickLineOpacity,
          LINEMouseClickLineStrokeColor : opts.LINEMouseClickLineStrokeColor,
          LINEMouseClickLineStrokeWidth : opts.LINEMouseClickLineStrokeWidth,
          LINEMouseDownDisplay : opts.LINEMouseDownDisplay,
          LINEMouseDownLineOpacity : opts.LINEMouseDownLineOpacity,
          LINEMouseDownLineStrokeColor : opts.LINEMouseDownLineStrokeColor,
          LINEMouseDownLineStrokeWidth : opts.LINEMouseDownLineStrokeWidth,
          LINEMouseEnterDisplay : opts.LINEMouseEnterDisplay,
          LINEMouseEnterLineOpacity : opts.LINEMouseEnterLineOpacity,
          LINEMouseEnterLineStrokeColor : opts.LINEMouseEnterLineStrokeColor,
          LINEMouseEnterLineStrokeWidth : opts.LINEMouseEnterLineStrokeWidth,
          LINEMouseLeaveDisplay : opts.LINEMouseLeaveDisplay,
          LINEMouseLeaveLineOpacity : opts.LINEMouseLeaveLineOpacity,
          LINEMouseLeaveLineStrokeColor : opts.LINEMouseLeaveLineStrokeColor,
          LINEMouseLeaveLineStrokeWidth : opts.LINEMouseLeaveLineStrokeWidth,
          LINEMouseMoveDisplay : opts.LINEMouseMoveDisplay,
          LINEMouseMoveLineOpacity : opts.LINEMouseMoveLineOpacity,
          LINEMouseMoveLineStrokeColor : opts.LINEMouseMoveLineStrokeColor,
          LINEMouseMoveLineStrokeWidth : opts.LINEMouseMoveLineStrokeWidth,
          LINEMouseOutDisplay : opts.LINEMouseOutDisplay,
          LINEMouseOutAnimationTime : opts.LINEMouseOutAnimationTime,
          LINEMouseOutLineOpacity : opts.LINEMouseOutLineOpacity,
          LINEMouseOutLineStrokeColor : opts.LINEMouseOutLineStrokeColor,
          LINEMouseOutLineStrokeWidth : opts.LINEMouseOutLineStrokeWidth,
          LINEMouseUpDisplay : opts.LINEMouseUpDisplay,
          LINEMouseUpLineOpacity : opts.LINEMouseUpLineOpacity,
          LINEMouseUpLineStrokeColor : opts.LINEMouseUpLineStrokeColor,
          LINEMouseUpLineStrokeWidth : opts.LINEMouseUpLineStrokeWidth,
          LINEMouseOverDisplay : opts.LINEMouseOverDisplay,
          LINEMouseOverLineOpacity : opts.LINEMouseOverLineOpacity,
          LINEMouseOverLineStrokeColor : opts.LINEMouseOverLineStrokeColor,
          LINEMouseOverLineStrokeWidth : opts.LINEMouseOverLineStrokeWidth,
          LINEMouseOverTooltipsHtml01 : opts.LINEMouseOverTooltipsHtml01,
          LINEMouseOverTooltipsPosition : opts.LINEMouseOverTooltipsPosition,
          LINEMouseOverTooltipsBackgroundColor : opts.LINEMouseOverTooltipsBackgroundColor,
          LINEMouseOverTooltipsBorderStyle : opts.LINEMouseOverTooltipsBorderStyle,
          LINEMouseOverTooltipsBorderWidth : opts.LINEMouseOverTooltipsBorderWidth,
          LINEMouseOverTooltipsPadding : opts.LINEMouseOverTooltipsPadding,
          LINEMouseOverTooltipsBorderRadius : opts.LINEMouseOverTooltipsBorderRadius,
          LINEMouseOverTooltipsOpacity : opts.LINEMouseOverTooltipsOpacity,


          // Link interaction options
          LINKMouseEvent : opts.LINKMouseEvent,
          LINKMouseClickDisplay : opts.LINKMouseClickDisplay,
          LINKMouseClickColor : opts.LINKMouseClickColor,
          LINKMouseClickTextFromData : opts.LINKMouseClickTextFromData,
          LINKMouseClickTextOpacity : opts.LINKMouseClickTextOpacity,
          LINKMouseClickTextColor : opts.LINKMouseClickTextColor,
          LINKMouseClickTextSize : opts.LINKMouseClickTextSize,
          LINKMouseClickTextPostionX : opts.LINKMouseClickTextPostionX,
          LINKMouseClickTextPostionY : opts.LINKMouseClickTextPostionY,
          LINKMouseClickTextDrag : opts.LINKMouseClickTextDrag,
          LINKMouseDownDisplay : opts.LINKMouseDownDisplay,
          LINKMouseDownColor : opts.LINKMouseDownColor,
          LINKMouseEnterDisplay : opts.LINKMouseEnterDisplay,
          LINKMouseEnterColor : opts.LINKMouseEnterColor,
          LINKMouseLeaveDisplay : opts.LINKMouseLeaveDisplay,
          LINKMouseLeaveColor : opts.LINKMouseLeaveColor,
          LINKMouseMoveDisplay : opts.LINKMouseMoveDisplay,
          LINKMouseMoveColor : opts.LINKMouseMoveColor,
          LINKMouseOutDisplay : opts.LINKMouseOutDisplay,
          LINKMouseOutAnimationTime : opts.LINKMouseOutAnimationTime,
          LINKMouseOutStrokeColor : opts.LINKMouseOutStrokeColor,
          LINKMouseOutStrokeWidth : opts.LINKMouseOutStrokeWidth,
          LINKMouseUpDisplay : opts.LINKMouseUpDisplay,
          LINKMouseUpColor : opts.LINKMouseUpColor,
          LINKMouseOverOpacity : opts.LINKMouseOverOpacity,
          LINKMouseOverDisplay : opts.LINKMouseOverDisplay,
          LINKMouseOverStrokeColor : opts.LINKMouseOverStrokeColor,
          
          LINKMouseOverTooltipsHtml01 : opts.LINKMouseOverTooltipsHtml01,
          LINKMouseOverTooltipsHtml02 : opts.LINKMouseOverTooltipsHtml02,
          LINKMouseOverTooltipsHtml03 : opts.LINKMouseOverTooltipsHtml03,
          LINKMouseOverTooltipsHtml04 : opts.LINKMouseOverTooltipsHtml04,
          LINKMouseOverTooltipsHtml05 : opts.LINKMouseOverTooltipsHtml05,
          LINKMouseOverTooltipsBorderWidth : opts.ARCMouseOverTooltipsBorderWidth,

          // Histogram interaction options
          HISTOGRAMMouseEvent : opts.HISTOGRAMMouseEvent,
          HISTOGRAMMouseClickDisplay : opts.HISTOGRAMMouseClickDisplay,
          HISTOGRAMMouseClickColor : opts.HISTOGRAMMouseClickColor,
          HISTOGRAMMouseClickOpacity : opts.HISTOGRAMMouseClickOpacity,
          HISTOGRAMMouseClickStrokeColor : opts.HISTOGRAMMouseClickStrokeColor,
          HISTOGRAMMouseClickStrokeWidth : opts.HISTOGRAMMouseClickStrokeWidth,
          HISTOGRAMMouseDownDisplay : opts.HISTOGRAMMouseDownDisplay,
          HISTOGRAMMouseDownColor : opts.HISTOGRAMMouseDownColor,
          HISTOGRAMMouseDownOpacity : opts.HISTOGRAMMouseDownOpacity,
          HISTOGRAMMouseDownStrokeColor : opts.HISTOGRAMMouseDownStrokeColor,
          HISTOGRAMMouseDownStrokeWidth : opts.HISTOGRAMMouseDownStrokeWidth,
          HISTOGRAMMouseEnterDisplay : opts.HISTOGRAMMouseEnterDisplay,
          HISTOGRAMMouseEnterColor : opts.HISTOGRAMMouseEnterColor,
          HISTOGRAMMouseEnterOpacity : opts.HISTOGRAMMouseEnterOpacity,
          HISTOGRAMMouseEnterStrokeColor : opts.HISTOGRAMMouseEnterStrokeColor,
          HISTOGRAMMouseEnterStrokeWidth : opts.HISTOGRAMMouseEnterStrokeWidth,
          HISTOGRAMMouseLeaveDisplay : opts.HISTOGRAMMouseLeaveDisplay,
          HISTOGRAMMouseLeaveColor : opts.HISTOGRAMMouseLeaveColor,
          HISTOGRAMMouseLeaveOpacity : opts.HISTOGRAMMouseLeaveOpacity,
          HISTOGRAMMouseLeaveStrokeColor : opts.HISTOGRAMMouseLeaveStrokeColor,
          HISTOGRAMMouseLeaveStrokeWidth : opts.HISTOGRAMMouseLeaveStrokeWidth,
          HISTOGRAMMouseMoveDisplay : opts.HISTOGRAMMouseMoveDisplay,
          HISTOGRAMMouseMoveColor : opts.HISTOGRAMMouseMoveColor,
          HISTOGRAMMouseMoveOpacity : opts.HISTOGRAMMouseMoveOpacity,
          HISTOGRAMMouseMoveStrokeColor : opts.HISTOGRAMMouseMoveStrokeColor,
          HISTOGRAMMouseMoveStrokeWidth : opts.HISTOGRAMMouseMoveStrokeWidth,
          HISTOGRAMMouseOutDisplay : opts.HISTOGRAMMouseOutDisplay,
          HISTOGRAMMouseOutAnimationTime : opts.HISTOGRAMMouseOutAnimationTime,
          HISTOGRAMMouseOutColor : opts.HISTOGRAMMouseOutColor,
          HISTOGRAMMouseOutOpacity : opts.HISTOGRAMMouseOutOpacity,
          HISTOGRAMMouseOutStrokeColor : opts.HISTOGRAMMouseOutStrokeColor,
          HISTOGRAMMouseOutStrokeWidth : opts.HISTOGRAMMouseOutStrokeWidth,
          HISTOGRAMMouseUpDisplay : opts.HISTOGRAMMouseUpDisplay,
          HISTOGRAMMouseUpColor : opts.HISTOGRAMMouseUpColor,
          HISTOGRAMMouseUpOpacity : opts.HISTOGRAMMouseUpOpacity,
          HISTOGRAMMouseUpStrokeColor : opts.HISTOGRAMMouseUpStrokeColor,
          HISTOGRAMMouseUpStrokeWidth : opts.HISTOGRAMMouseUpStrokeWidth,
          HISTOGRAMMouseOverDisplay : opts.HISTOGRAMMouseOverDisplay,
          HISTOGRAMMouseOverColor : opts.HISTOGRAMMouseOverColor,
          HISTOGRAMMouseOverOpacity : opts.HISTOGRAMMouseOverOpacity,
          HISTOGRAMMouseOverStrokeColor : opts.HISTOGRAMMouseOverStrokeColor,
          HISTOGRAMMouseOverStrokeWidth : opts.HISTOGRAMMouseOverStrokeWidth,
          HISTOGRAMMouseOverTooltipsHtml01 : opts.HISTOGRAMMouseOverTooltipsHtml01,
          HISTOGRAMMouseOverTooltipsHtml02 : opts.HISTOGRAMMouseOverTooltipsHtml02,
          HISTOGRAMMouseOverTooltipsHtml03 : opts.HISTOGRAMMouseOverTooltipsHtml03,
          HISTOGRAMMouseOverTooltipsHtml04 : opts.HISTOGRAMMouseOverTooltipsHtml04,
          HISTOGRAMMouseOverTooltipsHtml05 : opts.HISTOGRAMMouseOverTooltipsHtml05,
          HISTOGRAMMouseOverTooltipsHtml06 : opts.HISTOGRAMMouseOverTooltipsHtml06,
          HISTOGRAMMouseOverTooltipsPosition : opts.HISTOGRAMMouseOverTooltipsPosition,
          HISTOGRAMMouseOverTooltipsBackgroundColor : opts.HISTOGRAMMouseOverTooltipsBackgroundColor,
          HISTOGRAMMouseOverTooltipsBorderStyle : opts.HISTOGRAMMouseOverTooltipsBorderStyle,
          HISTOGRAMMouseOverTooltipsBorderWidth : opts.HISTOGRAMMouseOverTooltipsBorderWidth,
          HISTOGRAMMouseOverTooltipsPadding : opts.HISTOGRAMMouseOverTooltipsPadding,
          HISTOGRAMMouseOverTooltipsBorderRadius : opts.HISTOGRAMMouseOverTooltipsBorderRadius,
          HISTOGRAMMouseOverTooltipsOpacity : opts.HISTOGRAMMouseOverTooltipsOpacity,
          HEATMAPMouseEvent : opts.HEATMAPMouseEvent,
          HEATMAPMouseClickDisplay : opts.HEATMAPMouseClickDisplay,
          HEATMAPMouseClickColor : opts.HEATMAPMouseClickColor,
          HEATMAPMouseClickOpacity : opts.HEATMAPMouseClickOpacity,
          HEATMAPMouseClickStrokeColor : opts.HEATMAPMouseClickStrokeColor,
          HEATMAPMouseClickStrokeWidth : opts.HEATMAPMouseClickStrokeWidth,
          HEATMAPMouseDownDisplay : opts.HEATMAPMouseDownDisplay,
          HEATMAPMouseDownColor : opts.HEATMAPMouseDownColor,
          HEATMAPMouseDownOpacity : opts.HEATMAPMouseDownOpacity,
          HEATMAPMouseDownStrokeColor : opts.HEATMAPMouseDownStrokeColor,
          HEATMAPMouseDownStrokeWidth : opts.HEATMAPMouseDownStrokeWidth,
          HEATMAPMouseEnterDisplay : opts.HEATMAPMouseEnterDisplay,
          HEATMAPMouseEnterColor : opts.HEATMAPMouseEnterColor,
          HEATMAPMouseEnterOpacity : opts.HEATMAPMouseEnterOpacity,
          HEATMAPMouseEnterStrokeColor : opts.HEATMAPMouseEnterStrokeColor,
          HEATMAPMouseEnterStrokeWidth : opts.HEATMAPMouseEnterStrokeWidth,
          HEATMAPMouseLeaveDisplay : opts.HEATMAPMouseLeaveDisplay,
          HEATMAPMouseLeaveColor : opts.HEATMAPMouseLeaveColor,
          HEATMAPMouseLeaveOpacity : opts.HEATMAPMouseLeaveOpacity,
          HEATMAPMouseLeaveStrokeColor : opts.HEATMAPMouseLeaveStrokeColor,
          HEATMAPMouseLeaveStrokeWidth : opts.HEATMAPMouseLeaveStrokeWidth,
          HEATMAPMouseMoveDisplay : opts.HEATMAPMouseMoveDisplay,
          HEATMAPMouseMoveColor : opts.HEATMAPMouseMoveColor,
          HEATMAPMouseMoveOpacity : opts.HEATMAPMouseMoveOpacity,
          HEATMAPMouseMoveStrokeColor : opts.HEATMAPMouseMoveStrokeColor,
          HEATMAPMouseMoveStrokeWidth : opts.HEATMAPMouseMoveStrokeWidth,
          HEATMAPMouseOutDisplay : opts.HEATMAPMouseOutDisplay,
          HEATMAPMouseOutAnimationTime : opts.HEATMAPMouseOutAnimationTime,
          HEATMAPMouseOutColor : opts.HEATMAPMouseOutColor,
          HEATMAPMouseOutOpacity : opts.HEATMAPMouseOutOpacity,
          HEATMAPMouseOutStrokeColor : opts.HEATMAPMouseOutStrokeColor,
          HEATMAPMouseOutStrokeWidth : opts.HEATMAPMouseOutStrokeWidth,
          HEATMAPMouseUpDisplay : opts.HEATMAPMouseUpDisplay,
          HEATMAPMouseUpColor : opts.HEATMAPMouseUpColor,
          HEATMAPMouseUpOpacity : opts.HEATMAPMouseUpOpacity,
          HEATMAPMouseUpStrokeColor : opts.HEATMAPMouseUpStrokeColor,
          HEATMAPMouseUpStrokeWidth : opts.HEATMAPMouseUpStrokeWidth,
          HEATMAPMouseOverDisplay : opts.HEATMAPMouseOverDisplay,
          HEATMAPMouseOverColor : opts.HEATMAPMouseOverColor,
          HEATMAPMouseOverOpacity : opts.HEATMAPMouseOverOpacity,
          HEATMAPMouseOverStrokeColor : opts.HEATMAPMouseOverStrokeColor,
          HEATMAPMouseOverStrokeWidth : opts.HEATMAPMouseOverStrokeWidth,
          HEATMAPMouseOverTooltipsHtml01 : opts.HEATMAPMouseOverTooltipsHtml01,
          HEATMAPMouseOverTooltipsHtml02 : opts.HEATMAPMouseOverTooltipsHtml02,
          HEATMAPMouseOverTooltipsHtml03 : opts.HEATMAPMouseOverTooltipsHtml03,
          HEATMAPMouseOverTooltipsHtml04 : opts.HEATMAPMouseOverTooltipsHtml04,
          HEATMAPMouseOverTooltipsHtml05 : opts.HEATMAPMouseOverTooltipsHtml05,
          HEATMAPMouseOverTooltipsHtml06 : opts.HEATMAPMouseOverTooltipsHtml06,
          HEATMAPMouseOverTooltipsPosition : opts.HEATMAPMouseOverTooltipsPosition,
          HEATMAPMouseOverTooltipsBackgroundColor : opts.HEATMAPMouseOverTooltipsBackgroundColor,
          HEATMAPMouseOverTooltipsBorderStyle : opts.HEATMAPMouseOverTooltipsBorderStyle,
          HEATMAPMouseOverTooltipsBorderWidth : opts.HEATMAPMouseOverTooltipsBorderWidth,
          HEATMAPMouseOverTooltipsPadding : opts.HEATMAPMouseOverTooltipsPadding,
          HEATMAPMouseOverTooltipsBorderRadius : opts.HEATMAPMouseOverTooltipsBorderRadius,
          HEATMAPMouseOverTooltipsOpacity : opts.HEATMAPMouseOverTooltipsOpacity,
          
          // Genome options
          genomeFillColor: opts.genomeFillColor,
          genomeBorder : {
            display : opts.displayGenomeBorder,
            borderColor : opts.genomeBorderColor,
            borderSize : opts.genomeBorderSize
          },
          ticks : {
            display : opts.genomeTicksDisplay,
            len : opts.genomeTicksLen,
            color : opts.genomeTicksColor,
            textSize : opts.genomeTicksTextSize,
            textColor : opts.genomeTicksTextColor,
            scale : opts.genomeTicksScale
          },
          genomeLabel : {
            display : opts.genomeLabelDisplay,
            textSize : opts.genomeLabelTextSize,
            textColor : opts.genomeLabelTextColor,
            dx : opts.genomeLabelDx,
            dy : opts.genomeLabelDy,
            rotation : opts.genomeLabelOrientation
          },
          TEXTModuleDragEvent : opts.TEXTModuleDragEvent
        });

        BioCircos01.draw_genome(BioCircos01.genomeLength);
      },

      resize: function(width, height) {

        // code to re-render the widget with a new size

      }

    };
  }
});
```
